# A ribosome-dependent small-ORF switch controls flagellar gene expression and virulence

**DOI:** 10.64898/2026.11.10.765585

**Authors:** Filip Lastovka, Owain J. Bryant, Dominik Meggers, Elizaveta Alexeeva, Melina Knorr, Franz Narberhaus, Betty Y.-W. Chung

## Abstract

RNA structures are active regulators of bacterial translation. Here, we identified a naturally occurring translation-enhancing small open reading frame (tesORF) that controls the expression of the flagellar operon *fliFGHIJK* in *Salmonella enterica* and other motile Enterobacterales. Nucleotide-resolution ribosome profiling revealed this previously unannotated 54-nucleotide ORF immediately upstream of *fliF*. Genetic, transcriptomic, and biochemical analyses support a model in which tesORF translation counteracts an inhibitory RNA structure, exposing the *fliF* ribosome-binding site and promoting translation initiation independently of the encoded peptide sequence. Disruption of tesORF translation reduced expression of the downstream operon and impaired motility and virulence-associated phenotypes in *Escherichia coli*, *Salmonella*, and *Yersinia*. Together, our findings reveal how translation of a conserved small ORF regulates an entire flagellar operon through RNA structural remodelling, with consequences expending to the wider flagellar regulon, linking RNA structural dynamics to bacterial motility and virulence.

**Significance statement:** Bacteria organize functionally related genes into operons, allowing them to be transcribed together. Here, we show how translation of a previously unrecognized small open reading frame controls expression of the flagellar *fliFGHIJK* operon in *E. coli*, *Salmonella*, and *Yersinia*. Translation of this small ORF counteracts an inhibitory RNA structure, enabling translation of the first gene of the operon. Disrupting this mechanism collapses downstream gene expression, compromises motility, and alters virulence-associated phenotypes in *Salmonella.* Our findings reveal how a short coding sequence can regulate an entire operon through the process of translation rather than through its encoded peptide, expanding our understanding of how bacterial genomes encode regulatory information and coordinate production of functionally linked proteins.

## Introduction

Many bacterial genes are organized in operons: groups of neighbouring coding sequences transcribed together from a shared promoter into a single polycistronic mRNA. Translation of individual genes within an operon is modulated by RNA features, including mRNA secondary structure and Shine–Dalgarno (SD) strength, as well as by translation of neighbouring open reading frames (ORFs).

In addition to transcriptional coupling within operons, translational coupling has been observed in artificial constructs, native bacterial operons, and in bacteriophages. Translational coupling occurs when translation of an upstream ORF promotes translation of a downstream ORF on the same mRNA, as exemplified by *trpB*–*trpA* and *trpE–trpD* gene pairs in the *Escherichia coli* tryptophan biosynthesis operon [1] [2]. The mechanism of translational coupling can involve disruption of an inhibitory mRNA secondary structure by translating ribosomes, thereby exposing the SD sequence of the downstream gene and facilitating translation initiation [3] [4] [5] [6] [7] [8] [9] [10] [11]. Translational coupling can also occur independently of inhibitory mRNA secondary structures through 70S-scanning. In this mechanism, the ribosome remains associated with the mRNA after terminating translation of an upstream ORF and locates a downstream start codon by one-dimensional diffusion, enabling translation reinitiation [12] [13] [14] [15] [16] [17]. This mechanism parallels the scanning and reinitiation model for mammalian *ATF4* mRNA, but involves an intact 70S ribosome rather than a retained 40S subunit [18]. Translational coupling is most efficient when the distance between the termination and initiation sites is short [4] [13] [17] [19] [20] [21] [22] [23]. Although 70S-scanning initiation and *de novo* translation initiation are not mutually exclusive, reinitiation is likely favoured at intergenic distances below 30 nucleotides, where spatial constraints may limit simultaneous termination at the upstream ORF and initiation at the downstream ORF [15]. A strong downstream SD sequence can enhance coupling [24], although coupling can also promote translation of ORFs with weak SD sequences [14] [25].

Several naturally occurring bacterial small ORFs enhance translation of downstream genes through translational coupling [26] [27] [28]. In this study, using nucleotide-resolution ribosome profiling, we identified a highly conserved small ORF in *Salmonella enterica* serovar Typhimurium and *Escherichia coli*, which we termed a translation-enhancing small open reading frame (tesORF). We show that efficient ribosome recruitment to the tesORF SD sequence enables tesORF translation, which disrupts the inhibitory hairpin sequestering the *fliF* SD sequence and thereby promotes *fliF* translation. Loss of tesORF-mediated regulation results in loss of flagella, lack of motility, and impaired virulence.

## Results

### Nucleotide-resolution Ribo-Seq reveals a conserved open reading frame upstream of a flagellar operon in Enterobacterales

Analysis of ribosome profiles from *Salmonella* Typhimurium SL1344 and *E. coli* MG1655 (Ribo-Seq; Bryant *et al.*, 2023 [29] and this study; figures S1A–C) revealed an accumulation of highly phased ribosome-protected fragment (RPF) reads mapping to the leader region of the flagellar operon *fliFGHIJK* (figures 1A–B, S1D, and S1E). These reads define a 54-nt ORF, encoding 17 amino acids, that is out of frame with the *fliF* coding sequence (CDS). The small ORF begins with a non-standard UUG start codon preceded by a clear SD sequence, and its stop codon overlaps the *fliF* start codon in the sequence AUGA. DNA sequence alignments revealed that the small ORF is conserved upstream of *fliF* in motile Enterobacterales. Despite variation in the coding sequence, particularly in the more distantly related genera *Yersinia* or *Serratia*, all examined species retained a UUG codon preceded by a putative SD sequence and an uninterrupted ORF whose stop codon overlaps the *fliF* start (figures 1C, and S2). This conservation suggests a role for the small ORF in regulating expression of the downstream operon.

**Figure 1.**
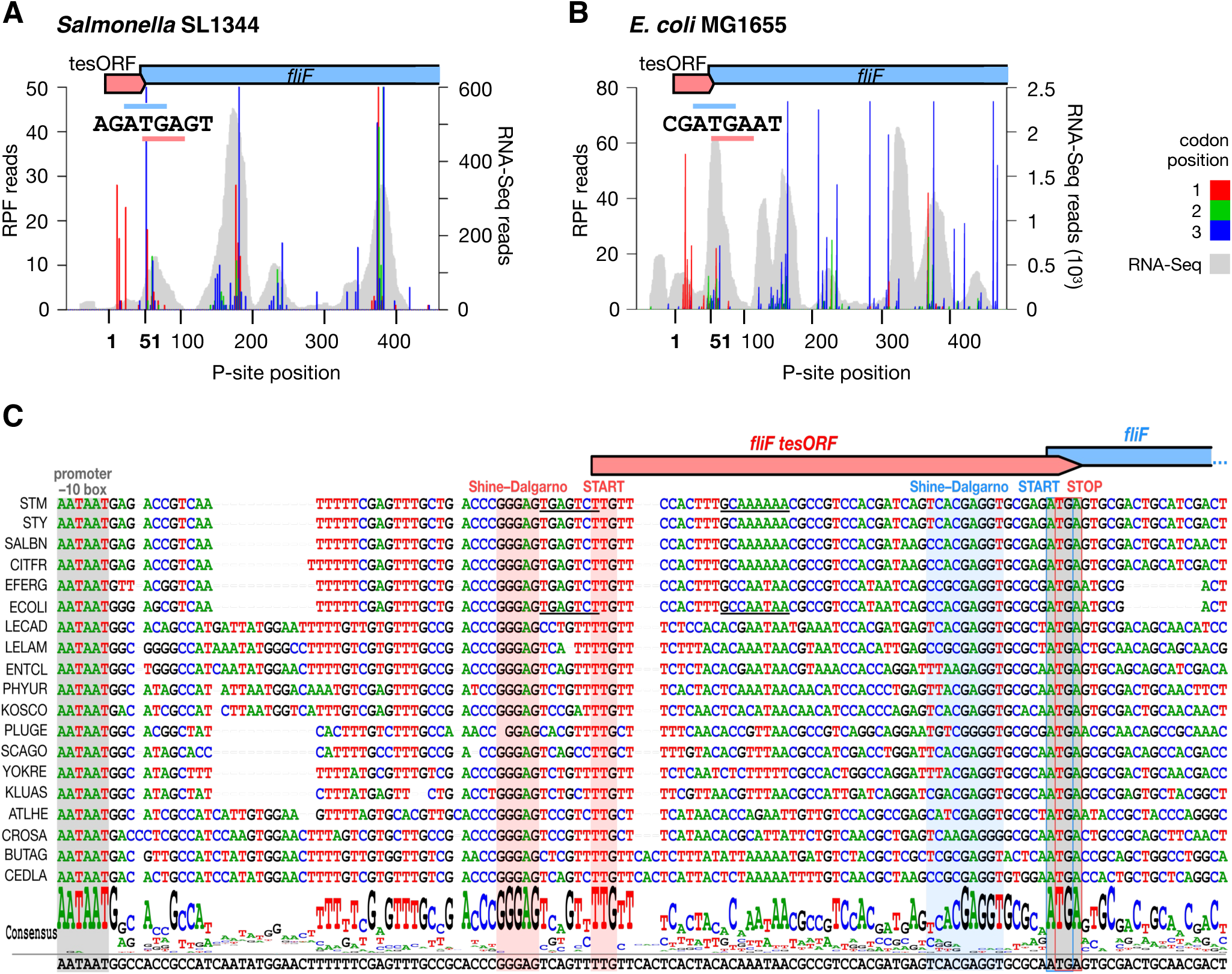
**A** and **B**, Visualization of translation of the novel *fliF* translation-enhancing small ORF (*fliF*-tesORF) in *Salmonella* SL1344 (**A**; RPF reads capped at 50) and *E. coli* MG1655 (**B**; RPF reads capped at 75). Red, green, and blue bars indicate Ribo-Seq reads mapping to codon positions 1, 2, and 3, respectively, relative to the first nucleotide of the tesORF. The positions correspond to the inferred P-site position. Grey shaded areas show the coverage of RNA-Seq reads. The overlap of the tesORF stop codon with the start codon of *fliF* is shown in the map. **C**, Alignment of the region upstream of *fliF* shows conservation of the tesORF among Enterobacterales. A potential σ^28^ promoter sequence is underlined in *Salmonella* SL1344 and *E. coli*. Species abbreviations are as follows: *Salmonella enterica* serovar Typhimurium (STM), *Salmonella enterica* serovar Typhi (STY), *Salmonella bongori* (SALBN), *Citrobacter freundii* (CITFR), *Escherichia fergusonii* (EFEG), *Escherichia coli* (ECOLI), *Leclercia adecarboxylata* (LECAD), *Lelliottia amnigena* (LELAM), *Enterobacter cloacae* (ENTCL), *Phytobacter ursingii* (PHYUR), *Kosakonia cowanii* (KOSCO), *Pluralibacter gergoviae* (PLUGE), *Scandinavium goeteborgense* (SCAGO), *Yokenella regensburgei* (YOKRE), *Kluyvera ascorbata* (KLUAS), *Atlantibacter hermanii* (ATLHE), *Cronobacter sakazakii* (CROSA), *Buttiauxella agrestis* (BUTAG) and *Cedecea lapagei* (CEDLA).

### Short upstream open reading frame regulates downstream gene translation

The strong conservation of the small ORF suggests a role in regulating expression of the *fliFGHIJK* flagellar operon. To test its effect on *fliF* translation, we generated an anhydrotetracycline-inducible luciferase reporter construct [29]. The reporter contains the *Salmonella fliFGHIJK* 5′ UTR, the small ORF, and the first ten codons of *fliF* fused in-frame to *luxA,* the first gene of the *luxAB* luciferase operon. The 5′-UTR boundary was defined using the transcription start site reported in the *Salmonella* compendium [30], which agrees with our RNA-Seq data [29]. This design provides a sensitive, luminescence-based readout of *fliF* expression.

Luminescence was assayed at intervals of 5 min for 20 min after transcription induction, comparing reporters containing either the intact small ORF or a disrupted start codon. In the mutant, the UUG start codon was replaced with UAAG, introducing a UAA stop codon and an additional nucleotide that shifts the reading frame. Disruption of the small-ORF start codon markedly reduced luminescence, supporting a role for small ORF translation in promoting *fliF* translation (figure 2A and S3A). We therefore named this element the **translation-enhancing small ORF** (tesORF).

**Figure 2.**
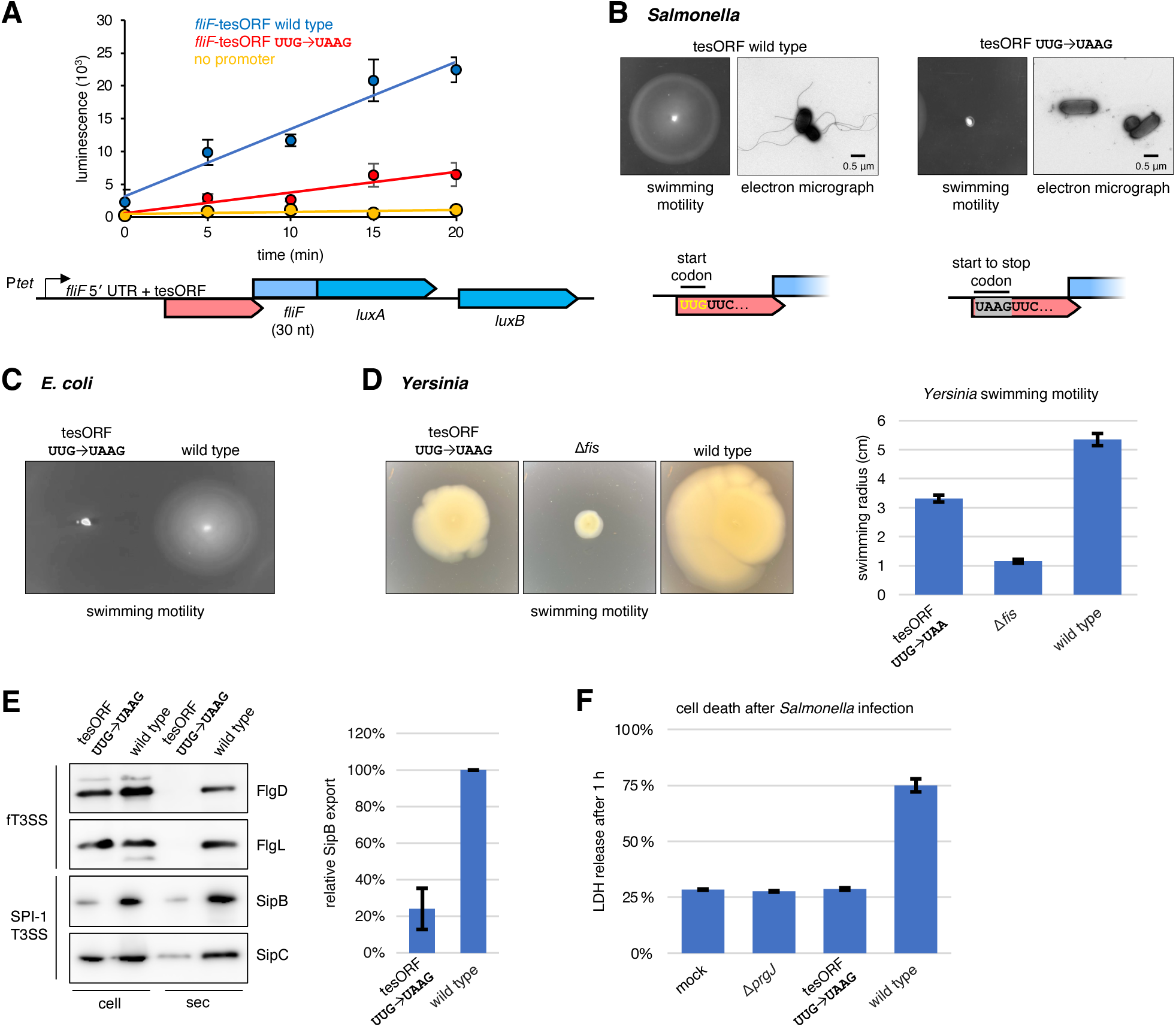
**A**, Induction time course of a *luxAB* luciferase reporter containing the *Salmonella fliF* 5′ UTR, tesORF (or its UUG→UAAG start codon mutant), and the first ten codons of the *fliF* CDS fused in-frame to *luxA.* Expression in *Salmonella* was measured by luminescence. A schematic of the construct is shown below. **B**, Swimming motility assay and negative-stain electron micrograph of wild-type and *fliF-*tesORF mutant *Salmonella*. The wild-type and mutant tesORF start codon sequences are shown below. **C**, Swimming motility assay of *fliF-*tesORF mutant (UUG→UAAG) and wild-type *E. coli* MG1655. **D**, Swimming motility assay of *fliF-*tesORF mutant (UUG→UAAG), wild-type, and non-motile mutant (Δ*fis*) *Yersinia pseudotuberculosis* YPIII. Results are from four independent replicates. **E**, Export assay of flagellar (fT3SS) and *Salmonella* pathogenicity island 1 injectisome (SPI-1 T3SS) proteins by wild-type and *fliF-*tesORF UUG→UAAG mutant *Salmonella*. Whole-cell fractions (cell) and secreted fractions (sec) were examined by Western blotting. SipB export was quantified from the secreted fraction using ImageJ and normalized to wild type (100%). Error bars represent the standard error of the mean from three independent experiments. **F**, Cell death assay in iBMDMs 1 h after infection by wild-type *Salmonella*, the *fliF-*tesORF UUG→UAAG mutant, or a SPI-1 injectisome mutant (Δ*prgJ*) at a multiplicity of infection (MOI) of 10. No bacteria were added to the mock group.

Reverse-transcription quantitative PCR showed similar RNA abundance between the constructs following transcription induction (figure S3B), consistent with the previous findings using this reporter system [29]. Furthermore, the UUG→UAAG mutation reduced the abundance of a tagged reporter protein below the detection limit of immunoblotting (figure S3C).

Swimming motility assays were employed to assess the function of *fliF*-tesORF. Chromosomal mutations disrupting the tesORF start codon completely abolished swimming in both *Salmonella* and *E. coli* (figures 2B and C). *Yersinia* possesses two flagellar clusters: one containing a tesORF-associated *fliFGHIJK* operon, which is expressed at a higher level at 25 °C than at 37 °C, and another containing a *fliEFGHIJ* operon, with approximately 50-fold lower transcript levels (figure S4A [31] [32]). Disruption of the *fliF*-tesORF in *Yersinia pseudotuberculosis* significantly reduced swimming at room temperature (figure 2D), despite this tesORF being longer and more divergent than its *Salmonella* counterpart (figure S2A). Together, these results establish a critical role for *fliF*-tesORF in motility.

Protein export assays further showed that the *Salmonella fliF-*tesORF start-codon mutant (UUG→UAAG) was defective in flagellar type III secretion (fT3SS), consistent with its lack of flagella (figures 2B and E). This strain also displayed reduced secretion of SipB and SipC, proteins exported by the *Salmonella* pathogenicity island 1 injectisome type III secretion system (SPI-1 T3SS; figures 2E and S4B). Changes in flagellar gene expression have previously been linked to altered SPI-1 T3SS activity [33] [34]. Our findings therefore suggest that tesORF-mediated regulation of the *fliFGHIJK* operon may affect secretion pathways beyond the flagellar system. The tesORF mutant also induced less host cell death than the wild type during infection of murine immortalized bone-marrow-derived macrophages (iBMDMs; figure 2F), potentially reflecting reduced bacterial contact with host cells owing to impaired motility.

Given that bacterial peptides as short as seven amino acid residues, such as cCF10, can be functional [35], we considered whether tesORF acts through its encoded peptide or through the process of translation itself. The amino acid sequence encoded by the *fliF-*tesORF shows a degree of conservation across species (figure S2B), suggesting the possibility of a functional peptide product. However, the plasmid-based expression of *fliF-*tesORF *in trans* did not restore motility in the tesORF mutant, arguing against a freely diffusible peptide mediating this phenotype (figure S4C). Likewise, ectopic expression of *fliF-*tesORF in wild-type *Salmonella* did not impair swimming (figure S4D), despite being detectable in whole cell fractions (figure S4E). Reporter experiments described below further indicate that downstream translation does not depend on the tesORF-encoded amino acid sequence. Together, these findings support a model in which tesORF translation, rather than its peptide product, regulates expression of the flagellar operon.

Transcription of the *fliFGHIJK* operon is driven by a σ^70^-dependent promoter controlled by FlhD4C2 [36] [37] [38]. *E. coli* and *Salmonella* also contain a sequence overlapping the *fliF-*tesORF CDS that resembles a promoter recognized by σ^28^, the sigma factor responsible for expression of late flagellar genes [39] [40] (figures 1C and S2A, underlined). However, the tesORF mutant reporter retained lower signal than the wild-type reporter in a σ^28^-deficient (Δ*fliA*) strain, indicating that the expression defect does not depend on σ^28^ and supporting a role for tesORF translation (figure S4F). This interpretation is consistent with previous regulon mapping, which found no evidence of σ^28^-dependent regulation of the *fliFGHIJK* operon, and with the poor conservation of the putative σ^28^ promoter across Enterobacterales (figures 1C and S2A) [41].

### Translation of *fliF*-tesORF is required for full flagellar gene expression

Ribo-Seq was used to assess the impact of *fliF-*tesORF translation on gene expression in *Salmonella* (figures S5A and B). As expected, the start codon mutation abolished *fliF*-tesORF translation (figure 3A and S5C). Translation was markedly reduced throughout the entire *fliFGHIJK* operon, which encodes components of the flagellar basal body and ATPase complex (figures 3B and S5D). RNA-Seq revealed a corresponding decrease in transcript abundance, with reduced coverage from within *fliF* onwards. This pattern may reflect mRNA destabilization from loss of translating ribosomes and consequent exposure of RNA cleavage sites [42]. Alternatively, exposure of cryptic Rho utilization sites could promote premature transcriptional termination [43] [44] [45].

**Figure 3.**
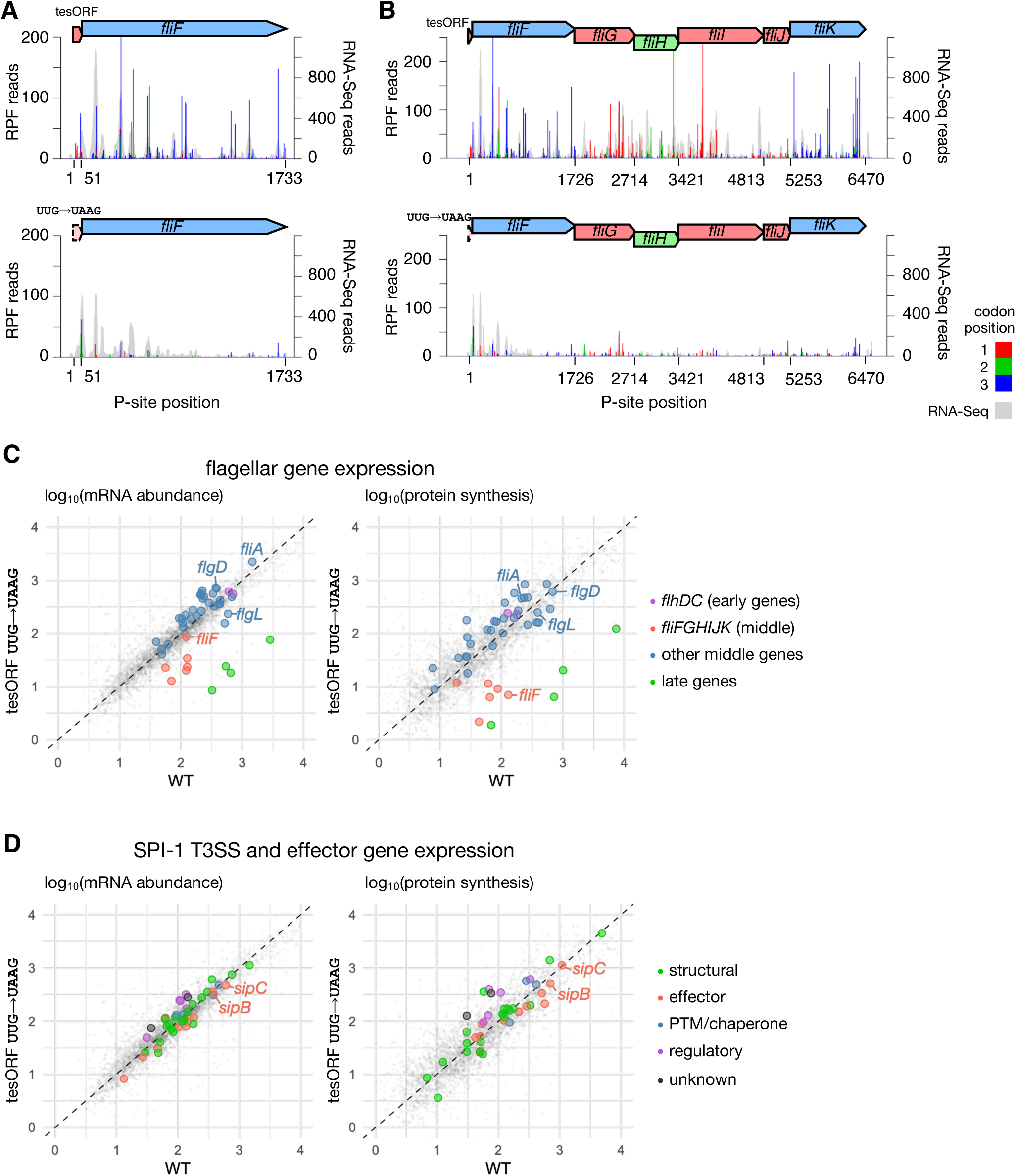
**A** and **B**, Ribo-Seq and parallel RNA-Seq read maps for *fliF*-tesORF and *fliF* (**A**) and the *fliFGHIJK* operon (**B**; Ribo-Seq display capped at 250 reads) in wild-type *Salmonella* and the *fliF*-tesORF UUG→UAAG mutant, with read counts normalized by library size. Red, green, and blue bars indicate Ribo-Seq reads mapping to codon positions 1, 2, and 3, respectively, relative to the first nucleotide of the tesORF. Read positions correspond to the inferred P-site position. Grey shaded areas show the coverage of RNA-Seq reads. **C**, Comparison of RNA abundance and translation of flagellar genes in wild-type *Salmonella* and the tesORF UUG→UAAG mutant. Flagellar genes are coloured according to their regulatory class: early genes (violet), the *fliFGHIJK* operon (red), other middle genes (blue), and late genes (regulated by σ^28^; green). Genes with both a class 2 (middle) and a class 3 (late) promoter are shown in blue. Non-flagellar genes are shown in grey. The dashed line indicates equal mRNA abundance or protein synthesis between the two strains (y = x). **D**, Comparison of RNA abundance and translation of *Salmonella* pathogenicity island 1 type III secretion system (SPI-1 T3SS) genes and secreted effector genes in wild-type *Salmonella* and the tesORF UUG→UAAG mutant. The SPI-1 T3SS and effector genes are displayed in colour corresponding to injectisome structural genes (green), effector genes (including those outside of SPI-1, red), post-translational modification (PTM) and chaperone genes (blue), and regulatory genes (violet). Genes outside of SPI-1 that do not encode secreted effectors are shown in grey. The dashed line indicates equal mRNA abundance or protein synthesis between the two strains (y = x).

Late flagellar genes, including genes encoding the filament and motor complex components under σ^28^ control, also showed a marked reduction in mRNA abundance and protein synthesis output, approximated from the total number of ribosome-protected fragments recovered, in the *fliF*-tesORF UUG→UAAG mutant (figure 3C; table S1). This broader effect is consistent with the hierarchical organization of flagellar gene regulation, in which the expression of late genes depends on successful completion of earlier assembly steps (reviewed by Minamino and Kinoshita, 2023 [46]). Although σ^28^ production was not reduced, its activity depends on release from inhibition by the anti-sigma factor FlgM during flagellar assembly. Reduced FliF synthesis in the *fliF*-tesORF mutant may therefore impair σ^28^ activation and consequently reduce transcription of late flagellar genes.

Ribo-Seq and RNA-Seq showed comparable transcript abundance and protein synthesis output for the SPI-1 effector genes *sipC* and *sipB* in the wild type and the tesORF UUG→UAAG mutant (figure 3D; table S1). Nevertheless, secretion of both effectors was markedly impaired (figure 2E and S4B), indicating that loss of *fliF*-tesORF translation affects SPI-1 secretion without reducing effector expression. These findings are consistent with coordination between flagellar function and virulence-associated type III secretion [47] [48].

### An RNA secondary structure overlapping tesORF limits access to the *fliF* Shine–Dalgarno sequence

RNA secondary structure predictions in *Salmonella* and *E. coli* revealed that the SD sequence and the start codon of *fliF* are embedded within an RNA stem–loop (figures S6A and B). Translation of tesORF likely disrupts the stem–loop, enabling ribosome access to the *fliF* translation initiation site. *Yersinia fliF* also possesses a predicted RNA secondary structure more stable than that at the average translation initiation site (figure S6C). Enzymatic probing of the *Salmonella fliFGHIJK* leader sequence at three different temperatures with RNases T1 and T2, which cleave unpaired bases with different sequence preferences, confirmed that the *fliF* SD sequence flanking the exposed GAG loop was largely protected (figures 4A, 4B, and S6D). The minimum free energy (MFE) of the *Salmonella fliF* stem–loop determined from the probing data was −11.8 kcal mol^−1^ (figure 4C), indicating greater stability than the average RNA structure around translation initiation sites in *Salmonella* (figure S6A). A stem–loop structure with an MFE of −13.0 kcal mol^−1^ is predicted to form in the same location in *E. coli* (figure S7A). In contrast, the tesORF SD sequence and start codon were exposed, allowing tesORF translation, which in turn likely relieves the structural inhibition of *fliF* translation and activates downstream gene expression. Structure predictions supported by transcriptome-wide RNA structure probing data [49] suggest that the divergent *Yersinia fliF* mRNA sequence may form inhibitory long-distance interactions (figures S2A and S7B).

**Figure 4.**
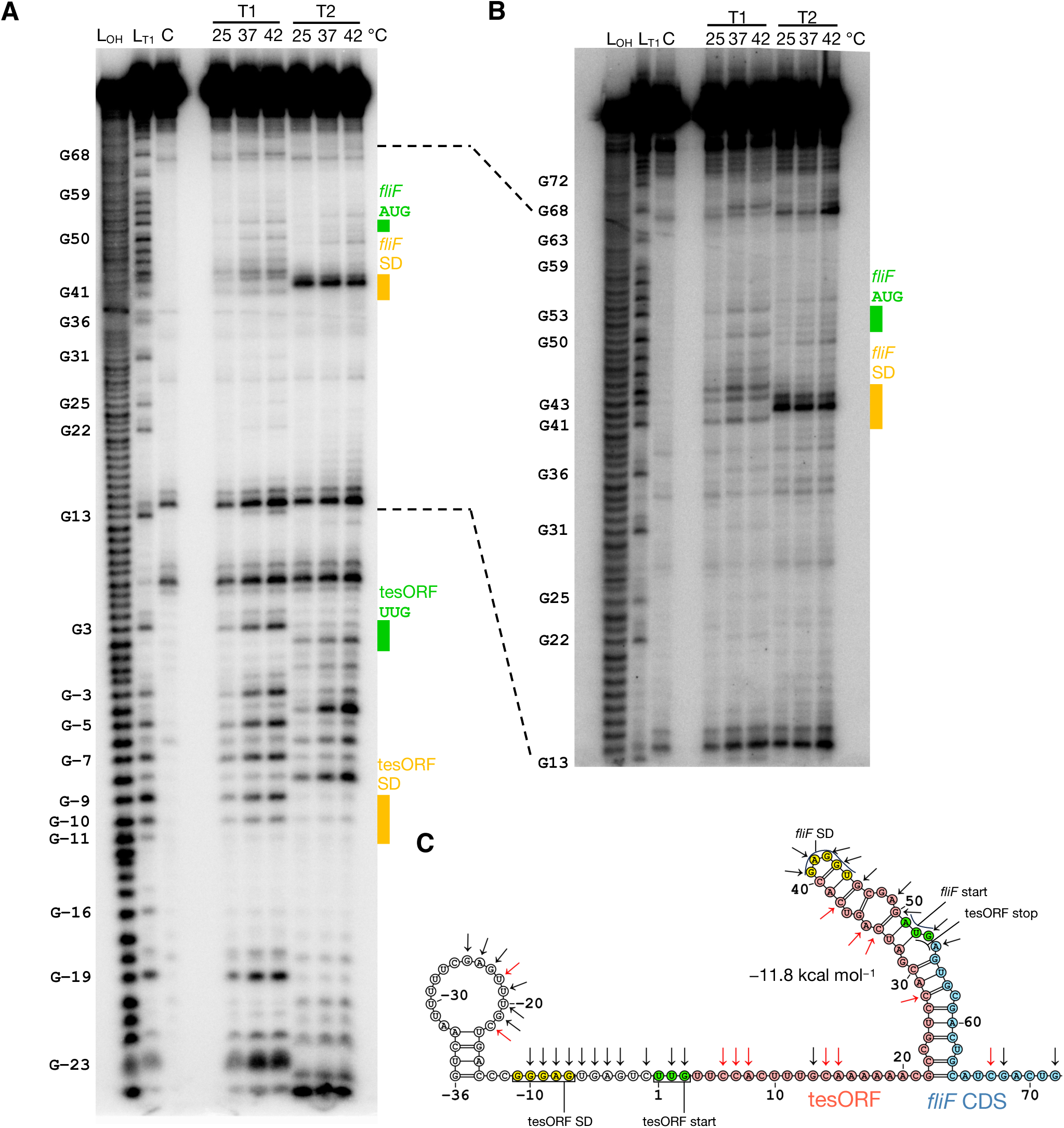
**A** and **B**, 8% polyacrylamide gel showing partially digested 5′-end-labelled RNA containing the *Salmonella fliF* 5′ UTR, tesORF, and the first 30 nucleotides of the *fliF* coding sequence. Products from the same probing reaction were resolved for longer in panel B. Limited digestion was performed at 25 °C, 37 °C, or 42 °C with 0.0016 U of RNase T1, which cleaves RNA 3′ of unpaired guanine, or 0.053 U of RNase T2, which cleaves RNA 3′ of unpaired bases, preferentially adenine. Nucleotide positions are numbered relative to the first nucleotide of the tesORF. An alkaline hydrolysis ladder and a T1 digestion ladder generated from the same RNA are shown in lanes L_OH_ and L_T1_, respectively. The lane labelled C contains the enzyme-free control group. **C**, The experimentally determined secondary structure of *Salmonella fliF* RNA. The Shine–Dalgarno sequences are highlighted in yellow and start codons in green. Cleavage sites detected by structure-probing RNases are indicated by black arrows. Red arrows indicate signals also present in the enzyme-free control.

### Destabilization of the RNA secondary structure rescues main ORF expression

To identify mechanisms capable of restoring motility when *fliF-*tesORF translation is abolished, motile suppressors were isolated from the non-motile *Salmonella fliF-*tesORF start codon mutant. Four motile suppressors were recovered. Each carried a single-nucleotide substitution within the tesORF or *fliF* coding region. Motility was partially restored, although it remained below wild-type levels (figure 5A), and luciferase reporters showed correspondingly intermediate activity (figure 5B). All substitutions mapped within the experimentally determined RNA stem–loop, and the predicted partial destabilization of this structure correlated with recovery of translation. Apart from suppressor A, all suppressor mutants exhibited higher predicted *fliF* ribosome-binding site strength than the parental sequence, consistent with enhanced initiation (table S2 [50]). The reduced swimming of suppressor D (*fliF*-tesORF UUG→UAAG; *fliF* 13G>A), despite its stronger ribosome-binding site and higher reporter activity, could not be explained solely by the alanine-to-threonine substitution at position 5 of FliF, since a strain carrying the same *fliF* allele but an intact *fliF*-tesORF displayed wild-type motility (figure S4G).

**Figure 5.**
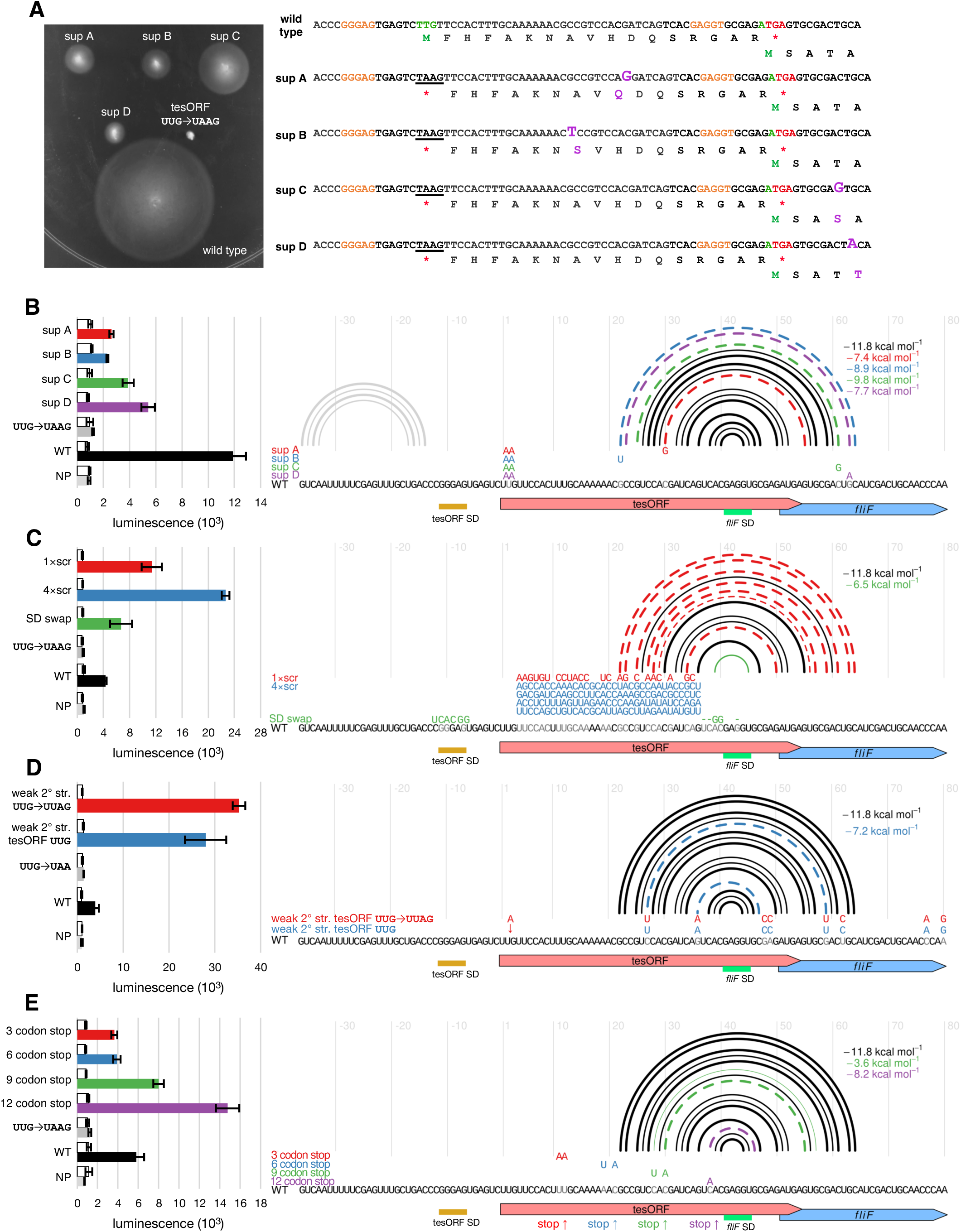
**A**, Swimming motility assays of wild-type *Salmonella* (WT), the tesORF mutant (UUG→UAAG), and the suppressor mutants. The suppressor mutant sequences are shown on the right, with Shine–Dalgarno sequences annotated in orange, suppressor mutations in purple, start codons in green, and stop codons in red. The encoded amino acid sequence is given below each nucleotide sequence. **B–E**, Luciferase reporter signal from constructs containing *Salmonella fliF*-tesORF sequence variants (left), with the variants and their effects on RNA secondary structure illustrated on the right. White and coloured bars represent the signal at 0 and 20 min after induction, respectively. The colours correspond to the sequences shown in the chart on the right. Error bars represent the standard error of the mean from three replicates. In the secondary structure diagrams, arcs illustrate GC (thick), AU (medium), and GU (thin) base pairs. Disrupted base pairs are shown as dashed arcs in the colour corresponding to the sequence variant. Sequence differences are highlighted in colour, and the substituted wild-type nucleotides are shown in grey. A downward arrow represents an insertion, while dashes represent deletions. The free energy of the stem– loop structure was calculated with *RNAeval* from the ViennaRNA package [93]. WT = wild-type *fliF* and tesORF sequence in the luciferase reporter construct. NP = luciferase reporter construct with no promoter. **B**, Suppressor mutations that partially restored the motility of the *fliF*-tesORF mutant. **C**, Scrambled sequence variants, in which a 32-nt region of the tesORF coding sequence—excluding the start codon and preceding the *fliF* Shine–Dalgarno sequence—was scrambled, preserving nucleotide composition and avoiding in-frame stop codons. In the 4× scrambled variant, the region was replaced with a fourfold longer sequence with nucleotide ratios corresponding to the original sequence. In the SD swap variant, the Shine–Dalgarno sequences of *fliF* and the tesORF were swapped. **D**, Weakly structured variants of the tesORF coding sequence with either a wild-type (WT) or a mutated *fliF*-tesORF start codon (UUG→UAA). The variants contain eight point mutations that disrupt the experimentally determined RNA stem–loop structure while preserving the amino acid sequence encoded by the tesORF and preventing the formation of alternative strong secondary structures. **E**, Stop codon position variants, in which stop codons were introduced by sequence substitution, yielding tesORF sequences three, six, nine, and twelve codons in length.

To further test the role of RNA structure in controlling downstream translation, luciferase reporters were constructed with scrambled or extended scrambled *Salmonella* tesORF sequences that disrupted the stem–loop while preserving the *fliF* SD sequence. In the extended construct, the scrambled region was increased to four times its original length while preserving nucleotide composition. Both constructs exhibited higher activity than the wild type, consistent with enhanced main ORF translation following destabilization of the secondary structure (figure 5C). These results also further suggest that neither the tesORF-encoded peptide sequence nor the short length of tesORF are required for enhanced downstream translation.

To confirm that the increase in main ORF expression resulted from RNA structural changes rather than changes to the encoded amino acid sequence, additional constructs containing targeted synonymous point mutations were produced that weakened or disrupted the experimentally determined stem–loop (figures 5D and S8A). These mutations produced similar derepression whether the *fliF*-tesORF start codon was intact or disrupted, whereas constructs predicted to form alternative strong secondary structures overlapping the *fliF* SD sequence restored repression (figure S8B).

Reporter assays using a heterologous *tetA* 5′ UTR in place of the native *fliF* leader upstream of the tesORF produced similar results, indicating that specific native upstream elements are not required (figure S8C). In these constructs, shortening the spacer between the *fliF* SD sequence and the *fliF* start codon reduced reporter activity. This was expected, given that most SD sequences in *Salmonella* and *E. coli* are located 4–8 nt upstream of the start codon (counting from the 3′ end of uaaggaggU, inclusive), and spacer lengths outside of this range tend to reduce translation initiation [29] [51] [52] [53]. Conversely, increasing the spacer length enhanced reporter expression despite a lower predicted ribosome-binding site strength. The accompanying decrease in predicted RNA stem–loop stability (MFE changing from −11.8 to −8.5 kcal mol^−1^) suggests that structural accessibility, rather than SD strength alone, is a key determinant of translational efficiency.

Collectively, these findings support a model in which tesORF translation promotes downstream gene expression by destabilizing an inhibitory RNA secondary structure, without requiring a specific encoded peptide sequence or native upstream elements.

### Position of the tesORF stop codon influences downstream translation

To assess how tesORF termination position affects *fliF* expression, we introduced in-frame stop codons at different positions within the tesORF. Reporter activity generally decreased as the tesORF was shortened, with both the native *fliF* and heterologous *tetA-*derived 5′ UTRs (figures 5E and S8D).

Three non-mutually exclusive mechanisms may contribute to this relationship. Firstly, ribosomes terminating closer to the main ORF directly disrupt the inhibitory stem–loop, exposing the *fliF* SD sequence and facilitating initiation. Secondly, translational coupling efficiency generally decreases as the gap between adjacent ORFs increases [3] [12] [13] [19] [20] [21] [22] [23] and is strongest when termination occurs immediately upstream (UGAUG, UAAUG [12]) or downstream of the initiation site (AUGA, as in tesORF [12] [20] [23] [13]), allowing efficient reinitiation. Thirdly, sequence substitutions can alter RNA secondary structure and SD–anti-SD pairing. A stop codon inserted immediately upstream of the *fliF* SD sequence (codon 12 stop, figure 5E) produced substantially higher reporter activity than the wild-type reporter, consistent with improved base-pairing compatibility with the 16S rRNA (UcAcGAGGU to UAAcGAGGU).

Additional experiments examined the effects of other translation control elements. Swapping the SD sequences of tesORF and *fliF* also disrupted the stem–loop and strengthened the predicted tesORF ribosome-binding site (table S2), resulting in increased main ORF expression (figure 5C). Unexpectedly, replacement of the native tesORF UUG start codon with AUG reduced reporter expression, despite the higher predicted ribosome-binding site strength of the UUG→AUG sequence (figure 6A and table S2). Substitution of the *fliF* AUG start codon with the non-canonical GUG reduced the reporter signal, as expected (figure S8C). In contrast, mutation of the tesORF stop codon (UGA→UGG) extended tesORF translation by 27 codons but yielded reporter activity comparable to the wild type, despite the limited opportunity for reinitiation at *fliF*. Maintenance of reporter activity when tesORF translation extends beyond the *fliF* start codon supports a model in which ribosome transit through the inhibitory stem–loop permits *de novo* initiation at *fliF*, without requiring reinitiation by the same ribosome.

**Figure 6.**
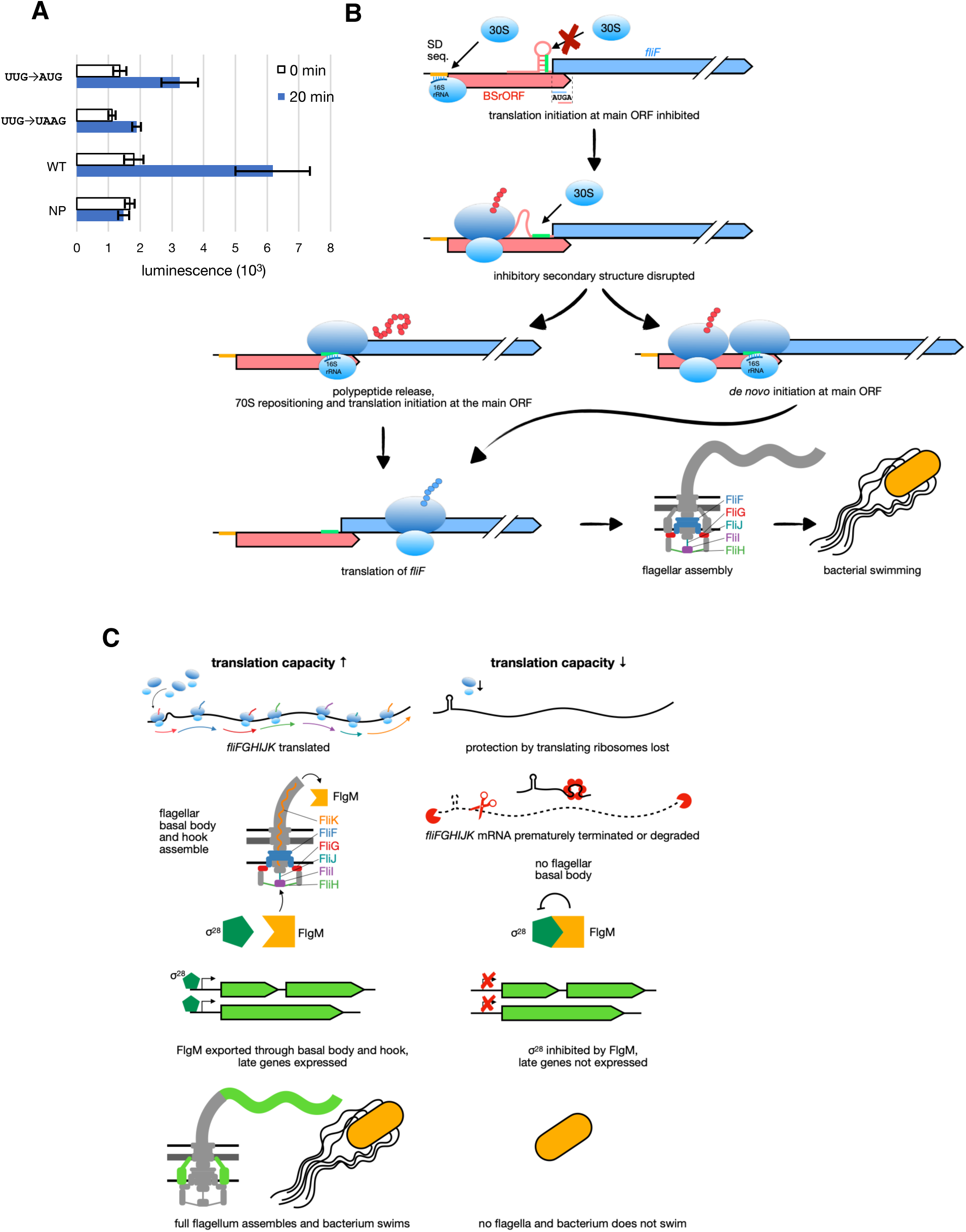
**A**, Luciferase reporter signals in wild-type *Salmonella* from constructs containing wild-type (WT) *Salmonella fliF-*tesORF sequence or the UUG→UAAG or UUG→AUG start codon mutation. NP = luciferase reporter construct with no promoter. Luminescence at 0 and 20 min after induction is shown in white and blue, respectively. **B**, Summary model for tesORF-mediated control of translation initiation. An inhibitory RNA secondary structure occludes the Shine–Dalgarno sequence of the main ORF and limits translation initiation. Translation of the upstream tesORF disrupts this structure, or prevents its formation, exposing the main ORF Shine–Dalgarno sequence. After tesORF termination, the ribosome may engage the adjacent Shine–Dalgarno sequence and initiate translation at the main ORF. Exposure of the Shine–Dalgarno sequence also permits *de novo* initiation by newly recruited ribosomes. **C**, Proposed model linking tesORF-mediated control of flagellar biosynthesis to cellular translation capacity. When translation capacity is sufficient, tesORF is translated, permitting *fliF* translation and potentially stabilizing the *fliFGHIJK* mRNA through ribosome loading. This enables production of the flagellar basal body and export ATPase complex. Completion of the hook-basal body permits efficient export of the anti-sigma factor FlgM, relieving inhibition of the flagellar sigma factor σ^28^ and allowing synthesis of late flagellar proteins, including the filament and stator complex. In this model, reduced cellular translation capacity limits tesORF translation. Consequently, *fliF* translation is inefficient, potentially exposing the *fliFGHIJK* mRNA to degradation or promoting premature transcription termination. Impaired flagellar assembly limits FlgM export, maintaining σ^28^ inhibition and reducing synthesis of late flagellar proteins. Thus, tesORF may link cellular translational capacity to hierarchical flagellar biosynthesis.

## Discussion

We identified a previously unannotated bacterial short regulatory open reading frame, tesORF, that activates expression of the middle flagellar operon *fliFGHIJK* in *Salmonella* and *E. coli*. This element initiates at a non-canonical UUG start codon, terminates with a UGA stop codon that overlaps the *fliF* start codon, and is embedded within an RNA structure that sequesters the downstream *fliF* SD sequence. Genetic, reporter, ribosome profiling, RNA-Seq, and phenotypic analyses support a model in which translation through tesORF remodels this inhibitory RNA structure or prevents its formation as the mRNA is synthesized, exposing the *fliF* ribosome-binding site and enabling efficient downstream translation initiation (figure 6B). Synonymous mutations disrupting this structure bypassed the requirement for tesORF translation (figures 5D and S8A). Conversely, premature stop codons generally reduced reporter activity, consistent with the requirement for ribosome progression through the hairpin-forming region (figures 5E and S8D). Together, these findings support a regulatory role for ribosome transit rather than a specific encoded peptide.

This architecture raises the question of why *fliF* initiation is gated by tesORF translation rather than by a constitutively accessible SD sequence. One possibility is that the tesORF couples flagellar biogenesis to translational capacity. Flagellum assembly is an energetically expensive, multi-operon programme requiring coordinated synthesis of basal body, hook, filament, export, motor, and regulatory components. Premature activation under conditions of limited ribosome availability could risk inefficient expression, incomplete assembly, and wasteful expenditure of cellular resources. By embedding the *fliF* SD sequence within an inhibitory RNA structure, *Salmonella* keeps the middle flagellar operon restrained unless ribosomes successfully initiate and translate the upstream tesORF. In this model, tesORF could act as a local translational checkpoint, potentially linking *fliF* initiation to ribosome availability and elongation capacity. Whether tesORF responds to changes in cellular translational capacity remains to be tested.

This mechanism is related to bacterial translational coupling but differs from canonical examples in which translation of one protein-coding gene promotes expression of the next. Overlapping stop–start codons are common in bacteria, including 444 of 4463 annotated genes on the *Salmonella* chromosome (FQ312003.1), and they often coordinate expression of adjacent genes. Here, however, the upstream ORF (uORF) appears to function primarily as a regulatory module that licenses expression of the downstream structural gene. The overlap between the tesORF stop codon and the *fliF* start codon may promote ribosome reinitiation, but reporter activity was maintained when tesORF translation extended beyond the *fliF* start codon. This supports a contribution from *de novo* initiation at *fliF* and argues against an exclusive requirement for reinitiation by the same ribosome.

The mechanism of *fliF*-tesORF regulation differs from many characterized eukaryotic uORFs and bacterial transcriptional attenuators. Eukaryotic uORFs regulate downstream translation through scanning, reinitiation, and context-dependent ribosome availability, often repressing main ORF expression, although exceptions such as *ATF4* permit stress-dependent activation [18] [54]. Bacterial transcriptional attenuators operate by a different logic, coupling translation of a short leader-peptide-encoding ORF to formation or prevention of transcriptional terminators, as seen in amino acid biosynthesis operons, pyrimidine synthesis operons, and the *Salmonella* virulence-associated operons *mgtCBR* and *mgtA* [55] [56] [57] [58] [59]. By contrast, the *fliF-*tesORF acts locally at the level of translation: its translation directly changes the accessibility of a downstream ribosome-binding site.

Like the *fliF*-tesORF, certain characterized bacterial ORFs control access to the downstream SD sequence. Some of these encode an arrest peptide, which induces programmed ribosome pausing and subsequent conditional mRNA secondary structure remodelling, allowing translation of the downstream gene. These include the well-characterized secretion monitor gene *secM* (figure S5B) [60] [61] [62], uORFs controlling antimicrobial resistance genes, such as *ermCL* [63] [64] [65] [66] [67] [68], and the *Bacillus* membrane insertion and folding monitor gene *mifM* [69]. Another noteworthy example is the dual-function *speFL* (a.k.a. *orf34*), located upstream of the putrescine–ornithine metabolism and transport operon *speF-potE*, which relieves transcriptional attenuation and exposes the occluded *speF* ribosome-binding site in response to ornithine and arginine levels, respectively [70] [71]. Furthermore, a number of small uORFs have been discovered in *E. coli* and *Sinorhizobium meliloti* that enhance the translation of the downstream gene by translational coupling, but they have not yet been mechanistically characterized [26] [27]. Another reported example of leader ORF-dependent translational activation is the *idpL*-mediated regulation of *iraD* in *E. coli*, which is itself regulated by the RNA-binding protein CsrA [28]. Whether additional regulatory inputs contribute to *fliF*-tesORF function remains to be explored.

The consequences of the tesORF-mediated regulatory event propagate through the flagellar gene expression hierarchy. Although tesORF acts locally on translation of the first gene in the operon, *fliF*, loss of this regulation affects expression of the rest of the operon and the assembly of the flagellar apparatus. *fliFGHIJK* encodes core components of the flagellar basal body and export machinery, and insufficient expression of this operon impairs assembly of the flagellar export apparatus. Defective flagellar assembly prevents efficient export of the anti-sigma factor FlgM, maintaining inhibition of σ^28^ and reducing expression of late flagellar genes [46]. Consistent with this model, disruption of tesORF translation reduces *fliF* translation, lowers *fliFGHIJK* transcript abundance, potentially through reduced protection by translating ribosomes [42] or premature termination due to exposure of cryptic Rho utilization sites [43] [44] [45], and impairs downstream flagellar function. A small upstream translational event is therefore amplified into operon-level and regulon-level effects on flagellar biogenesis (figure 6B).

The regulatory defect also has consequences beyond motility. Flagella contribute to swimming, swarming, chemotaxis, adhesion, and biofilm formation—key functions that facilitate host colonization as well as invasion of permissive host cells [72] [73] [74]. Mutation of the tesORF start codon reduced secretion of the SPI-1 injectisome translocon proteins SipB and SipC, despite no marked change in expression of SPI-1 injectisome and effector genes. Because previous work argues against direct secretion of SipB and SipC through the flagellar apparatus in *Salmonella*, this phenotype is unlikely to reflect simple rerouting through the flagellar type III secretion system [75] [76]. Instead, it suggests that impaired expression of *fliFGHIJK* affects SPI-1 secretion through an indirect mechanism that remains to be defined, placing the tesORF at a regulatory intersection between flagellar assembly, motility, and virulence-associated secretion.

The conservation of the *fliF-*tesORF across motile Enterobacterales suggests that this regulatory architecture has been maintained by selective pressure on flagellar gene expression. Its evolutionary origin may lie in reorganisation of the flagellar locus. In non-Enterobacterales Gammaproteobacteria, including *Vibrio parahaemolyticus* and *Pseudomonas aeruginosa*, *fliE* is located immediately upstream of *fliF* within the same operon, often separated by fewer than 40 nucleotides, a spacing compatible with translational coupling (table S3) [17] [77]. In Enterobacterales, by contrast, *fliE* is encoded on the opposite strand to *fliF*. One possibility is that tesORF preserves an upstream translation-dependent regulatory mechanism following reorganization of the locus, although this evolutionary relationship remains to be established.

Notably, *Yersinia* possesses both a *fliFGHIJK* operon with a tesORF and a *fliEFGHIJ* operon in a second flagellar cluster (NZ_CP009792.1; YPK-0710–YPK-0715). In our swimming assays, *fliF*-tesORF start codon disruption in *Y. pseudotuberculosis* reduced but did not abolish swimming motility (figure 2D). The presence of the second operon could allow a low level of swimming motility, even though its transcript abundance at 25 °C is about 50-fold lower than that of the tesORF-containing *fliFGHIJK* operon (figure S4A) [31] [32]. Alternatively, the mRNA secondary structure in *Yersinia* could have a weaker inhibitory effect, or tesORF or *fliF* translation could initiate at a different site in the tesORF mutant background; the sequence upstream of *fliF* contains multiple potential initiation sites (figure S2A), though their predicted ribosome-binding site strengths are low.

Together, our findings establish *fliF-*tesORF as a conserved regulatory element whose local effect on translation has consequences for an entire flagellar operon and the wider flagellar regulon, linking RNA structural regulation to motility and virulence-associated phenotypes. More broadly, our results suggest that conserved unannotated short ORFs may function through ribosome transit rather than their encoded peptides, highlighting a potentially underappreciated layer of translational control in bacterial genomes.

## Materials and Methods

### Strains and plasmids

The strains and plasmids used in this study are listed in table S4 [78] [79] [80] [81] [82] [32] [83] [84].

### Bacterial genome modification

Chromosomal modifications of *E. coli* and *Salmonella* were performed using the pWRG730 and pWRG717 constructs following a protocol based on Hoffmann *et al*., 2017 [85]. Genes were deleted by recombination with an *aph*-I-*Sce*I kanamycin resistance cassette amplified from pWRG717 with primers that incorporate regions homologous to the target gene. To introduce point mutations, the resistance cassette was replaced by recombination with a PCR product incorporating the point mutation and at least 400 bp flanking the gene of interest.

*Y. pseudotuberculosis fliF*-tesORF start codon mutant was generated by homologous recombination using the pDM4 plasmid [83]. First, the region of interest was cloned into pDM4. The *fliF*-tesORF start codon (TTG) was then replaced with a stop codon (TAA) using the Q5 Site-Directed Mutagenesis Kit (NEB) and mutagenic primers. The resulting recombinant plasmid pBO8751 was introduced into *E. coli* S17-1 *λ-pir* cells (donor) and subsequently transferred into *Y. pseudotuberculosis* (recipient) by conjugation. Putative tesORF mutants were confirmed by DNA sequencing.

### Luciferase reporter and swimming motility assays

The luciferase reporter assays were performed as described in Bryant *et al.*, 2023 [29]. For swimming motility, cultures were grown in LB broth to OD_600_ 1; *Salmonella* and *E. coli* at 37 °C and *Yersinia* at 25 °C. 2 μL of culture was inoculated into soft tryptone agar (0.3% agar, 10 g L^−1^ tryptone, 5 g L^−1^ NaCl). *Salmonella* and *E. coli* plates were incubated at 37 °C for 4–6 h, and *Yersinia* plates were incubated at room temperature for four days unless otherwise stated. For strains carrying pTrc99a vectors, LB broth and motility plates were supplemented with 100 μg mL^−1^ carbenicillin and 100 μM isopropyl β-D-1-thiogalactopyranoside (IPTG).

### Negative stain electron microscopy

The strains were grown to OD_600_ of 1.0, harvested by centrifugation (500 rcf, 3 min) and resuspended in phosphate-buffered saline (PBS). Carbon-coated copper grids (300 mesh size) were glow-discharged for 30 seconds at 15 mA. 20 μL of bacterial suspension was applied to the freshly glow-discharged grids and incubated for 2 min. Excess sample was blotted away, and the grids were washed thrice in 20 μL of distilled water with blotting in between. The grids were then incubated with 10 μL of uranyl acetate for 30 seconds, blotted once more to remove excess uranyl acetate, washed once with 10 μL distilled water, and left to air dry for 5 min before storage. The grids were imaged on a FEI Tecnai electron microscope with CCD camera image acquisition.

### Protein export assays

Export assays were performed as previously described [86]. Fractions were analysed by immunoblotting with anti-FlgD [87], anti-FlgL [88], anti-SipB [89], anti-SipC (tgcBIOMICS, #tgc-a201-1), anti-FlhA [86], anti-FLAG MR antibody (F3165-1MG), anti-Myc (9B11, 2276S) and anti-GroEL sera (Abcam, #ab90522) primary antibodies, and anti-Rabbit IgG, HRP conjugate (Promega, W4011) or anti-Mouse IgG, HRP conjugate (Promega, W4021) secondary antibodies. The secondary antibodies were detected using ECL western blot detection reagent (Amersham, RPN2232) and imaged with a BioRad ChemiDoc MP imaging system.

### Cytotoxicity assay

96-well plates were seeded with murine immortalized bone-marrow-derived macrophages (iBMDMs) in Dulbecco’s modified Eagle medium (Sigma) with 10% FBS and 2 mM L-glutamine. Conditioned medium was obtained from overnight iBMDM culture in T75 flasks seeded at the equivalent concentration. Exponential-phase *Salmonella* suspensions were washed in PBS and twice in the conditioned medium, added to the wells with iBMDMs for a multiplicity of infection (MOI) of 10, and incubated at 37 °C. After 15 min, gentamicin was added to a final concentration of 85 ng μL^−1^. One hour post infection, the medium in the wells was sampled and colorimetric assay was continued based on the CytoTox 96 (Promega) protocol. The experiment was carried out with negative and 0.8% Triton X-100 lysis control groups, the values of which were used for normalization. Equivalence of the bacterial suspension concentrations was verified by serial dilution and overnight culture of droplets on LB agar followed by colony counting.

### RNA structure probing

To perform enzymatic structure probing at different temperatures, the DNA region of interest was first modified at the 5′ end with a T7 promoter sequence and at the 3′ end with an *Eco*RV restriction site and then cloned into a pUC18 vector, generating pBO8349. Following digestion of the plasmid by *Eco*RV, RNA was synthesized by *in vitro* run-off transcription using T7 RNA polymerase (Thermo Scientific). To purify the RNA, samples were separated on a 6% polyacrylamide gel at 1000 V for approximately 3 h. RNA was detected on a silica gel plate using a UV lamp and excised from the gel. The RNA was eluted in elution buffer (10 mM EDTA, 0.5% (w/v) SDS, and 0.1 M sodium acetate; buffer adjusted to pH 5.6), precipitated in ethanol with sodium acetate and resuspended in nuclease-free water. Purified RNA was subsequently dephosphorylated using an alkaline phosphatase (FastAP, Thermo Scientific, Waltham, USA). Radioactive labelling with ^32^P at the 5′ end of the dephosphorylated RNA was performed according to Brantl and Wagner, 1994 [90]. As described in Waldminghaus *et al.*, 2007 [91], limited digestion of 30000 cpm of radiolabelled RNA was performed with ribonuclease T1 (0.0016 U; Invitrogen, AM2283) and T2 (0.053 U; Biozol) in 1×TN buffer (20 mM Tris acetate pH 7, 100 mM NaCl) at 25 °C, 37 °C, and 42 °C. In addition, an alkaline hydrolysis ladder [90] and a T1 ladder were prepared from the examined RNA. For the T1 ladder, 30000 cpm of labelled RNA was adjusted to 2 μL with nuclease-free water, heated with 2 μL of sequencing buffer (provided with RNase T1 in the kit) to 90 °C, and then incubated with 1 μL of the RNase T1 in a 5 μL reaction at 37 °C for 5 min. The ladders and the examined RNA were then applied to an 8% polyacrylamide gel and run at 1800 V for approximately 2 h. The gel was then transferred onto Whatman paper, covered with Saran wrap, and dried in a Bio-Rad Model 583 gel dryer at 80 °C for 1.5 h. For detection of the radiolabelled RNA, the dried gel was placed in an autoradiography cassette, and a storage phosphor screen (GE Healthcare) was positioned on top of the gel and incubated overnight. The following day, the radiolabelled RNA was detected using a GE Amersham Typhoon imager (GE Healthcare).

### RNA isolation and qPCR

*Salmonella* LB culture was mixed with acidic phenol–chloroform by vigorous shaking. RNA was isolated by acidic phenol–chloroform extraction and GlycoBlue-assisted precipitation, treated with DNase I (Thermo Fisher Scientific), and re-purified. RNA integrity was assessed by agarose gel electrophoresis. cDNA was synthesized from 500 ng RNA using random hexamers and M-MLV reverse transcriptase (Promega), with no-reverse-transcriptase controls. qPCR was performed using iTaq Universal SYBR Green Supermix (Bio-Rad) on a QuantStudio 6 Pro system with 500 nM primers targeting *luxA* (CCTTATGTTGCTGCCGCACACC; TTGTCGAACCGGATGGGCAGTC) or *lppA* (SL1344_1311; GTACTAAACTGGTACTGGGCG; GCTGATCGATTTTAGCGTTGC). Relative *luxA* transcript abundance was calculated using the ΔΔCq method, normalized to *lppA* and calibrated to the wild-type *fliF*-tesORF reporter at 0 min post-induction.

### Ribosome profiling, RNA-Seq, and Bioinformatic analysis

Ribo-Seq and RNA-Seq data from Bryant *et al.*, 2023 [29] and this study were processed as described therein, using the *Salmonella* SL1344 reference sequences FQ312003.1, HE654725.1, HE654726.1 and HE654724.1 and the *E. coli* reference sequence U00096.3. P-site offsets are listed in table S5. Only Ribo-Seq read lengths with clear periodicity were selected for analysis. CDS counts included RNA-Seq and phased Ribo-Seq reads, retaining genes with ≥50 Ribo-Seq reads across ≥10 positions in at least one compared sample. Counts were normalized by library size and CDS length using edgeR [92].

RNA structures were predicted with RNAfold and their free energies evaluated with RNAfold or RNAeval (ViennaRNA) [93]. Mutation effects were visualized with R4RNA [94], and ribosome-binding site strength was predicted using Ribosome Binding Site Calculator v1.0 [50]. GenBank sequences (table S6) were aligned using MAFFT, manually adjusted, and visualized in Jalview [95] [96].

## Data availability

Raw and processed data are available from ArrayExpress accessions E-MTAB-17328 and E-MTAB-17837 or can be found in the supplementary tables.

## Acknowledgements

We would also like to thank Nancy Standart and Ben Luisi for discussions and comments on the manuscript. FL was supported by a BBSRC project grant awarded to BYWC [BB/V006096/1]. BYWC and OJB were supported by a Medical Research Council Fellowship to BYWC [MR/R021821/1]. The BYWC laboratory is supported by a Medical Research Council Fellowship [MR/R021821/1] and BBSRC project grants [BB/X001261/1, BB/V017780/1 and BB/V006096/1]. FN is supported by the German Research Foundation (DFG grant NA 240/14-1).

## Author contributions

B.Y.W.C., O.J.B., F.L. conceived the research and designed experiments. O.J.B. generated the bacterial mutants, F.L. generated Ribo-Seq and RNA-Seq libraries, F.L. and O.J.B. performed molecular experiments. E.A. performed RT-qPCR. D.M performed the RNA structural probing, F.L. performed most of the bioinformatic analysis. B.Y.W.C., O.J.B., F.L., D.M and F.N wrote the manuscript.

## Supplementary information Supplementary figures S1 to S8

**Table S1**: Normalized Ribo-Seq and RNA-Seq counts from wild-type *Salmonella* and the tesORF UUG→UAAG mutant.

**Table S2**: Ribosome-binding site strengths of *fliF* and the *fliF*-tesORF in the examined strains and constructs calculated using the Ribosome Binding Site Calculator version 1.0 [50].

**Table S3**: Distances between the *fliF* and *fliE* coding sequences in motile non-Enterobacterales Gammaproteobacteria with *fliF* and *fliE* on the same strand.

**Table S4**: Strains and plasmids used in this study.

**Table S5**: Distances between inferred ribosome P-site positions and Ribo-Seq read 5′ ends.

**Table S6**: GenBank accession numbers of motile Enterobacterales genomes shown in figures 1C and S2.

