## Supplementary Information for "A ribosome-dependent small-ORF switch controls flagellar gene expression and virulence"

Betty Y-W Chung

Franz Narberhaus

'For the purpose of open access, the author has applied a Creative Commons Attribution (CC BY) licence to any Author Accepted Manuscript version arising.'

### Figure S1

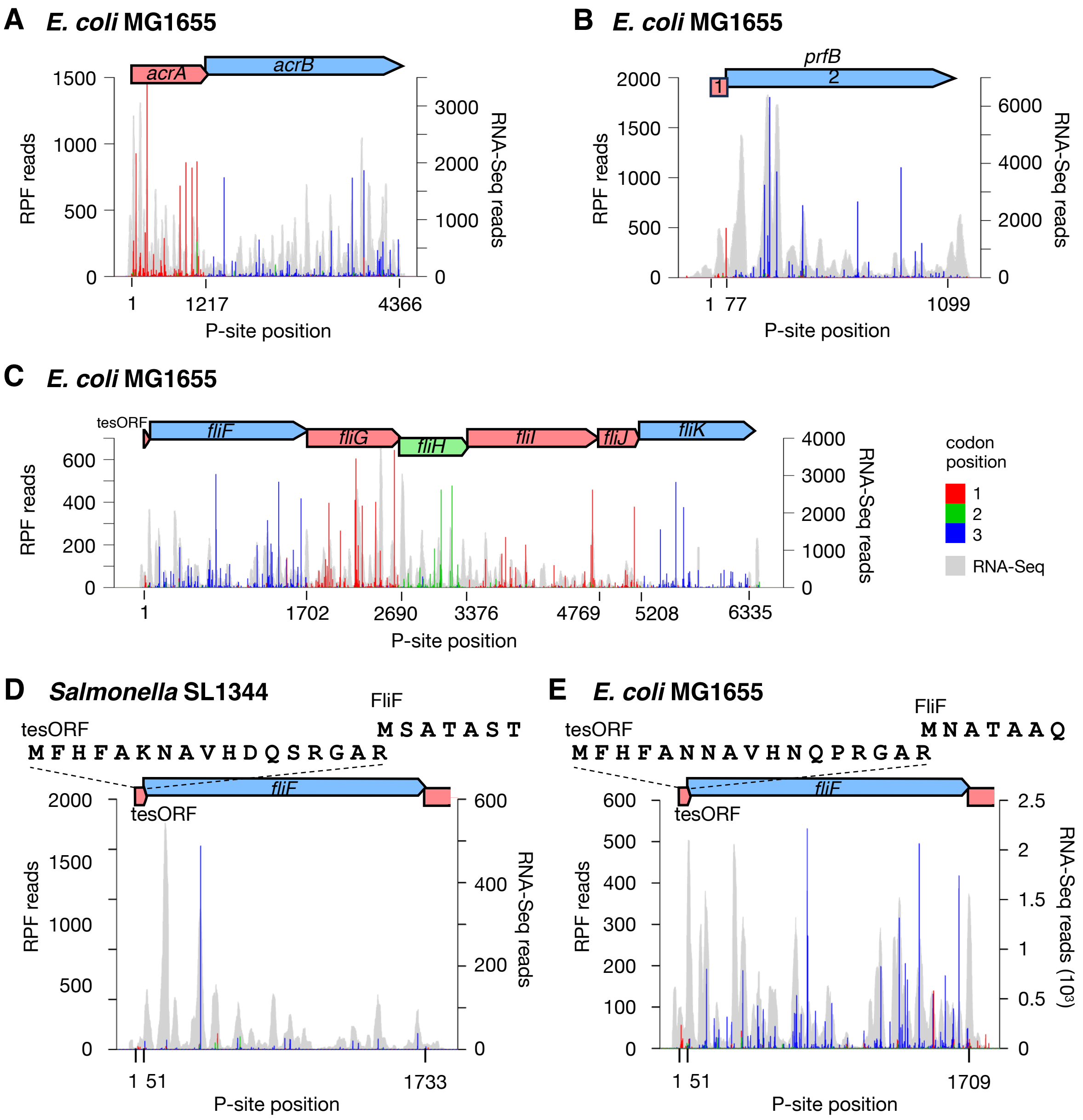

A
E. coli MG1655
acrA
acrB
1500
1000
500
0
RPF reads
3000
2000
1000
0
RNA-Seq reads
1
1217
4366
P-site position
B
E. coli MG1655
prfB
1
2
2000
1500
1000
500
0
RPF reads
6000
4000
2000
0
RNA-Seq reads
1
77
1099
P-site position
C
E. coli MG1655
tesORF
fliF
fliG
fliH
fliI
fliJ
fliK
600
400
200
0
RPF reads
4000
3000
2000
1000
0
RNA-Seq reads
1
1702
2690
3376
4769
5208
6335
P-site position
codon position
1
2
3
RNA-Seq
D
Salmonella SL1344
FliF
tesORF
M S A T A S T
M F H F A K N A V H D Q S R G A R
fliF
tesORF
2000
1500
1000
500
0
RPF reads
600
400
200
0
RNA-Seq reads
1
51
1733
P-site position
E
E. coli MG1655
FliF
tesORF
M N A T A A Q
M F H F A N N A V H N Q P R G A R
fliF
tesORF
600
500
400
300
200
100
0
RPF reads
2.5
2
1.5
1
0.5
0
RNA-Seq reads (10³)
1
51
1709
P-site position

**Figure S1**

**A–C**, Visualization of translation of the *acrAB* operon (**A**; two genes in different reading frames), the *prfB* gene (**B**; a single gene with a frameshift; the two frames are annotated 1 and 2), and the *fliFGHIJK* operon (**C**) in *E. coli* MG1655. RPF read counts are not capped. The coding sequences are coloured by their frames relative to the first nucleotide of the 5′-most coding sequence. Red, green, and blue bars indicate Ribo-Seq reads mapping to codon positions 1, 2, and 3, respectively, relative to the first nucleotide of the 5′-most coding sequence. Grey shaded peaks show RNA-Seq coverage.

**D** and **E,** Visualization of translation of *fliF* and the tesORF in *Salmonella* SL1344 (**D**) and *E. coli* MG1655 (**E**). RPF reads are not capped. The amino acid sequences encoded by the tesORF and *fliF* (partial) are displayed above.

### Figure S2

**A**

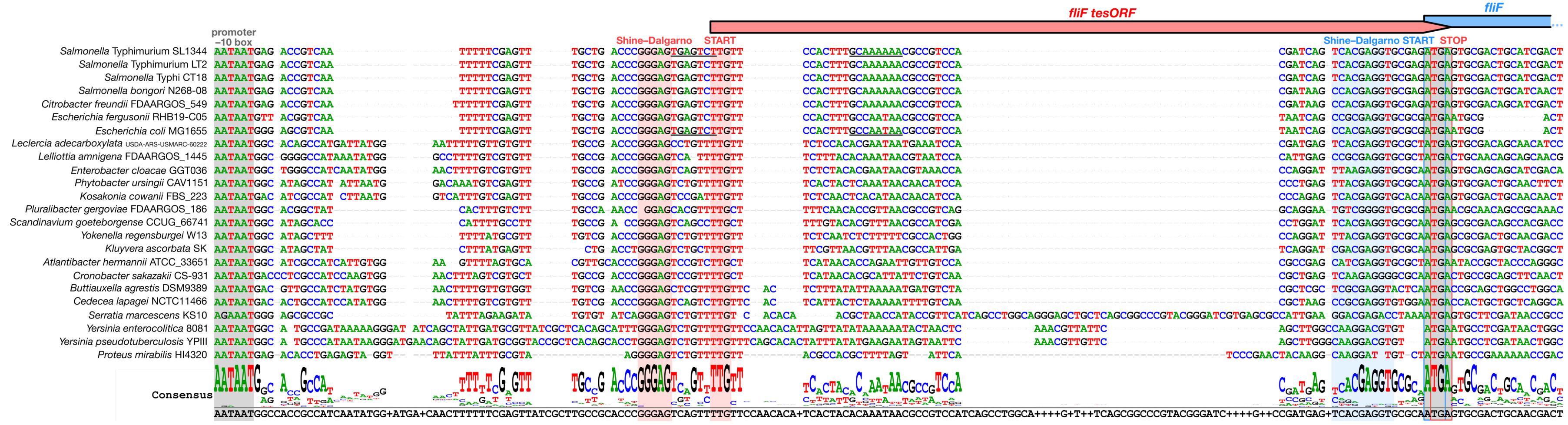
fliF tesORF
fliF
promoter
−10 box
Shine–Dalgarno
START
Shine–Dalgarno START
STOP
Salmonella Typhimurium SL1344
Salmonella Typhimurium LT2
Salmonella Typhi CT18
Salmonella bongori N268-08
Citrobacter freundii FDAARGOS_549
Escherichia fergusonii RHB19-C05
Escherichia coli MG1655
Leclercia adecarboxylata USDA-ARS-USMARC-60222
Lelliottia amnigena FDAARGOS_1445
Enterobacter cloacae GGT036
Phytobacter ursingii CAV1151
Kosakonia cowanii FBS_223
Pluralibacter gergoviae FDAARGOS_186
Scandinavium goeteborgense CCUG_66741
Yokenella regensburgei W13
Kluyvera ascorbata SK
Atlantibacter hermannii ATCC_33651
Cronobacter sakazakii CS-931
Buttiauxella agrestis DSM9389
Cedecea lapagei NCTC11466
Serratia marcescens KS10
Yersinia enterocolitica 8081
Yersinia pseudotuberculosis YPIII
Proteus mirabilis HI4320
Consensus

**B**

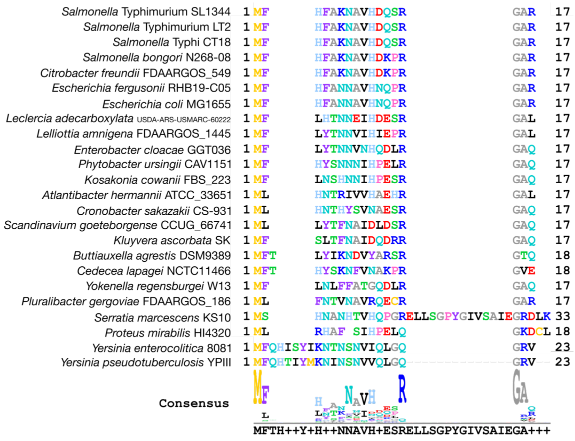
Salmonella Typhimurium SL1344
Salmonella Typhimurium LT2
Salmonella Typhi CT18
Salmonella bongori N268-08
Citrobacter freundii FDAARGOS_549
Escherichia fergusonii RHB19-C05
Escherichia coli MG1655
Leclercia adecarboxylata USDA-ARS-USMARC-60222
Lelliottia amnigena FDAARGOS_1445
Enterobacter cloacae GGT036
Phytobacter ursingii CAV1151
Kosakonia cowanii FBS_223
Atlantibacter hermannii ATCC_33651
Cronobacter sakazakii CS-931
Scandinavium goeteborgense CCUG_66741
Kluyvera ascorbata SK
Buttiauxella agrestis DSM9389
Cedecea lapagei NCTC11466
Yokenella regensburgei W13
Pluralibacter gergoviae FDAARGOS_186
Serratia marcescens KS10
Proteus mirabilis HI4320
Yersinia enterocolitica 8081
Yersinia pseudotuberculosis YPIII
Consensus

**Figure S2**

**A**, Alignment of the region upstream of *fliF* shows conservation of the translation-enhancing small ORF among Enterobacterales. A potential $\sigma^{28}$ promoter sequence is underlined in *Salmonella* SL1344 and *E. coli*.

**B**, Alignment of the amino acid sequences encoded by the *fliF*-tesORF.

### Figure S3

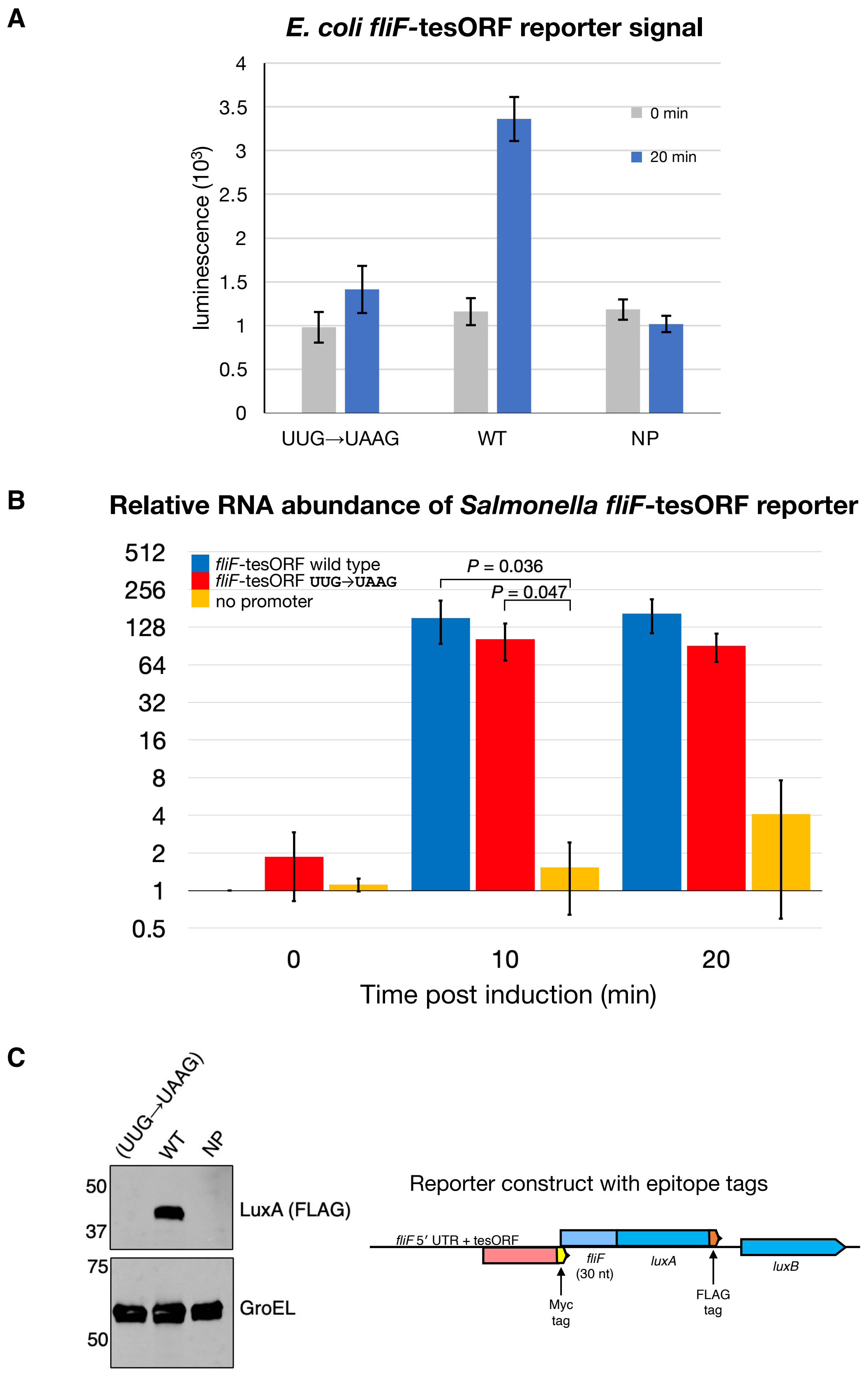
A
E. coli fliF-tesORF reporter signal
luminescence (10³)
0
0.5
1
1.5
2
2.5
3
3.5
4
0 min
20 min
UUG→UAAG
WT
NP
B
Relative RNA abundance of Salmonella fliF-tesORF reporter
fliF-tesORF wild type
fliF-tesORF UUG→UAAG
no promoter
P = 0.036
P = 0.047
512
256
128
64
32
16
8
4
2
1
0.5
0
10
20
Time post induction (min)
C
(UUG→UAAG)
WT
NP
50
37
75
50
LuxA (FLAG)
GroEL
Reporter construct with epitope tags
fliF 5′ UTR + tesORF
fliF
(30 nt)
luxA
luxB
Myc
tag
FLAG
tag

**Figure S3**

**A**, Luciferase reporter signals in wild-type *E. coli* from constructs containing the wild-type (WT) *E. coli fliF*-tesORF sequence or the UUG→UAAG start codon mutation. NP = luciferase reporter construct with no promoter. Luminescence at 0 and 20 min after induction is shown in white and blue, respectively.

**B,** Relative RNA abundance in *Salmonella* carrying *fliF-luxAB* luciferase reporter constructs corresponding to those in figure 2A, at 0, 10, and 20 min after induction with 50 ng $mL^{-1}$ anhydrotetracycline. Error bars represent the SEM of relative RNA abundance from two biological replicates. The low transcript abundance of the no-promoter construct resulted in greater variability between biological replicates than for the *fliF*-tesORF start codon mutant (UUG→UAAG) and wild-type constructs. The displayed *P* values were calculated using a two-tailed Student's *t*-test.

**C**, Design and validation of the epitope-tagged reporter construct. The reporter contains the *fliF* 5′ UTR and tesORF followed by a coding sequence encoding a Myc-tagged 30-residue segment of FliF and FLAG-tagged LuxA. Immunoblotting confirmed LuxA-FLAG expression from the wild-type (WT) reporter, whereas LuxA-FLAG was not detected following mutation of the tesORF start codon (UUG→UUAG) or in the no-plasmid (NP) control. GroEL was used as a loading control.

### Figure S4

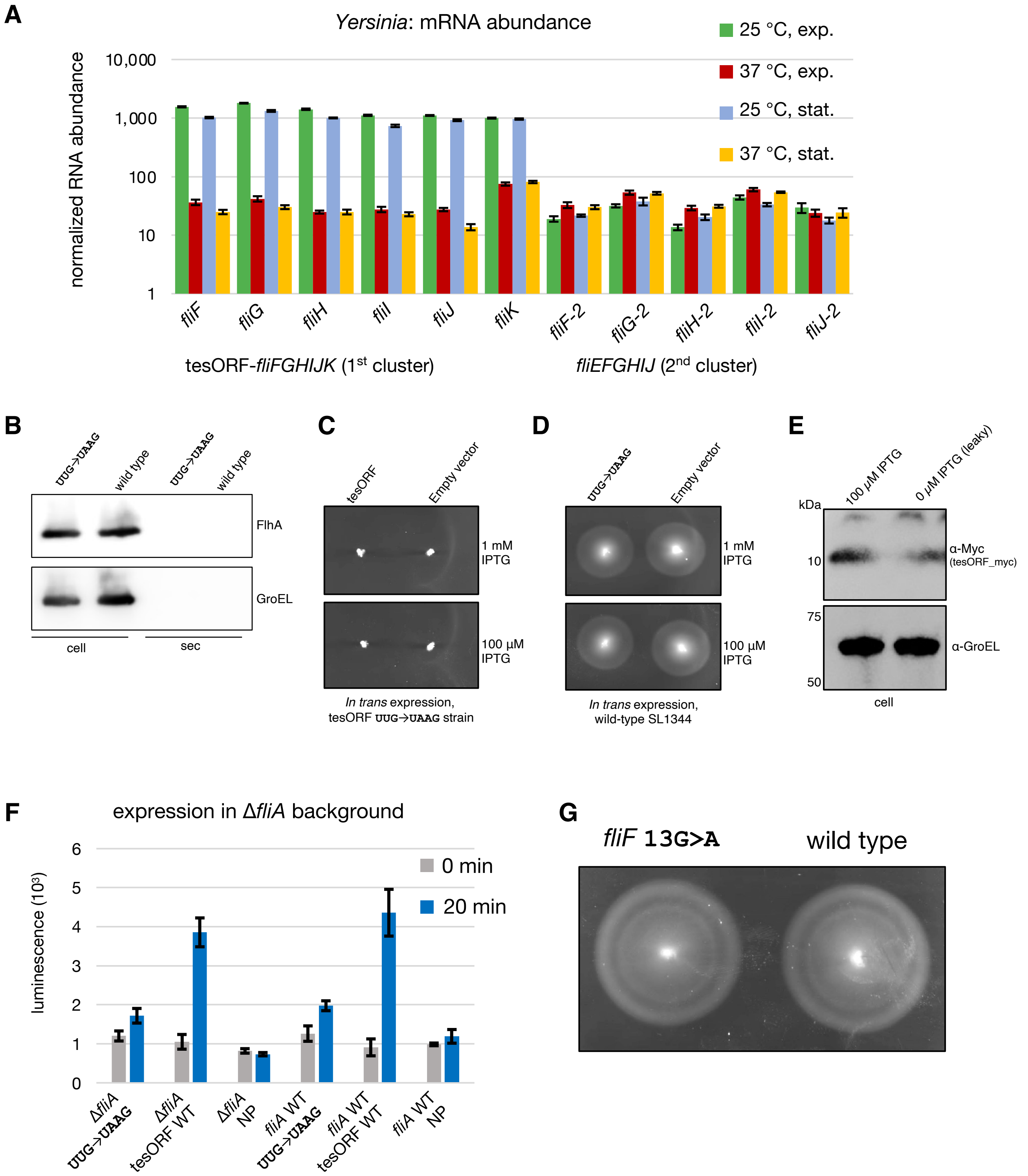
A
Yersinia: mRNA abundance
25 °C, exp.
37 °C, exp.
25 °C, stat.
37 °C, stat.
normalized RNA abundance
10,000
1,000
100
10
1
fliF
fliG
fliH
fliI
fliJ
fliK
fliF-2
fliG-2
fliH-2
fliI-2
fliJ-2
tesORF-fliFGHIJK (1st cluster)
fliEFGHIJ (2nd cluster)
B
UUG->UAAG
wild type
UUG->UAAG
wild type
FlhA
GroEL
cell
sec
C
tesORF
Empty vector
1 mM IPTG
100 µM IPTG
In trans expression, tesORF UUG→UAAG strain
D
UUG->UAAG
Empty vector
1 mM IPTG
100 µM IPTG
In trans expression, wild-type SL1344
E
100 µM IPTG
0 µM IPTG (leaky)
kDa
10
75
50
α-Myc (tesORF_myc)
α-GroEL
cell
F
expression in ΔfliA background
luminescence (10³)
6
5
4
3
2
1
0
0 min
20 min
ΔfliA UUG->UAAG
ΔfliA tesORF WT
ΔfliA NP
fliA WT UUG->UAAG
fliA WT tesORF WT
fliA WT NP
G
fliF 13G>A
wild type

**Figure S4**

**A**, Normalized mRNA abundance of *Yersinia fliFGHIJK* operon (with tesORF) and *fliEFGHIJ* operon (second flagellar cluster) in the exponential and stationary phase at 25 °C and 37 °C. Data derived from Javadi *et al.*, 2026, and normalized by library size and CDS length. Error bars represent the standard error of the mean from three replicates.

**B,** Analysis of non-exported control proteins in the export assay comparing wild-type *Salmonella* and the tesORF UUG→UAAG mutant. Corresponds to figure 2E.

**C**, Swimming motility assay of the *Salmonella fliF*-tesORF UUG→UAAG mutant expressing the wild-type or mutated *fliF*-tesORF CDS from the IPTG-inducible promoter in pTrc99a.

**D**, Swimming motility assay of wild-type *Salmonella* expressing the wild-type or UUG→ UAAG mutant *fliF*-tesORF CDS from the IPTG-inducible promoter in pTrc99a.

**E**, Detection of the Myc-tagged tesORF-encoded peptide (tesORF_myc) by Western blotting in *Salmonella* carrying pTrc99-tesORF_myc, with or without induction with 100 µM IPTG (uninduced cultures allow leaky expression). Equal levels of GroEL were detected in the samples.

**F**, Luciferase reporter signals from constructs containing the wild-type or UUG→UAAG mutant *fliF*-tesORF sequence in wild-type *Salmonella* or a $\sigma^{28}$-deficient strain (Δ*fliA*). WT = wild-type SL1344. NP = luciferase reporter construct with no promoter.

**G**, Swimming motility assay comparing *Salmonella fliF* 13G>A (FliF Ala5Thr; suppressor D mutation with an intact *fliF*-tesORF start codon) with wild-type *Salmonella*.

### Figure S5

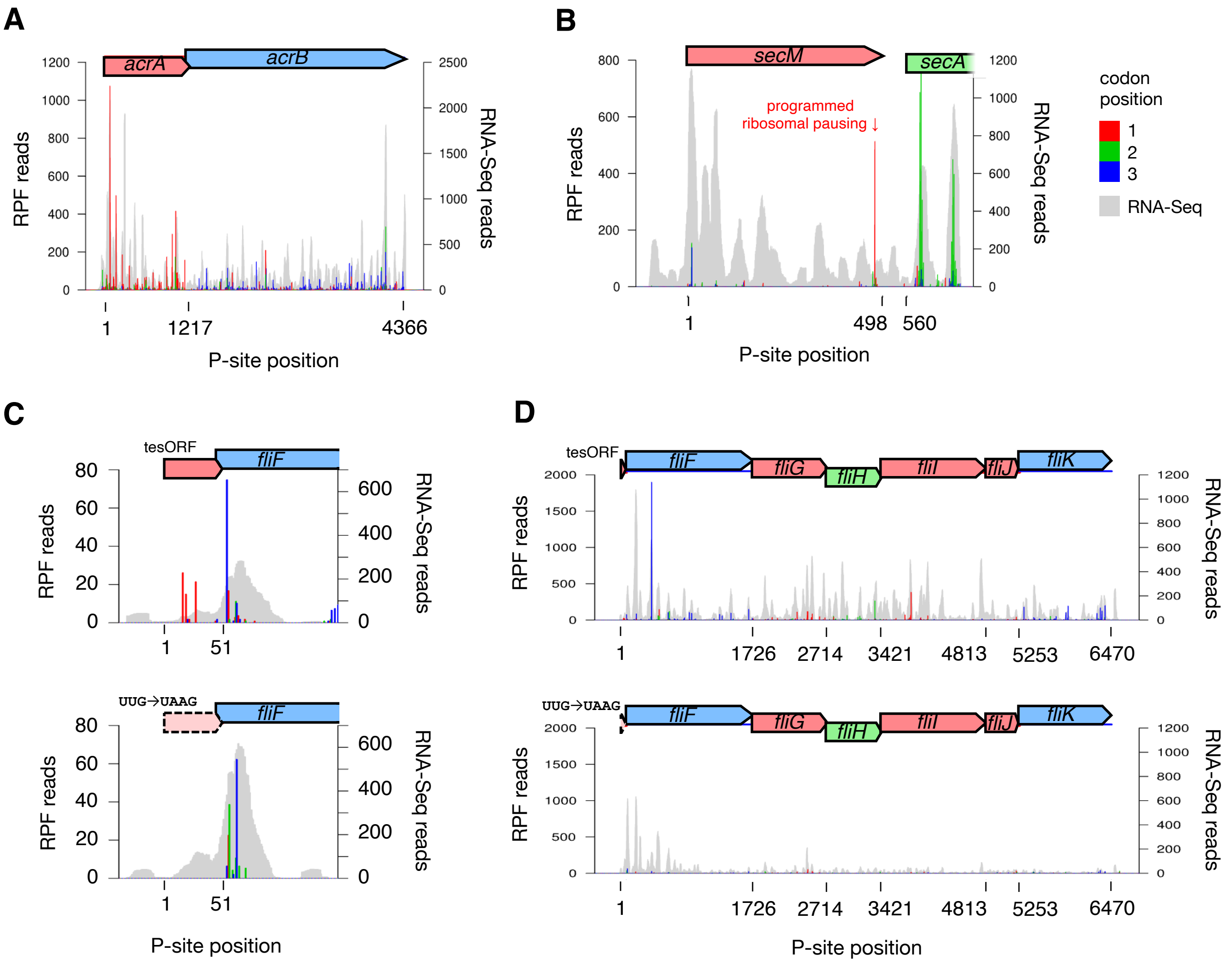
A
acrA
acrB
RPF reads
RNA-Seq reads
1 1217 4366
P-site position
B
secM
secA
programmed ribosomal pausing ↓
RPF reads
RNA-Seq reads
1 498 560
P-site position
codon position
1
2
3
RNA-Seq
C
tesORF
fliF
RPF reads
RNA-Seq reads
1 51
UUG→UAAG
fliF
RPF reads
RNA-Seq reads
1 51
P-site position
D
tesORF
fliF
fliG
fliH
fliI
fliJ
fliK
RPF reads
RNA-Seq reads
1 1726 2714 3421 4813 5253 6470
UUG→UAAG
fliF
fliG
fliH
fliI
fliJ
fliK
RPF reads
RNA-Seq reads
1 1726 2714 3421 4813 5253 6470
P-site position

**Figure S5**

**A**, Visualization of translation of the *acrAB* operon in the *Salmonella fliF*-tesORF UUG→UAAG mutant. Red, green, and blue bars indicate Ribo-Seq reads mapping to codon positions 1, 2, and 3, respectively, relative to the first nucleotide of the 5′-most coding sequence. Read positions correspond to inferred ribosomal P-site positions. Grey shaded peaks show RNA-Seq coverage.

**B**, Visualization of translation of *secM* in the *Salmonella fliF*-tesORF UUG→UAAG mutant. The programmed ribosomal pausing site is labelled with a red arrow.

**C**, Visualization of translation on *fliF*-tesORF in wild-type *Salmonella* and the *fliF*-tesORF UUG→UAAG mutant, with read counts normalized by library size.

**D**, Uncapped Ribo-Seq and parallel RNA-Seq read maps on the *fliFGHIJK* operon in wild-type *Salmonella* and the *fliF*-tesORF UUG→UAAG mutant, with read counts normalized by library size.

### Figure S6

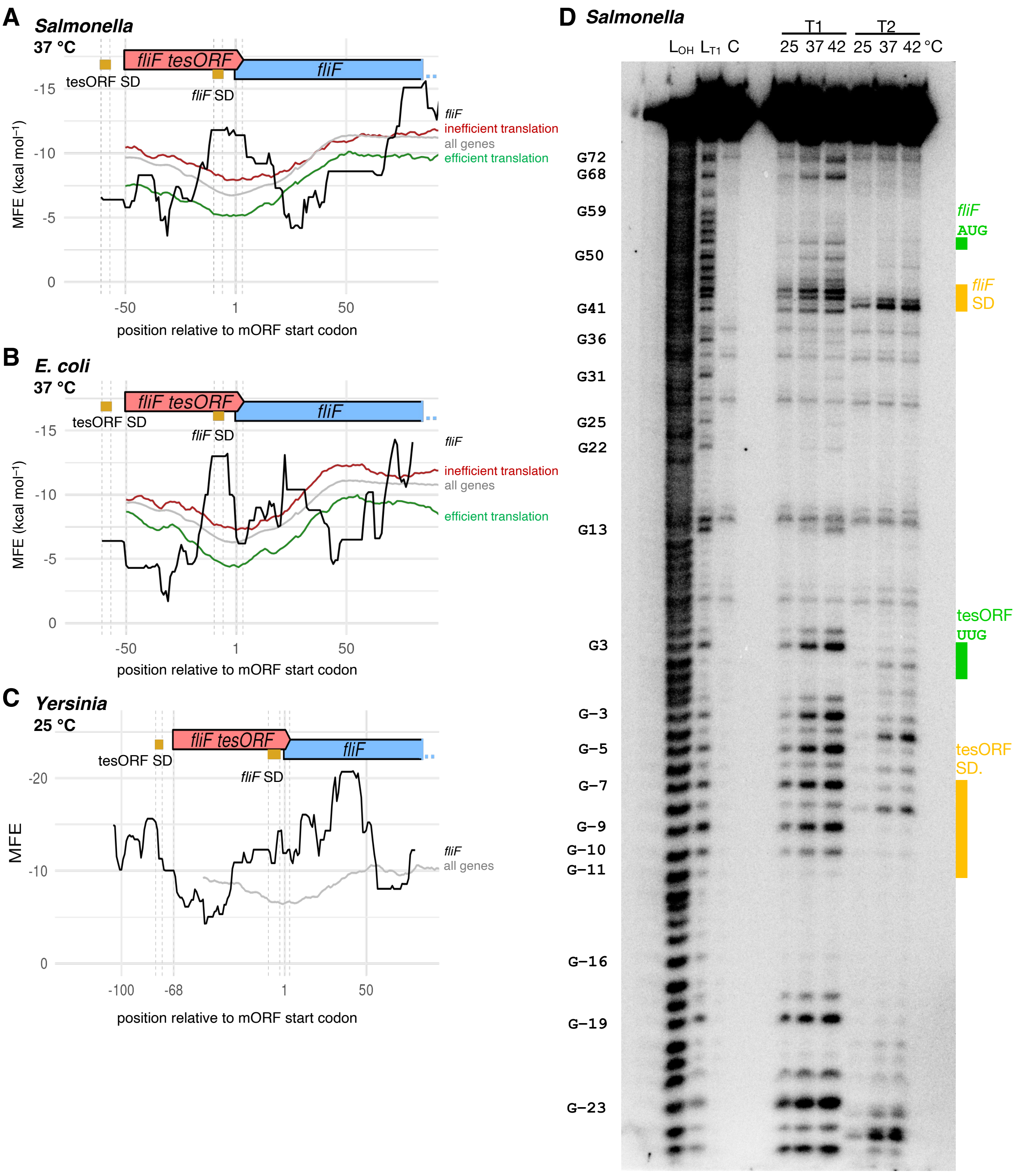
A
Salmonella
37 °C
fliF tesORF
fliF
tesORF SD
fliF SD
MFE (kcal mol⁻¹)
-15
-10
-5
0
-50
1
50
position relative to mORF start codon
fliF
inefficient translation
all genes
efficient translation
B
E. coli
37 °C
fliF tesORF
fliF
tesORF SD
fliF SD
MFE (kcal mol⁻¹)
-15
-10
-5
0
-50
1
50
position relative to mORF start codon
fliF
inefficient translation
all genes
efficient translation
C
Yersinia
25 °C
fliF tesORF
fliF
tesORF SD
fliF SD
MFE
-20
-10
0
-100
-68
1
50
position relative to mORF start codon
fliF
all genes
D
Salmonella
T1
T2
LOH LT1 C
25 37 42 25 37 42 °C
G72
G68
G59
G50
G41
G36
G31
G25
G22
G13
G3
G–3
G–5
G–7
G–9
G–10
G–11
G–16
G–19
G–23
fliF
AUG
fliF
SD
tesORF
UUG
tesORF
SD.

**Figure S6**

**A–C**, Minimum free energies (MFEs) of predicted local RNA secondary structures around *fliF*-tesORF in *Salmonella* (**A**), *E. coli* (**B**), and *Yersinia* (**C**). tesORF is labelled in red, the main ORF in blue, and the Shine–Dalgarno sequences in gold. For comparison, the graph shows the mean sliding-window MFE of predicted secondary structures surrounding the start codons of all *Salmonella* SL1344 or *E. coli* MG1655 genes (grey), and the 100 genes with the highest (green) or lowest (red) translational efficiency. The local MFE was predicted using a 51-nt sliding window moving in 1-nt increments and ViennaRNA RNAfold calculations at 37 °C (*Salmonella* and E*. coli*) or 25 °C (*Yersinia*). The mean profiles are aligned by their start codons to the *fliF* start codon.

**D**, 8% polyacrylamide gel showing partially digested 5′-end-labelled *Salmonella fliF* RNA, from a replicate of the RNA structure-probing experiment shown in figure 4. Limited digestion was performed at 25 °C, 37 °C, or 42 °C with 0.0016 U of RNase T1, which cleaves RNA 3′ of unpaired guanine, or 0.053 U of RNase T2, which cleaves RNA 3′ of unpaired bases, preferentially adenine. Nucleotide positions are numbered relative to the first nucleotide of tesORF. An alkaline hydrolysis ladder and a T1 digestion ladder generated from the same RNA are shown in lanes $L_{OH}$ and $L_{T1}$, respectively. The lane labelled C contains the enzyme-free control.

### Figure S7

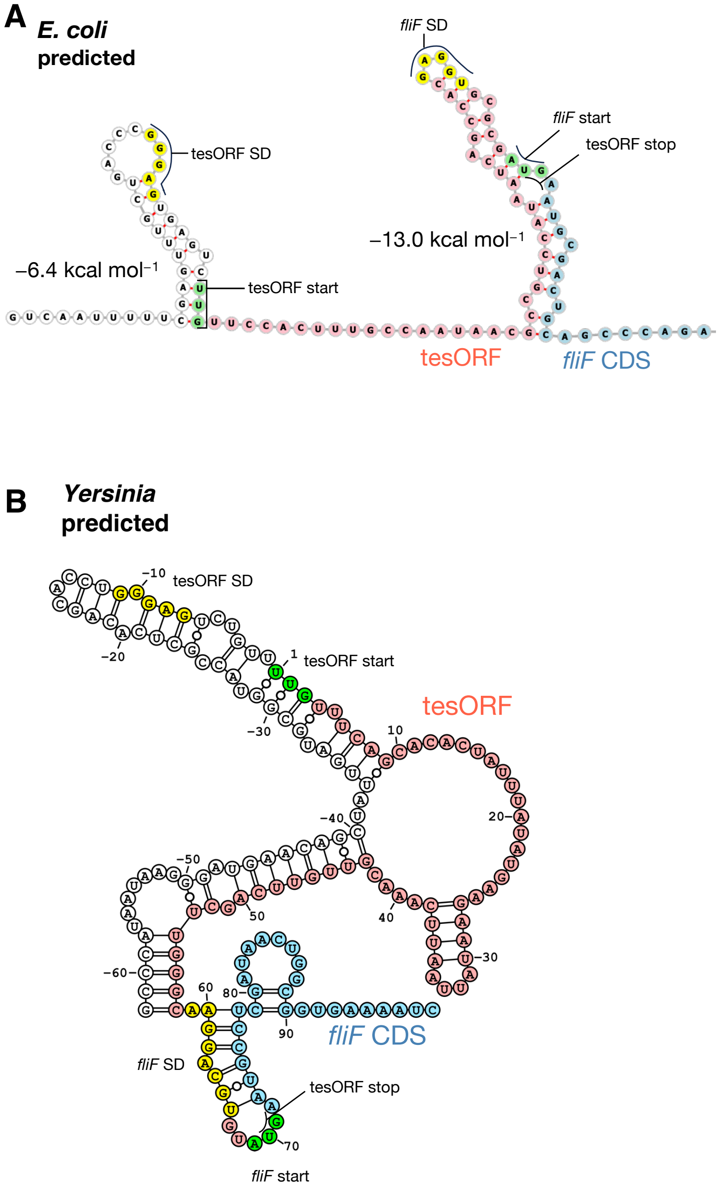

A
E. coli
predicted
tesORF SD
−6.4 kcal mol−1
tesORF start
fliF SD
fliF start
tesORF stop
−13.0 kcal mol−1
tesORF
fliF CDS
B
Yersinia
predicted
tesORF SD
tesORF start
tesORF
fliF CDS
fliF SD
tesORF stop
fliF start

**Figure S7**

Predicted secondary structures of *E. coli fliF* RNA (A) and *Yersinia pseudotuberculosis fliF* RNA (B). The *E. coli* structure was predicted using ViennaRNA RNAfold with a maximum base-pair span of 50 nt and coaxial stacking in multiloop helices allowed, and visualized with forna. The *Yersinia* structure was predicted using ViennaRNA RNAfold at 25 °C. The Shine–Dalgarno sequences are highlighted in yellow and start codon positions in green. Nucleotide positions are numbered relative to the first nucleotide of tesORF.

### Figure S8

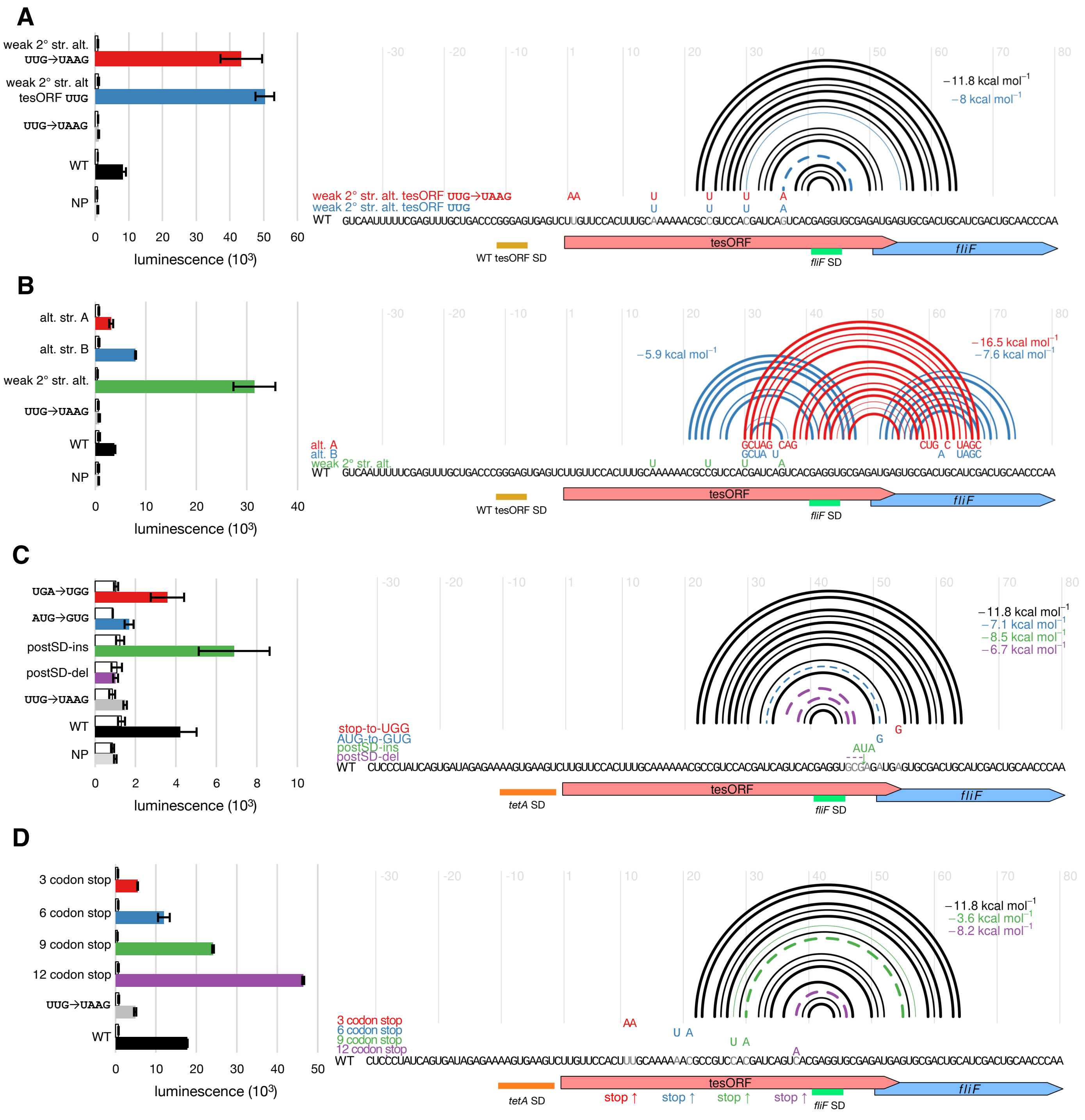

A
weak 2° str. alt. UUG→UAAG
weak 2° str. alt tesORF UUG
UUG→UAAG
WT
NP
luminescence (10³)
−11.8 kcal mol⁻¹
−8 kcal mol⁻¹
weak 2° str. alt. tesORF UUG→UAAG
weak 2° str. alt. tesORF UUG
WT GUCAAUUUUUCGAGUUUGCUGACCCGGGAGUGAGUCUUGUUCCACUUUGCAAAAAACGCCGUCCACGAUCAGUCACGAGGUGCGAGAUGAGUGCGACUGCAUCGACUGCAACCCAA
WT tesORF SD
tesORF
fliF
fliF SD
B
alt. str. A
alt. str. B
weak 2° str. alt.
UUG→UAAG
WT
NP
luminescence (10³)
−5.9 kcal mol⁻¹
−16.5 kcal mol⁻¹
−7.6 kcal mol⁻¹
alt. A
alt. B
weak 2° str. alt.
WT GUCAAUUUUUCGAGUUUGCUGACCCGGGAGUGAGUCUUGUUCCACUUUGCAAAAAACGCCGUCCACGAUCAGUCACGAGGUGCGAGAUGAGUGCGACUGCAUCGACUGCAACCCAA
WT tesORF SD
tesORF
fliF
fliF SD
C
UGA→UGG
AUG→GUG
postSD-ins
postSD-del
UUG→UAAG
WT
NP
luminescence (10³)
−11.8 kcal mol⁻¹
−7.1 kcal mol⁻¹
−8.5 kcal mol⁻¹
−6.7 kcal mol⁻¹
stop-to-UGG
AUG-to-GUG
postSD-ins
postSD-del
WT CUCCCUAUCAGUGAUAGAGAAAAGUGAAGUCUUGUUCCACUUUGCAAAAAACGCCGUCCACGAUCAGUCACGAGGUGCGAGAUGAGUGCGACUGCAUCGACUGCAACCCAA
tetA SD
tesORF
fliF
fliF SD
D
3 codon stop
6 codon stop
9 codon stop
12 codon stop
UUG→UAAG
WT
luminescence (10³)
−11.8 kcal mol⁻¹
−3.6 kcal mol⁻¹
−8.2 kcal mol⁻¹
3 codon stop
6 codon stop
9 codon stop
12 codon stop
WT CUCCCUAUCAGUGAUAGAGAAAAGUGAAGUCUUGUUCCACUUUGCAAAAAACGCCGUCCACGAUCAGUCACGAGGUGCGAGAUGAGUGCGACUGCAUCGACUGCAACCCAA
tetA SD
tesORF
fliF
stop ↑
stop ↑
stop ↑
stop ↑
fliF SD

**Figure S8**

Luciferase reporter signal from constructs containing *Salmonella fliF*-tesORF sequence variants (left), with the variants and their effects on the determined secondary structure illustrated on the right. White and coloured bars represent the signal 0 and 20 min after induction, respectively. The colours correspond to the sequences shown in the chart on the right. The error bars represent the standard error of the mean from three replicates. In the secondary structure diagrams, arcs illustrate GC (thick), AU (medium), and GU (thin) base pairs. Disrupted base pairs are shown as dashed arcs in the colour corresponding to the sequence variant. Sequence differences are highlighted in colour, and the corresponding wild-type nucleotides are shown in grey. A downward arrow represents an insertion, while dashes represent deletions. The free energy of the stem–loop structure was calculated with *RNAeval* from the ViennaRNA package. WT = wild-type *fliF* and *fliF*-tesORF sequence in the *fliF-luxAB* luciferase reporter construct. NP = luciferase reporter construct with no promoter.

**A**, Alternative weakly structured variants of the *fliF*-tesORF coding sequence with either a wild-type (WT) or a mutated start codon (UUG→UAAG). The variants contain four point mutations that destabilize the overall predicted secondary structure while preserving the amino acid sequence encoded by the tesORF.

**B**, Sequence variants that form alternative secondary structures. The displayed secondary structures on the right were predicted with ViennaRNA RNAfold and the MFE of the individual stem–loop structures was calculated with RNAeval.

**C**, Sequence variants affecting the *fliF*-tesORF stop codon, the *fliF* start codon, or the spacing between the *fliF* Shine–Dalgarno sequence and start codon. The post-SD ins and del variants respectively increase or decrease this spacing by three nucleotides. The AUG→GUG mutant contains a mutation of the fliF start codon, and the UGA→UGG mutant has a disrupted tesORF stop codon. These constructs contain a *tetA* 5′ UTR rather than the *fliF* 5′ UTR. The *tetA* Shine–Dalgarno sequence is labelled in orange.

**D**, Stop codon position variants, in which stop codons were introduced by sequence substitution, yielding tesORF sequences three, six, nine, and twelve codons in length. This construct has a *tetA* 5′ UTR rather than the *fliF* 5′ UTR. The *tetA* Shine–Dalgarno sequence is labelled in orange.

**Table S1: Normalized expression from wild-type *Salmonella* and the tesORF UUG→UAAG mutant.**

| | | translation | | RNA abundance | | translational efficiency | |
|---|---|---|---|---|---|---|---|
| **locus** | **gene** | **WT** | ***fliF* -tesORF mut.** | **WT** | ***fliF* -tesORF mut.** | **WT** | ***fliF* -tesORF mut.** |
| **SL1344_0001** | *thrL* | 146.965 | 82.095 | 394.672 | 789.286 | 0.372 | 0.104 |
| **SL1344_0002** | *thrA* | 50.783 | 20.796 | 55.152 | 47.288 | 0.921 | 0.44 |
| **SL1344_0003** | *thrB* | 39.908 | 16.237 | 69.063 | 68.273 | 0.578 | 0.238 |
| **SL1344_0004** | *thrC* | 77.638 | 60.199 | 114.126 | 120.011 | 0.68 | 0.502 |
| **SL1344_0005** | *yaaA* | 100.416 | 124.266 | 152.258 | 146.354 | 0.66 | 0.849 |
| **SL1344_0006** | *yaaJ* | 6.577 | 9.554 | 32.911 | 37.696 | 0.2 | 0.253 |
| **SL1344_0007** | *talB* | 2366.906 | 2858.533 | 1362.514 | 1525.572 | 1.74 | 1.87 |
| **SL1344_0008** | *mog* | 143.529 | 121.425 | 124.688 | 130.631 | 1.15 | 0.93 |
| **SL1344_0009** | *yaaH* | 71.722 | 14.516 | 79.396 | 17.74 | 0.903 | 0.818 |
| **SL1344_0012** | *dnaK* | 8493.265 | 1081.173 | 5777.315 | 847.621 | 1.47 | 1.28 |
| **SL1344_0013** | *dnaJ* | 672.998 | 33.116 | 669.11 | 122.591 | 1.01 | 0.27 |
| **SL1344_0028** | *bcfH* | 4.128 | 2.895 | 9.49 | 16.719 | 0.435 | 0.173 |
| **SL1344_0036** | *SL1344-0036* | 8.519 | 14.979 | 7.869 | 15.957 | 1.08 | 0.939 |
| **SL1344_0040** | *nhaA* | 242.661 | 153.412 | 187.225 | 179.197 | 1.3 | 0.856 |
| **SL1344_0041** | *nhaR* | 83.661 | 43.987 | 133.252 | 101.82 | 0.628 | 0.432 |
| **SL1344_0042** | *SL1344-0042* | 9.198 | 5.468 | 8.89 | 11.734 | 1.03 | 0.466 |
| **SL1344_0043** | *SL1344-0043* | 0.991 | 4.951 | 2.035 | 5.191 | 0.487 | 0.954 |
| **SL1344_0044** | *rpsT* | 2100.727 | 4124.562 | 9553.737 | 6378.539 | 0.22 | 0.647 |
| **SL1344_0046** | *ribF* | 129.61 | 80.626 | 419.49 | 391.445 | 0.309 | 0.206 |
| **SL1344_0047** | *ileS* | 526.882 | 233.264 | 428.028 | 288.73 | 1.23 | 0.808 |
| **SL1344_0048** | *lspA* | 262.889 | 166.591 | 603.229 | 512.244 | 0.436 | 0.325 |

| | | | | | | | |
|---|---|---|---|---|---|---|---|
| **SL1344_0049** | *fkpB* | 42.62 | 26.906 | 185.262 | 236.952 | 0.23 | 0.114 |
| **SL1344_0050** | *ispH* | 235.968 | 88.907 | 170.88 | 153.361 | 1.38 | 0.58 |
| **SL1344_0051** | *SL1344-0051* | 4.379 | 6.559 | 68.823 | 67.813 | 0.0636 | 0.0967 |
| **SL1344_0052** | *rihC* | 95.187 | 191.804 | 61.877 | 106.061 | 1.54 | 1.81 |
| **SL1344_0053** | *citB2* | 25.849 | 8.614 | 5.739 | 7.778 | 4.5 | 1.11 |
| **SL1344_0062** | *citF2* | 4.748 | 5.724 | 8.436 | 10.126 | 0.563 | 0.565 |
| **SL1344_0064** | *citG2* | 2.98 | 4.109 | 16.438 | 4.057 | 0.181 | 1.01 |
| **SL1344_0065** | *dapB* | 72.085 | 48.078 | 154.617 | 149.662 | 0.466 | 0.321 |
| **SL1344_0067** | *carA* | 20.762 | 18.056 | 20.276 | 20.307 | 1.02 | 0.889 |
| **SL1344_0068** | *carB* | 25.783 | 25.54 | 21.185 | 26.021 | 1.22 | 0.981 |
| **SL1344_0069** | *caiF* | 5.082 | 32.293 | 25.705 | 66.98 | 0.198 | 0.482 |
| **SL1344_0071** | *caiD* | 4.745 | 1.471 | 6.293 | 3.722 | 0.754 | 0.395 |
| **SL1344_0074** | *caiA* | 8.079 | 4.939 | 22.264 | 24.835 | 0.363 | 0.199 |
| **SL1344_0082** | *SL1344-0082* | 12.208 | 45.674 | 40.913 | 64.767 | 0.298 | 0.705 |
| **SL1344_0083** | *SL1344-0083* | 18.308 | 24.076 | 38.921 | 22.351 | 0.47 | 1.08 |
| **SL1344_0087** | *kefC* | 8.738 | 6.791 | 14.006 | 18.624 | 0.624 | 0.365 |
| **SL1344_0088** | *folA* | 206.317 | 176.855 | 317.797 | 228.843 | 0.649 | 0.773 |
| **SL1344_0089** | *apaH* | 109.185 | 58.247 | 93.467 | 94.551 | 1.17 | 0.616 |
| **SL1344_0090** | *apaG* | 10.649 | 35.09 | 41.721 | 61.79 | 0.255 | 0.568 |
| **SL1344_0091** | *ksgA* | 135.816 | 51.306 | 229.265 | 198.931 | 0.592 | 0.258 |
| **SL1344_0092** | *pdxAa* | 41.675 | 28.926 | 201.583 | 143.119 | 0.207 | 0.202 |
| **SL1344_0093** | *surA* | 288.567 | 250.68 | 372.955 | 369.277 | 0.774 | 0.679 |
| **SL1344_0094** | *imp* | 122.676 | 126.362 | 201.843 | 194.209 | 0.608 | 0.651 |
| **SL1344_0095** | *djlA* | 62.544 | 96.719 | 105.277 | 111.437 | 0.594 | 0.868 |

| | | | | | | | |
|---|---|---|---|---|---|---|---|
| **SL1344_0096** | *rluA* | 8.7 | 6.802 | 53.328 | 23.666 | 0.163 | 0.287 |
| **SL1344_0097** | *hepA* | 89.881 | 15.607 | 67.27 | 26.383 | 1.34 | 0.592 |
| **SL1344_0098** | *polB* | 3.7 | 3.181 | 24.321 | 24.631 | 0.152 | 0.129 |
| **SL1344_0100** | *SL1344-0100* | 6.483 | 9.07 | 9.899 | 10.075 | 0.655 | 0.9 |
| **SL1344_0102** | *araA* | 2.166 | 5.114 | 12.257 | 8.671 | 0.177 | 0.59 |
| **SL1344_0104** | *araC* | 43.941 | 69.951 | 184.547 | 166.994 | 0.238 | 0.419 |
| **SL1344_0105** | *yabI* | 13.257 | 15.145 | 170.529 | 175.611 | 0.0777 | 0.0862 |
| **SL1344_0106** | *thiQ* | 5.602 | 2.882 | 19.845 | 17.212 | 0.282 | 0.167 |
| **SL1344_0107** | *thiP* | 5.107 | 2.956 | 37.421 | 22.123 | 0.136 | 0.134 |
| **SL1344_0108** | *tbpA* | 76.038 | 34.011 | 88.147 | 78.114 | 0.863 | 0.435 |
| **SL1344_0109** | *yabN* | 8.67 | 8.241 | 14.734 | 13.918 | 0.588 | 0.592 |
| **SL1344_0110** | *leuD* | 16.41 | 12.908 | 25.314 | 17.037 | 0.648 | 0.758 |
| **SL1344_0111** | *leuC* | 32.703 | 19.129 | 25.276 | 12.616 | 1.29 | 1.52 |
| **SL1344_0112** | *leuB* | 60.983 | 23.11 | 25.273 | 16.761 | 2.41 | 1.38 |
| **SL1344_0113** | *leuA* | 21.313 | 11.683 | 22.846 | 23.333 | 0.933 | 0.501 |
| **SL1344_0114** | *leuL* | 351.084 | 1268.166 | 1719.582 | 1350.679 | 0.204 | 0.939 |
| **SL1344_0116** | *ilvI* | 46.729 | 38.144 | 29.545 | 26.521 | 1.58 | 1.44 |
| **SL1344_0117** | *ilvH* | 34.289 | 32.075 | 52.597 | 67.278 | 0.652 | 0.477 |
| **SL1344_0118** | *fruR* | 407.057 | 317.974 | 518.69 | 383.926 | 0.785 | 0.828 |
| **SL1344_0119** | *mraZ* | 53.778 | 64.613 | 1228.616 | 1328.835 | 0.0438 | 0.0486 |
| **SL1344_0120** | *mraW* | 112.231 | 137.631 | 389.085 | 459.988 | 0.288 | 0.299 |
| **SL1344_0121** | *ftsL* | 127.931 | 188.638 | 440.482 | 558.557 | 0.29 | 0.338 |
| **SL1344_0122** | *ftsI* | 38.692 | 29.449 | 192.454 | 157.511 | 0.201 | 0.187 |
| **SL1344_0123** | *murE* | 119.861 | 84.661 | 246.412 | 195.9 | 0.486 | 0.432 |

| | | | | | | | |
|---|---|---|---|---|---|---|---|
| **SL1344_0124** | *murF* | 67.993 | 39.091 | 188.308 | 138.704 | 0.361 | 0.282 |
| **SL1344_0125** | *mraY* | 172.227 | 82.656 | 373.708 | 214.35 | 0.461 | 0.386 |
| **SL1344_0126** | *murD* | 106.478 | 59.809 | 227.08 | 183.186 | 0.469 | 0.326 |
| **SL1344_0127** | *ftsW* | 48.562 | 42.528 | 154.386 | 110.016 | 0.315 | 0.387 |
| **SL1344_0128** | *murG* | 92.283 | 37.386 | 185.115 | 161.871 | 0.499 | 0.231 |
| **SL1344_0129** | *murC* | 79.688 | 55.901 | 138.656 | 125.317 | 0.575 | 0.446 |
| **SL1344_0130** | *ddlB* | 49.618 | 48.966 | 308.527 | 244.072 | 0.161 | 0.201 |
| **SL1344_0131** | *ftsQ* | 88.115 | 53.206 | 371.875 | 273.532 | 0.237 | 0.195 |
| **SL1344_0132** | *ftsA* | 516.392 | 342.907 | 611.382 | 521.06 | 0.845 | 0.658 |
| **SL1344_0133** | *ftsZ* | 1109.74 | 1196.985 | 768.172 | 781.887 | 1.44 | 1.53 |
| **SL1344_0134** | *lpxC* | 1503.46 | 921.621 | 2783.634 | 2506.986 | 0.54 | 0.368 |
| **SL1344_0135** | *secM* | 120.291 | 89.875 | 230.453 | 261.306 | 0.522 | 0.344 |
| **SL1344_0136** | *secA* | 484.821 | 383.393 | 170.917 | 260.526 | 2.84 | 1.47 |
| **SL1344_0137** | *mutT* | 44.695 | 24.048 | 71.865 | 57.394 | 0.622 | 0.419 |
| **SL1344_0138** | *yacG* | 405.728 | 1501.428 | 184.063 | 192.423 | 2.2 | 7.8 |
| **SL1344_0139** | *yacF* | 77.414 | 83.198 | 109.549 | 108.057 | 0.707 | 0.77 |
| **SL1344_0140** | *coaE* | 21.447 | 42.5 | 153.987 | 126.111 | 0.139 | 0.337 |
| **SL1344_0141** | *guaC* | 986.289 | 918.026 | 645.907 | 510.465 | 1.53 | 1.8 |
| **SL1344_0145** | *nadC* | 132.625 | 57.445 | 126.208 | 78.324 | 1.05 | 0.733 |
| **SL1344_0146** | *ampD* | 73.048 | 70.554 | 125.319 | 91.357 | 0.583 | 0.772 |
| **SL1344_0147** | *ampE* | 99.35 | 140.498 | 69.923 | 84.583 | 1.42 | 1.66 |
| **SL1344_0148** | *SL1344-0148* | 1.805 | 3.862 | 4.447 | 7.526 | 0.406 | 0.513 |
| **SL1344_0150** | *aroP* | 24.47 | 28.515 | 45.546 | 39.612 | 0.537 | 0.72 |
| **SL1344_0151** | *pdhR* | 12.226 | 17.694 | 68.404 | 132.02 | 0.179 | 0.134 |

| | | | | | | | |
|---|---|---|---|---|---|---|---|
| **SL1344_0152** | *aceE* | 736.352 | 461.237 | 462.445 | 358.553 | 1.59 | 1.29 |
| **SL1344_0153** | *aceF* | 707.858 | 397.331 | 499.737 | 375.143 | 1.42 | 1.06 |
| **SL1344_0154** | *lpdA* | 3390.695 | 4804.497 | 2922.879 | 3473.539 | 1.16 | 1.38 |
| **SL1344_0159** | *acnB* | 1457.895 | 1789.079 | 1057.319 | 1408.319 | 1.38 | 1.27 |
| **SL1344_0160** | *SL1344-0160* | 36.092 | 157.748 | 22.449 | 42.076 | 1.61 | 3.75 |
| **SL1344_0161** | *yacL* | 141.871 | 323.804 | 465.45 | 518.146 | 0.305 | 0.625 |
| **SL1344_0162** | *kdgT* | 830.148 | 23.886 | 806.177 | 46.457 | 1.03 | 0.514 |
| **SL1344_0163** | *dtnK* | 1062.101 | 35.829 | 798.494 | 43.377 | 1.33 | 0.826 |
| **SL1344_0164** | *pdxAb* | 6772.822 | 163.14 | 1215.707 | 91.653 | 5.57 | 1.78 |
| **SL1344_0165** | *SL1344-0165* | 224.754 | 86.444 | 1400.443 | 343.73 | 0.16 | 0.251 |
| **SL1344_0166** | *speD* | 324.936 | 158.46 | 239.396 | 108.272 | 1.36 | 1.46 |
| **SL1344_0167** | *speE* | 286.278 | 159.269 | 334.523 | 129.88 | 0.856 | 1.23 |
| **SL1344_0168** | *yacC* | 82.158 | 59.616 | 179.827 | 165.708 | 0.457 | 0.36 |
| **SL1344_0169** | *cueO* | 28.146 | 24.489 | 27.054 | 39.232 | 1.04 | 0.624 |
| **SL1344_0170** | *gcd* | 27.703 | 28.335 | 26.113 | 43.18 | 1.06 | 0.656 |
| **SL1344_0171** | *hpt* | 881.743 | 372.534 | 569.998 | 357.685 | 1.55 | 1.04 |
| **SL1344_0172** | *yadF* | 686.048 | 879.044 | 658.018 | 787.581 | 1.04 | 1.12 |
| **SL1344_0173** | *yadG* | 68.964 | 76.9 | 46.784 | 53.392 | 1.47 | 1.44 |
| **SL1344_0174** | *yadH* | 77.621 | 33.261 | 63.223 | 34.026 | 1.23 | 0.978 |
| **SL1344_0178** | *stiA* | 2.302 | 6.424 | 3.319 | 4.612 | 0.694 | 1.39 |
| **SL1344_0180** | *yadE* | 11.213 | 6.747 | 21.155 | 21.289 | 0.53 | 0.317 |
| **SL1344_0181** | *panD* | 299.235 | 302.436 | 146.377 | 234.617 | 2.04 | 1.29 |
| **SL1344_0182** | *panC* | 174.675 | 85.922 | 137.247 | 79.182 | 1.27 | 1.09 |
| **SL1344_0183** | *panB* | 56.504 | 26.796 | 61.275 | 51.136 | 0.922 | 0.524 |

| | | | | | | | |
|---|---|---|---|---|---|---|---|
| **SL1344_0184** | *folK* | 8.263 | 20.406 | 49.133 | 34.354 | 0.168 | 0.594 |
| **SL1344_0185** | *pcnB* | 107.626 | 58.29 | 217.425 | 174.227 | 0.495 | 0.335 |
| **SL1344_0186** | *yadB* | 32.99 | 12.564 | 83.859 | 68.712 | 0.393 | 0.183 |
| **SL1344_0187** | *dksA* | 2144.368 | 3148.817 | 3636.055 | 3478.676 | 0.59 | 0.905 |
| **SL1344_0188** | *sfsA* | 29.471 | 55.285 | 47.077 | 143.53 | 0.626 | 0.385 |
| **SL1344_0189** | *ligT* | 5.908 | 8.455 | 45.494 | 51.363 | 0.13 | 0.165 |
| **SL1344_0190** | *hrpB* | 11.815 | 8.905 | 65.312 | 70.583 | 0.181 | 0.126 |
| **SL1344_0191** | *mrcB* | 81.178 | 67.401 | 166.864 | 164.086 | 0.486 | 0.411 |
| **SL1344_0192** | *fhuA* | 66.546 | 2.858 | 55.579 | 8.015 | 1.2 | 0.357 |
| **SL1344_0193** | *fhuC* | 10.014 | 0.085 | 15.451 | 4.454 | 0.648 | 0.0191 |
| **SL1344_0194** | *fhuD* | 4.119 | 0.153 | 18.746 | 8.494 | 0.22 | 0.018 |
| **SL1344_0195** | *fhuB* | 4.487 | 0.76 | 21.665 | 12.947 | 0.207 | 0.0587 |
| **SL1344_0203** | *hemL* | 385.156 | 370.32 | 334.747 | 281.18 | 1.15 | 1.32 |
| **SL1344_0204** | *clcA* | 18.857 | 8.802 | 55.552 | 97.276 | 0.339 | 0.0905 |
| **SL1344_0205** | *yadR* | 1404.022 | 1620.483 | 535.653 | 580.642 | 2.62 | 2.79 |
| **SL1344_0206** | *yadS* | 38.514 | 80.23 | 127.859 | 93.189 | 0.301 | 0.861 |
| **SL1344_0207** | *btuF* | 9.385 | 4.925 | 34.723 | 28.586 | 0.27 | 0.172 |
| **SL1344_0208** | *mtn* | 445.863 | 250.579 | 331.139 | 206.54 | 1.35 | 1.21 |
| **SL1344_0209** | *dgt* | 21.587 | 24.598 | 40.719 | 59.319 | 0.53 | 0.415 |
| **SL1344_0210** | *htrA* | 220.984 | 227.452 | 122.384 | 183.644 | 1.81 | 1.24 |
| **SL1344_0211** | *cdaR* | 42.121 | 75.07 | 85.117 | 166.37 | 0.495 | 0.451 |
| **SL1344_0212** | *yaeH* | 1648.418 | 6176.225 | 1538.882 | 2121.426 | 1.07 | 2.91 |
| **SL1344_0213** | *SL1344-0213* | 9.693 | 13.193 | 35.842 | 42.16 | 0.27 | 0.313 |
| **SL1344_0214** | *dapD* | 525.359 | 688.128 | 532.022 | 418.572 | 0.987 | 1.64 |

| | | | | | | | |
|---|---|---|---|---|---|---|---|
| **SL1344_0215** | *glnD* | 19.82 | 16.846 | 34.032 | 31.361 | 0.582 | 0.537 |
| **SL1344_0216** | *map* | 296.94 | 310.245 | 319.195 | 309.398 | 0.93 | 1 |
| **SL1344_0217** | *rpsB* | 7379.809 | 3638.294 | 7077.618 | 4842.493 | 1.04 | 0.751 |
| **SL1344_0218** | *tsf* | 4463.686 | 3947.468 | 5091.974 | 3244.191 | 0.877 | 1.22 |
| **SL1344_0219** | *pyrH* | 427.814 | 403.818 | 500.205 | 502.127 | 0.855 | 0.804 |
| **SL1344_0220** | *frr* | 888.87 | 1522.311 | 869.838 | 1411.948 | 1.02 | 1.08 |
| **SL1344_0221** | *dxr* | 28.039 | 17.503 | 93.482 | 72.476 | 0.3 | 0.241 |
| **SL1344_0222** | *uppS* | 61.456 | 76.625 | 341.04 | 452.644 | 0.18 | 0.169 |
| **SL1344_0223** | *cdsA* | 198.97 | 113.448 | 579.15 | 425.891 | 0.344 | 0.266 |
| **SL1344_0224** | *yaeL* | 77.788 | 149.164 | 307.079 | 415.222 | 0.253 | 0.359 |
| **SL1344_0225** | *yaeT* | 876.542 | 374.356 | 540.194 | 419.15 | 1.62 | 0.893 |
| **SL1344_0226** | *hlpA* | 3066.293 | 11221.268 | 1474.086 | 1579.435 | 2.08 | 7.1 |
| **SL1344_0227** | *lpxD* | 420.303 | 119.534 | 876.622 | 787.768 | 0.479 | 0.152 |
| **SL1344_0228** | *fabZ* | 840.404 | 917.392 | 1872.576 | 963.919 | 0.449 | 0.952 |
| **SL1344_0229** | *lpxA* | 209.243 | 117.076 | 851.844 | 754.11 | 0.246 | 0.155 |
| **SL1344_0230** | *lpxB* | 28.18 | 9.294 | 93.644 | 95.032 | 0.301 | 0.0978 |
| **SL1344_0231** | *rnhB* | 56.021 | 32.472 | 116.951 | 102.061 | 0.479 | 0.318 |
| **SL1344_0232** | *dnaE* | 40.874 | 18.475 | 216.553 | 155.41 | 0.189 | 0.119 |
| **SL1344_0233** | *accA* | 339.258 | 369.44 | 791.357 | 622.025 | 0.429 | 0.594 |
| **SL1344_0234** | *SL1344-0234* | 14.899 | 3.085 | 25.479 | 27.194 | 0.585 | 0.113 |
| **SL1344_0235** | *ldcC* | 31.891 | 29.057 | 41.899 | 43.357 | 0.761 | 0.67 |
| **SL1344_0236** | *yaeR* | 52.212 | 52.324 | 89.696 | 88.346 | 0.582 | 0.592 |
| **SL1344_0237** | *mesJ* | 19.39 | 16.276 | 46.843 | 56.856 | 0.414 | 0.286 |
| **SL1344_0238** | *rof* | 87.283 | 723.96 | 316.813 | 540.847 | 0.276 | 1.34 |

| | | | | | | | |
|---|---|---|---|---|---|---|---|
| **SL1344_0239** | *yaeP* | 434.092 | 815.944 | 463.488 | 778.801 | 0.937 | 1.05 |
| **SL1344_0240** | *yaeQ* | 73.071 | 76.742 | 93.871 | 116.023 | 0.778 | 0.661 |
| **SL1344_0242** | *cutF* | 89.635 | 172.96 | 91.187 | 135.222 | 0.983 | 1.28 |
| **SL1344_0243** | *proS* | 279.753 | 180.203 | 246.702 | 216.737 | 1.13 | 0.831 |
| **SL1344_0244** | *yaeB* | 9.949 | 14.123 | 61.356 | 72.878 | 0.162 | 0.194 |
| **SL1344_0245** | *rcsF* | 78.049 | 159.724 | 126.552 | 171.103 | 0.617 | 0.933 |
| **SL1344_0246** | *metQ* | 735.142 | 553.59 | 411.39 | 245.403 | 1.79 | 2.26 |
| **SL1344_0247** | *yaeE* | 41.273 | 20.074 | 113.444 | 45.955 | 0.364 | 0.437 |
| **SL1344_0248** | *abc* | 69.634 | 16.214 | 57.444 | 25.538 | 1.21 | 0.635 |
| **SL1344_0249** | *yaeD* | 122.878 | 65.502 | 216.062 | 150.83 | 0.569 | 0.434 |
| **SL1344_0250** | *dkgB* | 50.948 | 64.806 | 64.64 | 51.726 | 0.788 | 1.25 |
| **SL1344_0251** | *yafC* | 25.489 | 20.89 | 23.424 | 21.14 | 1.09 | 0.988 |
| **SL1344_0252** | *SL1344-0252* | 1.812 | 3.586 | 8.595 | 10.383 | 0.211 | 0.345 |
| **SL1344_0253** | *yafD* | 29.673 | 32.703 | 92.178 | 84.037 | 0.322 | 0.389 |
| **SL1344_0254** | *yafE* | 18.349 | 6.088 | 21.012 | 20.51 | 0.873 | 0.297 |
| **SL1344_0255** | *dniR* | 110.557 | 75.828 | 480.353 | 468.2 | 0.23 | 0.162 |
| **SL1344_0256** | *gloB* | 30.537 | 88.536 | 32.049 | 44.904 | 0.953 | 1.97 |
| **SL1344_0257** | *yafS* | 5.24 | 5.927 | 19.433 | 23.242 | 0.27 | 0.255 |
| **SL1344_0258** | *rnhA* | 45.787 | 26.889 | 92.668 | 67.835 | 0.494 | 0.396 |
| **SL1344_0259** | *dnaQ* | 67.2 | 77.407 | 75.307 | 111.48 | 0.892 | 0.694 |
| **SL1344_0260** | *sciA* | 2.466 | 4.316 | 3.666 | 4.053 | 0.673 | 1.06 |
| **SL1344_0264** | *sciE* | 2.798 | 4.205 | 8.515 | 14.8 | 0.329 | 0.284 |
| **SL1344_0266** | *sciG* | 1.457 | 1.031 | 4.182 | 5.669 | 0.349 | 0.182 |
| **SL1344_0267** | *sciH* | 7.74 | 14.657 | 8.053 | 7.881 | 0.961 | 1.86 |

| | | | | | | | |
|---|---|---|---|---|---|---|---|
| **SL1344_0268** | *sciI* | 10.341 | 6.146 | 4.174 | 4.386 | 2.48 | 1.4 |
| **SL1344_0282** | *sciU* | 7.24 | 9.95 | 13.924 | 25.107 | 0.52 | 0.396 |
| **SL1344_0284** | *vgrS* | 1.865 | 1.056 | 3.928 | 4.14 | 0.475 | 0.255 |
| **SL1344_0286** | *SL1344-0286* | 2.448 | 14.763 | 16.732 | 20.273 | 0.146 | 0.728 |
| **SL1344_0286A** | *SL1344-0286A* | 87.895 | 160.52 | 79.015 | 100.196 | 1.11 | 1.6 |
| **SL1344_0288** | *sciX* | 10.594 | 44.587 | 35.601 | 49.169 | 0.298 | 0.907 |
| **SL1344_0300** | *sinR* | 8.867 | 13.776 | 16.938 | 24.765 | 0.523 | 0.556 |
| **SL1344_0302** | *pagN* | 7.975 | 55.362 | 12.146 | 41.138 | 0.657 | 1.35 |
| **SL1344_0303** | *sciZ* | 30.903 | 57.351 | 31.449 | 25.127 | 0.983 | 2.28 |
| **SL1344_0304** | *yafV* | 60.119 | 99.729 | 97.258 | 101.847 | 0.618 | 0.979 |
| **SL1344_0305** | *fadE* | 147.175 | 325.667 | 71.947 | 225.966 | 2.05 | 1.44 |
| **SL1344_0306** | *gmhA* | 652.263 | 537.826 | 361.389 | 420.018 | 1.8 | 1.28 |
| **SL1344_0307** | *yafJ* | 49.483 | 42.69 | 127.406 | 134.117 | 0.388 | 0.318 |
| **SL1344_0308** | *yafK* | 76.13 | 148.343 | 69.846 | 62.617 | 1.09 | 2.37 |
| **SL1344_0309** | *dbh* | 6.222 | 3.156 | 56.071 | 30.636 | 0.111 | 0.103 |
| **SL1344_0310** | *rtcB2* | 2.879 | 9.304 | 132.981 | 126.382 | 0.0216 | 0.0736 |
| **SL1344_0312** | *pepD* | 318.22 | 457.954 | 343.914 | 404.506 | 0.925 | 1.13 |
| **SL1344_0313** | *gpt* | 272.887 | 201.1 | 328.901 | 281.77 | 0.83 | 0.714 |
| **SL1344_0314** | *frsA* | 55.628 | 62.503 | 91.087 | 102.619 | 0.611 | 0.609 |
| **SL1344_0315** | *crl* | 632.585 | 1294.773 | 730.21 | 774.865 | 0.866 | 1.67 |
| **SL1344_0316** | *phoE* | 3.373 | 2.39 | 10.007 | 11.389 | 0.337 | 0.21 |
| **SL1344_0317** | *proB* | 148.361 | 119.592 | 328.355 | 238.525 | 0.452 | 0.501 |
| **SL1344_0318** | *proA* | 189.602 | 139.359 | 167.262 | 144.976 | 1.13 | 0.961 |
| **SL1344_0322** | *SL1344-0322* | 342.377 | 1033.965 | 424.792 | 1214.016 | 0.806 | 0.852 |

| | | | | | | | |
|---|---|---|---|---|---|---|---|
| **SL1344_0324** | *leuC2* | 9.646 | 5.542 | 4.661 | 4.497 | 2.07 | 1.23 |
| **SL1344_0327** | *SL1344-0327* | 3.882 | 3.122 | 8.539 | 6.765 | 0.455 | 0.462 |
| **SL1344_0328** | *SL1344-0328* | 17.893 | 12.393 | 20.778 | 21.225 | 0.861 | 0.584 |
| **SL1344_0336** | *SL1344-0336* | 3.714 | 11.292 | 18.272 | 22.156 | 0.203 | 0.51 |
| **SL1344_0337** | *SL1344-0337* | 3.19 | 9.368 | 6.868 | 9.363 | 0.464 | 1 |
| **SL1344_0338** | *SL1344-0338* | 2.856 | 9.458 | 14.564 | 18.637 | 0.196 | 0.507 |
| **SL1344_0339** | *SL1344-0339* | 14.958 | 25.782 | 35.36 | 46.375 | 0.423 | 0.556 |
| **SL1344_0340** | *SL1344-0340* | 11.621 | 40.201 | 7.036 | 13.202 | 1.65 | 3.05 |
| **SL1344_0344** | *SL1344-0344* | 19.074 | 15.305 | 10.354 | 5.843 | 1.84 | 2.62 |
| **SL1344_0348** | *SL1344-0348* | 2.974 | 3.685 | 12.401 | 7.838 | 0.24 | 0.47 |
| **SL1344_0349** | *SL1344-0349* | 5.474 | 19.309 | 11.562 | 14.351 | 0.473 | 1.35 |
| **SL1344_0350** | *SL1344-0350* | 16.999 | 17.441 | 23.895 | 21.823 | 0.711 | 0.799 |
| **SL1344_0351** | *SL1344-0351* | 78.265 | 132.083 | 84.717 | 111.87 | 0.924 | 1.18 |
| **SL1344_0352** | *mod* | 55.085 | 45.348 | 52.839 | 48.901 | 1.04 | 0.927 |
| **SL1344_0353** | *res* | 57.404 | 31.154 | 61.639 | 35.09 | 0.931 | 0.888 |
| **SL1344_0355** | *SL1344-0355* | 4.68 | 2.81 | 13.122 | 12.158 | 0.357 | 0.231 |
| **SL1344_0360** | *yahN* | 20.573 | 27.294 | 75.421 | 73.606 | 0.273 | 0.371 |
| **SL1344_0361** | *yahO* | 72.921 | 169.314 | 51.685 | 105.038 | 1.41 | 1.61 |
| **SL1344_0362** | *prpR* | 39.026 | 24.138 | 147.776 | 135.318 | 0.264 | 0.178 |
| **SL1344_0363** | *prpB* | 5.599 | 9.881 | 6.7 | 8.468 | 0.836 | 1.17 |
| **SL1344_0364** | *prpC* | 4.149 | 6.918 | 4.289 | 4.588 | 0.967 | 1.51 |
| **SL1344_0365** | *prpD* | 7.257 | 3.701 | 4.937 | 4.912 | 1.47 | 0.753 |
| **SL1344_0366** | *prpE* | 1.819 | 2.848 | 5.546 | 10.084 | 0.328 | 0.282 |
| **SL1344_0367** | *hemB* | 153.541 | 171.344 | 153.587 | 133.542 | 1 | 1.28 |

| | | | | | | | |
|---|---|---|---|---|---|---|---|
| **SL1344_0368** | *yaiU* | 0.967 | 1.876 | 35.805 | 52.671 | 0.027 | 0.0356 |
| **SL1344_0370** | *ampH* | 45.534 | 43.724 | 41.641 | 34.376 | 1.09 | 1.27 |
| **SL1344_0371** | *sbmA* | 53.375 | 18.723 | 52.081 | 43.323 | 1.02 | 0.432 |
| **SL1344_0372** | *yaiW* | 6.974 | 4.597 | 25.4 | 19.564 | 0.275 | 0.235 |
| **SL1344_0373** | *yaiY* | 22.797 | 51.511 | 54.285 | 52.114 | 0.42 | 0.988 |
| **SL1344_0374** | *yaiZ* | 747.329 | 348.294 | 818.387 | 302.235 | 0.913 | 1.15 |
| **SL1344_0375** | *ddlA* | 476.798 | 214.624 | 231.81 | 131.25 | 2.06 | 1.64 |
| **SL1344_0377** | *SL1344-0377* | 3.321 | 0.561 | 5.678 | 4.908 | 0.585 | 0.114 |
| **SL1344_0378** | *yaiB* | 321.246 | 681.996 | 173.974 | 507.031 | 1.85 | 1.35 |
| **SL1344_0379** | *psiF* | 17.887 | 54.247 | 10.942 | 49.563 | 1.63 | 1.09 |
| **SL1344_0381** | *proC* | 292.683 | 317.097 | 150.801 | 146.267 | 1.94 | 2.17 |
| **SL1344_0382** | *yaiI* | 18.433 | 3.729 | 86.303 | 64 | 0.214 | 0.0583 |
| **SL1344_0383** | *aroL* | 48.136 | 44.351 | 84.419 | 91.003 | 0.57 | 0.487 |
| **SL1344_0384** | *yaiA* | 12.024 | 66.604 | 92.591 | 134.117 | 0.13 | 0.497 |
| **SL1344_0385** | *aroM* | 60.941 | 70.63 | 224.143 | 151.349 | 0.272 | 0.467 |
| **SL1344_0386** | *yaiE* | 178.206 | 235.568 | 237.939 | 275.978 | 0.749 | 0.854 |
| **SL1344_0387** | *rdgC* | 74.058 | 73.615 | 120.652 | 109.733 | 0.614 | 0.671 |
| **SL1344_0388** | *yajF* | 124.771 | 121.6 | 71.132 | 96.449 | 1.75 | 1.26 |
| **SL1344_0390** | *sbcC* | 14.021 | 8.099 | 19.307 | 16.55 | 0.726 | 0.489 |
| **SL1344_0391** | *sbcD* | 14.959 | 13.796 | 33.488 | 18.129 | 0.447 | 0.761 |
| **SL1344_0392** | *phoB* | 46.927 | 120.96 | 326.939 | 535.368 | 0.144 | 0.226 |
| **SL1344_0393** | *phoR* | 7.719 | 2.782 | 39.492 | 46.343 | 0.195 | 0.06 |
| **SL1344_0394** | *brnQ* | 163.906 | 100.125 | 198.216 | 149.872 | 0.827 | 0.668 |
| **SL1344_0395** | *proY* | 24.913 | 11.759 | 76.442 | 32.344 | 0.326 | 0.364 |

| | | | | | | | |
|---|---|---|---|---|---|---|---|
| **SL1344_0396** | *malZ* | 2.8 | 3.293 | 14.116 | 15.601 | 0.198 | 0.211 |
| **SL1344_0397** | *tsaA* | 1673.542 | 2428.348 | 899.908 | 1337.935 | 1.86 | 1.81 |
| **SL1344_0398** | *yajB* | 15.256 | 14.025 | 39.29 | 106.271 | 0.388 | 0.132 |
| **SL1344_0399** | *queA* | 110.663 | 19.033 | 99.213 | 49.606 | 1.12 | 0.384 |
| **SL1344_0400** | *tgt* | 164.464 | 33.046 | 276.694 | 186.078 | 0.594 | 0.178 |
| **SL1344_0401** | *yajC* | 2494.34 | 3093.435 | 1729.878 | 2070.666 | 1.44 | 1.49 |
| **SL1344_0402** | *secD* | 185.335 | 116.203 | 449.419 | 329.999 | 0.412 | 0.352 |
| **SL1344_0403** | *secF* | 234.768 | 157.457 | 882.18 | 618.227 | 0.266 | 0.255 |
| **SL1344_0404** | *SL1344-0404* | 5.337 | 9.85 | 22.132 | 21.47 | 0.241 | 0.459 |
| **SL1344_0406** | *yajD* | 25.855 | 101.641 | 85.279 | 133.192 | 0.303 | 0.763 |
| **SL1344_0407** | *tsx* | 9996.928 | 960.328 | 5766.877 | 772.071 | 1.73 | 1.24 |
| **SL1344_0408** | *yajI* | 4.166 | 19.525 | 12.346 | 24.806 | 0.337 | 0.787 |
| **SL1344_0409** | *nrdR* | 44.725 | 131.81 | 208.997 | 276.122 | 0.214 | 0.477 |
| **SL1344_0410** | *ribD* | 61.393 | 34.75 | 133.628 | 150.461 | 0.459 | 0.231 |
| **SL1344_0411** | *ribH* | 394.126 | 808.023 | 304.084 | 378.237 | 1.3 | 2.14 |
| **SL1344_0412** | *nusB* | 116.416 | 86.808 | 477.72 | 618.633 | 0.244 | 0.14 |
| **SL1344_0413** | *thiL* | 37.647 | 29.42 | 89.348 | 58.692 | 0.421 | 0.501 |
| **SL1344_0414** | *pgpA* | 31.777 | 17.533 | 119.889 | 57.261 | 0.265 | 0.306 |
| **SL1344_0415** | *yajO* | 27.867 | 51.208 | 35.511 | 52.375 | 0.785 | 0.978 |
| **SL1344_0416** | *dxs* | 170.561 | 76.821 | 121.281 | 87.02 | 1.41 | 0.883 |
| **SL1344_0417** | *ispA* | 81.491 | 66.812 | 241.494 | 217.583 | 0.337 | 0.307 |
| **SL1344_0418** | *xseB* | 128.863 | 182.79 | 348.683 | 478.701 | 0.37 | 0.382 |
| **SL1344_0419** | *thiI* | 82.725 | 33.987 | 104.631 | 66.646 | 0.791 | 0.51 |
| **SL1344_0424** | *phnR* | 4.933 | 6.708 | 33.452 | 29.854 | 0.147 | 0.225 |

| | | | | | | | |
|---|---|---|---|---|---|---|---|
| **SL1344_0427** | *thiJ* | 16.426 | 20.257 | 46.091 | 49.79 | 0.356 | 0.407 |
| **SL1344_0428** | *apbA* | 41.8 | 62.129 | 63.509 | 74.79 | 0.658 | 0.831 |
| **SL1344_0429** | *yajQ* | 596.122 | 735.83 | 608.186 | 512.07 | 0.98 | 1.44 |
| **SL1344_0430** | *yajR* | 40.113 | 26.76 | 69.268 | 62.05 | 0.579 | 0.431 |
| **SL1344_0431** | *SL1344-0431* | 1.997 | 3.419 | 6.922 | 10.762 | 0.288 | 0.318 |
| **SL1344_0432** | *SL1344-0432* | 6.178 | 3.39 | 8.542 | 7.909 | 0.723 | 0.429 |
| **SL1344_0433** | *cyoE* | 758.033 | 480.806 | 1451.049 | 1962.121 | 0.522 | 0.245 |
| **SL1344_0434** | *cyoD* | 1562.731 | 1388.664 | 1754.293 | 2359.853 | 0.891 | 0.588 |
| **SL1344_0435** | *cyoC* | 4634.775 | 3845.247 | 3157.458 | 3191.526 | 1.47 | 1.2 |
| **SL1344_0436** | *cyoB* | 3004.59 | 1884.585 | 2341.807 | 1886.919 | 1.28 | 0.999 |
| **SL1344_0437** | *cyoA* | 4828.454 | 3952.625 | 4319.067 | 4269.849 | 1.12 | 0.926 |
| **SL1344_0438** | *ampG* | 18.849 | 8.618 | 71.441 | 38.463 | 0.264 | 0.224 |
| **SL1344_0439** | *yajG* | 320.753 | 342.255 | 194.667 | 152.216 | 1.65 | 2.25 |
| **SL1344_0440** | *bolA* | 137.934 | 354.652 | 451.066 | 559.459 | 0.306 | 0.634 |
| **SL1344_0441** | *tig* | 5158.067 | 2972.147 | 5498.334 | 2936.732 | 0.938 | 1.01 |
| **SL1344_0442** | *clpP* | 424.416 | 515.61 | 937.63 | 790.384 | 0.453 | 0.652 |
| **SL1344_0443** | *clpX* | 555.366 | 532.459 | 836.308 | 759.162 | 0.664 | 0.701 |
| **SL1344_0444** | *lon* | 870.346 | 247.418 | 1145.141 | 471.03 | 0.76 | 0.525 |
| **SL1344_0445** | *hupB* | 10703.556 | 25368.939 | 10410.398 | 11085.416 | 1.03 | 2.29 |
| **SL1344_0446** | *cypD* | 552.259 | 283.604 | 517.791 | 491.739 | 1.07 | 0.577 |
| **SL1344_0447** | *comE1* | 16.732 | 66.207 | 10.896 | 42.232 | 1.54 | 1.57 |
| **SL1344_0448** | *ybaW* | 2.67 | 26.083 | 4.671 | 17.028 | 0.572 | 1.53 |
| **SL1344_0449** | *ybaX* | 200.887 | 129.983 | 325.563 | 249.362 | 0.617 | 0.521 |
| **SL1344_0450** | *ybaE* | 10.336 | 22.514 | 29.499 | 41.918 | 0.35 | 0.537 |

| | | | | | | | |
|---|---|---|---|---|---|---|---|
| **SL1344_0451** | *cof* | 4.915 | 4.817 | 16.717 | 13.639 | 0.294 | 0.353 |
| **SL1344_0452** | *SL1344-0452* | 2.018 | 4.38 | 14.798 | 12.891 | 0.136 | 0.34 |
| **SL1344_0453** | *ybaO* | 24.503 | 102.699 | 167.73 | 170.305 | 0.146 | 0.603 |
| **SL1344_0454** | *mdlA* | 13.522 | 11.625 | 26.199 | 36.902 | 0.516 | 0.315 |
| **SL1344_0455** | *mdlB* | 7.474 | 5.077 | 32.221 | 31.682 | 0.232 | 0.16 |
| **SL1344_0457** | *amtB* | 2.438 | 1.321 | 37.039 | 15.743 | 0.0658 | 0.0839 |
| **SL1344_0458** | *tesB* | 120.039 | 109.972 | 56.947 | 77.563 | 2.11 | 1.42 |
| **SL1344_0459** | *ybaY* | 100.111 | 446.315 | 51.688 | 139.219 | 1.94 | 3.21 |
| **SL1344_0460** | *ybaZ* | 11.573 | 34.011 | 75.819 | 56.882 | 0.153 | 0.598 |
| **SL1344_0461** | *ylaB* | 7.213 | 8.377 | 20.659 | 19.05 | 0.349 | 0.44 |
| **SL1344_0462** | *rpmE2* | 12.247 | 28.147 | 32.134 | 23.067 | 0.381 | 1.22 |
| **SL1344_0464** | *ylaC* | 60.577 | 195.11 | 77.771 | 253.082 | 0.779 | 0.771 |
| **SL1344_0465** | *maa* | 11.367 | 11.583 | 178.56 | 97.854 | 0.0637 | 0.118 |
| **SL1344_0466** | *hha* | 264.888 | 926.828 | 648.753 | 618.161 | 0.408 | 1.5 |
| **SL1344_0467** | *ybaJ* | 3.157 | 29.385 | 265.325 | 315.872 | 0.0119 | 0.093 |
| **SL1344_0468** | *acrB* | 252.318 | 94.301 | 285.71 | 176.528 | 0.883 | 0.534 |
| **SL1344_0469** | *acrA* | 400.975 | 345.632 | 255.575 | 297.072 | 1.57 | 1.16 |
| **SL1344_0470** | *acrR* | 82.094 | 59.493 | 106.539 | 99.377 | 0.771 | 0.599 |
| **SL1344_0471** | *aefA* | 29.166 | 22.35 | 36.982 | 32.886 | 0.789 | 0.68 |
| **SL1344_0472** | *SL1344-0472* | 3.099 | 18.677 | 61.804 | 73.725 | 0.0501 | 0.253 |
| **SL1344_0473** | *ybaM* | 50.033 | 56.279 | 28.588 | 27.345 | 1.75 | 2.06 |
| **SL1344_0474** | *priC* | 11.128 | 14.501 | 73.628 | 65.601 | 0.151 | 0.221 |
| **SL1344_0476** | *apt* | 375.541 | 54.959 | 350.366 | 119.142 | 1.07 | 0.461 |
| **SL1344_0477** | *dnaX* | 23.506 | 23.626 | 158.938 | 139.859 | 0.148 | 0.169 |

| | | | | | | | |
|---|---|---|---|---|---|---|---|
| **SL1344_0478** | *ybaB* | 262.428 | 283.01 | 408.379 | 342.02 | 0.643 | 0.827 |
| **SL1344_0479** | *recR* | 57.045 | 26.602 | 650.357 | 530.019 | 0.0877 | 0.0502 |
| **SL1344_0480** | *htpG* | 2143.941 | 433.885 | 1511.587 | 348.549 | 1.42 | 1.24 |
| **SL1344_0481** | *adk* | 1753.897 | 839.773 | 1175.166 | 729.413 | 1.49 | 1.15 |
| **SL1344_0482** | *hemH* | 74.254 | 64.136 | 92.824 | 121.12 | 0.8 | 0.53 |
| **SL1344_0483** | *aes* | 6.212 | 7.348 | 5.457 | 13.259 | 1.14 | 0.554 |
| **SL1344_0484** | *gsk* | 118.162 | 22.986 | 75.693 | 50.896 | 1.56 | 0.452 |
| **SL1344_0485** | *ybaL* | 31.768 | 27.987 | 90.149 | 69.177 | 0.352 | 0.405 |
| **SL1344_0486** | *fsr* | 29.621 | 21.225 | 30.118 | 24.1 | 0.984 | 0.881 |
| **SL1344_0487** | *ushA* | 244.405 | 243.979 | 118.562 | 196.078 | 2.06 | 1.24 |
| **SL1344_0488** | *ybaK* | 76.583 | 93.246 | 133.511 | 88.656 | 0.574 | 1.05 |
| **SL1344_0489** | *ybaP* | 26.954 | 11.893 | 66.815 | 53.315 | 0.403 | 0.223 |
| **SL1344_0491** | *copA* | 12.421 | 37.382 | 19.683 | 23.29 | 0.631 | 1.61 |
| **SL1344_0492** | *cueR* | 58.059 | 53.014 | 185.483 | 251.587 | 0.313 | 0.211 |
| **SL1344_0493** | *ybbJ* | 44.559 | 23.875 | 60.29 | 67.626 | 0.739 | 0.353 |
| **SL1344_0494** | *ybbK* | 189.641 | 96.919 | 103.387 | 81.992 | 1.83 | 1.18 |
| **SL1344_0495** | *ybbL* | 13.62 | 24.781 | 21.463 | 20.434 | 0.635 | 1.21 |
| **SL1344_0496** | *ybbM* | 12.825 | 11.773 | 29.592 | 30.936 | 0.433 | 0.381 |
| **SL1344_0497** | *ybbN* | 661.661 | 149.249 | 987.473 | 183.476 | 0.67 | 0.813 |
| **SL1344_0498** | *ybbO* | 80.922 | 59.993 | 64.432 | 69.306 | 1.26 | 0.866 |
| **SL1344_0499** | *tesA* | 62.756 | 74.879 | 36.599 | 43.601 | 1.71 | 1.72 |
| **SL1344_0500** | *ybbA* | 15.682 | 11.386 | 48.832 | 56.452 | 0.321 | 0.202 |
| **SL1344_0501** | *ybbP* | 8.775 | 7.351 | 33.482 | 25.5 | 0.262 | 0.288 |
| **SL1344_0502** | *SL1344-0502* | 49.903 | 27.952 | 29.331 | 25.65 | 1.7 | 1.09 |

| | | | | | | | |
|---|---|---|---|---|---|---|---|
| **SL1344_0503** | *sfbA* | 19.447 | 17.926 | 18.201 | 12.162 | 1.07 | 1.47 |
| **SL1344_0504** | *sfbB* | 10.069 | 7.491 | 16.141 | 9.177 | 0.624 | 0.816 |
| **SL1344_0505** | *sfbC* | 15.157 | 4.638 | 16.074 | 17.914 | 0.943 | 0.259 |
| **SL1344_0506** | *ybbB* | 23.137 | 24.04 | 66.512 | 82.893 | 0.348 | 0.29 |
| **SL1344_0507** | *ybbS* | 2.937 | 7.705 | 11.058 | 17.684 | 0.266 | 0.436 |
| **SL1344_0508** | *allA* | 148.537 | 167.87 | 47.641 | 57.819 | 3.12 | 2.9 |
| **SL1344_0509** | *allR* | 134 | 87.788 | 206.386 | 194.08 | 0.649 | 0.452 |
| **SL1344_0510** | *gcl* | 6.079 | 4.733 | 2.655 | 5.712 | 2.29 | 0.829 |
| **SL1344_0512** | *garRa* | 6.532 | 5.494 | 10.683 | 106.123 | 0.611 | 0.0518 |
| **SL1344_0516** | *allB* | 3.477 | 7.342 | 3.263 | 4.19 | 1.07 | 1.75 |
| **SL1344_0518** | *glxK* | 5.77 | 4.677 | 3.868 | 4.482 | 1.49 | 1.04 |
| **SL1344_0519** | *ylbA* | 48.35 | 39.809 | 51.711 | 46.266 | 0.935 | 0.86 |
| **SL1344_0520** | *allC* | 8.764 | 11.942 | 8.177 | 9.488 | 1.07 | 1.26 |
| **SL1344_0521** | *allD* | 6.427 | 7.191 | 5.462 | 6.678 | 1.18 | 1.08 |
| **SL1344_0522** | *fdrA* | 1.955 | 0.327 | 7.534 | 5.446 | 0.26 | 0.06 |
| **SL1344_0525** | *arcC* | 11.653 | 4.641 | 12.108 | 11.63 | 0.962 | 0.399 |
| **SL1344_0526** | *purK* | 14.466 | 10.7 | 44.03 | 33.665 | 0.329 | 0.318 |
| **SL1344_0527** | *purE* | 8.241 | 3.735 | 20.24 | 23.42 | 0.407 | 0.159 |
| **SL1344_0528** | *lpxH* | 24.234 | 9.126 | 169.74 | 121.663 | 0.143 | 0.075 |
| **SL1344_0529** | *ppiB* | 2128.366 | 2109.346 | 921.46 | 768.642 | 2.31 | 2.74 |
| **SL1344_0530** | *cysS* | 282.305 | 176.777 | 289.01 | 212.969 | 0.977 | 0.83 |
| **SL1344_0531** | *SL1344-0531* | 4.305 | 7.091 | 13.902 | 10.873 | 0.31 | 0.652 |
| **SL1344_0532** | *SL1344-0532* | 9.595 | 30.283 | 7.856 | 13.469 | 1.22 | 2.25 |
| **SL1344_0534** | *ybcJ* | 109.496 | 49.18 | 227.166 | 195.02 | 0.482 | 0.252 |

| | | | | | | | |
|---|---|---|---|---|---|---|---|
| **SL1344_0535** | *folD* | 124.943 | 86.851 | 191.735 | 182.248 | 0.652 | 0.477 |
| **SL1344_0536** | *fimA* | 594.669 | 699.254 | 247.79 | 232.705 | 2.4 | 3 |
| **SL1344_0538** | *fimC* | 13.581 | 21.496 | 6.827 | 7.99 | 1.99 | 2.69 |
| **SL1344_0539** | *fimD* | 3.262 | 2.603 | 9.218 | 10.031 | 0.354 | 0.26 |
| **SL1344_0540** | *fimH* | 4.228 | 4.791 | 6.898 | 8.467 | 0.613 | 0.566 |
| **SL1344_0542** | *fimZ* | 37.406 | 9.779 | 5.662 | 6.341 | 6.61 | 1.54 |
| **SL1344_0545** | *fimW* | 1.686 | 6.267 | 6.244 | 9.437 | 0.27 | 0.664 |
| **SL1344_0547** | *SL1344-0547* | 15.971 | 13.785 | 12.012 | 11.073 | 1.33 | 1.24 |
| **SL1344_0548** | *gtrBa* | 25.734 | 14.089 | 9.589 | 6.051 | 2.68 | 2.33 |
| **SL1344_0551** | *ykgD* | 15.231 | 4.614 | 7.881 | 9.247 | 1.93 | 0.499 |
| **SL1344_0552** | *ykgC* | 3.795 | 3.232 | 7.406 | 13.639 | 0.512 | 0.237 |
| **SL1344_0555** | *SL1344-0555* | 19.379 | 15.154 | 49.252 | 52.901 | 0.393 | 0.286 |
| **SL1344_0556** | *pheP* | 95.517 | 23.064 | 138.126 | 45.896 | 0.692 | 0.503 |
| **SL1344_0557** | *ybdG* | 47.889 | 46.129 | 50.705 | 49.555 | 0.944 | 0.931 |
| **SL1344_0558** | *apeE* | 7.688 | 18.947 | 16.184 | 20.327 | 0.475 | 0.932 |
| **SL1344_0559** | *SL1344-0559* | 14.181 | 15.448 | 23.238 | 33.735 | 0.61 | 0.458 |
| **SL1344_0560** | *SL1344-0560* | 4.318 | 4.066 | 6.536 | 4.459 | 0.661 | 0.912 |
| **SL1344_0566** | *nfnB* | 307.105 | 325.441 | 156.52 | 126.07 | 1.96 | 2.58 |
| **SL1344_0567** | *ybdF* | 59.676 | 50.693 | 208.446 | 101.105 | 0.286 | 0.501 |
| **SL1344_0568** | *ramR* | 26.139 | 57.386 | 194.359 | 202.155 | 0.134 | 0.284 |
| **SL1344_0569** | *ramA* | 2.423 | 21.48 | 355.483 | 423.412 | 0.00682 | 0.0507 |
| **SL1344_0570** | *ybdJ* | 3.566 | 13.932 | 59.592 | 43.114 | 0.0598 | 0.323 |
| **SL1344_0571** | *ybdK* | 8.464 | 12.765 | 7.879 | 11.863 | 1.07 | 1.08 |
| **SL1344_0576** | *entF* | 2.788 | 0.735 | 6.071 | 4.96 | 0.459 | 0.148 |

| | | | | | | | |
|---|---|---|---|---|---|---|---|
| **SL1344_0577** | *fepE* | 66.327 | 33.741 | 39.341 | 37.81 | 1.69 | 0.892 |
| **SL1344_0580** | *fepD* | 5.285 | 0 | 13.867 | 4.39 | 0.381 | 0 |
| **SL1344_0582** | *fepB* | 3.464 | 2.203 | 7.266 | 2.855 | 0.477 | 0.772 |
| **SL1344_0583** | *entC* | 4.329 | 0.347 | 6.217 | 3.927 | 0.696 | 0.0884 |
| **SL1344_0584** | *entE* | 2.535 | 1.267 | 4.539 | 3.122 | 0.559 | 0.406 |
| **SL1344_0585** | *entB* | 11.039 | 6.025 | 8.772 | 6.904 | 1.26 | 0.873 |
| **SL1344_0586** | *entA* | 4.306 | 1.889 | 9.197 | 6.397 | 0.468 | 0.295 |
| **SL1344_0588** | *cstAa* | 756.429 | 1072.288 | 461.723 | 906.235 | 1.64 | 1.18 |
| **SL1344_0589** | *ybdD* | 28.401 | 271.741 | 32.945 | 175.602 | 0.862 | 1.55 |
| **SL1344_0590** | *ybdH* | 14.785 | 23.673 | 21.92 | 10.191 | 0.675 | 2.32 |
| **SL1344_0591** | *ybdL* | 10.605 | 5.097 | 18.214 | 9.705 | 0.582 | 0.525 |
| **SL1344_0592** | *ybdM* | 18.199 | 18.491 | 34.914 | 33.53 | 0.521 | 0.551 |
| **SL1344_0593** | *ybdN* | 6.705 | 8.75 | 11.657 | 10.818 | 0.575 | 0.809 |
| **SL1344_0594** | *ybdO* | 26.549 | 24.557 | 11.828 | 11.059 | 2.24 | 2.22 |
| **SL1344_0595** | *dsbG* | 89.386 | 71.39 | 52.395 | 40.978 | 1.71 | 1.74 |
| **SL1344_0596** | *ahpC* | 4916.386 | 2222.631 | 1538.147 | 909.793 | 3.2 | 2.44 |
| **SL1344_0597** | *ahpF* | 208.881 | 127.486 | 107.8 | 97.642 | 1.94 | 1.31 |
| **SL1344_0602** | *ybdQ* | 367.999 | 1317.615 | 942.581 | 1216.775 | 0.39 | 1.08 |
| **SL1344_0603** | *ybdR* | 1.624 | 7.247 | 9.199 | 17.856 | 0.177 | 0.406 |
| **SL1344_0604** | *rnk* | 367.984 | 288.304 | 534.226 | 254.672 | 0.689 | 1.13 |
| **SL1344_0605** | *rna* | 66.676 | 60.267 | 68.663 | 92.82 | 0.971 | 0.649 |
| **SL1344_0614** | *citB* | 5.302 | 9.289 | 14.105 | 14.628 | 0.376 | 0.635 |
| **SL1344_0616** | *pagP* | 8.264 | 14.958 | 30.15 | 39.792 | 0.274 | 0.376 |
| **SL1344_0617** | *cspE* | 8555.67 | 11684.647 | 12052.601 | 9597.632 | 0.71 | 1.22 |

| | | | | | | | |
|---|---|---|---|---|---|---|---|
| **SL1344_0618** | *ccrB* | 56.882 | 31.531 | 110.512 | 84.626 | 0.515 | 0.373 |
| **SL1344_0619** | *SL1344-0619* | 3.601 | 14.828 | 21.896 | 16.763 | 0.164 | 0.885 |
| **SL1344_0620** | *tatE* | 366.483 | 490.153 | 848.958 | 553.268 | 0.432 | 0.886 |
| **SL1344_0621** | *lipA* | 350.815 | 292.717 | 275.529 | 271.924 | 1.27 | 1.08 |
| **SL1344_0622** | *ybeF* | 21.717 | 21.818 | 18.409 | 18.78 | 1.18 | 1.16 |
| **SL1344_0623** | *lipB* | 49.329 | 32.527 | 47.566 | 54.497 | 1.04 | 0.597 |
| **SL1344_0624** | *ybeD* | 299.559 | 246.319 | 2095.387 | 478.356 | 0.143 | 0.515 |
| **SL1344_0625** | *dacA* | 335.874 | 127.063 | 242.376 | 151.517 | 1.39 | 0.839 |
| **SL1344_0626** | *rlpA* | 135.959 | 125.343 | 145.045 | 83.924 | 0.937 | 1.49 |
| **SL1344_0627** | *mrdB* | 25.954 | 16.44 | 51.009 | 47.576 | 0.509 | 0.346 |
| **SL1344_0628** | *mrdA* | 18.891 | 10.765 | 36.106 | 23.645 | 0.523 | 0.455 |
| **SL1344_0629** | *ybeA* | 40.222 | 26.453 | 150.413 | 130.245 | 0.267 | 0.203 |
| **SL1344_0630** | *ybeB* | 482.304 | 180.321 | 662.96 | 286.8 | 0.728 | 0.629 |
| **SL1344_0632** | *cobD* | 15.569 | 10.126 | 91.127 | 86.955 | 0.171 | 0.116 |
| **SL1344_0633** | *nadD* | 36.697 | 23.309 | 37.628 | 40.524 | 0.975 | 0.575 |
| **SL1344_0634** | *holA* | 18.871 | 11.864 | 68.141 | 87.227 | 0.277 | 0.136 |
| **SL1344_0635** | *rlpB* | 117.087 | 161.248 | 210.563 | 208.404 | 0.556 | 0.774 |
| **SL1344_0636** | *leuS* | 550.945 | 194.557 | 251.406 | 180.831 | 2.19 | 1.08 |
| **SL1344_0640** | *SL1344-0640* | 9.482 | 20.981 | 23.374 | 30.083 | 0.406 | 0.697 |
| **SL1344_0641** | *ybeL* | 70.684 | 140.348 | 163.027 | 296.777 | 0.434 | 0.473 |
| **SL1344_0642** | *ybeQ* | 1.605 | 4.576 | 11.803 | 10.317 | 0.136 | 0.444 |
| **SL1344_0648** | *hscC* | 1.656 | 2.146 | 11.777 | 11.14 | 0.141 | 0.193 |
| **SL1344_0649** | *SL1344-0649* | 19.397 | 11.694 | 86.379 | 31.419 | 0.225 | 0.372 |
| **SL1344_0650** | *rihA* | 337.585 | 81.829 | 219.722 | 98.73 | 1.54 | 0.829 |

| | | | | | | | |
|---|---|---|---|---|---|---|---|
| **SL1344_0651** | *gltL* | 388.269 | 839.959 | 381.128 | 582.257 | 1.02 | 1.44 |
| **SL1344_0652** | *gltK* | 116.723 | 137.151 | 333.037 | 400.974 | 0.35 | 0.342 |
| **SL1344_0653** | *gltJ* | 1273.518 | 3136.689 | 407.658 | 669.47 | 3.12 | 4.69 |
| **SL1344_0654** | *gltI* | 2121.547 | 2902.901 | 1143.071 | 1582.839 | 1.86 | 1.83 |
| **SL1344_0655** | *lnt* | 34.347 | 19.712 | 94.134 | 46.221 | 0.365 | 0.426 |
| **SL1344_0656** | *ybeX* | 129.366 | 81.254 | 150.87 | 135.61 | 0.857 | 0.599 |
| **SL1344_0657** | *ybeY* | 212.55 | 54.819 | 265.108 | 165.732 | 0.802 | 0.331 |
| **SL1344_0658** | *phoL* | 269.264 | 86.123 | 369.969 | 161.816 | 0.728 | 0.532 |
| **SL1344_0659** | *miaB* | 303.657 | 135.899 | 334.674 | 131.126 | 0.907 | 1.04 |
| **SL1344_0660** | *ubiF* | 21.644 | 24.64 | 90.458 | 93.611 | 0.239 | 0.263 |
| **SL1344_0662** | *asnB* | 53.613 | 86.201 | 53.601 | 60.963 | 1 | 1.41 |
| **SL1344_0663** | *nagD* | 115.166 | 61.879 | 129.754 | 105.77 | 0.888 | 0.585 |
| **SL1344_0664** | *nagC* | 35.439 | 35.71 | 97.046 | 183.648 | 0.365 | 0.194 |
| **SL1344_0665** | *nagA* | 214.996 | 304.653 | 135.113 | 237.56 | 1.59 | 1.28 |
| **SL1344_0666** | *nagB* | 638.501 | 1024.055 | 215.23 | 417.922 | 2.97 | 2.45 |
| **SL1344_0667** | *nagE* | 439.032 | 622.222 | 352.068 | 558.032 | 1.25 | 1.12 |
| **SL1344_0668** | *glnS* | 145.857 | 140.161 | 221.153 | 194.039 | 0.66 | 0.722 |
| **SL1344_0669** | *ybfM* | 31.511 | 48.683 | 279.092 | 347.147 | 0.113 | 0.14 |
| **SL1344_0670** | *ybfN* | 75.726 | 66.591 | 47.789 | 121.838 | 1.58 | 0.547 |
| **SL1344_0671** | *SL1344-0671* | 4.944 | 6.307 | 15.271 | 12.34 | 0.324 | 0.511 |
| **SL1344_0672** | *SL1344-0672* | 3.946 | 3.461 | 5.848 | 6.766 | 0.675 | 0.512 |
| **SL1344_0674** | *SL1344-0674* | 13.984 | 29.278 | 23.276 | 35.134 | 0.601 | 0.833 |
| **SL1344_0675** | *fur* | 363.529 | 604.233 | 516.976 | 934.434 | 0.703 | 0.647 |
| **SL1344_0676** | *fldA* | 891.594 | 947.686 | 493.955 | 603.241 | 1.81 | 1.57 |

| | | | | | | | |
|---|---|---|---|---|---|---|---|
| **SL1344_0677** | *ybfE* | 115.87 | 117.711 | 71.429 | 90.542 | 1.62 | 1.3 |
| **SL1344_0678** | *ybfF* | 63.647 | 59.993 | 64.803 | 53.877 | 0.982 | 1.11 |
| **SL1344_0679** | *seqA* | 533.948 | 833.166 | 424.425 | 468.704 | 1.26 | 1.78 |
| **SL1344_0680** | *pgm* | 372.733 | 327.049 | 317.574 | 348.11 | 1.17 | 0.939 |
| **SL1344_0682** | *potE* | 0.852 | 38.545 | 7.494 | 56.307 | 0.114 | 0.685 |
| **SL1344_0683** | *speF* | 2.369 | 57.442 | 4.107 | 31.359 | 0.577 | 1.83 |
| **SL1344_0684** | *kdpE* | 29.597 | 59.995 | 52.124 | 56.275 | 0.568 | 1.07 |
| **SL1344_0685** | *kdpD* | 8.576 | 5.143 | 23.307 | 22.009 | 0.368 | 0.234 |
| **SL1344_0689** | *ybfA* | 20.875 | 62.106 | 108.737 | 95.314 | 0.192 | 0.652 |
| **SL1344_0692** | *ybgI* | 146.236 | 575.986 | 143.464 | 175.004 | 1.02 | 3.29 |
| **SL1344_0693** | *ybgJ* | 70.367 | 32.716 | 167.48 | 143.895 | 0.42 | 0.227 |
| **SL1344_0694** | *ybgK* | 58.243 | 49.868 | 116.015 | 101.043 | 0.502 | 0.494 |
| **SL1344_0695** | *ybgL* | 14.819 | 9.902 | 41.84 | 59.575 | 0.354 | 0.166 |
| **SL1344_0697** | *fimB* | 17.226 | 19.675 | 23.136 | 31.172 | 0.745 | 0.631 |
| **SL1344_0710** | *nei* | 23.767 | 51.875 | 37.561 | 58.646 | 0.633 | 0.885 |
| **SL1344_0711** | *abrB* | 2.213 | 12.079 | 14.441 | 35.051 | 0.153 | 0.345 |
| **SL1344_0712** | *gltA* | 3573.699 | 5892.05 | 1211.866 | 2256.107 | 2.95 | 2.61 |
| **SL1344_0714** | *sdhC* | 495.715 | 1981.512 | 1314.934 | 1837.467 | 0.377 | 1.08 |
| **SL1344_0715** | *sdhD* | 1778.382 | 2486.294 | 3510.023 | 2045.821 | 0.507 | 1.22 |
| **SL1344_0716** | *sdhA* | 2601.269 | 1663.653 | 1215.903 | 1489.18 | 2.14 | 1.12 |
| **SL1344_0717** | *sdhB* | 8072.961 | 4200.218 | 2480.849 | 3384.498 | 3.25 | 1.24 |
| **SL1344_0718** | *sucA* | 1142.861 | 1246.595 | 1474.221 | 1789.527 | 0.775 | 0.697 |
| **SL1344_0719** | *sucB* | 5501.355 | 8122.548 | 2380.915 | 2558.616 | 2.31 | 3.17 |
| **SL1344_0720** | *sucC* | 2441.932 | 2125.679 | 2463.102 | 2085.512 | 0.991 | 1.02 |

| | | | | | | | |
|---|---|---|---|---|---|---|---|
| **SL1344_0721** | *sucD* | 1849.314 | 1812.336 | 1071.052 | 1455.704 | 1.73 | 1.24 |
| **SL1344_0722** | *cydA* | 485.024 | 701.326 | 457.24 | 726.536 | 1.06 | 0.965 |
| **SL1344_0723** | *cydB* | 243.112 | 711.478 | 283.087 | 559.528 | 0.859 | 1.27 |
| **SL1344_0724** | *ybgT* | 2041.167 | 2989.355 | 396.146 | 612.954 | 5.15 | 4.88 |
| **SL1344_0725** | *ybgE* | 35.894 | 88.042 | 198.782 | 708.302 | 0.181 | 0.124 |
| **SL1344_0726** | *ybgC* | 19.585 | 38.965 | 130.446 | 148.954 | 0.15 | 0.262 |
| **SL1344_0727** | *tolQ* | 117.364 | 90.008 | 176.054 | 164.577 | 0.667 | 0.547 |
| **SL1344_0728** | *tolR* | 74.51 | 87.682 | 235.269 | 232.94 | 0.317 | 0.376 |
| **SL1344_0729** | *tolA* | 30.758 | 20.006 | 88.96 | 91.066 | 0.346 | 0.22 |
| **SL1344_0730** | *tolB* | 955.396 | 427.224 | 795.172 | 926.537 | 1.2 | 0.461 |
| **SL1344_0731** | *pal* | 4865.551 | 7764.918 | 2328.283 | 3190.868 | 2.09 | 2.43 |
| **SL1344_0732** | *ybgF* | 119.889 | 105.868 | 582.554 | 799.066 | 0.206 | 0.132 |
| **SL1344_0733** | *nadA* | 3.459 | 3.844 | 11.055 | 15.957 | 0.313 | 0.241 |
| **SL1344_0734** | *pnuC* | 11.51 | 12.187 | 21.306 | 31.768 | 0.54 | 0.384 |
| **SL1344_0735** | *ybgR* | 43.308 | 58.532 | 36.109 | 35.174 | 1.2 | 1.66 |
| **SL1344_0736** | *ybgS* | 2.6 | 42.008 | 6.298 | 15.994 | 0.413 | 2.63 |
| **SL1344_0737** | *aroG* | 266.852 | 168.923 | 175.567 | 166.519 | 1.52 | 1.01 |
| **SL1344_0738** | *SL1344-0738* | 35.994 | 71.385 | 11.422 | 11.957 | 3.15 | 5.97 |
| **SL1344_0739** | *SL1344-0739* | 31.976 | 18.654 | 14.15 | 11.975 | 2.26 | 1.56 |
| **SL1344_0740** | *SL1344-0740* | 18.203 | 42.981 | 11.907 | 19.973 | 1.53 | 2.15 |
| **SL1344_0741** | *SL1344-0741* | 4.917 | 5.723 | 5.8 | 4.904 | 0.848 | 1.17 |
| **SL1344_0744** | *dcoA* | 7.733 | 3.677 | 2.341 | 2.342 | 3.3 | 1.57 |
| **SL1344_0746** | *SL1344-0746* | 3.557 | 5.255 | 17.126 | 12.902 | 0.208 | 0.407 |
| **SL1344_0749** | *gpmA* | 2926.78 | 2606.94 | 1418.059 | 1006.873 | 2.06 | 2.59 |

| | | | | | | | |
|---|---|---|---|---|---|---|---|
| **SL1344_0750** | *galM* | 352.721 | 328.998 | 178.142 | 278.765 | 1.98 | 1.18 |
| **SL1344_0751** | *galK* | 148.836 | 170.793 | 111.3 | 171.735 | 1.34 | 0.995 |
| **SL1344_0752** | *galT* | 62.813 | 54.248 | 106.19 | 87.039 | 0.592 | 0.623 |
| **SL1344_0753** | *galE* | 195.161 | 174.768 | 169.94 | 141.317 | 1.15 | 1.24 |
| **SL1344_0755** | *modF* | 35.653 | 26.13 | 357.835 | 109.706 | 0.0996 | 0.238 |
| **SL1344_0756** | *modE* | 43.814 | 36.037 | 58.601 | 70.912 | 0.748 | 0.508 |
| **SL1344_0757** | *ybhT* | 327.542 | 569.565 | 1045.624 | 538.865 | 0.313 | 1.06 |
| **SL1344_0758** | *modA* | 65.542 | 29.792 | 357.77 | 81.44 | 0.183 | 0.366 |
| **SL1344_0759** | *modB* | 42.894 | 13.013 | 156.665 | 43.031 | 0.274 | 0.302 |
| **SL1344_0760** | *modC* | 42.649 | 8.093 | 62.207 | 20.023 | 0.686 | 0.404 |
| **SL1344_0761** | *ybhA* | 38.306 | 60.629 | 29.759 | 45.819 | 1.29 | 1.32 |
| **SL1344_0762** | *ybhE* | 313.148 | 268.261 | 134.083 | 130.07 | 2.34 | 2.06 |
| **SL1344_0763** | *ybhC* | 244.661 | 206.607 | 155.929 | 148.124 | 1.57 | 1.39 |
| **SL1344_0764** | *hutI* | 34.627 | 52.461 | 114.787 | 159.098 | 0.302 | 0.33 |
| **SL1344_0765** | *hutG* | 32.174 | 23.396 | 61.867 | 106.392 | 0.52 | 0.22 |
| **SL1344_0766** | *hutC* | 17.985 | 32.606 | 49.798 | 98.641 | 0.361 | 0.331 |
| **SL1344_0767** | *hutU* | 51.751 | 83.756 | 100.893 | 138.075 | 0.513 | 0.607 |
| **SL1344_0768** | *hutH* | 45.884 | 48.389 | 77.103 | 80.405 | 0.595 | 0.602 |
| **SL1344_0769** | *ybhB* | 49.515 | 68.449 | 64.62 | 54.035 | 0.766 | 1.27 |
| **SL1344_0770** | *bioA* | 27.624 | 6.486 | 14.059 | 6.073 | 1.96 | 1.07 |
| **SL1344_0771** | *bioB* | 22.006 | 4.509 | 19.074 | 6.62 | 1.15 | 0.681 |
| **SL1344_0773** | *bioC* | 5.089 | 1.17 | 8.154 | 3.07 | 0.624 | 0.381 |
| **SL1344_0774** | *bioDa* | 5.859 | 0.891 | 14.504 | 6.757 | 0.404 | 0.132 |
| **SL1344_0775** | *uvrB* | 52.637 | 41.008 | 108.27 | 104.034 | 0.486 | 0.394 |

| | | | | | | | |
|---|---|---|---|---|---|---|---|
| **SL1344_0776** | *slrP* | 31.168 | 30.695 | 24.05 | 29.187 | 1.3 | 1.05 |
| **SL1344_0777** | *ybhK* | 26.83 | 21.851 | 55.99 | 52.348 | 0.479 | 0.417 |
| **SL1344_0778** | *moaA* | 20.868 | 8.726 | 53.944 | 54.463 | 0.387 | 0.16 |
| **SL1344_0779** | *moaB* | 71.079 | 63.115 | 106.757 | 213.32 | 0.666 | 0.296 |
| **SL1344_0780** | *moaC* | 38.732 | 12.876 | 84.811 | 54.228 | 0.457 | 0.237 |
| **SL1344_0781** | *moaD* | 458.287 | 177.881 | 121.748 | 87.312 | 3.76 | 2.04 |
| **SL1344_0782** | *moaE* | 45.735 | 28.079 | 121.213 | 93.246 | 0.377 | 0.301 |
| **SL1344_0783** | *ybhL* | 203.612 | 135.464 | 383.939 | 338.469 | 0.53 | 0.4 |
| **SL1344_0788** | *ybhO* | 2.669 | 5.422 | 5.945 | 12.829 | 0.449 | 0.423 |
| **SL1344_0789** | *ybhP* | 4.757 | 9.769 | 10.2 | 14.335 | 0.466 | 0.681 |
| **SL1344_0790** | *ybhQ* | 157.42 | 231.041 | 634.513 | 806.784 | 0.248 | 0.286 |
| **SL1344_0791** | *ybhR* | 12.085 | 19.479 | 23.959 | 24.397 | 0.504 | 0.798 |
| **SL1344_0792** | *ybhS* | 6.438 | 9.021 | 21.93 | 28.112 | 0.294 | 0.321 |
| **SL1344_0793** | *ybhF* | 17.619 | 14.215 | 33.799 | 43.43 | 0.521 | 0.327 |
| **SL1344_0794** | *SL1344-0794* | 37.086 | 72.734 | 64.99 | 70.983 | 0.571 | 1.02 |
| **SL1344_0795** | *ybiH* | 25.607 | 11.992 | 41.63 | 46.961 | 0.615 | 0.255 |
| **SL1344_0796** | *rhlE* | 28.405 | 5.631 | 38.809 | 13.002 | 0.732 | 0.433 |
| **SL1344_0797** | *dinG* | 5.786 | 3.458 | 29.36 | 29.367 | 0.197 | 0.118 |
| **SL1344_0798** | *ybiB* | 15.482 | 32.79 | 30.732 | 53.094 | 0.504 | 0.618 |
| **SL1344_0800** | *ybiI* | 7.538 | 29.807 | 9.665 | 28.616 | 0.78 | 1.04 |
| **SL1344_0801** | *ybiN* | 10.342 | 6.707 | 25.124 | 15.264 | 0.412 | 0.439 |
| **SL1344_0802** | *ybiO* | 1.438 | 1.591 | 8.062 | 8.549 | 0.178 | 0.186 |
| **SL1344_0803** | *glnQ* | 880.793 | 785.584 | 367.242 | 440.202 | 2.4 | 1.78 |
| **SL1344_0804** | *glnP* | 141.349 | 142.02 | 393.391 | 423.826 | 0.359 | 0.335 |

| | | | | | | | |
|---|---|---|---|---|---|---|---|
| **SL1344_0805** | *glnH* | 2034.955 | 1705.904 | 1455.263 | 1479.44 | 1.4 | 1.15 |
| **SL1344_0806** | *dps* | 338.371 | 1629.406 | 188.881 | 528.528 | 1.79 | 3.08 |
| **SL1344_0807** | *ybiF* | 37.196 | 23.057 | 59.171 | 31.013 | 0.629 | 0.743 |
| **SL1344_0808** | *ompX* | 4215.075 | 1663.884 | 2462.525 | 1484.795 | 1.71 | 1.12 |
| **SL1344_0809** | *ybiP* | 5.429 | 3.055 | 26.796 | 13.886 | 0.203 | 0.22 |
| **SL1344_0810** | *SL1344-0810* | 109.397 | 54.388 | 182.234 | 177.158 | 0.6 | 0.307 |
| **SL1344_0811** | *ybiR* | 16.7 | 10.879 | 80.314 | 82.16 | 0.208 | 0.132 |
| **SL1344_0812** | *ybiS* | 135.935 | 60.34 | 106.319 | 54.396 | 1.28 | 1.11 |
| **SL1344_0813** | *ybiT* | 232.876 | 58.414 | 155.967 | 102.696 | 1.49 | 0.569 |
| **SL1344_0817** | *ybiU* | 24.033 | 14.238 | 11.381 | 15.031 | 2.11 | 0.947 |
| **SL1344_0818** | *ybiV1* | 75.491 | 59.959 | 115.579 | 95.133 | 0.653 | 0.63 |
| **SL1344_0819** | *pflF* | 1.192 | 1.594 | 3.329 | 3.796 | 0.358 | 0.42 |
| **SL1344_0821** | *moeB* | 20.757 | 22.039 | 58.303 | 49.905 | 0.356 | 0.442 |
| **SL1344_0822** | *moeA* | 50.472 | 54.384 | 42.941 | 49.097 | 1.18 | 1.11 |
| **SL1344_0823** | *ybiK* | 63.153 | 56.829 | 138.193 | 238.991 | 0.457 | 0.238 |
| **SL1344_0824** | *yliA* | 24.601 | 26.126 | 57.324 | 109.722 | 0.429 | 0.238 |
| **SL1344_0825** | *yliB* | 42.655 | 84.861 | 38.101 | 71.374 | 1.12 | 1.19 |
| **SL1344_0826** | *yliC* | 15.104 | 7.386 | 62.344 | 133.259 | 0.242 | 0.0554 |
| **SL1344_0827** | *yliD* | 15.513 | 15.364 | 37.178 | 42.233 | 0.417 | 0.364 |
| **SL1344_0828** | *rimO* | 319.453 | 48.784 | 96.173 | 43.07 | 3.32 | 1.13 |
| **SL1344_0829** | *bssR* | 3.237 | 20.725 | 44.429 | 107.986 | 0.0729 | 0.192 |
| **SL1344_0831** | *SL1344-0831* | 2.077 | 5.41 | 8.133 | 2.8 | 0.255 | 1.93 |
| **SL1344_0833** | *SL1344-0833* | 4.729 | 5.727 | 10.654 | 34.17 | 0.444 | 0.168 |
| **SL1344_0834** | *SL1344-0834* | 5.196 | 9.118 | 84.184 | 82.776 | 0.0617 | 0.11 |

| | | | | | | | |
|---|---|---|---|---|---|---|---|
| **SL1344_0835** | *SL1344-0835* | 4.983 | 5.649 | 6.999 | 9.661 | 0.712 | 0.585 |
| **SL1344_0838** | *yliJ* | 742.715 | 560.66 | 356.932 | 297.203 | 2.08 | 1.89 |
| **SL1344_0839** | *dacC* | 102.348 | 137.513 | 104.635 | 166.84 | 0.978 | 0.824 |
| **SL1344_0840** | *deoR* | 12.712 | 13.084 | 91.045 | 90.057 | 0.14 | 0.145 |
| **SL1344_0841** | *ybjG* | 40.338 | 46.576 | 51.673 | 76.904 | 0.781 | 0.606 |
| **SL1344_0842** | *mdfA* | 10.082 | 1.765 | 51.626 | 40.141 | 0.195 | 0.044 |
| **SL1344_0843** | *SL1344-0843* | 12.115 | 27.509 | 32.855 | 24.208 | 0.369 | 1.14 |
| **SL1344_0844** | *SL1344-0844* | 13.675 | 3.985 | 32.707 | 22.224 | 0.418 | 0.179 |
| **SL1344_0845** | *SL1344-0845* | 2.192 | 7.229 | 18.7 | 14.912 | 0.117 | 0.485 |
| **SL1344_0846** | *SL1344-0846* | 39.638 | 17.55 | 38.648 | 25.756 | 1.03 | 0.681 |
| **SL1344_0848** | *grxA* | 3388.429 | 2552.085 | 1907.489 | 1443.677 | 1.78 | 1.77 |
| **SL1344_0852** | *rimK* | 28.384 | 9.265 | 72.398 | 54.998 | 0.392 | 0.168 |
| **SL1344_0853** | *ybjN* | 173.116 | 158.574 | 104.144 | 88.757 | 1.66 | 1.79 |
| **SL1344_0854** | *potF* | 20.902 | 15.34 | 24.023 | 22.05 | 0.87 | 0.696 |
| **SL1344_0855** | *potG* | 12.841 | 4.499 | 16.056 | 18.891 | 0.8 | 0.238 |
| **SL1344_0857** | *potI* | 5.668 | 2.412 | 33.046 | 27.866 | 0.172 | 0.0866 |
| **SL1344_0858** | *ybjO* | 9.741 | 10.476 | 99.51 | 98.654 | 0.0979 | 0.106 |
| **SL1344_0859** | *rumB* | 10.625 | 6.556 | 50.831 | 31.191 | 0.209 | 0.21 |
| **SL1344_0863** | *artJ* | 20.459 | 35.405 | 17.823 | 17.705 | 1.15 | 2 |
| **SL1344_0864** | *artM* | 11.591 | 16.472 | 73.291 | 93.029 | 0.158 | 0.177 |
| **SL1344_0865** | *artQ* | 7.513 | 7.684 | 120.572 | 116.645 | 0.0623 | 0.0659 |
| **SL1344_0866** | *artI* | 187.125 | 273.851 | 78.049 | 94.866 | 2.4 | 2.89 |
| **SL1344_0867** | *artP* | 50.587 | 80.991 | 127.536 | 156.482 | 0.397 | 0.518 |
| **SL1344_0868** | *ybjP* | 257.312 | 274.985 | 65.015 | 101.542 | 3.96 | 2.71 |

| | | | | | | | |
|---|---|---|---|---|---|---|---|
| **SL1344_0869** | *SL1344-0869* | 61.387 | 117.358 | 130.977 | 175.968 | 0.469 | 0.667 |
| **SL1344_0870** | *ybjR* | 5.271 | 7.285 | 43.821 | 68.899 | 0.12 | 0.106 |
| **SL1344_0871** | *SL1344-0871* | 15.937 | 6.306 | 69.846 | 66.074 | 0.228 | 0.0954 |
| **SL1344_0872** | *ybjT* | 6.316 | 5.787 | 32.493 | 30.552 | 0.194 | 0.189 |
| **SL1344_0873** | *ltaA* | 84.184 | 67.071 | 107.668 | 85.977 | 0.782 | 0.78 |
| **SL1344_0874** | *poxB* | 18.492 | 42.736 | 8.048 | 19.242 | 2.3 | 2.22 |
| **SL1344_0876** | *hcp* | 1.003 | 4.815 | 26.15 | 25.919 | 0.0383 | 0.186 |
| **SL1344_0877** | *ybjE* | 73.32 | 69.395 | 70.688 | 68.507 | 1.04 | 1.01 |
| **SL1344_0878** | *ybjD* | 50.595 | 54.819 | 70.906 | 82.578 | 0.714 | 0.664 |
| **SL1344_0879** | *ybjX* | 157.302 | 128.953 | 219.859 | 112.81 | 0.715 | 1.14 |
| **SL1344_0880** | *macA* | 49.778 | 44.132 | 78.538 | 105.812 | 0.634 | 0.417 |
| **SL1344_0881** | *macB* | 10.671 | 16.455 | 45.985 | 55.324 | 0.232 | 0.297 |
| **SL1344_0882** | *cspD* | 531.149 | 2929.51 | 1031.016 | 3191.278 | 0.515 | 0.918 |
| **SL1344_0883** | *clpS* | 344.286 | 151.3 | 451.986 | 610.873 | 0.762 | 0.248 |
| **SL1344_0884** | *clpA* | 409.135 | 529.443 | 477.103 | 816.834 | 0.858 | 0.648 |
| **SL1344_0885** | *tnp* | 51.07 | 80.618 | 96.359 | 121.474 | 0.53 | 0.664 |
| **SL1344_0888** | *SL1344-0888* | 15.366 | 15.25 | 16.282 | 22.431 | 0.944 | 0.68 |
| **SL1344_0890** | *SL1344-0890* | 21.549 | 33.19 | 42.366 | 55.251 | 0.509 | 0.601 |
| **SL1344_0891** | *infA* | 2377.505 | 605.979 | 1328.928 | 984.597 | 1.79 | 0.615 |
| **SL1344_0892** | *SL1344-0892* | 12.949 | 85.381 | 216.544 | 440.634 | 0.0598 | 0.194 |
| **SL1344_0893** | *aat* | 7.473 | 10.517 | 21.454 | 26.82 | 0.348 | 0.392 |
| **SL1344_0894** | *cydC* | 30.732 | 18.21 | 47.414 | 37.392 | 0.648 | 0.487 |
| **SL1344_0895** | *cydD* | 35.543 | 26.446 | 50.385 | 46.004 | 0.705 | 0.575 |
| **SL1344_0896** | *trxB* | 888.771 | 592.676 | 705.073 | 641.45 | 1.26 | 0.924 |

| | | | | | | | |
|---|---|---|---|---|---|---|---|
| **SL1344_0897** | *lrp* | 2837.383 | 2622.597 | 1416.294 | 1238.835 | 2 | 2.12 |
| **SL1344_0898** | *ftsK* | 69.755 | 56.235 | 106.438 | 114.385 | 0.655 | 0.492 |
| **SL1344_0899** | *lolA* | 120.41 | 188.358 | 136.84 | 180.535 | 0.88 | 1.04 |
| **SL1344_0900** | *ycaJ* | 24.664 | 34.213 | 37.389 | 61.419 | 0.66 | 0.557 |
| **SL1344_0901** | *serS* | 1262.31 | 560.741 | 628.409 | 552.093 | 2.01 | 1.02 |
| **SL1344_0902** | *dmsA* | 7.505 | 28.85 | 9.968 | 27.917 | 0.753 | 1.03 |
| **SL1344_0903** | *dmsB* | 4.789 | 25.205 | 5.452 | 20.071 | 0.878 | 1.26 |
| **SL1344_0904** | *dmsC* | 6.235 | 24.642 | 11.118 | 22.947 | 0.561 | 1.07 |
| **SL1344_0906** | *ycaM* | 69.976 | 14.733 | 66.088 | 23.397 | 1.06 | 0.63 |
| **SL1344_0907** | *pflA* | 156.56 | 134.558 | 129.899 | 117.7 | 1.21 | 1.14 |
| **SL1344_0909** | *sopD2* | 6.536 | 64.053 | 11.201 | 27.075 | 0.584 | 2.37 |
| **SL1344_0910** | *pflB* | 807.693 | 765.783 | 336.941 | 388.545 | 2.4 | 1.97 |
| **SL1344_0911** | *focA* | 128.668 | 83.798 | 63.412 | 68.591 | 2.03 | 1.22 |
| **SL1344_0912** | *ycaO* | 179.869 | 19.893 | 105.958 | 20.884 | 1.7 | 0.953 |
| **SL1344_0913** | *ycaP* | 12.3 | 34.845 | 14.068 | 26.621 | 0.874 | 1.31 |
| **SL1344_0914** | *serC* | 218.894 | 133.232 | 185.824 | 172.405 | 1.18 | 0.773 |
| **SL1344_0915** | *aroA* | 40.708 | 29.667 | 45.444 | 35.684 | 0.896 | 0.831 |
| **SL1344_0916** | *ycaL* | 10.332 | 49.454 | 40.075 | 33.539 | 0.258 | 1.47 |
| **SL1344_0917** | *cmk* | 68.8 | 40.674 | 297.529 | 257.448 | 0.231 | 0.158 |
| **SL1344_0918** | *rpsA* | 6013.709 | 3873.23 | 8388.606 | 5031.146 | 0.717 | 0.77 |
| **SL1344_0919** | *himD* | 560.996 | 1124.618 | 773.931 | 745.217 | 0.725 | 1.51 |
| **SL1344_0921** | *msbA* | 122.789 | 77.9 | 162.548 | 145.432 | 0.755 | 0.536 |
| **SL1344_0922** | *lpxK* | 17.431 | 10.224 | 68.459 | 45.688 | 0.255 | 0.224 |
| **SL1344_0923** | *ycaQ* | 9.362 | 9.047 | 23.429 | 20.316 | 0.4 | 0.445 |

| | | | | | | | |
|---|---|---|---|---|---|---|---|
| **SL1344_0924** | *ycaR* | 151.706 | 405.526 | 410.124 | 275.348 | 0.37 | 1.47 |
| **SL1344_0925** | *kdsB* | 89.307 | 55.091 | 168.509 | 193.884 | 0.53 | 0.284 |
| **SL1344_0926** | *SL1344-0926* | 4.105 | 8.37 | 19.164 | 30.751 | 0.214 | 0.272 |
| **SL1344_0927** | *ycbC* | 19.351 | 7.091 | 25.463 | 19.791 | 0.76 | 0.358 |
| **SL1344_0928** | *smtA* | 33.426 | 18.19 | 263.018 | 217.128 | 0.127 | 0.0838 |
| **SL1344_0929** | *mukF* | 87.158 | 90.181 | 213.1 | 185.846 | 0.409 | 0.485 |
| **SL1344_0930** | *mukE* | 245.762 | 149.164 | 226.439 | 171.207 | 1.09 | 0.871 |
| **SL1344_0931** | *mukB* | 220.279 | 98.781 | 173.793 | 163.179 | 1.27 | 0.605 |
| **SL1344_0932** | *ycbB* | 173.92 | 148.561 | 123.831 | 155.298 | 1.4 | 0.957 |
| **SL1344_0933** | *ycbK* | 243.247 | 511.956 | 474.496 | 449.62 | 0.513 | 1.14 |
| **SL1344_0934** | *ycbL* | 189.367 | 242.168 | 405.433 | 512.569 | 0.467 | 0.472 |
| **SL1344_0935** | *aspC* | 269.183 | 295.501 | 361.969 | 351.801 | 0.744 | 0.84 |
| **SL1344_0936** | *ompF* | 652.602 | 440.207 | 384.676 | 265.812 | 1.7 | 1.66 |
| **SL1344_0937** | *asnS* | 1138.213 | 490.714 | 456.81 | 320.469 | 2.49 | 1.53 |
| **SL1344_0938** | *SL1344-0938* | 10.25 | 20.171 | 38.014 | 54.429 | 0.27 | 0.371 |
| **SL1344_0939** | *dpaL* | 2.144 | 5.151 | 3.481 | 4.716 | 0.616 | 1.09 |
| **SL1344_0941** | *pncB* | 19.781 | 18.546 | 48.683 | 42.489 | 0.406 | 0.436 |
| **SL1344_0942** | *SL1344-0942* | 6.73 | 10.784 | 49.009 | 85.159 | 0.137 | 0.127 |
| **SL1344_0945** | *SL1344-0945* | 0.427 | 3.37 | 3.1 | 4.683 | 0.138 | 0.72 |
| **SL1344_0946** | *recEa* | 0.328 | 3.229 | 2.409 | 2.89 | 0.136 | 1.12 |
| **SL1344_0947** | *SL1344-0947* | 1.184 | 21.54 | 2.628 | 19.182 | 0.45 | 1.12 |
| **SL1344_0950** | *SL1344-0950* | 41.911 | 54.45 | 212.958 | 338.327 | 0.197 | 0.161 |
| **SL1344_0956** | *SL1344-0956* | 24.41 | 230.711 | 25.373 | 140.503 | 0.962 | 1.64 |
| **SL1344_0957** | *SL1344-0957* | 652.15 | 570.041 | 321.963 | 219.366 | 2.03 | 2.6 |

| | | | | | | | |
|---|---|---|---|---|---|---|---|
| **SL1344_0962** | *SL1344-0962* | 757.055 | 151.871 | 220.461 | 298.206 | 3.43 | 0.509 |
| **SL1344_0964** | *SL1344-0964* | 170.348 | 97.648 | 107.047 | 27.886 | 1.59 | 3.5 |
| **SL1344_0965** | *gtgA* | 4.825 | 15.446 | 5.426 | 7.18 | 0.889 | 2.15 |
| **SL1344_0971** | *SL1344-0971* | 4.245 | 3.955 | 8.752 | 10.746 | 0.485 | 0.368 |
| **SL1344_0972** | *SL1344-0972* | 11.742 | 11.566 | 12.842 | 13.959 | 0.914 | 0.829 |
| **SL1344_0977** | *SL1344-0977* | 5.084 | 5.446 | 9.913 | 11.72 | 0.513 | 0.465 |
| **SL1344_0978** | *SL1344-0978* | 15.577 | 8.013 | 9.163 | 10.362 | 1.7 | 0.773 |
| **SL1344_0980** | *SL1344-0980* | 1.686 | 1.292 | 8.614 | 12.042 | 0.196 | 0.107 |
| **SL1344_0982** | *SL1344-0982* | 78.213 | 72.256 | 55.418 | 33.209 | 1.41 | 2.18 |
| **SL1344_0983** | *sodCa* | 203.411 | 500.988 | 104.572 | 221.639 | 1.95 | 2.26 |
| **SL1344_0984** | *SL1344-0984* | 4.508 | 7.525 | 16.685 | 17.509 | 0.27 | 0.43 |
| **SL1344_0985** | *SL1344-0985* | 8.422 | 5.162 | 13.404 | 13.891 | 0.628 | 0.372 |
| **SL1344_0986** | *SL1344-0986* | 11.083 | 38.208 | 10.778 | 13.067 | 1.03 | 2.92 |
| **SL1344_0989** | *SL1344-0989* | 3.255 | 7.714 | 2.217 | 3.282 | 1.47 | 2.35 |
| **SL1344_0991** | *sseI* | 4.459 | 14.741 | 4.882 | 5.664 | 0.913 | 2.6 |
| **SL1344_0995** | *gtgE* | 64.537 | 121.884 | 19.929 | 23.862 | 3.24 | 5.11 |
| **SL1344_0997** | *pepN* | 156.674 | 180.401 | 103.397 | 133.516 | 1.52 | 1.35 |
| **SL1344_0998** | *pyrD* | 39.349 | 43.971 | 37.016 | 26.066 | 1.06 | 1.69 |
| **SL1344_0999** | *ycbW* | 59.085 | 83.555 | 184.952 | 215.249 | 0.319 | 0.388 |
| **SL1344_1000** | *SL1344-1000* | 23.571 | 16.178 | 45.722 | 26.575 | 0.516 | 0.609 |
| **SL1344_1001** | *rlmL* | 57.482 | 30.479 | 130.281 | 102.436 | 0.441 | 0.298 |
| **SL1344_1002** | *uup* | 76.351 | 48.378 | 115.903 | 67.189 | 0.659 | 0.72 |
| **SL1344_1003** | *pqiA* | 17.654 | 22.62 | 40.987 | 45.697 | 0.431 | 0.495 |
| **SL1344_1004** | *pqiB* | 118.497 | 67.648 | 79.372 | 62.694 | 1.49 | 1.08 |

| | | | | | | | |
|---|---|---|---|---|---|---|---|
| **SL1344_1005** | *ymbA* | 14.379 | 31.84 | 37.24 | 69.321 | 0.386 | 0.459 |
| **SL1344_1006** | *rmf* | 12.685 | 119.847 | 102.832 | 405.139 | 0.123 | 0.296 |
| **SL1344_1007** | *fabA* | 685.354 | 413.763 | 841.971 | 383.554 | 0.814 | 1.08 |
| **SL1344_1008** | *lonH* | 93.447 | 30.245 | 126.271 | 76.271 | 0.74 | 0.397 |
| **SL1344_1009** | *ycbG* | 136.16 | 139.796 | 102.699 | 175.283 | 1.33 | 0.798 |
| **SL1344_1010** | *ompA* | 18983.572 | 15357.753 | 5896.229 | 4391.017 | 3.22 | 3.5 |
| **SL1344_1011** | *sulA* | 68.248 | 67.488 | 139.994 | 97.332 | 0.488 | 0.693 |
| **SL1344_1012** | *sxy* | 14.359 | 62.409 | 17.389 | 50.911 | 0.826 | 1.23 |
| **SL1344_1013** | *yccS* | 10.69 | 9.379 | 25.126 | 19.273 | 0.425 | 0.487 |
| **SL1344_1014** | *yccF* | 27.147 | 39.565 | 73.608 | 33.915 | 0.369 | 1.17 |
| **SL1344_1015** | *helD* | 35.632 | 32.637 | 53.44 | 63.489 | 0.667 | 0.514 |
| **SL1344_1016** | *mgsA* | 1019.716 | 1461.346 | 675.294 | 423.208 | 1.51 | 3.45 |
| **SL1344_1018** | *SL1344-1018* | 78.642 | 179.596 | 55.181 | 73.348 | 1.43 | 2.45 |
| **SL1344_1019** | *yccV* | 176.28 | 46.631 | 509.675 | 223.464 | 0.346 | 0.209 |
| **SL1344_1020** | *yccW* | 84.884 | 20.485 | 82.211 | 36.787 | 1.03 | 0.557 |
| **SL1344_1021** | *SL1344-1021* | 4.636 | 8.797 | 19.455 | 17.881 | 0.238 | 0.492 |
| **SL1344_1022** | *SL1344-1022* | 1.821 | 4.884 | 17.553 | 29.883 | 0.104 | 0.163 |
| **SL1344_1023** | *yccX* | 33.166 | 37.146 | 21.353 | 38.069 | 1.55 | 0.976 |
| **SL1344_1024** | *yccK* | 147.627 | 254.358 | 219.395 | 309.855 | 0.673 | 0.821 |
| **SL1344_1025** | *yccA* | 214.695 | 220.511 | 537.317 | 463.574 | 0.4 | 0.476 |
| **SL1344_1027** | *pipB* | 1.892 | 11.492 | 5.564 | 13.193 | 0.34 | 0.871 |
| **SL1344_1029** | *pipC* | 68.541 | 257.566 | 203.313 | 154.617 | 0.337 | 1.67 |
| **SL1344_1030** | *sopB* | 214.764 | 149.397 | 99.235 | 75.907 | 2.16 | 1.97 |
| **SL1344_1031** | *orfX* | 40.848 | 358.802 | 51.562 | 325.354 | 0.792 | 1.1 |

| | | | | | | | |
|---|---|---|---|---|---|---|---|
| **SL1344_1033** | *pipD* | 8.577 | 14.761 | 9.85 | 17.386 | 0.871 | 0.849 |
| **SL1344_1034** | *copS* | 4.987 | 2.84 | 9.873 | 9.122 | 0.505 | 0.311 |
| **SL1344_1035** | *copR* | 17.354 | 20.944 | 22.551 | 28.095 | 0.77 | 0.745 |
| **SL1344_1036** | *SL1344-1036* | 4.609 | 15.226 | 18.139 | 30.768 | 0.254 | 0.495 |
| **SL1344_1038** | *hpaB* | 2.424 | 11.446 | 3.944 | 8.199 | 0.615 | 1.4 |
| **SL1344_1039** | *hpaR* | 13.02 | 26.838 | 40.962 | 50.386 | 0.318 | 0.533 |
| **SL1344_1040** | *hpaG* | 31.983 | 40.971 | 17.782 | 29.97 | 1.8 | 1.37 |
| **SL1344_1041** | *hpaE* | 11.903 | 17.063 | 10.164 | 16.729 | 1.17 | 1.02 |
| **SL1344_1042** | *hpaD* | 9.518 | 7.984 | 13.041 | 23.725 | 0.73 | 0.337 |
| **SL1344_1044** | *hpaH* | 8.025 | 21.828 | 8.47 | 13.413 | 0.947 | 1.63 |
| **SL1344_1045** | *hpaI* | 3.513 | 9.104 | 7.241 | 9.433 | 0.485 | 0.965 |
| **SL1344_1046** | *hpaX* | 2.794 | 1.778 | 4.321 | 7.849 | 0.647 | 0.227 |
| **SL1344_1048** | *SL1344-1048* | 2.708 | 4.965 | 4.607 | 6.19 | 0.588 | 0.802 |
| **SL1344_1050** | *yccD* | 30.951 | 62.686 | 63.016 | 108.015 | 0.491 | 0.58 |
| **SL1344_1051** | *cbpA* | 77.319 | 260.49 | 77.521 | 107.111 | 0.997 | 2.43 |
| **SL1344_1052** | *scsA* | 8.643 | 8.62 | 102.292 | 90.987 | 0.0845 | 0.0947 |
| **SL1344_1056** | *agp* | 198.458 | 375.705 | 79.648 | 209.219 | 2.49 | 1.8 |
| **SL1344_1057** | *yccJ* | 273.125 | 307.587 | 105.325 | 194.915 | 2.59 | 1.58 |
| **SL1344_1058** | *wrbA* | 162.611 | 664.261 | 130.519 | 300.433 | 1.25 | 2.21 |
| **SL1344_1059** | *ymdF* | 2.819 | 72.88 | 34.562 | 171.267 | 0.0816 | 0.426 |
| **SL1344_1060** | *ycdC* | 10.097 | 5.748 | 31.298 | 38.671 | 0.323 | 0.149 |
| **SL1344_1061** | *SL1344-1061* | 23.917 | 80.217 | 105.172 | 332.642 | 0.227 | 0.241 |
| **SL1344_1062** | *putA* | 652.452 | 861.688 | 246.723 | 721.277 | 2.64 | 1.19 |
| **SL1344_1063** | *putP* | 849.473 | 1690.84 | 635.271 | 1415.735 | 1.34 | 1.19 |

| | | | | | | | |
|---|---|---|---|---|---|---|---|
| **SL1344_1064** | *phoH* | 87.095 | 375.032 | 59.024 | 233.248 | 1.48 | 1.61 |
| **SL1344_1065** | *SL1344-1065* | 61.006 | 23.676 | 53.072 | 39.53 | 1.15 | 0.599 |
| **SL1344_1066** | *SL1344-1066* | 4.508 | 597.06 | 4.98 | 231.08 | 0.905 | 2.58 |
| **SL1344_1067** | *SL1344-1067* | 8.084 | 562.249 | 23.895 | 777.734 | 0.338 | 0.723 |
| **SL1344_1068** | *SL1344-1068* | 4.079 | 577.8 | 3.828 | 319.739 | 1.07 | 1.81 |
| **SL1344_1069** | *SL1344-1069* | 8.2 | 1892.228 | 4.138 | 473.528 | 1.98 | 4 |
| **SL1344_1070** | *SL1344-1070* | 1.109 | 209.852 | 4.98 | 227.104 | 0.223 | 0.924 |
| **SL1344_1071** | *SL1344-1071* | 14.705 | 398.358 | 12.303 | 223.878 | 1.2 | 1.78 |
| **SL1344_1072** | *ycdW* | 175.629 | 102.213 | 129.168 | 87.471 | 1.36 | 1.17 |
| **SL1344_1073** | *ycdX* | 85.583 | 111.802 | 122.775 | 107.036 | 0.697 | 1.04 |
| **SL1344_1074** | *ycdY* | 53.542 | 46.328 | 39.523 | 48.619 | 1.35 | 0.953 |
| **SL1344_1075** | *ycdZ* | 107.899 | 19.086 | 542.617 | 119.67 | 0.199 | 0.159 |
| **SL1344_1083** | *ymdA* | 12.355 | 59.545 | 59.178 | 155.618 | 0.209 | 0.383 |
| **SL1344_1084** | *SL1344-1084* | 15.456 | 46.985 | 65.046 | 174.222 | 0.238 | 0.27 |
| **SL1344_1085** | *ymdC* | 4.058 | 8.32 | 43.55 | 75.624 | 0.0932 | 0.11 |
| **SL1344_1086** | *mdoC* | 11.275 | 8.598 | 16.757 | 11.681 | 0.673 | 0.736 |
| **SL1344_1087** | *mdoG* | 83.281 | 116.159 | 145.281 | 134.4 | 0.573 | 0.864 |
| **SL1344_1088** | *mdoH* | 150.894 | 88.823 | 167.205 | 135.749 | 0.902 | 0.654 |
| **SL1344_1090** | *msyB* | 47.513 | 122.257 | 16.248 | 80.08 | 2.92 | 1.53 |
| **SL1344_1092** | *htrB* | 40.17 | 68.317 | 122.664 | 140.321 | 0.327 | 0.487 |
| **SL1344_1093** | *trhO* | 49.357 | 14.341 | 76.789 | 41.653 | 0.643 | 0.344 |
| **SL1344_1094** | *yceI* | 55.597 | 53.614 | 14.312 | 16.791 | 3.88 | 3.19 |
| **SL1344_1095** | *yceJ* | 17.045 | 5.342 | 10.759 | 8.228 | 1.58 | 0.649 |
| **SL1344_1097** | *solA* | 80.989 | 82.124 | 71.812 | 64.39 | 1.13 | 1.28 |

| | | | | | | | |
|---|---|---|---|---|---|---|---|
| **SL1344_1098** | *bssS* | 219.34 | 611.928 | 1081.922 | 1166.863 | 0.203 | 0.524 |
| **SL1344_1099** | *dinI* | 67.135 | 145.167 | 242.442 | 190.089 | 0.277 | 0.764 |
| **SL1344_1100** | *pyrC* | 94.643 | 167.032 | 97.974 | 101.311 | 0.966 | 1.65 |
| **SL1344_1101** | *yceB* | 77.027 | 176.298 | 126.5 | 192.181 | 0.609 | 0.917 |
| **SL1344_1102** | *grxB* | 134.192 | 397.42 | 73.564 | 133.506 | 1.82 | 2.98 |
| **SL1344_1104** | *rimJ* | 29.344 | 86.858 | 232.451 | 292.212 | 0.126 | 0.297 |
| **SL1344_1105** | *yceH* | 272.678 | 444.343 | 207.307 | 348.951 | 1.32 | 1.27 |
| **SL1344_1106** | *mviM* | 27.867 | 78.18 | 100 | 179.928 | 0.279 | 0.435 |
| **SL1344_1107** | *mviN* | 21.799 | 11.099 | 48.017 | 33.436 | 0.454 | 0.332 |
| **SL1344_1108** | *flgN* | 263.926 | 145.53 | 437.727 | 421.043 | 0.603 | 0.346 |
| **SL1344_1109** | *flgM* | 230.133 | 465.969 | 347.69 | 325.098 | 0.662 | 1.43 |
| **SL1344_1110** | *flgA* | 26.637 | 84.511 | 206.579 | 533.766 | 0.129 | 0.158 |
| **SL1344_1111** | *flgB* | 237.345 | 839.581 | 216.77 | 567.478 | 1.09 | 1.48 |
| **SL1344_1112** | *flgC* | 75.418 | 170.809 | 225.671 | 449.786 | 0.334 | 0.38 |
| **SL1344_1113** | *flgD* | 674.511 | 604.309 | 368.673 | 725.226 | 1.83 | 0.833 |
| **SL1344_1114** | *flgE* | 542.81 | 852.006 | 332.335 | 561.001 | 1.63 | 1.52 |
| **SL1344_1115** | *flgF* | 66.398 | 114.988 | 237.618 | 493.395 | 0.279 | 0.233 |
| **SL1344_1116** | *flgG* | 160.422 | 574.054 | 380.573 | 700.26 | 0.422 | 0.82 |
| **SL1344_1117** | *flgH* | 70.627 | 58.387 | 178.645 | 214.496 | 0.395 | 0.272 |
| **SL1344_1118** | *flgI* | 81.19 | 166.46 | 117.579 | 220.78 | 0.691 | 0.754 |
| **SL1344_1119** | *flgJ* | 25.458 | 36.55 | 102.588 | 187.71 | 0.248 | 0.195 |
| **SL1344_1120** | *flgK* | 371.3 | 162.441 | 517.053 | 156.028 | 0.718 | 1.04 |
| **SL1344_1121** | *flgL* | 384.081 | 247.272 | 586.768 | 233.399 | 0.655 | 1.06 |
| **SL1344_1122** | *rne* | 310.475 | 316.413 | 170.685 | 203.12 | 1.82 | 1.56 |

| | | | | | | | |
|---|---|---|---|---|---|---|---|
| **SL1344_1123** | *rluC* | 34.16 | 6.023 | 79.748 | 53.999 | 0.428 | 0.112 |
| **SL1344_1124** | *SL1344-1124* | 3.182 | 8.96 | 27.17 | 44.328 | 0.117 | 0.202 |
| **SL1344_1126** | *maf-a* | 90.967 | 73.951 | 53.426 | 45.96 | 1.7 | 1.61 |
| **SL1344_1127** | *yceD* | 871.926 | 522.669 | 4175.774 | 2857.409 | 0.209 | 0.183 |
| **SL1344_1128** | *rpmF* | 3984.733 | 10240.718 | 7691.153 | 4867.448 | 0.518 | 2.1 |
| **SL1344_1129** | *plsX* | 15.292 | 12.597 | 486.917 | 459.487 | 0.0314 | 0.0274 |
| **SL1344_1130** | *fabH* | 524.931 | 255.828 | 872.526 | 445.281 | 0.602 | 0.575 |
| **SL1344_1131** | *fabD* | 1207.247 | 500.871 | 1141.465 | 567.89 | 1.06 | 0.882 |
| **SL1344_1132** | *fabG* | 814.144 | 917.871 | 1890.008 | 1644.358 | 0.431 | 0.558 |
| **SL1344_1133** | *acpP* | 11928.539 | 21804.974 | 6759.126 | 6335.363 | 1.76 | 3.44 |
| **SL1344_1134** | *fabF* | 1113.207 | 707.432 | 1586.273 | 1205.885 | 0.702 | 0.587 |
| **SL1344_1135** | *pabC* | 10.889 | 9.489 | 24.691 | 37.014 | 0.441 | 0.256 |
| **SL1344_1136** | *yceG* | 33.525 | 24.309 | 85.241 | 71.374 | 0.393 | 0.341 |
| **SL1344_1137** | *tmk* | 111.197 | 74.908 | 163.465 | 124.547 | 0.68 | 0.601 |
| **SL1344_1138** | *holB* | 11.662 | 5.821 | 60.129 | 66.185 | 0.194 | 0.0879 |
| **SL1344_1139** | *ycfH* | 83.747 | 74.926 | 79.948 | 51.63 | 1.05 | 1.45 |
| **SL1344_1140** | *ptsG* | 265.755 | 272.275 | 393.81 | 419.285 | 0.675 | 0.649 |
| **SL1344_1142** | *ycfF* | 810.963 | 750.312 | 420.544 | 370.138 | 1.93 | 2.03 |
| **SL1344_1143** | *ycfL* | 32.044 | 51.696 | 209.698 | 232.826 | 0.153 | 0.222 |
| **SL1344_1144** | *ycfM* | 71.237 | 103.15 | 87.165 | 176.025 | 0.817 | 0.586 |
| **SL1344_1145** | *thiK* | 11.767 | 3.71 | 84.978 | 92.935 | 0.138 | 0.0399 |
| **SL1344_1146** | *nagZ* | 97.042 | 80.087 | 149.236 | 172.386 | 0.65 | 0.465 |
| **SL1344_1147** | *ycfP* | 270.136 | 627.85 | 300.068 | 383.611 | 0.9 | 1.64 |
| **SL1344_1148** | *ndh* | 35.698 | 24.185 | 64.323 | 35.87 | 0.555 | 0.674 |

| | | | | | | | |
|---|---|---|---|---|---|---|---|
| **SL1344_1149** | *ycfJ* | 44.725 | 110.598 | 93.454 | 120.625 | 0.479 | 0.917 |
| **SL1344_1150** | *ycfQ* | 27.643 | 51.016 | 113.386 | 182.064 | 0.244 | 0.28 |
| **SL1344_1151** | *ycfR* | 26.156 | 56.948 | 61.403 | 39.548 | 0.426 | 1.44 |
| **SL1344_1152** | *ycfS* | 4.412 | 13.309 | 12.17 | 28.234 | 0.363 | 0.471 |
| **SL1344_1153** | *mfd* | 50.9 | 27.785 | 53.487 | 50.686 | 0.952 | 0.548 |
| **SL1344_1154** | *ycfU* | 48.786 | 27.038 | 127.776 | 107.032 | 0.382 | 0.253 |
| **SL1344_1155** | *lolD* | 64.422 | 51.743 | 67.191 | 63.168 | 0.959 | 0.819 |
| **SL1344_1156** | *ycfW* | 21.3 | 23.712 | 71.741 | 68.536 | 0.297 | 0.346 |
| **SL1344_1157** | *ycfX* | 194.004 | 114.115 | 140.695 | 151.754 | 1.38 | 0.752 |
| **SL1344_1158** | *cobB* | 119.325 | 101.204 | 222.725 | 170.546 | 0.536 | 0.593 |
| **SL1344_1159** | *potD* | 306.714 | 207.442 | 197.524 | 118.1 | 1.55 | 1.76 |
| **SL1344_1160** | *potC* | 91.144 | 22.499 | 260.45 | 88.811 | 0.35 | 0.253 |
| **SL1344_1161** | *sifA* | 2.576 | 6.93 | 13.472 | 7.151 | 0.191 | 0.969 |
| **SL1344_1162** | *potB* | 24.116 | 18.58 | 58.243 | 16.147 | 0.414 | 1.15 |
| **SL1344_1163** | *potA* | 76.843 | 17.648 | 91.543 | 43.296 | 0.839 | 0.408 |
| **SL1344_1164** | *pepT* | 67.761 | 88.759 | 51.344 | 84.606 | 1.32 | 1.05 |
| **SL1344_1166** | *SL1344-1166* | 55.712 | 81.092 | 102.606 | 130.044 | 0.543 | 0.624 |
| **SL1344_1167** | *ycfD* | 52.019 | 37.891 | 88.934 | 95.896 | 0.585 | 0.395 |
| **SL1344_1168** | *phoQ* | 22.764 | 31.873 | 71.782 | 105.105 | 0.317 | 0.303 |
| **SL1344_1169** | *phoP* | 946.847 | 1791.126 | 281 | 473.511 | 3.37 | 3.78 |
| **SL1344_1170** | *purB* | 177.929 | 94.912 | 94.585 | 42.749 | 1.88 | 2.22 |
| **SL1344_1171** | *ycfC* | 19.64 | 80.618 | 123.787 | 84.29 | 0.159 | 0.956 |
| **SL1344_1172** | *mnmA* | 112.239 | 57.637 | 197.114 | 196.816 | 0.569 | 0.293 |
| **SL1344_1173** | *ymfB* | 18.066 | 9.57 | 20.016 | 23.655 | 0.903 | 0.405 |

| | | | | | | | |
|---|---|---|---|---|---|---|---|
| **SL1344_1174** | *SL1344-1174* | 5.74 | 12.574 | 18.464 | 38.393 | 0.311 | 0.327 |
| **SL1344_1175** | *rluB* | 3.911 | 7.456 | 21.957 | 32.347 | 0.178 | 0.23 |
| **SL1344_1176** | *icdA* | 2610.893 | 2840.2 | 1966.113 | 1755.043 | 1.33 | 1.62 |
| **SL1344_1177** | *SL1344-1177* | 20.047 | 15.72 | 21.792 | 17.968 | 0.92 | 0.875 |
| **SL1344_1180** | *msgA* | 116.169 | 273.219 | 143.367 | 155.147 | 0.81 | 1.76 |
| **SL1344_1181** | *envE* | 138.687 | 61.115 | 46.278 | 23.206 | 3 | 2.63 |
| **SL1344_1184** | *pagC* | 112.231 | 748.234 | 25.416 | 115.145 | 4.42 | 6.5 |
| **SL1344_1185** | *pliC* | 99.687 | 204.458 | 25.557 | 88.235 | 3.9 | 2.32 |
| **SL1344_1187** | *SL1344-1187* | 80.317 | 40.842 | 413.559 | 225.101 | 0.194 | 0.181 |
| **SL1344_1189** | *SL1344-1189* | 27.758 | 95.179 | 15.795 | 42.347 | 1.76 | 2.25 |
| **SL1344_1190** | *SL1344-1190* | 33.074 | 87.24 | 84.427 | 130.182 | 0.392 | 0.67 |
| **SL1344_1191** | *SL1344-1191* | 42.106 | 255.361 | 47.877 | 198.947 | 0.879 | 1.28 |
| **SL1344_1192** | *SL1344-1192* | 6.193 | 87.207 | 26.247 | 83.82 | 0.236 | 1.04 |
| **SL1344_1193** | *SL1344-1193* | 1.893 | 5.522 | 8.376 | 30.751 | 0.226 | 0.18 |
| **SL1344_1194** | *SL1344-1194* | 7.492 | 18.753 | 6.558 | 32.935 | 1.14 | 0.569 |
| **SL1344_1195** | *SL1344-1195* | 2.304 | 12.776 | 9.339 | 31.625 | 0.247 | 0.404 |
| **SL1344_1196** | *SL1344-1196* | 8.352 | 22.554 | 13.338 | 16.098 | 0.626 | 1.4 |
| **SL1344_1197** | *SL1344-1197* | 13.215 | 36.195 | 21.702 | 46.065 | 0.609 | 0.786 |
| **SL1344_1198** | *SL1344-1198* | 11.327 | 12.911 | 37.059 | 45.596 | 0.306 | 0.283 |
| **SL1344_1199** | *aadA* | 4.877 | 6.38 | 14.446 | 21.789 | 0.338 | 0.293 |
| **SL1344_1201** | *SL1344-1201* | 23.706 | 14.582 | 74.892 | 51.69 | 0.317 | 0.282 |
| **SL1344_1202** | *SL1344-1202* | 9.985 | 154.619 | 21.879 | 93.511 | 0.456 | 1.65 |
| **SL1344_1204** | *aroQ* | 15.07 | 48.337 | 22.45 | 38.261 | 0.671 | 1.26 |
| **SL1344_1205** | *yeaS* | 84.855 | 43.857 | 186.22 | 130.354 | 0.456 | 0.336 |

| | | | | | | | |
|---|---|---|---|---|---|---|---|
| **SL1344_1207** | *yoaG* | 74.721 | 173.213 | 85.394 | 105.964 | 0.875 | 1.63 |
| **SL1344_1208** | *SL1344-1208* | 8.022 | 7.629 | 19.205 | 34.385 | 0.418 | 0.222 |
| **SL1344_1210** | *yeaQ* | 310.711 | 603.176 | 172.732 | 227.902 | 1.8 | 2.65 |
| **SL1344_1211** | *yaoF* | 24.899 | 142.251 | 64.288 | 98.25 | 0.387 | 1.45 |
| **SL1344_1212** | *SL1344-1212* | 35.099 | 17.005 | 157.152 | 146.778 | 0.223 | 0.116 |
| **SL1344_1213** | *yeaO* | 52.05 | 58.053 | 114.323 | 128.537 | 0.455 | 0.452 |
| **SL1344_1214** | *yeaN* | 2.997 | 4.305 | 14.216 | 13.385 | 0.211 | 0.322 |
| **SL1344_1215** | *yeaM* | 12.658 | 11.465 | 20.107 | 33.637 | 0.63 | 0.341 |
| **SL1344_1216** | *yeaL* | 26.75 | 17.652 | 28.706 | 22.015 | 0.932 | 0.802 |
| **SL1344_1217** | *yeaK* | 16.652 | 45.872 | 22.099 | 45.749 | 0.754 | 1 |
| **SL1344_1218** | *yeaJ* | 2.932 | 9.971 | 17.657 | 24.017 | 0.166 | 0.415 |
| **SL1344_1219** | *yeaH* | 7.405 | 30.126 | 28.629 | 75.016 | 0.259 | 0.402 |
| **SL1344_1220** | *yeaG* | 22.252 | 68.364 | 23.091 | 64.084 | 0.964 | 1.07 |
| **SL1344_1221** | *mipA* | 1441.664 | 1314.896 | 1134.558 | 1026.322 | 1.27 | 1.28 |
| **SL1344_1223** | *SL1344-1223* | 97.76 | 83.798 | 43.277 | 52.275 | 2.26 | 1.6 |
| **SL1344_1224** | *yeaD* | 190.559 | 93.155 | 62.693 | 50.98 | 3.04 | 1.83 |
| **SL1344_1225** | *gapA* | 4862.734 | 3754.213 | 2746.215 | 2054.253 | 1.77 | 1.83 |
| **SL1344_1226** | *yeaA* | 505.552 | 673.472 | 511.149 | 755.97 | 0.989 | 0.891 |
| **SL1344_1227** | *yeaC* | 179.917 | 908.169 | 336.065 | 688.88 | 0.535 | 1.32 |
| **SL1344_1228** | *nam* | 45.229 | 30.232 | 64.919 | 55.608 | 0.697 | 0.544 |
| **SL1344_1229** | *ansA* | 22.642 | 17.524 | 33.128 | 29.219 | 0.683 | 0.6 |
| **SL1344_1230** | *sppA* | 56.102 | 77.325 | 97.663 | 80.817 | 0.574 | 0.957 |
| **SL1344_1231** | *ydjA* | 177.047 | 743.798 | 224.401 | 186.028 | 0.789 | 4 |
| **SL1344_1232** | *selD* | 121.507 | 102.488 | 173.579 | 151.118 | 0.7 | 0.678 |

| | | | | | | | |
|---|---|---|---|---|---|---|---|
| **SL1344_1233** | *topB* | 20.217 | 16.22 | 68.007 | 79.046 | 0.297 | 0.205 |
| **SL1344_1234** | *gdhA* | 59.106 | 79.611 | 53.016 | 63.595 | 1.11 | 1.25 |
| **SL1344_1235** | *SL1344-1235* | 246.214 | 11.09 | 401.792 | 22.252 | 0.613 | 0.498 |
| **SL1344_1237** | *xthA* | 151.03 | 234.155 | 134.573 | 129.523 | 1.12 | 1.81 |
| **SL1344_1238** | *astC* | 134.791 | 434.405 | 66.835 | 247.345 | 2.02 | 1.76 |
| **SL1344_1239** | *astA* | 20.647 | 56.257 | 28.189 | 83.82 | 0.732 | 0.671 |
| **SL1344_1240** | *astD* | 33.019 | 35.367 | 32.619 | 67.632 | 1.01 | 0.523 |
| **SL1344_1241** | *astB* | 16.208 | 16.904 | 19.521 | 34.469 | 0.83 | 0.49 |
| **SL1344_1242** | *astE* | 6.964 | 5.405 | 26.41 | 39.949 | 0.264 | 0.135 |
| **SL1344_1243** | *SL1344-1243* | 14.738 | 41.848 | 24.78 | 36.467 | 0.595 | 1.15 |
| **SL1344_1244** | *SL1344-1244* | 19.664 | 5.63 | 41.368 | 32.54 | 0.475 | 0.173 |
| **SL1344_1245** | *nadE* | 292.826 | 364.752 | 182.499 | 159.966 | 1.6 | 2.28 |
| **SL1344_1246** | *osmE* | 429.419 | 2170.912 | 543.496 | 977.334 | 0.79 | 2.22 |
| **SL1344_1247** | *celA* | 115.438 | 342.649 | 156.096 | 218.363 | 0.74 | 1.57 |
| **SL1344_1248** | *celB* | 4.922 | 5.806 | 14.084 | 21.67 | 0.349 | 0.268 |
| **SL1344_1249** | *celC* | 87.771 | 187.254 | 89.604 | 120.616 | 0.98 | 1.55 |
| **SL1344_1250** | *celD* | 4.643 | 8.625 | 13.681 | 22.188 | 0.339 | 0.389 |
| **SL1344_1251** | *celF* | 40.773 | 22.323 | 26.855 | 28.316 | 1.52 | 0.788 |
| **SL1344_1252** | *celG* | 28.154 | 21.778 | 57.611 | 54.888 | 0.489 | 0.397 |
| **SL1344_1253** | *katE* | 22.201 | 89.246 | 13.999 | 41.554 | 1.59 | 2.15 |
| **SL1344_1254** | *cedA* | 18.757 | 22.674 | 86.138 | 79.9 | 0.218 | 0.284 |
| **SL1344_1255** | *ydjN* | 70.804 | 353.74 | 101.964 | 293.759 | 0.694 | 1.2 |
| **SL1344_1257** | *yniC* | 269.87 | 214.638 | 193.299 | 139.037 | 1.4 | 1.54 |
| **SL1344_1258** | *yniB* | 33.069 | 51.554 | 83.164 | 65.467 | 0.398 | 0.787 |

| | | | | | | | |
|---|---|---|---|---|---|---|---|
| **SL1344_1259** | *SL1344-1259* | 29.494 | 58.225 | 130.629 | 120.922 | 0.226 | 0.482 |
| **SL1344_1260** | *ydiZ* | 20.749 | 42.776 | 31.285 | 40.713 | 0.663 | 1.05 |
| **SL1344_1261** | *pfkB* | 80.131 | 121.461 | 94.81 | 109.699 | 0.845 | 1.11 |
| **SL1344_1262** | *ydiY* | 46.95 | 21.598 | 95.011 | 50.301 | 0.494 | 0.429 |
| **SL1344_1263** | *SL1344-1263* | 113.764 | 61.077 | 85.274 | 43.9 | 1.33 | 1.39 |
| **SL1344_1265** | *SL1344-1265* | 9.161 | 22.854 | 22.946 | 35.661 | 0.399 | 0.641 |
| **SL1344_1266** | *rfc* | 13.928 | 24.452 | 8.785 | 13.848 | 1.59 | 1.77 |
| **SL1344_1267** | *thrS* | 419.853 | 565.574 | 748.534 | 664.999 | 0.561 | 0.85 |
| **SL1344_1268** | *infC* | 760.37 | 576.991 | 4456.131 | 4322.622 | 0.171 | 0.133 |
| **SL1344_1269** | *rpmI* | 711.528 | 1181.783 | 1730.543 | 2650.879 | 0.411 | 0.446 |
| **SL1344_1270** | *rplT* | 2447.53 | 3254.16 | 7992.819 | 5915.685 | 0.306 | 0.55 |
| **SL1344_1271** | *pheS* | 172.65 | 101.271 | 151.162 | 99.811 | 1.14 | 1.01 |
| **SL1344_1272** | *pheT* | 285.486 | 132.938 | 324.167 | 258.13 | 0.881 | 0.515 |
| **SL1344_1273** | *himA* | 277.758 | 647.292 | 1084.674 | 1323.093 | 0.256 | 0.489 |
| **SL1344_1274** | *btuC* | 18.163 | 8.945 | 68.322 | 53.681 | 0.266 | 0.167 |
| **SL1344_1275** | *btuE* | 43.216 | 107.454 | 38.439 | 40.692 | 1.12 | 2.64 |
| **SL1344_1276** | *btuD* | 27.15 | 17.051 | 34.599 | 33.85 | 0.785 | 0.504 |
| **SL1344_1277** | *nlpC* | 238.56 | 131.215 | 90.645 | 92.088 | 2.63 | 1.42 |
| **SL1344_1278** | *ydiV* | 7.627 | 9.432 | 10.14 | 9.008 | 0.752 | 1.05 |
| **SL1344_1279** | *ydiU* | 27.567 | 15.462 | 57.377 | 46.045 | 0.48 | 0.336 |
| **SL1344_1281** | *aroH* | 43.873 | 35.862 | 61.208 | 49.628 | 0.717 | 0.723 |
| **SL1344_1282** | *ppsR* | 56.355 | 71.773 | 45.039 | 48.619 | 1.25 | 1.48 |
| **SL1344_1283** | *pps* | 931.877 | 663.636 | 615.395 | 593.725 | 1.51 | 1.12 |
| **SL1344_1289** | *ydiP* | 5.647 | 4.046 | 3.459 | 2.415 | 1.63 | 1.68 |

| | | | | | | | |
|---|---|---|---|---|---|---|---|
| **SL1344_1290** | *ydiO* | 5.19 | 9.329 | 21.53 | 29.279 | 0.241 | 0.319 |
| **SL1344_1291** | *ydiF* | 2.745 | 0.597 | 3.818 | 3.257 | 0.719 | 0.183 |
| **SL1344_1292** | *aroD* | 186.786 | 212.488 | 59.784 | 59.635 | 3.12 | 3.56 |
| **SL1344_1297** | *ydiL* | 29.268 | 51.961 | 8.96 | 9.201 | 3.27 | 5.65 |
| **SL1344_1298** | *ydiK* | 41.156 | 35.439 | 55.861 | 52.463 | 0.737 | 0.676 |
| **SL1344_1299** | *ydiJ* | 24.611 | 26.145 | 30.648 | 40.187 | 0.803 | 0.651 |
| **SL1344_1300** | *SL1344-1300* | 67.98 | 83.247 | 70.463 | 107.658 | 0.965 | 0.773 |
| **SL1344_1301** | *ydiH* | 259.641 | 1358.626 | 52.34 | 143.666 | 4.96 | 9.46 |
| **SL1344_1302** | *SL1344-1302* | 15.029 | 24.744 | 15.646 | 24.61 | 0.961 | 1.01 |
| **SL1344_1303** | *sufA* | 15.561 | 16.775 | 6.799 | 15.857 | 2.29 | 1.06 |
| **SL1344_1304** | *sufB* | 6.325 | 6.354 | 19.848 | 12.398 | 0.319 | 0.513 |
| **SL1344_1305** | *sufC* | 21.316 | 11.2 | 17.273 | 14.015 | 1.23 | 0.799 |
| **SL1344_1306** | *sufD* | 9.68 | 5.882 | 12.68 | 12.203 | 0.763 | 0.482 |
| **SL1344_1309** | *ynhG* | 23.985 | 38.559 | 27.972 | 49.178 | 0.857 | 0.784 |
| **SL1344_1310** | *lppB* | 8224.071 | 12903.066 | 1705.472 | 1623.81 | 4.82 | 7.95 |
| **SL1344_1311** | *lppA* | 47685.933 | 96159.097 | 46667.218 | 37294.36 | 1.02 | 2.58 |
| **SL1344_1312** | *pykF* | 217.591 | 175.565 | 203.129 | 113.878 | 1.07 | 1.54 |
| **SL1344_1313** | *orf48* | 3.612 | 6.276 | 4.907 | 8.365 | 0.736 | 0.75 |
| **SL1344_1314** | *orf32* | 1.192 | 7.989 | 2.887 | 5.68 | 0.413 | 1.41 |
| **SL1344_1315** | *orf245* | 31.282 | 86.824 | 9.616 | 27.291 | 3.25 | 3.18 |
| **SL1344_1316** | *orf408* | 3.763 | 13.139 | 2.395 | 11.429 | 1.57 | 1.15 |
| **SL1344_1320** | *ttrS* | 23.292 | 23.018 | 87.197 | 105.862 | 0.267 | 0.217 |
| **SL1344_1321** | *ttrR* | 21.306 | 47.264 | 36.01 | 73.028 | 0.592 | 0.647 |
| **SL1344_1322** | *orf70* | 24.178 | 36.406 | 161.54 | 183.668 | 0.15 | 0.198 |

| | | | | | | | |
|---|---|---|---|---|---|---|---|
| **SL1344_1323** | *orf319* | 24.603 | 98.843 | 75.716 | 120.642 | 0.325 | 0.819 |
| **SL1344_1325** | *ssrB* | 2.131 | 80.157 | 5.497 | 15.173 | 0.388 | 5.28 |
| **SL1344_1326** | *ssrA* | 1.543 | 8.641 | 4.203 | 6.196 | 0.367 | 1.39 |
| **SL1344_1328** | *ssaC* | 0.158 | 5.372 | 0.816 | 3.382 | 0.194 | 1.59 |
| **SL1344_1329** | *ssaD* | 0.391 | 6.286 | 4.14 | 4.489 | 0.0944 | 1.4 |
| **SL1344_1330** | *ssaE* | 2.923 | 16.795 | 14.16 | 15.92 | 0.206 | 1.05 |
| **SL1344_1331** | *sseA* | 3.62 | 70.933 | 7.234 | 58.488 | 0.5 | 1.21 |
| **SL1344_1332** | *sseB* | 3.105 | 16.574 | 6.186 | 28.802 | 0.502 | 0.575 |
| **SL1344_1334** | *sseC* | 0.773 | 9.911 | 2.66 | 13.028 | 0.291 | 0.761 |
| **SL1344_1335** | *sseD* | 6.04 | 28.805 | 5.486 | 23.809 | 1.1 | 1.21 |
| **SL1344_1337** | *sscB* | 0.408 | 16.263 | 2.637 | 8.115 | 0.155 | 2 |
| **SL1344_1340** | *ssaG* | 0 | 37.475 | 3.319 | 56.305 | 0 | 0.666 |
| **SL1344_1342** | *ssaI* | 0.238 | 456.753 | 2.591 | 53.601 | 0.0918 | 8.52 |
| **SL1344_1343** | *ssaJ* | 0.237 | 7.437 | 3.058 | 27.822 | 0.0774 | 0.267 |
| **SL1344_1345** | *ssaK* | 0 | 7.256 | 1.381 | 18.054 | 0 | 0.402 |
| **SL1344_1346** | *ssaL* | 0.698 | 22.808 | 10.925 | 29.552 | 0.0639 | 0.772 |
| **SL1344_1348** | *ssaV* | 0.608 | 7.248 | 6.377 | 12.527 | 0.0953 | 0.579 |
| **SL1344_1349** | *ssaN* | 1.318 | 6.426 | 6.717 | 16.212 | 0.196 | 0.396 |
| **SL1344_1352** | *ssaQ* | 7.086 | 15.654 | 19.53 | 23.929 | 0.363 | 0.654 |
| **SL1344_1353** | *ssaR* | 0.548 | 10.497 | 5.089 | 10.634 | 0.108 | 0.987 |
| **SL1344_1357** | *mdtK* | 26.323 | 18.812 | 63.701 | 49.185 | 0.413 | 0.382 |
| **SL1344_1358** | *ribE* | 427.453 | 264.033 | 269.539 | 238.323 | 1.59 | 1.11 |
| **SL1344_1359** | *cfa* | 410.342 | 290.556 | 169.757 | 182.888 | 2.42 | 1.59 |
| **SL1344_1360** | *ydhC* | 86.288 | 10.152 | 65.264 | 16.119 | 1.32 | 0.63 |

| | | | | | | | |
|---|---|---|---|---|---|---|---|
| **SL1344_1361** | *ydhB* | 13.07 | 18.737 | 40.029 | 29.206 | 0.327 | 0.642 |
| **SL1344_1362** | *purR* | 68.829 | 51.911 | 117.936 | 85.309 | 0.584 | 0.609 |
| **SL1344_1362A** | *ynhF* | 1079.311 | 1260.661 | 817.989 | 1002.884 | 1.32 | 1.26 |
| **SL1344_1363** | *sodB* | 1630.286 | 2077.22 | 980.291 | 1757.701 | 1.66 | 1.18 |
| **SL1344_1364** | *ydhO* | 39.532 | 81.785 | 47.286 | 63.6 | 0.836 | 1.29 |
| **SL1344_1365** | *ydhD* | 3590.615 | 2189.972 | 1135.402 | 846.12 | 3.16 | 2.59 |
| **SL1344_1366** | *rnt* | 76.733 | 36.74 | 171.797 | 191.714 | 0.447 | 0.192 |
| **SL1344_1367** | *gloA* | 583.239 | 865.604 | 279.355 | 444.76 | 2.09 | 1.95 |
| **SL1344_1368** | *nemA* | 79.681 | 95.155 | 34.079 | 51.463 | 2.34 | 1.85 |
| **SL1344_1369** | *ydhM* | 7.005 | 4.988 | 17.682 | 20.31 | 0.396 | 0.246 |
| **SL1344_1371** | *ydhF* | 67.311 | 92.667 | 117.874 | 81.513 | 0.571 | 1.14 |
| **SL1344_1372** | *sodCb* | 18.144 | 49.257 | 31.859 | 50.21 | 0.569 | 0.981 |
| **SL1344_1373** | *SL1344-1373* | 1.741 | 2.868 | 8.714 | 9.138 | 0.2 | 0.314 |
| **SL1344_1374** | *ydhJ* | 10.559 | 28.058 | 15.024 | 20.459 | 0.703 | 1.37 |
| **SL1344_1375** | *ydhI* | 24.227 | 27.84 | 31.154 | 31.932 | 0.778 | 0.872 |
| **SL1344_1376** | *slyA* | 469.26 | 547.101 | 332.248 | 416.249 | 1.41 | 1.31 |
| **SL1344_1377** | *slyB* | 1878.662 | 3931.571 | 1184.158 | 1416.166 | 1.59 | 2.78 |
| **SL1344_1378** | *anmK* | 33.237 | 20.309 | 44.658 | 39.393 | 0.744 | 0.516 |
| **SL1344_1379** | *mliC* | 16.144 | 62.662 | 50.396 | 69.972 | 0.32 | 0.896 |
| **SL1344_1380** | *pdxH* | 26.219 | 40.274 | 51.39 | 61.57 | 0.51 | 0.654 |
| **SL1344_1381** | *tyrS* | 378.751 | 174.028 | 472.606 | 319.369 | 0.801 | 0.545 |
| **SL1344_1382** | *pdxY* | 84.77 | 89.352 | 81.758 | 84.668 | 1.04 | 1.06 |
| **SL1344_1383** | *gst* | 176.118 | 464.924 | 82.093 | 150.459 | 2.15 | 3.09 |
| **SL1344_1384** | *tppB* | 515.769 | 400.178 | 441.097 | 336.834 | 1.17 | 1.19 |

| | | | | | | | |
|---|---|---|---|---|---|---|---|
| **SL1344_1385** | *nth* | 47.839 | 39.037 | 61.765 | 41.591 | 0.775 | 0.939 |
| **SL1344_1386** | *rnfE* | 25.113 | 26.109 | 74.27 | 54.987 | 0.338 | 0.475 |
| **SL1344_1387** | *rnfG* | 31.456 | 19.059 | 59.448 | 42.206 | 0.529 | 0.452 |
| **SL1344_1388** | *rnfD* | 17.44 | 4.239 | 31.814 | 15.16 | 0.548 | 0.28 |
| **SL1344_1389** | *rnfC* | 21.286 | 11.337 | 27.368 | 16.798 | 0.778 | 0.675 |
| **SL1344_1390** | *rnfB* | 18.096 | 11.866 | 72.922 | 60.217 | 0.248 | 0.197 |
| **SL1344_1391** | *rnfA* | 31.733 | 40.088 | 230.2 | 173.531 | 0.138 | 0.231 |
| **SL1344_1392** | *ydgK* | 11.275 | 21.131 | 106.956 | 155.49 | 0.105 | 0.136 |
| **SL1344_1393** | *ydgT* | 176.302 | 780.036 | 248.255 | 297.368 | 0.71 | 2.62 |
| **SL1344_1393A** | *blr* | 267.836 | 1448.601 | 228.768 | 255.967 | 1.17 | 5.66 |
| **SL1344_1394** | *ydgJ* | 68.008 | 45.87 | 61.148 | 63.432 | 1.11 | 0.723 |
| **SL1344_1395** | *add* | 281.675 | 214.654 | 160.036 | 128.519 | 1.76 | 1.67 |
| **SL1344_1396** | *ydgA* | 312.447 | 179.181 | 166.502 | 120.381 | 1.88 | 1.49 |
| **SL1344_1397** | *manA* | 148.137 | 106.601 | 150.56 | 146.944 | 0.984 | 0.725 |
| **SL1344_1398** | *fumA* | 1650.362 | 1267.541 | 778.03 | 1013.661 | 2.12 | 1.25 |
| **SL1344_1400** | *fumC* | 148.576 | 227.27 | 107.934 | 270.427 | 1.38 | 0.84 |
| **SL1344_1401** | *tus* | 21.386 | 11.556 | 28.442 | 15.417 | 0.752 | 0.75 |
| **SL1344_1402** | *rstB* | 14.321 | 7.993 | 16.352 | 15.915 | 0.876 | 0.502 |
| **SL1344_1404** | *ompN* | 25.265 | 22.854 | 24.969 | 19.019 | 1.01 | 1.2 |
| **SL1344_1405** | *rstA* | 30.082 | 64.676 | 143.563 | 169.648 | 0.21 | 0.381 |
| **SL1344_1406** | *ydgC* | 10.042 | 16.803 | 42.669 | 32.526 | 0.235 | 0.517 |
| **SL1344_1407** | *ydgI* | 13.482 | 2.754 | 20.266 | 8.112 | 0.665 | 0.34 |
| **SL1344_1408** | *ydgH* | 265.529 | 307.139 | 426.385 | 298.564 | 0.623 | 1.03 |
| **SL1344_1409** | *pntA* | 524.857 | 164.096 | 213.083 | 165.633 | 2.46 | 0.991 |

| | | | | | | | |
|---|---|---|---|---|---|---|---|
| **SL1344_1410** | *pntB* | 219.646 | 165.572 | 166.797 | 169.025 | 1.32 | 0.98 |
| **SL1344_1411** | *tqsA* | 9.78 | 11.304 | 15.306 | 13.479 | 0.639 | 0.839 |
| **SL1344_1412** | *mdtJ* | 124.259 | 203.502 | 122.435 | 103.243 | 1.01 | 1.97 |
| **SL1344_1413** | *mdtI* | 87.357 | 95.642 | 116.649 | 99.353 | 0.749 | 0.963 |
| **SL1344_1414** | *SL1344-1414* | 3.961 | 13.406 | 17.616 | 29.915 | 0.225 | 0.448 |
| **SL1344_1415** | *asr* | 50.679 | 31.505 | 27.919 | 23.33 | 1.82 | 1.35 |
| **SL1344_1416** | *ynfM* | 4.484 | 5.045 | 10.575 | 12.147 | 0.424 | 0.415 |
| **SL1344_1418** | *mlc* | 27.343 | 28.356 | 87.594 | 101.527 | 0.312 | 0.279 |
| **SL1344_1419** | *bioDb* | 29.767 | 41.927 | 36.769 | 65.589 | 0.81 | 0.639 |
| **SL1344_1421** | *SL1344-1421* | 17.362 | 23.739 | 13.039 | 15.109 | 1.33 | 1.57 |
| **SL1344_1422** | *SL1344-1422* | 5.39 | 8.818 | 153.987 | 166.304 | 0.035 | 0.053 |
| **SL1344_1423** | *SL1344-1423* | 3.212 | 10.998 | 5.081 | 8.97 | 0.632 | 1.23 |
| **SL1344_1424** | *SL1344-1424* | 9.991 | 17.412 | 12.502 | 16.663 | 0.799 | 1.04 |
| **SL1344_1425** | *ynfI* | 107.031 | 49.772 | 93.597 | 62.983 | 1.14 | 0.79 |
| **SL1344_1428** | *dmsA2* | 1.361 | 1.564 | 3.355 | 2.779 | 0.406 | 0.563 |
| **SL1344_1429** | *dmsA1* | 1.454 | 5.989 | 4.256 | 7.208 | 0.342 | 0.831 |
| **SL1344_1430** | *ynfD* | 301.719 | 156.295 | 45.469 | 78.718 | 6.64 | 1.99 |
| **SL1344_1431** | *ynfC* | 39.216 | 23.996 | 42.413 | 53.63 | 0.925 | 0.447 |
| **SL1344_1432** | *speG* | 191.617 | 418.555 | 550.469 | 394.102 | 0.348 | 1.06 |
| **SL1344_1433** | *ynfB* | 178.968 | 259.555 | 586.254 | 483.223 | 0.305 | 0.537 |
| **SL1344_1434** | *ynfA* | 10.318 | 20.802 | 41.87 | 25.658 | 0.246 | 0.811 |
| **SL1344_1435** | *rspA* | 83.213 | 99.373 | 34.75 | 35.223 | 2.39 | 2.82 |
| **SL1344_1436** | *rspB* | 32.615 | 26.008 | 17.78 | 22.757 | 1.83 | 1.14 |
| **SL1344_1438** | *ydfI* | 10.451 | 5.61 | 18.862 | 19.992 | 0.554 | 0.281 |

| | | | | | | | |
|---|---|---|---|---|---|---|---|
| **SL1344_1439** | *ydfZ* | 62.967 | 121.038 | 68.521 | 162.496 | 0.919 | 0.745 |
| **SL1344_1440** | *ydfH* | 33.518 | 39.011 | 40.902 | 39.383 | 0.819 | 0.991 |
| **SL1344_1441** | *ydfG* | 95.091 | 157.624 | 208.429 | 153.748 | 0.456 | 1.03 |
| **SL1344_1442** | *dcp* | 54.645 | 31.797 | 48.07 | 45.008 | 1.14 | 0.706 |
| **SL1344_1444** | *ydeJ* | 5.045 | 12.496 | 4.616 | 5.587 | 1.09 | 2.24 |
| **SL1344_1445** | *ydeI* | 27.865 | 107.83 | 9.485 | 24.856 | 2.94 | 4.34 |
| **SL1344_1446** | *ydeE* | 10.015 | 3.893 | 15.206 | 14.064 | 0.659 | 0.277 |
| **SL1344_1447** | *ydeD* | 30.089 | 25.386 | 49.853 | 41.878 | 0.604 | 0.606 |
| **SL1344_1449** | *marA* | 37.034 | 36.801 | 41.172 | 78.922 | 0.9 | 0.466 |
| **SL1344_1450** | *marR* | 5.987 | 15.793 | 205.658 | 178.479 | 0.0291 | 0.0885 |
| **SL1344_1451** | *marC* | 42.307 | 29.108 | 65.764 | 62.444 | 0.643 | 0.466 |
| **SL1344_1452** | *sotB* | 3.777 | 2.913 | 8.908 | 6.111 | 0.424 | 0.477 |
| **SL1344_1453** | *yneJ* | 5.56 | 7.324 | 23.73 | 15.482 | 0.234 | 0.473 |
| **SL1344_1454** | *yneI* | 9.376 | 13.124 | 13.573 | 20.21 | 0.691 | 0.649 |
| **SL1344_1455** | *yneH* | 13.09 | 5.797 | 40.443 | 35.108 | 0.324 | 0.165 |
| **SL1344_1457** | *SL1344-1457* | 4.658 | 4.832 | 13.005 | 11.733 | 0.358 | 0.412 |
| **SL1344_1459** | *SL1344-1459* | 16.821 | 26.887 | 23.542 | 23.727 | 0.715 | 1.13 |
| **SL1344_1462** | *hyaF2* | 33.722 | 23.25 | 27.877 | 23.245 | 1.21 | 1 |
| **SL1344_1463** | *hyaE2* | 13.106 | 12.247 | 17.616 | 18.767 | 0.744 | 0.653 |
| **SL1344_1465** | *hyaD2* | 18.371 | 20.216 | 39.432 | 46.452 | 0.466 | 0.435 |
| **SL1344_1466** | *hyaC2* | 133.267 | 38.856 | 64.361 | 52.615 | 2.07 | 0.739 |
| **SL1344_1467** | *hyaB2* | 137.464 | 29.502 | 36.697 | 43.677 | 3.75 | 0.675 |
| **SL1344_1468** | *hyaA2* | 76.513 | 40.11 | 46.101 | 38.612 | 1.66 | 1.04 |
| **SL1344_1469** | *SL1344-1469* | 21.032 | 30.647 | 12.144 | 16.208 | 1.73 | 1.89 |

| | | | | | | | |
|---|---|---|---|---|---|---|---|
| **SL1344_1470** | *SL1344-1470* | 114.513 | 64.782 | 79.044 | 69.154 | 1.45 | 0.937 |
| **SL1344_1471** | *SL1344-1471* | 210.065 | 60.264 | 19.144 | 27.667 | 11 | 2.18 |
| **SL1344_1472** | *SL1344-1472* | 24.607 | 44.29 | 84.606 | 36.447 | 0.291 | 1.22 |
| **SL1344_1473** | *pqaA* | 0.19 | 3.32 | 2.578 | 3.121 | 0.0737 | 1.06 |
| **SL1344_1475** | *SL1344-1475* | 19.182 | 12.441 | 11.306 | 7.996 | 1.7 | 1.56 |
| **SL1344_1476** | *SL1344-1476* | 33.773 | 72.055 | 42.952 | 49.248 | 0.786 | 1.46 |
| **SL1344_1477** | *SL1344-1477* | 2.302 | 13.163 | 3.252 | 10.388 | 0.708 | 1.27 |
| **SL1344_1478** | *SL1344-1478* | 12.598 | 51.975 | 15.991 | 33.54 | 0.788 | 1.55 |
| **SL1344_1479** | *SL1344-1479* | 10.385 | 26.254 | 21.379 | 36.395 | 0.486 | 0.721 |
| **SL1344_1480** | *SL1344-1480* | 45.406 | 199.42 | 76.578 | 107.105 | 0.593 | 1.86 |
| **SL1344_1482** | *SL1344-1482* | 1.628 | 5.762 | 21.765 | 21.306 | 0.0748 | 0.27 |
| **SL1344_1484** | *SL1344-1484* | 28.73 | 44.281 | 15.246 | 20.602 | 1.88 | 2.15 |
| **SL1344_1485** | *SL1344-1485* | 17.676 | 10.257 | 26.455 | 31.399 | 0.668 | 0.327 |
| **SL1344_1486** | *SL1344-1486* | 2.528 | 7.121 | 7.8 | 7.843 | 0.324 | 0.908 |
| **SL1344_1487** | *SL1344-1487* | 14.712 | 16.058 | 9.355 | 8.542 | 1.57 | 1.88 |
| **SL1344_1488** | *SL1344-1488* | 7.67 | 8.519 | 12.465 | 17.715 | 0.615 | 0.481 |
| **SL1344_1489** | *SL1344-1489* | 1.872 | 3.228 | 5.952 | 8.299 | 0.315 | 0.389 |
| **SL1344_1490** | *SL1344-1490* | 9.153 | 7.85 | 5.582 | 10.024 | 1.64 | 0.783 |
| **SL1344_1491** | *SL1344-1491* | 4.669 | 9.433 | 3.739 | 5.783 | 1.25 | 1.63 |
| **SL1344_1492** | *hdeB* | 47.176 | 84.511 | 18.464 | 15.533 | 2.56 | 5.44 |
| **SL1344_1493** | *osmC* | 12.88 | 56.527 | 11.615 | 51.996 | 1.11 | 1.09 |
| **SL1344_1494** | *yddX* | 34.53 | 62.983 | 23.231 | 37.947 | 1.49 | 1.66 |
| **SL1344_1495** | *sra* | 164.429 | 576.764 | 228.989 | 617.232 | 0.718 | 0.934 |
| **SL1344_1496** | *sfcA* | 172.493 | 146.939 | 149.615 | 134.436 | 1.15 | 1.09 |

| | | | | | | | |
|---|---|---|---|---|---|---|---|
| **SL1344_1497** | *adhP* | 356.337 | 231.313 | 164.426 | 116.757 | 2.17 | 1.98 |
| **SL1344_1498** | *fdnI* | 39.002 | 43.97 | 58.843 | 85.153 | 0.663 | 0.516 |
| **SL1344_1499** | *fdnH* | 32.841 | 42.581 | 76.138 | 95.923 | 0.431 | 0.444 |
| **SL1344_1500** | *fdnG* | 34.083 | 50.592 | 75.705 | 112.739 | 0.45 | 0.449 |
| **SL1344_1502** | *yddG* | 4.951 | 1.567 | 18.768 | 17.76 | 0.264 | 0.0882 |
| **SL1344_1503** | *ompD* | 18300.681 | 18274.919 | 8655.216 | 8389.076 | 2.11 | 2.18 |
| **SL1344_1504** | *SL1344-1504* | 17.986 | 11.26 | 53.316 | 29.991 | 0.337 | 0.375 |
| **SL1344_1505** | *smvA* | 26.494 | 20.342 | 109.019 | 70.522 | 0.243 | 0.288 |
| **SL1344_1506** | *SL1344-1506* | 6.645 | 7.871 | 70.446 | 77.506 | 0.0943 | 0.102 |
| **SL1344_1508** | *narZ* | 4.51 | 4.582 | 7.665 | 9.081 | 0.588 | 0.505 |
| **SL1344_1509** | *narY* | 2.107 | 4.623 | 6.681 | 6.886 | 0.315 | 0.671 |
| **SL1344_1512** | *yddE* | 15.825 | 26.706 | 19.003 | 26.721 | 0.833 | 0.999 |
| **SL1344_1513** | *nhoA* | 36.524 | 31.277 | 54.991 | 32.01 | 0.664 | 0.977 |
| **SL1344_1514** | *steA* | 69.387 | 382.445 | 97.164 | 171.505 | 0.714 | 2.23 |
| **SL1344_1515** | *ansP* | 61.215 | 22.856 | 63.095 | 22.059 | 0.97 | 1.04 |
| **SL1344_1517** | *SL1344-1517* | 22.63 | 8.455 | 34.964 | 20.673 | 0.647 | 0.409 |
| **SL1344_1518** | *yncD* | 24.029 | 19.274 | 8.314 | 11.525 | 2.89 | 1.67 |
| **SL1344_1519** | *yncC* | 5.688 | 7.252 | 6.889 | 11.073 | 0.826 | 0.655 |
| **SL1344_1520** | *yncB* | 73.451 | 111.403 | 68.576 | 79.129 | 1.07 | 1.41 |
| **SL1344_1521** | *yncA* | 84.318 | 90.431 | 81.825 | 87.765 | 1.03 | 1.03 |
| **SL1344_1522** | *ydcZ* | 11.181 | 16.476 | 134.287 | 109.933 | 0.0833 | 0.15 |
| **SL1344_1523** | *ydcY* | 25.803 | 41.278 | 142.448 | 131.95 | 0.181 | 0.313 |
| **SL1344_1524** | *srfA* | 32.871 | 50.037 | 63.809 | 126.829 | 0.515 | 0.395 |
| **SL1344_1525** | *srfB* | 28.347 | 51.484 | 39.88 | 69.966 | 0.711 | 0.736 |

| | | | | | | | |
|---|---|---|---|---|---|---|---|
| **SL1344_1526** | *srfC* | 20.559 | 36.246 | 43.779 | 78.838 | 0.47 | 0.46 |
| **SL1344_1528** | *ydcW* | 18.708 | 53.768 | 464.357 | 134.121 | 0.0403 | 0.401 |
| **SL1344_1529** | *ydcR* | 13.667 | 13.27 | 17.053 | 25.061 | 0.801 | 0.53 |
| **SL1344_1530** | *pdgL* | 9.52 | 55.582 | 8.833 | 22.705 | 1.08 | 2.45 |
| **SL1344_1531** | *ugtL* | 1.038 | 15.684 | 5.929 | 23.815 | 0.175 | 0.659 |
| **SL1344_1534** | *ydcP* | 22.864 | 15.958 | 65.336 | 46.426 | 0.35 | 0.344 |
| **SL1344_1535** | *ydcN* | 17.086 | 14.567 | 47.522 | 52.5 | 0.36 | 0.277 |
| **SL1344_1536** | *SL1344-1536* | 11.146 | 16.789 | 28.211 | 43.19 | 0.395 | 0.389 |
| **SL1344_1537** | *SL1344-1537* | 123.963 | 197.557 | 171.762 | 248.8 | 0.722 | 0.794 |
| **SL1344_1538** | *tehB* | 458.186 | 323.357 | 140.245 | 111.741 | 3.27 | 2.89 |
| **SL1344_1539** | *tehA* | 10.683 | 28.711 | 46.163 | 44.733 | 0.231 | 0.642 |
| **SL1344_1540** | *ydcK* | 4.405 | 14.908 | 10.157 | 14.32 | 0.434 | 1.04 |
| **SL1344_1541** | *rimL* | 80.57 | 43.206 | 27.08 | 38.149 | 2.98 | 1.13 |
| **SL1344_1542** | *SL1344-1542* | 4.814 | 5.349 | 3.78 | 8.384 | 1.27 | 0.638 |
| **SL1344_1544** | *SL1344-1544* | 1.532 | 9.318 | 4.91 | 7.526 | 0.312 | 1.24 |
| **SL1344_1549** | *aac* | 10.271 | 27.177 | 9.82 | 14.187 | 1.05 | 1.92 |
| **SL1344_1552** | *ydcG* | 169.219 | 97.103 | 147.35 | 105.417 | 1.15 | 0.921 |
| **SL1344_1553** | *SL1344-1553* | 43.307 | 37.955 | 56.767 | 61.016 | 0.763 | 0.622 |
| **SL1344_1554** | *SL1344-1554* | 17.133 | 49.852 | 1209.66 | 635.234 | 0.0142 | 0.0785 |
| **SL1344_1555** | *ydcI* | 431.017 | 344.965 | 498.682 | 779.662 | 0.864 | 0.442 |
| **SL1344_1556** | *trg* | 158.106 | 2.259 | 164.043 | 5.056 | 0.964 | 0.447 |
| **SL1344_1557** | *adhC* | 91.093 | 106.013 | 82.766 | 65.989 | 1.1 | 1.61 |
| **SL1344_1559** | *steB* | 1.62 | 16.582 | 12.661 | 13.533 | 0.128 | 1.23 |
| **SL1344_1561** | *sseJ* | 0.386 | 4.712 | 0.935 | 2.187 | 0.413 | 2.15 |

| | | | | | | | |
|---|---|---|---|---|---|---|---|
| **SL1344_1563** | *SL1344-1563* | 6.448 | 6.874 | 5.362 | 5.14 | 1.2 | 1.34 |
| **SL1344_1568** | *SL1344-1568* | 107.318 | 271.059 | 85.739 | 119.472 | 1.25 | 2.27 |
| **SL1344_1569** | *cybB* | 54.401 | 45.988 | 78.838 | 59.195 | 0.69 | 0.777 |
| **SL1344_1570** | *ydcF* | 58.455 | 48.235 | 59.334 | 48.78 | 0.985 | 0.989 |
| **SL1344_1571** | *hrpA* | 61.939 | 32.015 | 53.556 | 53.301 | 1.16 | 0.601 |
| **SL1344_1572** | *acpD* | 27.741 | 41.082 | 21.41 | 22.742 | 1.3 | 1.81 |
| **SL1344_1573** | *SL1344-1573* | 18.942 | 31.139 | 32.337 | 50.238 | 0.586 | 0.62 |
| **SL1344_1574** | *ydbL* | 8.404 | 13.226 | 20.355 | 18.059 | 0.413 | 0.732 |
| **SL1344_1575** | *ynbE* | 40.696 | 75.815 | 60.483 | 53.899 | 0.673 | 1.41 |
| **SL1344_1576** | *ydbH* | 8.171 | 9.725 | 26.015 | 20.667 | 0.314 | 0.471 |
| **SL1344_1577** | *ldhA* | 290.531 | 108.971 | 162.845 | 53.584 | 1.78 | 2.03 |
| **SL1344_1578** | *hslJ* | 43.064 | 23.17 | 48.138 | 17.825 | 0.895 | 1.3 |
| **SL1344_1579** | *SL1344-1579* | 213.72 | 133.495 | 187.666 | 185.371 | 1.14 | 0.72 |
| **SL1344_1581** | *nifJ* | 35.399 | 20.686 | 21.678 | 18.815 | 1.63 | 1.1 |
| **SL1344_1582** | *uspF* | 930.236 | 1449.713 | 351.826 | 961.761 | 2.64 | 1.51 |
| **SL1344_1583** | *SL1344-1583* | 31.256 | 90.093 | 75.701 | 107.343 | 0.413 | 0.839 |
| **SL1344_1585** | *ydaO* | 94.863 | 62.353 | 55.754 | 43.605 | 1.7 | 1.43 |
| **SL1344_1586** | *dbpA* | 17.75 | 3.317 | 30.572 | 17.562 | 0.581 | 0.189 |
| **SL1344_1587** | *zntB* | 28.935 | 56.823 | 48.518 | 46.982 | 0.596 | 1.21 |
| **SL1344_1588** | *SL1344-1588* | 6.048 | 18.728 | 10.551 | 16.538 | 0.573 | 1.13 |
| **SL1344_1589** | *ydaL* | 34.95 | 63.8 | 121.125 | 99.502 | 0.289 | 0.641 |
| **SL1344_1590** | *ogt* | 18.125 | 42.843 | 64.599 | 84.485 | 0.281 | 0.507 |
| **SL1344_1591** | *fnr* | 141.107 | 242.185 | 414.584 | 482.806 | 0.34 | 0.502 |
| **SL1344_1592** | *ydaA* | 164.346 | 261.824 | 125.673 | 216.079 | 1.31 | 1.21 |

| | | | | | | | |
|---|---|---|---|---|---|---|---|
| **SL1344_1593** | *ynaJ* | 473.787 | 944.295 | 493.071 | 606.746 | 0.961 | 1.56 |
| **SL1344_1594** | *ynaI* | 28.966 | 30.122 | 7.154 | 8.434 | 4.05 | 3.57 |
| **SL1344_1595** | *SL1344-1595* | 41.23 | 54.597 | 60.985 | 53.45 | 0.676 | 1.02 |
| **SL1344_1596** | *SL1344-1596* | 47.821 | 65.348 | 48.311 | 51.032 | 0.99 | 1.28 |
| **SL1344_1602** | *SL1344-1602* | 7.233 | 71.994 | 12.817 | 36.605 | 0.564 | 1.97 |
| **SL1344_1603** | *SL1344-1603* | 5.107 | 22.14 | 24.457 | 32.902 | 0.209 | 0.673 |
| **SL1344_1604** | *SL1344-1604* | 33.253 | 21.638 | 35.781 | 33.507 | 0.929 | 0.646 |
| **SL1344_1605** | *SL1344-1605* | 33.991 | 40.394 | 28.914 | 25.669 | 1.18 | 1.57 |
| **SL1344_1606** | *SL1344-1606* | 15.513 | 17.592 | 14.666 | 21.233 | 1.06 | 0.829 |
| **SL1344_1607** | *SL1344-1607* | 8.036 | 10.886 | 15.745 | 18.948 | 0.51 | 0.575 |
| **SL1344_1609** | *mppA* | 57.544 | 115.35 | 50.543 | 83.338 | 1.14 | 1.38 |
| **SL1344_1610** | *ycjI* | 1.868 | 13.903 | 14.455 | 19.104 | 0.129 | 0.728 |
| **SL1344_1611** | *ycjG* | 18.69 | 30.49 | 45.266 | 48.233 | 0.413 | 0.632 |
| **SL1344_1612** | *tpx* | 1703.91 | 6141.635 | 1034.11 | 672.62 | 1.65 | 9.13 |
| **SL1344_1613** | *tyrR* | 67.179 | 59.331 | 108.455 | 280.064 | 0.619 | 0.212 |
| **SL1344_1614** | *ycjF* | 80.709 | 21.329 | 137.562 | 47.743 | 0.587 | 0.447 |
| **SL1344_1615** | *ycjX* | 66.783 | 26.404 | 144.031 | 50.39 | 0.464 | 0.524 |
| **SL1344_1616** | *pspE* | 48.859 | 145.112 | 75.325 | 161.039 | 0.649 | 0.901 |
| **SL1344_1618** | *pspC* | 31.735 | 29.665 | 99.561 | 62.462 | 0.319 | 0.475 |
| **SL1344_1619** | *pspB* | 37.621 | 36.278 | 75.188 | 69.528 | 0.5 | 0.522 |
| **SL1344_1620** | *pspA* | 93.702 | 110.318 | 147.439 | 117.75 | 0.636 | 0.937 |
| **SL1344_1621** | *pspF* | 24.981 | 22.882 | 31.932 | 26.545 | 0.782 | 0.862 |
| **SL1344_1622** | *sapA* | 21.705 | 26.755 | 51.525 | 41.983 | 0.421 | 0.637 |
| **SL1344_1623** | *sapB* | 23.976 | 14.032 | 30.567 | 22.932 | 0.784 | 0.612 |

| | | | | | | | |
|---|---|---|---|---|---|---|---|
| **SL1344_1624** | *sapC* | 29.298 | 15.116 | 50.846 | 35.305 | 0.576 | 0.428 |
| **SL1344_1625** | *sapD* | 39.642 | 17.605 | 40.209 | 30.29 | 0.986 | 0.581 |
| **SL1344_1626** | *sapF* | 52.079 | 22.674 | 21.319 | 18.606 | 2.44 | 1.22 |
| **SL1344_1627** | *SL1344-1627* | 52.84 | 16.237 | 91.933 | 58.637 | 0.575 | 0.277 |
| **SL1344_1628** | *steC* | 2.369 | 17.525 | 4.122 | 6.933 | 0.575 | 2.53 |
| **SL1344_1631** | *fabI* | 1571.764 | 814.962 | 1024.557 | 748.839 | 1.53 | 1.09 |
| **SL1344_1632** | *yciW* | 6.577 | 19.795 | 10.809 | 24.968 | 0.608 | 0.793 |
| **SL1344_1633** | *rnb* | 269.051 | 119.239 | 251.726 | 129.491 | 1.07 | 0.921 |
| **SL1344_1634** | *yciR* | 4.478 | 10.634 | 10.772 | 16.302 | 0.416 | 0.652 |
| **SL1344_1635** | *SL1344-1635* | 30.014 | 181.39 | 69.664 | 163.008 | 0.431 | 1.11 |
| **SL1344_1636** | *yciT* | 184.58 | 153.296 | 104.146 | 109.527 | 1.77 | 1.4 |
| **SL1344_1637** | *osmB* | 192.72 | 259.661 | 88.05 | 82.031 | 2.19 | 3.17 |
| **SL1344_1638** | *yciH* | 13.396 | 21.426 | 60.504 | 49.023 | 0.221 | 0.437 |
| **SL1344_1639** | *pyrF* | 147.585 | 54.657 | 69.45 | 57.826 | 2.13 | 0.945 |
| **SL1344_1640** | *yciM* | 36.599 | 21.759 | 87.164 | 84.019 | 0.42 | 0.259 |
| **SL1344_1641** | *yciS* | 197.889 | 111.167 | 664.639 | 326.61 | 0.298 | 0.34 |
| **SL1344_1642** | *pgpB* | 13.309 | 20.451 | 28.111 | 30.089 | 0.473 | 0.68 |
| **SL1344_1643** | *ribA* | 699.916 | 331.474 | 371.639 | 211.718 | 1.88 | 1.57 |
| **SL1344_1644** | *acnA* | 175.636 | 231.364 | 88.345 | 139.462 | 1.99 | 1.66 |
| **SL1344_1644A** | *ymiA* | 74.796 | 70.658 | 406.763 | 274.027 | 0.184 | 0.258 |
| **SL1344_1645** | *cysB* | 60.591 | 103.881 | 111.385 | 156.431 | 0.544 | 0.664 |
| **SL1344_1646** | *topA* | 350.017 | 120.543 | 353.148 | 185.008 | 0.991 | 0.652 |
| **SL1344_1647** | *yciN* | 38.288 | 283.962 | 440.342 | 587.584 | 0.087 | 0.483 |
| **SL1344_1648** | *sohB* | 114.94 | 105.573 | 121.116 | 116.299 | 0.949 | 0.908 |

| | | | | | | | |
|---|---|---|---|---|---|---|---|
| **SL1344_1649** | *yciK* | 104.95 | 175.677 | 93.32 | 127.113 | 1.12 | 1.38 |
| **SL1344_1650** | *btuR* | 222.955 | 108.88 | 151.13 | 167.861 | 1.48 | 0.649 |
| **SL1344_1651** | *yciL* | 217.249 | 48.842 | 414.308 | 163.842 | 0.524 | 0.298 |
| **SL1344_1652** | *yciO* | 41.846 | 126.294 | 86.69 | 129.655 | 0.483 | 0.974 |
| **SL1344_1653** | *trpH* | 26.309 | 22.905 | 86.557 | 93.809 | 0.304 | 0.244 |
| **SL1344_1654** | *trpE* | 3.674 | 3.83 | 7.843 | 4.966 | 0.468 | 0.771 |
| **SL1344_1655** | *trpD* | 9.68 | 1.364 | 6.153 | 7.348 | 1.57 | 0.186 |
| **SL1344_1656** | *trpC* | 7.231 | 6.257 | 23.842 | 26.99 | 0.303 | 0.232 |
| **SL1344_1657** | *trpB* | 18.591 | 11.109 | 17.291 | 12.94 | 1.08 | 0.859 |
| **SL1344_1658** | *trpA* | 51.052 | 18.375 | 20.608 | 12.134 | 2.48 | 1.51 |
| **SL1344_1663** | *ompW* | 2360.364 | 3293.017 | 618.902 | 1488.492 | 3.81 | 2.21 |
| **SL1344_1664** | *SL1344-1664* | 35.078 | 142.761 | 27.257 | 77.611 | 1.29 | 1.84 |
| **SL1344_1665** | *yciC* | 137.245 | 142.534 | 156.567 | 178.871 | 0.877 | 0.797 |
| **SL1344_1666** | *ispZ* | 71.033 | 81.5 | 161.421 | 161.64 | 0.44 | 0.504 |
| **SL1344_1667** | *yciA* | 93.504 | 83.25 | 333.453 | 119.691 | 0.28 | 0.696 |
| **SL1344_1668** | *tonB* | 13.56 | 9.704 | 28.909 | 15.489 | 0.469 | 0.627 |
| **SL1344_1669** | *yciI* | 1434.218 | 1550.289 | 211.913 | 255.791 | 6.77 | 6.06 |
| **SL1344_1670** | *cls* | 22.243 | 32.265 | 28.458 | 36.79 | 0.782 | 0.877 |
| **SL1344_1671** | *yciU* | 340.099 | 358.658 | 561.087 | 342.24 | 0.606 | 1.05 |
| **SL1344_1672** | *SL1344-1672* | 11.491 | 24.151 | 38.726 | 47.184 | 0.297 | 0.512 |
| **SL1344_1673** | *oppF* | 120.038 | 246.839 | 142.939 | 243.424 | 0.84 | 1.01 |
| **SL1344_1674** | *oppD* | 164.194 | 147.919 | 133.624 | 176.736 | 1.23 | 0.837 |
| **SL1344_1675** | *oppC* | 94.815 | 140.981 | 136.506 | 199.442 | 0.695 | 0.707 |
| **SL1344_1676** | *oppB* | 35.735 | 99.853 | 46.777 | 129.873 | 0.764 | 0.769 |

| | | | | | | | |
|---|---|---|---|---|---|---|---|
| **SL1344_1677** | *oppA* | 183.677 | 657.58 | 122.503 | 313.243 | 1.5 | 2.1 |
| **SL1344_1679** | *ychE* | 7.765 | 10.602 | 14.823 | 10.821 | 0.524 | 0.98 |
| **SL1344_1680** | *adh* | 138.783 | 146.326 | 93.839 | 86.318 | 1.48 | 1.7 |
| **SL1344_1681** | *tdk* | 87.546 | 63.288 | 105.669 | 101.019 | 0.828 | 0.626 |
| **SL1344_1682** | *hns* | 31095.201 | 19760.504 | 9390.375 | 6034.019 | 3.31 | 3.27 |
| **SL1344_1683** | *galU* | 1004.872 | 538.857 | 395.561 | 220.748 | 2.54 | 2.44 |
| **SL1344_1684** | *hnr* | 37.478 | 31.596 | 102.365 | 87.32 | 0.366 | 0.362 |
| **SL1344_1685** | *ychK* | 59.913 | 44.296 | 125.881 | 155.161 | 0.476 | 0.285 |
| **SL1344_1686** | *ychJ* | 39.721 | 65.947 | 120.253 | 119.366 | 0.33 | 0.552 |
| **SL1344_1687** | *purU* | 116.423 | 161.782 | 124.915 | 155.714 | 0.932 | 1.04 |
| **SL1344_1688** | *SL1344-1688* | 2.486 | 2.166 | 8.305 | 16.25 | 0.299 | 0.133 |
| **SL1344_1689** | *narI* | 1.31 | 55.681 | 6.344 | 64.548 | 0.206 | 0.863 |
| **SL1344_1690** | *narJ* | 0.999 | 39.799 | 5.142 | 96.749 | 0.194 | 0.411 |
| **SL1344_1691** | *narH* | 1.464 | 118.329 | 7.98 | 111.937 | 0.184 | 1.06 |
| **SL1344_1692** | *narG* | 0.443 | 113.623 | 4.825 | 98.846 | 0.0918 | 1.15 |
| **SL1344_1693** | *narK* | 1.27 | 237.393 | 5.487 | 228.782 | 0.232 | 1.04 |
| **SL1344_1694** | *narX* | 0.56 | 2.006 | 13.483 | 28.242 | 0.0415 | 0.071 |
| **SL1344_1695** | *narL* | 22.005 | 33.227 | 16.077 | 26.853 | 1.37 | 1.24 |
| **SL1344_1696** | *ychP* | 6.771 | 4.869 | 29.981 | 32.035 | 0.226 | 0.152 |
| **SL1344_1697** | *ychN* | 712.172 | 765.72 | 383.73 | 280.925 | 1.86 | 2.73 |
| **SL1344_1699** | *chaA* | 17.366 | 24.157 | 37.307 | 44.602 | 0.465 | 0.542 |
| **SL1344_1700** | *kdsA* | 659.792 | 632.001 | 221.758 | 253.355 | 2.98 | 2.49 |
| **SL1344_1701** | *ychA* | 54.152 | 63.57 | 137.792 | 128.357 | 0.393 | 0.495 |
| **SL1344_1702** | *sirC* | 8.5 | 5.232 | 88.961 | 70.119 | 0.0955 | 0.0746 |

| | | | | | | | |
|---|---|---|---|---|---|---|---|
| **SL1344_1703** | *hemK* | 17.815 | 5.791 | 74.606 | 39.834 | 0.239 | 0.145 |
| **SL1344_1704** | *prfA* | 65.699 | 50.812 | 69.433 | 57.712 | 0.946 | 0.88 |
| **SL1344_1705** | *hemA* | 64.61 | 51.895 | 197.258 | 263.89 | 0.328 | 0.197 |
| **SL1344_1706** | *hemM* | 37.376 | 89.932 | 50.431 | 53.937 | 0.741 | 1.67 |
| **SL1344_1707** | *ipk* | 47.87 | 27.863 | 121.997 | 137.893 | 0.392 | 0.202 |
| **SL1344_1708** | *prs* | 295.098 | 311.836 | 567.419 | 573.329 | 0.52 | 0.544 |
| **SL1344_1709** | *ychM* | 17.773 | 12.36 | 17.425 | 13.137 | 1.02 | 0.941 |
| **SL1344_1710** | *ychH* | 302.973 | 505.161 | 777.728 | 685.587 | 0.39 | 0.737 |
| **SL1344_1711** | *pth* | 39.463 | 42.557 | 198.876 | 113.909 | 0.198 | 0.374 |
| **SL1344_1712** | *ychF* | 420.54 | 172.669 | 250.893 | 170.403 | 1.68 | 1.01 |
| **SL1344_1713** | *SL1344-1713* | 8.557 | 18.053 | 17.156 | 11.765 | 0.499 | 1.53 |
| **SL1344_1723** | *SL1344-1723* | 3773.351 | 5020.851 | 2060.768 | 2330.43 | 1.83 | 2.15 |
| **SL1344_1724** | *treA* | 107.434 | 132.31 | 43.228 | 63.743 | 2.49 | 2.08 |
| **SL1344_1725** | *ymgE* | 45.498 | 131.241 | 26.987 | 39.634 | 1.69 | 3.31 |
| **SL1344_1726** | *ycgR* | 27.302 | 0 | 90.507 | 5.724 | 0.302 | 0 |
| **SL1344_1727** | *emtA* | 95.756 | 133.042 | 124.041 | 73.722 | 0.772 | 1.8 |
| **SL1344_1728** | *ldcA* | 64.952 | 51.443 | 61.656 | 43.839 | 1.05 | 1.17 |
| **SL1344_1729** | *cvrA* | 91.181 | 37.58 | 88.633 | 53.043 | 1.03 | 0.708 |
| **SL1344_1730** | *alr-a* | 249.213 | 213.209 | 328.566 | 499.583 | 0.758 | 0.427 |
| **SL1344_1731** | *dadA* | 1124.328 | 1130.598 | 815.946 | 1125.665 | 1.38 | 1 |
| **SL1344_1732** | *ycgB* | 10.854 | 61.651 | 10.703 | 30.592 | 1.01 | 2.02 |
| **SL1344_1733** | *fadR* | 83.53 | 142.75 | 217.54 | 155.987 | 0.384 | 0.915 |
| **SL1344_1734** | *nhaB* | 254.478 | 97.123 | 92.609 | 58.358 | 2.75 | 1.66 |
| **SL1344_1735** | *dsbB* | 121.622 | 146.931 | 296.049 | 259.228 | 0.411 | 0.567 |

| | | | | | | | |
|---|---|---|---|---|---|---|---|
| **SL1344_1736** | *SL1344-1736* | 8.654 | 11.735 | 9.851 | 5.797 | 0.878 | 2.02 |
| **SL1344_1737** | *SL1344-1737* | 78.586 | 666.921 | 122.356 | 267.774 | 0.642 | 2.49 |
| **SL1344_1739** | *ycgN* | 125.179 | 77.002 | 205.586 | 200.13 | 0.609 | 0.385 |
| **SL1344_1740** | *ycgM* | 91.662 | 85.542 | 132.506 | 151.667 | 0.692 | 0.564 |
| **SL1344_1741** | *ycgL* | 139.542 | 703.908 | 108.709 | 168.09 | 1.28 | 4.19 |
| **SL1344_1742** | *minC* | 65.381 | 60.143 | 418.357 | 521.86 | 0.156 | 0.115 |
| **SL1344_1743** | *minD* | 1289.826 | 1204.219 | 1325.926 | 1218.547 | 0.973 | 0.988 |
| **SL1344_1744** | *minE* | 298.411 | 1008.09 | 408.623 | 559.101 | 0.73 | 1.8 |
| **SL1344_1745** | *rnd* | 33.638 | 29.669 | 45.056 | 52.559 | 0.747 | 0.564 |
| **SL1344_1747** | *fadD* | 72.852 | 244.409 | 77.721 | 282.015 | 0.937 | 0.867 |
| **SL1344_1748** | *yeaY* | 18.511 | 12.856 | 83.754 | 83.421 | 0.221 | 0.154 |
| **SL1344_1749** | *yeaZ* | 70.506 | 22.283 | 128.639 | 146.08 | 0.548 | 0.153 |
| **SL1344_1750** | *yoaA* | 21.683 | 14.38 | 44.638 | 43.79 | 0.486 | 0.328 |
| **SL1344_1751** | *yoaB* | 1793.675 | 653.398 | 658.658 | 380.414 | 2.72 | 1.72 |
| **SL1344_1752** | *yoaH* | 80.241 | 304.962 | 66.108 | 156.759 | 1.21 | 1.95 |
| **SL1344_1753** | *pabB* | 35.387 | 34.982 | 52.778 | 47.224 | 0.67 | 0.741 |
| **SL1344_1754** | *yeaB* | 17.176 | 19.267 | 24.761 | 35.746 | 0.694 | 0.539 |
| **SL1344_1755** | *sdaA* | 297.013 | 271.487 | 332.843 | 233.002 | 0.892 | 1.17 |
| **SL1344_1757** | *yoaE* | 8.765 | 16.569 | 18.105 | 25.558 | 0.484 | 0.648 |
| **SL1344_1759** | *SL1344-1759* | 550.709 | 705.624 | 520.057 | 670.994 | 1.06 | 1.05 |
| **SL1344_1760** | *SL1344-1760* | 777.951 | 675.287 | 747.711 | 359.09 | 1.04 | 1.88 |
| **SL1344_1761** | *SL1344-1761* | 681.663 | 735.751 | 831.147 | 679.731 | 0.82 | 1.08 |
| **SL1344_1762** | *SL1344-1762* | 30.951 | 30.824 | 12.338 | 18.068 | 2.51 | 1.71 |
| **SL1344_1763** | *yebN* | 64.623 | 59.744 | 170.928 | 195.264 | 0.378 | 0.306 |

| | | | | | | | |
|---|---|---|---|---|---|---|---|
| **SL1344_1764** | *rrmA* | 41.509 | 17.215 | 88.056 | 63.223 | 0.471 | 0.272 |
| **SL1344_1765** | *ftsI2* | 3.322 | 1.987 | 14.246 | 17.559 | 0.233 | 0.113 |
| **SL1344_1766** | *cspC* | 3858.909 | 25118.955 | 3905.731 | 9213.169 | 0.988 | 2.73 |
| **SL1344_1767** | *yobF* | 49.329 | 2588.114 | 1586.497 | 4933.32 | 0.0311 | 0.525 |
| **SL1344_1768** | *SL1344-1768* | 19.115 | 38.971 | 266.822 | 284.101 | 0.0716 | 0.137 |
| **SL1344_1769** | *yobG* | 132.776 | 153.52 | 447.027 | 484.08 | 0.297 | 0.317 |
| **SL1344_1770** | *SL1344-1770* | 27.377 | 31.46 | 67.801 | 118.577 | 0.404 | 0.265 |
| **SL1344_1771** | *kdgR* | 183.712 | 277.238 | 470.831 | 440.897 | 0.39 | 0.629 |
| **SL1344_1772** | *SL1344-1772* | 5.558 | 4.208 | 8.243 | 9.467 | 0.674 | 0.444 |
| **SL1344_1773** | *htpX* | 355.166 | 234.835 | 423.925 | 292.339 | 0.838 | 0.803 |
| **SL1344_1774** | *prc* | 159.989 | 72.271 | 274.77 | 216.713 | 0.582 | 0.333 |
| **SL1344_1775** | *proQ* | 584.275 | 347.532 | 748.973 | 584.726 | 0.78 | 0.594 |
| **SL1344_1776** | *yebR* | 227.974 | 174.297 | 305.605 | 316.598 | 0.746 | 0.551 |
| **SL1344_1777** | *yebS* | 13.139 | 17.482 | 38.522 | 42.351 | 0.341 | 0.413 |
| **SL1344_1778** | *SL1344-1778* | 36.862 | 26.874 | 34.484 | 37.138 | 1.07 | 0.724 |
| **SL1344_1779** | *yebU* | 14.059 | 6.755 | 24.044 | 13.046 | 0.585 | 0.518 |
| **SL1344_1780** | *SL1344-1780* | 2.22 | 38.262 | 13.142 | 57.223 | 0.169 | 0.669 |
| **SL1344_1783** | *SL1344-1783* | 5.381 | 63.487 | 24.329 | 22.86 | 0.221 | 2.78 |
| **SL1344_1784** | *sopE2* | 49.861 | 51.181 | 45.608 | 30.533 | 1.09 | 1.68 |
| **SL1344_1786** | *SL1344-1786* | 23.206 | 41.212 | 33.362 | 47.192 | 0.696 | 0.873 |
| **SL1344_1787** | *SL1344-1787* | 343.75 | 492.668 | 591.048 | 590.309 | 0.582 | 0.835 |
| **SL1344_1797** | *pagM* | 21.349 | 186.222 | 11.36 | 21.008 | 1.88 | 8.86 |
| **SL1344_1799** | *pagK* | 13.841 | 145.857 | 3.566 | 37.05 | 3.88 | 3.94 |
| **SL1344_1807** | *SL1344-1807* | 148.655 | 220.204 | 312.249 | 273.48 | 0.476 | 0.805 |

| | | | | | | | |
|---|---|---|---|---|---|---|---|
| **SL1344_1808** | *SL1344-1808* | 30.746 | 36.262 | 297.209 | 281.617 | 0.103 | 0.129 |
| **SL1344_1809** | *yobA* | 112.796 | 201.921 | 135.089 | 239.313 | 0.835 | 0.844 |
| **SL1344_1810** | *holE* | 237.546 | 277.68 | 323.973 | 202.118 | 0.733 | 1.37 |
| **SL1344_1811** | *SL1344-1811* | 11.983 | 36.133 | 42.661 | 74.487 | 0.281 | 0.485 |
| **SL1344_1812** | *exoX* | 244.145 | 118.818 | 136.701 | 116.743 | 1.79 | 1.02 |
| **SL1344_1813** | *opdB* | 15.606 | 16.972 | 18.41 | 22.918 | 0.848 | 0.741 |
| **SL1344_1814** | *yebE* | 8.879 | 17.005 | 88.953 | 133.496 | 0.0998 | 0.127 |
| **SL1344_1815** | *yebF* | 32.44 | 263.438 | 217.886 | 245.613 | 0.149 | 1.07 |
| **SL1344_1816** | *yebG* | 354.759 | 236.321 | 221.948 | 191.603 | 1.6 | 1.23 |
| **SL1344_1817** | *purT* | 12.803 | 13.097 | 14.714 | 9.864 | 0.87 | 1.33 |
| **SL1344_1818** | *eda* | 346.591 | 619.078 | 263.175 | 234.595 | 1.32 | 2.64 |
| **SL1344_1819** | *edd* | 19.829 | 3.191 | 45.89 | 47.944 | 0.432 | 0.0666 |
| **SL1344_1820** | *zwf* | 652.261 | 334.899 | 282.509 | 192.956 | 2.31 | 1.74 |
| **SL1344_1821** | *hexR* | 48.648 | 55.433 | 78.852 | 96.243 | 0.617 | 0.576 |
| **SL1344_1822** | *pykA* | 1045.316 | 401.717 | 256.73 | 276.724 | 4.07 | 1.45 |
| **SL1344_1823** | *msbB* | 50.729 | 38.909 | 117.113 | 91.541 | 0.433 | 0.425 |
| **SL1344_1824** | *yebA* | 86.46 | 112.905 | 172.638 | 156.448 | 0.501 | 0.722 |
| **SL1344_1825** | *znuA* | 53.582 | 46.841 | 109.6 | 93.771 | 0.489 | 0.5 |
| **SL1344_1826** | *znuC* | 8.3 | 18.085 | 42.669 | 35.021 | 0.195 | 0.516 |
| **SL1344_1827** | *znuB* | 12.426 | 8.741 | 27.725 | 23.717 | 0.448 | 0.369 |
| **SL1344_1828** | *ruvB* | 103.877 | 75.92 | 145.371 | 121.128 | 0.715 | 0.627 |
| **SL1344_1829** | *ruvA* | 236.197 | 136.821 | 198.301 | 147.009 | 1.19 | 0.931 |
| **SL1344_1833** | *ruvC* | 20.412 | 8.47 | 115.628 | 90.138 | 0.177 | 0.094 |
| **SL1344_1834** | *yebC* | 1353.083 | 863.255 | 898.318 | 385.784 | 1.51 | 2.24 |

| | | | | | | | |
|---|---|---|---|---|---|---|---|
| **SL1344_1835** | *ntpA* | 14.505 | 5.706 | 158.242 | 162.26 | 0.0917 | 0.0352 |
| **SL1344_1836** | *aspS* | 580.258 | 180.584 | 433.134 | 212.305 | 1.34 | 0.851 |
| **SL1344_1837** | *yecD* | 123.504 | 87.816 | 73.706 | 71.001 | 1.68 | 1.24 |
| **SL1344_1838** | *yecE* | 10.625 | 10.216 | 20.831 | 24.385 | 0.51 | 0.419 |
| **SL1344_1839** | *yecN* | 30.644 | 39.164 | 120.197 | 121.627 | 0.255 | 0.322 |
| **SL1344_1840** | *yecO* | 98.339 | 34.285 | 161.384 | 99.543 | 0.609 | 0.344 |
| **SL1344_1841** | *yecP* | 35.687 | 19.735 | 76.035 | 41.791 | 0.469 | 0.472 |
| **SL1344_1842** | *cutC* | 24.486 | 18.485 | 33.395 | 39.715 | 0.733 | 0.465 |
| **SL1344_1843** | *yecM* | 118.596 | 26.731 | 68.037 | 48.061 | 1.74 | 0.556 |
| **SL1344_1844** | *argS* | 612.631 | 132.512 | 176.77 | 109.362 | 3.47 | 1.21 |
| **SL1344_1846** | *SL1344-1846* | 12.254 | 6.325 | 36.093 | 36.035 | 0.34 | 0.176 |
| **SL1344_1847** | *flhE* | 30.124 | 63.175 | 108.711 | 130.123 | 0.277 | 0.486 |
| **SL1344_1848** | *flhA* | 26.451 | 36.252 | 38.411 | 69.361 | 0.689 | 0.523 |
| **SL1344_1849** | *flhB* | 18.858 | 34.483 | 92.902 | 193.2 | 0.203 | 0.178 |
| **SL1344_1850** | *cheZ* | 336.536 | 16.663 | 603.809 | 41.685 | 0.557 | 0.4 |
| **SL1344_1851** | *cheY* | 728.546 | 52.324 | 771.242 | 27.465 | 0.945 | 1.91 |
| **SL1344_1852** | *cheB* | 98.77 | 3.628 | 216.28 | 9.533 | 0.457 | 0.381 |
| **SL1344_1853** | *cheR* | 262.381 | 2.04 | 220.508 | 5.689 | 1.19 | 0.359 |
| **SL1344_1854** | *cheM* | 291.983 | 4.216 | 250.202 | 6.838 | 1.17 | 0.617 |
| **SL1344_1855** | *cheW* | 1570.295 | 44.538 | 915.672 | 42.025 | 1.71 | 1.06 |
| **SL1344_1856** | *cheA* | 300.083 | 7.558 | 484.966 | 15.448 | 0.619 | 0.489 |
| **SL1344_1857** | *motB* | 66.641 | 1.902 | 321.882 | 8.476 | 0.207 | 0.224 |
| **SL1344_1858** | *motA* | 713.265 | 6.434 | 647.412 | 18.27 | 1.1 | 0.352 |
| **SL1344_1859** | *flhC* | 175.947 | 226.62 | 706.065 | 543.627 | 0.249 | 0.417 |

| | | | | | | | |
|---|---|---|---|---|---|---|---|
| **SL1344_1860** | *flhD* | 124.46 | 239.722 | 602.265 | 614.184 | 0.207 | 0.39 |
| **SL1344_1861** | *yecG* | 30.908 | 79.437 | 25.9 | 46.949 | 1.19 | 1.69 |
| **SL1344_1862** | *otsA* | 15.319 | 32.432 | 19.206 | 25.675 | 0.798 | 1.26 |
| **SL1344_1863** | *otsB* | 6.184 | 23.604 | 27.283 | 49.711 | 0.227 | 0.475 |
| **SL1344_1864** | *araH* | 9.335 | 7.436 | 12.333 | 15.513 | 0.757 | 0.479 |
| **SL1344_1865** | *ftnB* | 9.396 | 67.886 | 20.623 | 22.068 | 0.456 | 3.08 |
| **SL1344_1866** | *SL1344-1866* | 9.727 | 9.155 | 10.433 | 22.552 | 0.932 | 0.406 |
| **SL1344_1867** | *SL1344-1867* | 194.672 | 191.107 | 88.964 | 141.259 | 2.19 | 1.35 |
| **SL1344_1868** | *ftn* | 662.549 | 1973.163 | 659.259 | 764.886 | 1 | 2.58 |
| **SL1344_1869** | *yecH* | 5.673 | 19.84 | 8.064 | 12.593 | 0.703 | 1.58 |
| **SL1344_1870** | *tyrP* | 27.497 | 9.429 | 42.052 | 15.66 | 0.654 | 0.602 |
| **SL1344_1871** | *yecA* | 298.638 | 240.015 | 495.219 | 446.001 | 0.603 | 0.538 |
| **SL1344_1872** | *SL1344-1872* | 23.6 | 53.572 | 68.872 | 108.489 | 0.343 | 0.494 |
| **SL1344_1873** | *SL1344-1873* | 15.333 | 21.965 | 19.215 | 20.115 | 0.798 | 1.09 |
| **SL1344_1874** | *SL1344-1874* | 43.196 | 32.07 | 138.416 | 84.299 | 0.312 | 0.38 |
| **SL1344_1875** | *pgsA* | 82.376 | 98.129 | 114.772 | 137.234 | 0.718 | 0.715 |
| **SL1344_1876** | *uvrC* | 20.474 | 25.123 | 56.745 | 54.623 | 0.361 | 0.46 |
| **SL1344_1877** | *sirA* | 53.158 | 120.098 | 371.293 | 362.903 | 0.143 | 0.331 |
| **SL1344_1878** | *yecF* | 130.228 | 165.67 | 496.05 | 368.163 | 0.263 | 0.45 |
| **SL1344_1879** | *sdiA* | 129.523 | 33.117 | 85.267 | 33.041 | 1.52 | 1 |
| **SL1344_1880** | *yecC* | 45.909 | 39.747 | 42.839 | 51.055 | 1.07 | 0.779 |
| **SL1344_1881** | *yecS* | 20.262 | 23.589 | 32.252 | 32.491 | 0.628 | 0.726 |
| **SL1344_1882** | *dcyD* | 82.224 | 47.484 | 64.857 | 58.23 | 1.27 | 0.815 |
| **SL1344_1883** | *fliY* | 263.825 | 472.158 | 286.466 | 374.062 | 0.921 | 1.26 |

| | | | | | | | |
|---|---|---|---|---|---|---|---|
| **SL1344_1884** | *fliZ* | 25.951 | 177.324 | 215.181 | 521.816 | 0.121 | 0.34 |
| **SL1344_1885** | *fliA* | 210.14 | 453.853 | 1460.254 | 2228.18 | 0.144 | 0.204 |
| **SL1344_1887** | *fliB* | 107.394 | 191.824 | 60.39 | 62.833 | 1.78 | 3.05 |
| **SL1344_1888** | *fliC* | 7366.636 | 123.334 | 2834.729 | 76.095 | 2.6 | 1.62 |
| **SL1344_1889** | *fliD* | 400.658 | 160.218 | 433.267 | 356.844 | 0.925 | 0.449 |
| **SL1344_1890** | *fliS* | 186.143 | 268.084 | 214.875 | 264.011 | 0.866 | 1.02 |
| **SL1344_1891** | *fliT* | 159.456 | 195.4 | 411.258 | 356.261 | 0.388 | 0.548 |
| **SL1344_1892** | *amyA* | 3.468 | 7.604 | 12.068 | 17.487 | 0.287 | 0.435 |
| **SL1344_1893** | *yedD* | 300.14 | 216.359 | 308.441 | 164.2 | 0.973 | 1.32 |
| **SL1344_1894** | *yedE* | 62.14 | 41.855 | 58.774 | 41.501 | 1.06 | 1.01 |
| **SL1344_1895** | *yedF* | 25.044 | 84.009 | 40.437 | 48.151 | 0.619 | 1.74 |
| **SL1344_1896** | *SL1344-1896* | 32.58 | 104.756 | 20.375 | 32.176 | 1.6 | 3.26 |
| **SL1344_1897** | *fliE* | 112 | 376.168 | 221.878 | 642.773 | 0.505 | 0.585 |
| **SL1344_1898** | *fliF* | 127.815 | 6.992 | 120.367 | 86.386 | 1.06 | 0.0809 |
| **SL1344_1899** | *fliG* | 59.729 | 11.405 | 126.166 | 33.938 | 0.473 | 0.336 |
| **SL1344_1900** | *fliH* | 63.291 | 6.341 | 126.155 | 23.564 | 0.502 | 0.269 |
| **SL1344_1901** | *fliI* | 41.924 | 2.183 | 121.146 | 20.228 | 0.346 | 0.108 |
| **SL1344_1902** | *fliJ* | 17.465 | 11.95 | 55.054 | 22.872 | 0.317 | 0.522 |
| **SL1344_1903** | *fliK* | 85.487 | 9.103 | 69.153 | 12.824 | 1.24 | 0.71 |
| **SL1344_1904** | *fliL* | 57.803 | 118.02 | 155.621 | 276.093 | 0.371 | 0.427 |
| **SL1344_1905** | *fliM* | 103.31 | 101.795 | 143.367 | 190.544 | 0.721 | 0.534 |
| **SL1344_1906** | *fliN* | 616.822 | 287.694 | 673.375 | 485.153 | 0.916 | 0.593 |
| **SL1344_1907** | *fliO* | 26.935 | 17.815 | 105.629 | 156.907 | 0.255 | 0.114 |
| **SL1344_1908** | *fliP* | 7.058 | 8.94 | 47.498 | 51.142 | 0.149 | 0.175 |

| | | | | | | | |
|---|---|---|---|---|---|---|---|
| **SL1344_1909** | *fliQ* | 6.577 | 22.422 | 48.32 | 40.477 | 0.136 | 0.554 |
| **SL1344_1911** | *rcsA* | 3.984 | 7.304 | 6.203 | 4.921 | 0.642 | 1.48 |
| **SL1344_1912** | *dsrB* | 94.407 | 115.462 | 245.195 | 274.895 | 0.385 | 0.42 |
| **SL1344_1913** | *yodD* | 13.76 | 36.696 | 10.375 | 30.966 | 1.33 | 1.19 |
| **SL1344_1914** | *mngB* | 4.425 | 6.419 | 6.149 | 10.46 | 0.72 | 0.614 |
| **SL1344_1915** | *gcpA* | 3.006 | 2.7 | 18.998 | 33.495 | 0.158 | 0.0806 |
| **SL1344_1916** | *SL1344-1916* | 32.461 | 281.228 | 246.268 | 505.155 | 0.132 | 0.557 |
| **SL1344_1917** | *yedI* | 21.614 | 31.475 | 36.628 | 47.377 | 0.59 | 0.664 |
| **SL1344_1918** | *yedA* | 9.641 | 5.539 | 12.453 | 18.193 | 0.774 | 0.304 |
| **SL1344_1919** | *vsr* | 22.999 | 21.23 | 55.399 | 54.107 | 0.415 | 0.392 |
| **SL1344_1920** | *dcm* | 32.348 | 22.008 | 88.715 | 65.203 | 0.365 | 0.338 |
| **SL1344_1921** | *yedJ* | 19.731 | 12.607 | 49.334 | 31.578 | 0.4 | 0.399 |
| **SL1344_1923** | *ompS* | 15.577 | 9.49 | 29.703 | 47.428 | 0.524 | 0.2 |
| **SL1344_1924** | *cspB* | 32.515 | 76.644 | 86.491 | 364.045 | 0.376 | 0.211 |
| **SL1344_1925** | *umuC* | 3.172 | 2.519 | 16.89 | 14.443 | 0.188 | 0.174 |
| **SL1344_1926** | *umuD* | 25.087 | 19.111 | 18.262 | 13.471 | 1.37 | 1.42 |
| **SL1344_1929** | *SL1344-1929* | 81.688 | 6.009 | 88.29 | 12.13 | 0.925 | 0.495 |
| **SL1344_1930** | *SL1344-1930* | 6.046 | 3.921 | 32.294 | 12.998 | 0.187 | 0.302 |
| **SL1344_1933** | *SL1344-1933* | 21.358 | 12.583 | 12.21 | 11.469 | 1.75 | 1.1 |
| **SL1344_1937** | *SL1344-1937* | 6.489 | 3.754 | 15.983 | 11.612 | 0.406 | 0.323 |
| **SL1344_1938** | *SL1344-1938* | 1.473 | 2.257 | 30.992 | 19.391 | 0.0475 | 0.116 |
| **SL1344_1939** | *SL1344-1939* | 88.52 | 33.075 | 64.449 | 30.834 | 1.37 | 1.07 |
| **SL1344_1940** | *SL1344-1940* | 12.436 | 6.097 | 45.38 | 23.569 | 0.274 | 0.259 |
| **SL1344_1941** | *SL1344-1941* | 21.669 | 7.225 | 38.547 | 17.783 | 0.562 | 0.406 |

| | | | | | | | |
|---|---|---|---|---|---|---|---|
| **SL1344_1944** | *SL1344-1944* | 33.909 | 13.604 | 41.417 | 18.567 | 0.819 | 0.733 |
| **SL1344_1945** | *SL1344-1945* | 15.529 | 5.039 | 86.065 | 21.865 | 0.18 | 0.23 |
| **SL1344_1946** | *SL1344-1946* | 48.592 | 7.784 | 42.796 | 17.082 | 1.14 | 0.456 |
| **SL1344_1947** | *SL1344-1947* | 108.768 | 23.238 | 80.064 | 21.813 | 1.36 | 1.07 |
| **SL1344_1948** | *SL1344-1948* | 36.361 | 10.297 | 109.873 | 25.446 | 0.331 | 0.405 |
| **SL1344_1949** | *SL1344-1949* | 9.294 | 2.519 | 59.275 | 49.759 | 0.157 | 0.0506 |
| **SL1344_1950** | *SL1344-1950* | 16.488 | 1.491 | 32.989 | 10.346 | 0.5 | 0.144 |
| **SL1344_1953** | *SL1344-1953* | 51.38 | 30.755 | 72.866 | 37.824 | 0.705 | 0.813 |
| **SL1344_1954** | *SL1344-1954* | 24.664 | 10.203 | 46.082 | 19.401 | 0.535 | 0.526 |
| **SL1344_1956** | *SL1344-1956* | 44.853 | 13.051 | 91.032 | 18.36 | 0.493 | 0.711 |
| **SL1344_1957** | *SL1344-1957* | 10.123 | 5.323 | 73.761 | 24.389 | 0.137 | 0.218 |
| **SL1344_1958** | *SL1344-1958* | 10.688 | 7.369 | 72.878 | 70.253 | 0.147 | 0.105 |
| **SL1344_1959** | *SL1344-1959* | 26.846 | 6.977 | 18.151 | 8.757 | 1.48 | 0.797 |
| **SL1344_1960** | *SL1344-1960* | 3.923 | 1.639 | 17.417 | 9.807 | 0.225 | 0.167 |
| **SL1344_1961** | *SL1344-1961* | 19.209 | 18.695 | 50.637 | 205.867 | 0.379 | 0.0908 |
| **SL1344_1962** | *SL1344-1962* | 4.436 | 4.261 | 17.801 | 8.573 | 0.249 | 0.497 |
| **SL1344_1963** | *SL1344-1963* | 25.191 | 7.815 | 30.738 | 12.755 | 0.82 | 0.613 |
| **SL1344_1964** | *SL1344-1964* | 11.564 | 5.98 | 13.041 | 8.089 | 0.887 | 0.739 |
| **SL1344_1965** | *SL1344-1965* | 5.826 | 7.595 | 2330.635 | 1513.847 | 0.0025 | 0.00502 |
| **SL1344_1966** | *SL1344-1966* | 23.145 | 8.323 | 57.266 | 24.961 | 0.404 | 0.333 |
| **SL1344_1967** | *SL1344-1967* | 91.853 | 331.799 | 293.532 | 430.043 | 0.313 | 0.772 |
| **SL1344_1968** | *SL1344-1968* | 62.199 | 67.446 | 378.067 | 165.652 | 0.165 | 0.407 |
| **SL1344_1969** | *SL1344-1969* | 135.975 | 80.015 | 87.527 | 49.292 | 1.55 | 1.62 |
| **SL1344_1970** | *SL1344-1970* | 15.127 | 10.077 | 11.151 | 5.463 | 1.36 | 1.84 |

| | | | | | | | |
|---|---|---|---|---|---|---|---|
| **SL1344_1972** | *SL1344-1972* | 14.463 | 13.868 | 15.775 | 13.889 | 0.917 | 0.999 |
| **SL1344_1973** | *SL1344-1973* | 26.758 | 24.223 | 17.018 | 15.887 | 1.57 | 1.52 |
| **SL1344_1974** | *SL1344-1974* | 9.866 | 16.946 | 13.582 | 12.217 | 0.726 | 1.39 |
| **SL1344_1976** | *SL1344-1976* | 5.8 | 3.848 | 8.834 | 8.597 | 0.657 | 0.448 |
| **SL1344_1977** | *yeeI* | 151.324 | 257.339 | 99.531 | 230.759 | 1.52 | 1.12 |
| **SL1344_1982** | *SL1344-1982* | 4.639 | 3.262 | 4.594 | 6.911 | 1.01 | 0.472 |
| **SL1344_1983** | *SL1344-1983* | 1.06 | 2.043 | 3.189 | 4.568 | 0.332 | 0.447 |
| **SL1344_1986** | *amn* | 161.961 | 145.767 | 100.997 | 106.553 | 1.6 | 1.37 |
| **SL1344_1987** | *SL1344-1987* | 2.172 | 10.817 | 34.636 | 74.459 | 0.0627 | 0.145 |
| **SL1344_1988** | *SL1344-1988* | 8.26 | 3.164 | 83.353 | 68.038 | 0.0991 | 0.0465 |
| **SL1344_1991** | *erfK* | 5.665 | 19.09 | 17.728 | 28.417 | 0.32 | 0.672 |
| **SL1344_1992** | *cobT* | 13.818 | 11.94 | 21.418 | 27.633 | 0.645 | 0.432 |
| **SL1344_1993** | *cobS* | 49.647 | 16.822 | 53.474 | 42.443 | 0.928 | 0.396 |
| **SL1344_1994** | *cobU* | 16.587 | 10.465 | 26.126 | 36.268 | 0.635 | 0.289 |
| **SL1344_1995** | *cbiP* | 13.894 | 13.193 | 21.632 | 28.137 | 0.642 | 0.469 |
| **SL1344_1996** | *cbiO* | 7.327 | 9.503 | 29.605 | 28.327 | 0.247 | 0.335 |
| **SL1344_1997** | *cbiQ* | 3.056 | 7.123 | 26.326 | 58.022 | 0.116 | 0.123 |
| **SL1344_1999** | *cbiM* | 25.186 | 17.236 | 23.409 | 28.503 | 1.08 | 0.605 |
| **SL1344_2000** | *cbiL* | 11.192 | 23.055 | 11.144 | 19.777 | 1 | 1.17 |
| **SL1344_2001** | *cbiK* | 14.743 | 28.492 | 15.058 | 19.951 | 0.979 | 1.43 |
| **SL1344_2003** | *cbiH* | 21.525 | 25.203 | 25.376 | 34.137 | 0.848 | 0.738 |
| **SL1344_2004** | *cbiG* | 2.691 | 10.628 | 10.318 | 20.95 | 0.261 | 0.507 |
| **SL1344_2005** | *cbiF* | 19.502 | 29.177 | 22.783 | 41.735 | 0.856 | 0.699 |
| **SL1344_2006** | *cbiT* | 16.46 | 15.038 | 34.542 | 43.388 | 0.477 | 0.347 |

| | | | | | | | |
|---|---|---|---|---|---|---|---|
| **SL1344_2007** | *cbiE* | 7.228 | 11.225 | 17.507 | 37.425 | 0.413 | 0.3 |
| **SL1344_2008** | *cbiD* | 5.504 | 9.308 | 18.361 | 33.405 | 0.3 | 0.279 |
| **SL1344_2009** | *cbiC* | 8.51 | 21.169 | 14.609 | 31.704 | 0.583 | 0.668 |
| **SL1344_2010** | *cbiB* | 6.413 | 13.321 | 27.852 | 48.811 | 0.23 | 0.273 |
| **SL1344_2011** | *cbiA* | 8.193 | 20.998 | 31.53 | 73.64 | 0.26 | 0.285 |
| **SL1344_2012** | *pocR* | 40.566 | 83.684 | 44.252 | 40.537 | 0.917 | 2.06 |
| **SL1344_2013** | *pduF* | 3.127 | 6.588 | 16.411 | 17.792 | 0.191 | 0.37 |
| **SL1344_2015** | *pduB* | 13.761 | 0.586 | 20.809 | 2.468 | 0.661 | 0.237 |
| **SL1344_2016** | *pduC* | 12.017 | 0.449 | 11.323 | 1.089 | 1.06 | 0.413 |
| **SL1344_2017** | *pduD* | 5.613 | 0.504 | 8.39 | 1.719 | 0.669 | 0.293 |
| **SL1344_2018** | *pduE* | 15.762 | 0.521 | 18.264 | 1.76 | 0.863 | 0.296 |
| **SL1344_2021** | *pduJ* | 32.6 | 2.465 | 20.518 | 4.293 | 1.59 | 0.574 |
| **SL1344_2026** | *pduO* | 5.328 | 1.009 | 40.486 | 2.057 | 0.132 | 0.491 |
| **SL1344_2027** | *pduP* | 15.955 | 3.218 | 5.447 | 1.404 | 2.93 | 2.29 |
| **SL1344_2030** | *pduT* | 7.359 | 0 | 6.587 | 2.875 | 1.12 | 0 |
| **SL1344_2033** | *pduW* | 3.946 | 1.176 | 4.248 | 2.786 | 0.929 | 0.422 |
| **SL1344_2035** | *yeeX* | 275.712 | 629.399 | 451.863 | 454.648 | 0.61 | 1.38 |
| **SL1344_2036** | *yeeA* | 7.993 | 8.992 | 58.078 | 67.011 | 0.138 | 0.134 |
| **SL1344_2037** | *gyrI* | 534.394 | 805.209 | 694.626 | 862.636 | 0.769 | 0.933 |
| **SL1344_2038** | *dacD* | 9.588 | 4.349 | 11.489 | 9.523 | 0.835 | 0.457 |
| **SL1344_2039** | *phsC* | 62.909 | 10.314 | 47.321 | 15.582 | 1.33 | 0.662 |
| **SL1344_2040** | *phsB* | 27.501 | 5.522 | 30.456 | 12.904 | 0.903 | 0.428 |
| **SL1344_2041** | *phsA* | 50.174 | 13.801 | 23.328 | 10.247 | 2.15 | 1.35 |
| **SL1344_2043** | *sopA* | 49.442 | 27.162 | 12.115 | 8.235 | 4.08 | 3.3 |

| | | | | | | | |
|---|---|---|---|---|---|---|---|
| **SL1344_2044** | *sbcB* | 27.674 | 37.124 | 16.631 | 25.193 | 1.66 | 1.47 |
| **SL1344_2045** | *yeeF* | 611.111 | 71.559 | 331.11 | 86.619 | 1.85 | 0.826 |
| **SL1344_2046** | *yeeY* | 1.488 | 7.88 | 20.526 | 34.643 | 0.0725 | 0.227 |
| **SL1344_2047** | *yeeZ* | 436.245 | 117.409 | 127.727 | 110.109 | 3.42 | 1.07 |
| **SL1344_2048** | *hisG* | 44.988 | 36.202 | 42.214 | 34.576 | 1.07 | 1.05 |
| **SL1344_2049** | *hisD* | 76.885 | 50.143 | 42.626 | 28.644 | 1.8 | 1.75 |
| **SL1344_2050** | *hisC* | 58.866 | 32.31 | 26.085 | 19.948 | 2.26 | 1.62 |
| **SL1344_2051** | *hisB* | 74.381 | 69.996 | 68.663 | 58.251 | 1.08 | 1.2 |
| **SL1344_2052** | *hisH* | 76.335 | 53.02 | 84.717 | 48.928 | 0.901 | 1.08 |
| **SL1344_2053** | *hisA* | 61.761 | 19.448 | 70.421 | 45.114 | 0.877 | 0.431 |
| **SL1344_2054** | *hisF* | 40.91 | 51.826 | 55.815 | 49.136 | 0.733 | 1.05 |
| **SL1344_2055** | *hisI* | 110.941 | 60.797 | 61.376 | 60.21 | 1.81 | 1.01 |
| **SL1344_2056** | *wzzB* | 122.84 | 202.958 | 254.098 | 295.748 | 0.483 | 0.686 |
| **SL1344_2057** | *udg* | 34.593 | 46.338 | 41.831 | 68.538 | 0.827 | 0.676 |
| **SL1344_2058** | *gnd* | 686.646 | 530.73 | 505.453 | 399.2 | 1.36 | 1.33 |
| **SL1344_2059** | *rfbP* | 212.454 | 96.637 | 220.361 | 124.121 | 0.964 | 0.779 |
| **SL1344_2060** | *rfbK* | 257.252 | 241.442 | 126.171 | 134.855 | 2.04 | 1.79 |
| **SL1344_2061** | *cpsB2* | 183.256 | 122.911 | 142.67 | 126.015 | 1.28 | 0.975 |
| **SL1344_2062** | *rfbN* | 40.465 | 97.965 | 93.378 | 173.525 | 0.433 | 0.565 |
| **SL1344_2063** | *rfbU* | 86.06 | 133.352 | 137.697 | 131.572 | 0.625 | 1.01 |
| **SL1344_2064** | *rfbV* | 31.724 | 85.671 | 60.166 | 104.799 | 0.527 | 0.817 |
| **SL1344_2065** | *rfbX* | 63.223 | 39.561 | 81.441 | 68.927 | 0.776 | 0.574 |
| **SL1344_2066** | *rfbJ* | 196.196 | 134.077 | 170.209 | 168.822 | 1.15 | 0.794 |
| **SL1344_2067** | *rfbH* | 1826.421 | 1202.227 | 998.224 | 730.425 | 1.83 | 1.65 |

| | | | | | | | |
|---|---|---|---|---|---|---|---|
| **SL1344_2068** | *rfbG* | 486.928 | 262.323 | 495.413 | 417.98 | 0.983 | 0.628 |
| **SL1344_2069** | *rfbF* | 492.063 | 400.306 | 502.619 | 364.463 | 0.979 | 1.1 |
| **SL1344_2070** | *rfbI* | 405.002 | 156.661 | 380.363 | 222.625 | 1.06 | 0.704 |
| **SL1344_2071** | *rfbC* | 458.756 | 214.415 | 345.951 | 179.458 | 1.33 | 1.19 |
| **SL1344_2072** | *rfbA* | 79.667 | 64.462 | 130.808 | 104.775 | 0.609 | 0.615 |
| **SL1344_2073** | *rfbD* | 109.246 | 59.027 | 275.345 | 211.081 | 0.397 | 0.28 |
| **SL1344_2074** | *rfbB* | 406.021 | 176.505 | 398.484 | 277.322 | 1.02 | 0.636 |
| **SL1344_2075** | *galF* | 175.616 | 177.467 | 395.696 | 296.906 | 0.444 | 0.598 |
| **SL1344_2078** | *wcaK* | 3.004 | 1.54 | 4.197 | 3.228 | 0.716 | 0.477 |
| **SL1344_2082** | *cpsB* | 3.337 | 0.379 | 5.038 | 3.567 | 0.662 | 0.106 |
| **SL1344_2086** | *gmd* | 6.595 | 0.485 | 4.728 | 3.793 | 1.39 | 0.128 |
| **SL1344_2096** | *yegH* | 26.359 | 20.738 | 33.371 | 34.915 | 0.79 | 0.594 |
| **SL1344_2097** | *asmA* | 34.267 | 31.978 | 123.835 | 126.454 | 0.277 | 0.253 |
| **SL1344_2098** | *dcd* | 151.647 | 113.486 | 246.951 | 171.994 | 0.614 | 0.66 |
| **SL1344_2099** | *udk* | 60.577 | 61.452 | 67.441 | 84.362 | 0.898 | 0.728 |
| **SL1344_2100** | *yegE* | 3.503 | 3.252 | 10.066 | 12.958 | 0.348 | 0.251 |
| **SL1344_2101** | *alkA* | 5.987 | 5.004 | 15.161 | 22.9 | 0.395 | 0.219 |
| **SL1344_2102** | *yegD* | 31.807 | 6.234 | 26.808 | 11.151 | 1.19 | 0.559 |
| **SL1344_2103** | *SL1344-2103* | 4.814 | 3.122 | 8.657 | 16.859 | 0.556 | 0.185 |
| **SL1344_2104** | *yegN* | 0.967 | 1.002 | 5.715 | 6.682 | 0.169 | 0.15 |
| **SL1344_2106** | *yegB* | 5.823 | 3.899 | 16.792 | 15.674 | 0.347 | 0.249 |
| **SL1344_2107** | *baeS* | 4.174 | 4.263 | 12.1 | 13.76 | 0.345 | 0.31 |
| **SL1344_2108** | *baeR* | 27.018 | 41.584 | 42.336 | 37.522 | 0.638 | 1.11 |
| **SL1344_2111** | *SL1344-2111* | 2.943 | 2.126 | 9.537 | 6.949 | 0.309 | 0.306 |

| | | | | | | | |
|---|---|---|---|---|---|---|---|
| **SL1344_2112** | *yegQ* | 35.812 | 15.282 | 29.368 | 18.321 | 1.22 | 0.834 |
| **SL1344_2114** | *srcA* | 3.123 | 17.617 | 5.157 | 12.408 | 0.606 | 1.42 |
| **SL1344_2115** | *steD* | 11.275 | 14.981 | 23.895 | 42.025 | 0.472 | 0.356 |
| **SL1344_2117** | *yegS* | 10.458 | 24.11 | 16.806 | 36.725 | 0.622 | 0.656 |
| **SL1344_2118** | *fbaB* | 30.974 | 112.464 | 16.815 | 49.552 | 1.84 | 2.27 |
| **SL1344_2119** | *yegT* | 17.451 | 25.668 | 26.825 | 33.113 | 0.651 | 0.775 |
| **SL1344_2120** | *yegU* | 10.896 | 11.98 | 28.388 | 31.324 | 0.384 | 0.382 |
| **SL1344_2121** | *yegV* | 10.931 | 11.692 | 18.4 | 24.291 | 0.594 | 0.481 |
| **SL1344_2122** | *yegW* | 11.411 | 34.603 | 56.042 | 53.213 | 0.204 | 0.65 |
| **SL1344_2123** | *thiD* | 8.203 | 3.142 | 16.467 | 15.365 | 0.498 | 0.205 |
| **SL1344_2124** | *thiM* | 22.773 | 8.524 | 30.901 | 24.815 | 0.737 | 0.343 |
| **SL1344_2125** | *SL1344-2125* | 35.141 | 69.533 | 53.251 | 74.532 | 0.66 | 0.933 |
| **SL1344_2129** | *stcA* | 10.813 | 46.5 | 5.4 | 9.244 | 2 | 5.03 |
| **SL1344_2131** | *mrp* | 416.12 | 517.084 | 306.689 | 328.2 | 1.36 | 1.58 |
| **SL1344_2132** | *metG* | 203.135 | 158.75 | 149.358 | 102.695 | 1.36 | 1.55 |
| **SL1344_2133** | *yehR* | 7.738 | 10.966 | 12.025 | 9.482 | 0.643 | 1.16 |
| **SL1344_2135** | *yehS* | 153.045 | 28.488 | 73.981 | 64.27 | 2.07 | 0.443 |
| **SL1344_2136** | *yehT* | 154.481 | 66.415 | 136 | 64.544 | 1.14 | 1.03 |
| **SL1344_2137** | *yehU* | 49.153 | 14.201 | 131.845 | 69.554 | 0.373 | 0.204 |
| **SL1344_2138** | *yehV* | 2.264 | 14.961 | 14.493 | 44.328 | 0.156 | 0.338 |
| **SL1344_2139** | *yohO* | 566.731 | 377.896 | 252.22 | 248.952 | 2.25 | 1.52 |
| **SL1344_2141** | *yehX* | 10.927 | 17.508 | 22.004 | 40.63 | 0.497 | 0.431 |
| **SL1344_2142** | *yehY* | 3.44 | 6.163 | 27.387 | 54.909 | 0.126 | 0.112 |
| **SL1344_2143** | *yehZ* | 70.672 | 90.473 | 23.66 | 68.138 | 2.99 | 1.33 |

| | | | | | | | |
|---|---|---|---|---|---|---|---|
| **SL1344_2144** | *bglX* | 110.996 | 94.513 | 79.42 | 58.353 | 1.4 | 1.62 |
| **SL1344_2145** | *dld* | 49.448 | 68.964 | 39.797 | 60.007 | 1.24 | 1.15 |
| **SL1344_2146** | *pbpG* | 23.443 | 31.728 | 55.36 | 68.467 | 0.423 | 0.463 |
| **SL1344_2149** | *yohF* | 3.806 | 8.48 | 11.195 | 13.327 | 0.34 | 0.636 |
| **SL1344_2151** | *yohI* | 21.434 | 3.622 | 63.515 | 31.234 | 0.337 | 0.116 |
| **SL1344_2151A** | *yohP* | 78.926 | 133.613 | 69.977 | 101.321 | 1.13 | 1.32 |
| **SL1344_2152** | *SL1344-2152* | 18.889 | 15.496 | 19.452 | 23.632 | 0.971 | 0.656 |
| **SL1344_2153** | *SL1344-2153* | 5.69 | 3.48 | 14.892 | 13.795 | 0.382 | 0.252 |
| **SL1344_2154** | *SL1344-2154* | 20.912 | 12.79 | 14.909 | 25.66 | 1.4 | 0.498 |
| **SL1344_2155** | *gtdA* | 7.357 | 8.06 | 5.11 | 9.97 | 1.44 | 0.808 |
| **SL1344_2157** | *SL1344-2157* | 13.61 | 20.055 | 27.443 | 32.238 | 0.496 | 0.622 |
| **SL1344_2158** | *yohJ* | 12.462 | 28.129 | 24.254 | 34.784 | 0.514 | 0.809 |
| **SL1344_2159** | *yohK* | 10.546 | 14.269 | 11.741 | 11.638 | 0.898 | 1.23 |
| **SL1344_2160** | *cdd* | 984.366 | 94.615 | 1007.215 | 92.863 | 0.977 | 1.02 |
| **SL1344_2161** | *sanA* | 52.535 | 46.009 | 83.73 | 109.846 | 0.627 | 0.419 |
| **SL1344_2162** | *b2145* | 11.309 | 205.17 | 18.941 | 41.183 | 0.597 | 4.98 |
| **SL1344_2163** | *SL1344-2163* | 13.583 | 25.193 | 31.571 | 22.777 | 0.43 | 1.11 |
| **SL1344_2164** | *yeiA* | 85.439 | 45.843 | 44.077 | 33.96 | 1.94 | 1.35 |
| **SL1344_2165** | *mglC* | 811.039 | 692.39 | 736.405 | 815.623 | 1.1 | 0.849 |
| **SL1344_2166** | *mglA* | 344.736 | 570.108 | 610.889 | 952.894 | 0.564 | 0.598 |
| **SL1344_2167** | *mglB* | 9133.294 | 8189.718 | 2147.35 | 2689.731 | 4.25 | 3.04 |
| **SL1344_2168** | *galS* | 177.988 | 316.768 | 232.218 | 460.627 | 0.766 | 0.688 |
| **SL1344_2169** | *yeiB* | 4.35 | 3.868 | 37.127 | 38.415 | 0.117 | 0.101 |
| **SL1344_2170** | *folE* | 109.894 | 132.484 | 107.043 | 100.908 | 1.03 | 1.31 |

| | | | | | | | |
|---|---|---|---|---|---|---|---|
| **SL1344_2171** | *yeiG* | 108.04 | 270.975 | 142.532 | 189.993 | 0.758 | 1.43 |
| **SL1344_2172** | *SL1344-2172* | 4.553 | 9.418 | 73.889 | 60.447 | 0.0616 | 0.156 |
| **SL1344_2173** | *SL1344-2173* | 4.5 | 1.492 | 14.777 | 15.518 | 0.305 | 0.0961 |
| **SL1344_2177** | *lysP* | 149.436 | 48.633 | 128.543 | 79.889 | 1.16 | 0.609 |
| **SL1344_2178** | *yeiE* | 50.973 | 41.49 | 81.971 | 80.932 | 0.622 | 0.513 |
| **SL1344_2179** | *yeiH* | 25.595 | 21.184 | 43.215 | 28.14 | 0.592 | 0.753 |
| **SL1344_2180** | *nfo* | 49.191 | 68.021 | 53.637 | 47.371 | 0.917 | 1.44 |
| **SL1344_2181** | *fruA* | 4.907 | 3.826 | 10.398 | 11.91 | 0.472 | 0.321 |
| **SL1344_2183** | *fruB* | 7.798 | 13.713 | 13.183 | 11.865 | 0.592 | 1.16 |
| **SL1344_2187** | *SL1344-2187* | 33.195 | 106.167 | 201.838 | 56.702 | 0.164 | 1.87 |
| **SL1344_2188** | *yeiP* | 340.393 | 176.167 | 210.422 | 159.42 | 1.62 | 1.11 |
| **SL1344_2189** | *yeiR* | 63.333 | 51.826 | 36.75 | 42.282 | 1.72 | 1.23 |
| **SL1344_2190** | *yeiU* | 84.27 | 37.79 | 168.556 | 102.659 | 0.5 | 0.368 |
| **SL1344_2191** | *spr* | 2157.753 | 1476.643 | 2104.345 | 2796.515 | 1.03 | 0.528 |
| **SL1344_2192** | *rtn* | 13.42 | 14.941 | 26.243 | 27.316 | 0.511 | 0.547 |
| **SL1344_2193** | *yejA* | 11.996 | 12.542 | 13.733 | 13.094 | 0.874 | 0.958 |
| **SL1344_2194** | *yejB* | 4.973 | 7.268 | 11.915 | 19.012 | 0.417 | 0.382 |
| **SL1344_2195** | *yejE* | 12.577 | 16.641 | 12.366 | 13.268 | 1.02 | 1.25 |
| **SL1344_2196** | *yejF* | 9.382 | 6.588 | 11.992 | 14.34 | 0.782 | 0.459 |
| **SL1344_2197** | *yejG* | 86.818 | 175.278 | 228.349 | 391.487 | 0.38 | 0.448 |
| **SL1344_2198** | *bcr* | 9.394 | 12.051 | 37.978 | 49.555 | 0.247 | 0.243 |
| **SL1344_2199** | *rsuA* | 76.544 | 39.679 | 111.233 | 91.122 | 0.688 | 0.435 |
| **SL1344_2200** | *yejH* | 3.798 | 5.717 | 27.477 | 24.714 | 0.138 | 0.231 |
| **SL1344_2201** | *rplY* | 1980 | 2699.131 | 3826.645 | 3365.014 | 0.517 | 0.802 |

| | | | | | | | |
|---|---|---|---|---|---|---|---|
| **SL1344_2202** | *SL1344-2202* | 27.548 | 14.459 | 184.577 | 117.351 | 0.149 | 0.123 |
| **SL1344_2203** | *yejK* | 248.875 | 87.321 | 241.505 | 154.836 | 1.03 | 0.564 |
| **SL1344_2204** | *yejL* | 151.621 | 392.912 | 397.404 | 598.532 | 0.382 | 0.656 |
| **SL1344_2205** | *yejM* | 32.639 | 16.494 | 95.293 | 79.621 | 0.343 | 0.207 |
| **SL1344_2208** | *oafA* | 365 | 126.007 | 90.29 | 77.808 | 4.04 | 1.62 |
| **SL1344_2210** | *SL1344-2210* | 6.839 | 16.619 | 8.96 | 9.25 | 0.763 | 1.8 |
| **SL1344_2211** | *SL1344-2211* | 3.67 | 2.355 | 5.945 | 5.062 | 0.617 | 0.465 |
| **SL1344_2214** | *SL1344-2214* | 46.262 | 143.6 | 34.765 | 52.406 | 1.33 | 2.74 |
| **SL1344_2217** | *sspH2* | 10.303 | 11.897 | 23.319 | 23.913 | 0.442 | 0.498 |
| **SL1344_2221** | *SL1344-2221* | 2.624 | 13.749 | 7.626 | 5.702 | 0.344 | 2.41 |
| **SL1344_2222** | *narP* | 81.758 | 61.828 | 111.95 | 124.402 | 0.73 | 0.497 |
| **SL1344_2223** | *ccmH2* | 4.933 | 13.031 | 17.234 | 69.665 | 0.286 | 0.187 |
| **SL1344_2223A** | *ccmG2* | 6.047 | 27.428 | 22.738 | 79.643 | 0.266 | 0.344 |
| **SL1344_2223B** | *ccmF2* | 5.055 | 12.217 | 13.357 | 58.933 | 0.378 | 0.207 |
| **SL1344_2223C** | *ccmE2* | 9.989 | 53 | 18.369 | 140.64 | 0.544 | 0.377 |
| **SL1344_2223E** | *ccmC2* | 7.053 | 58.096 | 20.056 | 139.959 | 0.352 | 0.415 |
| **SL1344_2223F** | *ccmB2* | 3.153 | 9.214 | 24.113 | 121.373 | 0.131 | 0.0759 |
| **SL1344_2224** | *ccmA2* | 1.628 | 13.758 | 34.45 | 211.741 | 0.0473 | 0.065 |
| **SL1344_2225** | *napC* | 52.912 | 310.326 | 55.635 | 345.841 | 0.951 | 0.897 |
| **SL1344_2226** | *napB* | 33.149 | 297.026 | 46.196 | 278.916 | 0.718 | 1.06 |
| **SL1344_2227** | *napH* | 8.709 | 40.969 | 41.445 | 317.16 | 0.21 | 0.129 |
| **SL1344_2228** | *napG* | 1.871 | 17.689 | 30.898 | 262.736 | 0.0606 | 0.0673 |
| **SL1344_2229** | *napA* | 111.462 | 538.782 | 115.12 | 467.895 | 0.968 | 1.15 |
| **SL1344_2230** | *napD* | 15.023 | 54.365 | 200.931 | 602.731 | 0.0748 | 0.0902 |

| | | | | | | | |
|---|---|---|---|---|---|---|---|
| **SL1344_2231** | *napF* | 16.603 | 60.97 | 51.432 | 197.018 | 0.323 | 0.309 |
| **SL1344_2232** | *eco* | 350.263 | 727.896 | 183.915 | 225.669 | 1.9 | 3.23 |
| **SL1344_2233** | *yojI* | 27.761 | 4.717 | 33.4 | 8.971 | 0.831 | 0.526 |
| **SL1344_2234** | *alkB* | 16.367 | 13.583 | 45.256 | 38.107 | 0.362 | 0.356 |
| **SL1344_2235** | *ada* | 6.577 | 13.643 | 30.374 | 27.321 | 0.217 | 0.499 |
| **SL1344_2236** | *apbE* | 53.292 | 33.849 | 69.165 | 59.747 | 0.771 | 0.567 |
| **SL1344_2237** | *ompC* | 17381.434 | 17773.829 | 9783.768 | 7521.359 | 1.78 | 2.36 |
| **SL1344_2238** | *yojN* | 30.417 | 27.107 | 70.771 | 85.767 | 0.43 | 0.316 |
| **SL1344_2239** | *rcsB* | 360.349 | 615.848 | 333.642 | 334.864 | 1.08 | 1.84 |
| **SL1344_2240** | *rcsC* | 38.361 | 22.315 | 48.796 | 45.547 | 0.786 | 0.49 |
| **SL1344_2241** | *gyrA* | 440.332 | 226.892 | 474.221 | 351.936 | 0.929 | 0.645 |
| **SL1344_2242** | *SL1344-2242* | 5.609 | 5.315 | 8.104 | 7.658 | 0.692 | 0.694 |
| **SL1344_2245** | *ubiG* | 120.013 | 156.29 | 263.528 | 173.431 | 0.455 | 0.901 |
| **SL1344_2246** | *nrdA* | 441.498 | 351.265 | 223.486 | 297.624 | 1.98 | 1.18 |
| **SL1344_2247** | *nrdB* | 481.981 | 673.356 | 177.846 | 221.693 | 2.71 | 3.04 |
| **SL1344_2250** | *SL1344-2250* | 114.416 | 67.17 | 44.609 | 51.493 | 2.56 | 1.3 |
| **SL1344_2251** | *glpQ* | 3591.457 | 5192.226 | 2995.182 | 3880.623 | 1.2 | 1.34 |
| **SL1344_2252** | *glpT* | 3938.93 | 3682.057 | 5049.024 | 4636.588 | 0.78 | 0.794 |
| **SL1344_2253** | *glpA* | 656.952 | 481.285 | 429.133 | 638.03 | 1.53 | 0.754 |
| **SL1344_2254** | *glpB* | 351.267 | 439.979 | 361.49 | 411.462 | 0.972 | 1.07 |
| **SL1344_2255** | *glpC* | 310.435 | 619.33 | 354.626 | 641.967 | 0.875 | 0.965 |
| **SL1344_2256** | *sseL* | 1.117 | 41.426 | 2.855 | 13.585 | 0.391 | 3.05 |
| **SL1344_2257** | *SL1344-2257* | 27.089 | 13.066 | 26.729 | 140.838 | 1.01 | 0.0928 |
| **SL1344_2258** | *SL1344-2258* | 24.149 | 20.559 | 18.099 | 52.989 | 1.33 | 0.388 |

| | | | | | | | |
|---|---|---|---|---|---|---|---|
| **SL1344_2259** | *yfaV* | 4.13 | 6.117 | 10.225 | 12.764 | 0.404 | 0.479 |
| **SL1344_2260** | *yfaW* | 30.083 | 38.925 | 14.949 | 20.903 | 2.01 | 1.86 |
| **SL1344_2261** | *yfaX* | 3.024 | 11.033 | 5.951 | 16.304 | 0.508 | 0.677 |
| **SL1344_2262** | *SL1344-2262* | 237.668 | 80.125 | 89.769 | 76.455 | 2.65 | 1.05 |
| **SL1344_2263** | *yfaZ* | 439.625 | 124.766 | 413.16 | 211.256 | 1.06 | 0.591 |
| **SL1344_2264** | *yfaO* | 73.368 | 133.488 | 190.147 | 137.921 | 0.386 | 0.968 |
| **SL1344_2265** | *ais* | 21.49 | 23.796 | 83.158 | 104.057 | 0.258 | 0.229 |
| **SL1344_2266** | *yfbE* | 51.936 | 76.95 | 131.853 | 123.734 | 0.394 | 0.622 |
| **SL1344_2267** | *pmrF* | 51.013 | 114.406 | 82.174 | 108.043 | 0.621 | 1.06 |
| **SL1344_2268** | *yfbG* | 129.613 | 108.841 | 88.023 | 90.143 | 1.47 | 1.21 |
| **SL1344_2269** | *SL1344-2269* | 15.522 | 18.97 | 48.187 | 72.375 | 0.322 | 0.262 |
| **SL1344_2270** | *arnT* | 25.23 | 15.075 | 49.443 | 41.502 | 0.51 | 0.363 |
| **SL1344_2271** | *SL1344-2271* | 11.472 | 15.292 | 62.793 | 49.201 | 0.183 | 0.311 |
| **SL1344_2273** | *pmrD* | 209.934 | 136.306 | 179.765 | 298.299 | 1.17 | 0.457 |
| **SL1344_2274** | *menE* | 13.371 | 12.978 | 59.632 | 66.598 | 0.224 | 0.195 |
| **SL1344_2275** | *menC* | 47.392 | 29.384 | 98.407 | 104.474 | 0.482 | 0.281 |
| **SL1344_2276** | *menB* | 151.573 | 177.743 | 159.074 | 152.738 | 0.953 | 1.16 |
| **SL1344_2277** | *yfbB* | 5.303 | 6.184 | 45.806 | 60.367 | 0.116 | 0.102 |
| **SL1344_2278** | *menD* | 16.543 | 13.23 | 45.558 | 47.707 | 0.363 | 0.277 |
| **SL1344_2279** | *menF* | 8.13 | 6.036 | 38.386 | 51.044 | 0.212 | 0.118 |
| **SL1344_2280** | *elaB* | 377.744 | 881.878 | 230.444 | 394.766 | 1.64 | 2.23 |
| **SL1344_2281** | *elaA* | 170.28 | 89.37 | 98.216 | 93.889 | 1.73 | 0.952 |
| **SL1344_2282** | *elaC* | 10.317 | 9.04 | 29.985 | 29.578 | 0.344 | 0.306 |
| **SL1344_2283** | *SL1344-2283* | 170.848 | 13.917 | 184.288 | 20.704 | 0.927 | 0.672 |

| | | | | | | | |
|---|---|---|---|---|---|---|---|
| **SL1344_2284** | *yfbK* | 2.724 | 2.634 | 7.482 | 7.49 | 0.364 | 0.352 |
| **SL1344_2285** | *nuoN* | 420.816 | 333.341 | 531.088 | 429.083 | 0.792 | 0.777 |
| **SL1344_2286** | *nuoM* | 545.633 | 560.131 | 785.474 | 624.651 | 0.695 | 0.897 |
| **SL1344_2287** | *nuoL* | 473.234 | 275.039 | 945.429 | 660.284 | 0.501 | 0.417 |
| **SL1344_2288** | *nuoK* | 467.108 | 424.516 | 1515.528 | 814.739 | 0.308 | 0.521 |
| **SL1344_2289** | *nuoJ* | 500.219 | 289.611 | 876.994 | 739.698 | 0.57 | 0.392 |
| **SL1344_2290** | *nuoI* | 461.781 | 450.844 | 1029.049 | 879.922 | 0.449 | 0.512 |
| **SL1344_2291** | *nuoH* | 463.81 | 382.602 | 778.185 | 506.223 | 0.596 | 0.756 |
| **SL1344_2292** | *nuoG* | 1120.525 | 523.966 | 611.926 | 588.635 | 1.83 | 0.89 |
| **SL1344_2293** | *nuoF* | 584.069 | 470.709 | 684.476 | 642.365 | 0.853 | 0.733 |
| **SL1344_2294** | *nuoE* | 354.457 | 338.07 | 354.555 | 446.029 | 1 | 0.758 |
| **SL1344_2295** | *nuoC* | 713.517 | 384.435 | 639.984 | 516.995 | 1.11 | 0.744 |
| **SL1344_2296** | *nuoB* | 540.338 | 335.9 | 467.403 | 607.425 | 1.16 | 0.553 |
| **SL1344_2297** | *nuoA* | 802.057 | 937.743 | 2323.097 | 1497.399 | 0.345 | 0.626 |
| **SL1344_2299** | *lrhA* | 122.549 | 35.568 | 93.746 | 50.289 | 1.31 | 0.707 |
| **SL1344_2300** | *yfbQ* | 89.206 | 29.672 | 49.264 | 25.353 | 1.81 | 1.17 |
| **SL1344_2301** | *SL1344-2301* | 22.987 | 31.517 | 18.638 | 26.315 | 1.23 | 1.2 |
| **SL1344_2302** | *yfbS* | 44.096 | 26.099 | 43.042 | 58.257 | 1.02 | 0.448 |
| **SL1344_2303** | *yfbT* | 119.436 | 77.552 | 171.068 | 167.79 | 0.698 | 0.462 |
| **SL1344_2304** | *yfbU* | 235.462 | 194.445 | 132.216 | 158.554 | 1.78 | 1.23 |
| **SL1344_2305** | *SL1344-2305* | 43.487 | 18.497 | 57.064 | 81.285 | 0.762 | 0.228 |
| **SL1344_2306** | *ackA* | 438.225 | 216.786 | 323.559 | 142.52 | 1.35 | 1.52 |
| **SL1344_2307** | *pta* | 569.094 | 167.405 | 358.953 | 177.289 | 1.59 | 0.944 |
| **SL1344_2308** | *yfcC* | 2.802 | 0.939 | 5.373 | 3.99 | 0.522 | 0.235 |

| | | | | | | | |
|---|---|---|---|---|---|---|---|
| **SL1344_2309** | *SL1344-2309* | 1268.273 | 4075.001 | 1567.12 | 3391.22 | 0.809 | 1.2 |
| **SL1344_2310** | *SL1344-2310* | 1045.696 | 2578.587 | 1055.672 | 2559.173 | 0.991 | 1.01 |
| **SL1344_2311** | *SL1344-2311* | 1216.504 | 2734.875 | 985.855 | 2969.427 | 1.23 | 0.921 |
| **SL1344_2312** | *SL1344-2312* | 646.151 | 4098.468 | 1380.368 | 4036.181 | 0.468 | 1.02 |
| **SL1344_2313** | *SL1344-2313* | 291.039 | 2735.404 | 653.063 | 2943.119 | 0.446 | 0.929 |
| **SL1344_2314** | *SL1344-2314* | 107.827 | 105.9 | 180.474 | 271.158 | 0.597 | 0.391 |
| **SL1344_2315** | *SL1344-2315* | 102.817 | 106.75 | 334.782 | 299.774 | 0.307 | 0.356 |
| **SL1344_2316** | *yfcE* | 522.776 | 235.363 | 263.749 | 224.136 | 1.98 | 1.05 |
| **SL1344_2317** | *yfcF* | 57.91 | 47.668 | 96.245 | 76.21 | 0.602 | 0.625 |
| **SL1344_2318** | *yfcG* | 1.918 | 11.127 | 19.691 | 22.276 | 0.0974 | 0.5 |
| **SL1344_2319** | *yfcH* | 17.811 | 14.533 | 47.308 | 73.672 | 0.376 | 0.197 |
| **SL1344_2320** | *hisP* | 62.775 | 94.985 | 48.066 | 76.737 | 1.31 | 1.24 |
| **SL1344_2321** | *hisM* | 18.143 | 48.614 | 40.094 | 72.946 | 0.453 | 0.666 |
| **SL1344_2322** | *hisQ* | 48.51 | 52.773 | 59.267 | 113.116 | 0.819 | 0.467 |
| **SL1344_2323** | *hisJ* | 588.39 | 432.973 | 166.895 | 235.18 | 3.53 | 1.84 |
| **SL1344_2324** | *argT* | 130.182 | 215.618 | 51.176 | 162.458 | 2.54 | 1.33 |
| **SL1344_2325** | *ubiX* | 19.524 | 29.595 | 111.55 | 162.168 | 0.175 | 0.182 |
| **SL1344_2327** | *SL1344-2327* | 11.796 | 10.659 | 41.621 | 23.062 | 0.283 | 0.462 |
| **SL1344_2328** | *SL1344-2328* | 5.536 | 3.97 | 17.24 | 12.038 | 0.321 | 0.33 |
| **SL1344_2330** | *SL1344-2330* | 91.32 | 126.182 | 43.02 | 64.714 | 2.12 | 1.95 |
| **SL1344_2331** | *purF* | 58.96 | 34.907 | 52.417 | 47.466 | 1.12 | 0.735 |
| **SL1344_2332** | *cvpA* | 102.41 | 98.485 | 356.805 | 310.715 | 0.287 | 0.317 |
| **SL1344_2333** | *dedD* | 71.472 | 121.733 | 249.884 | 219.58 | 0.286 | 0.554 |
| **SL1344_2334** | *folC* | 123.986 | 70.701 | 211.323 | 171.653 | 0.587 | 0.412 |

| | | | | | | | |
|---|---|---|---|---|---|---|---|
| **SL1344_2335** | *accD* | 434.545 | 595.762 | 746.449 | 787.516 | 0.582 | 0.757 |
| **SL1344_2336** | *dedA* | 143.053 | 72.659 | 179.535 | 101.478 | 0.797 | 0.716 |
| **SL1344_2337** | *truA* | 33.929 | 16.482 | 116.916 | 107.898 | 0.29 | 0.153 |
| **SL1344_2338** | *usg* | 172.329 | 94.72 | 215.192 | 168.68 | 0.801 | 0.562 |
| **SL1344_2339** | *pdxB* | 151.969 | 129.641 | 203.135 | 181.65 | 0.748 | 0.714 |
| **SL1344_2340** | *div* | 25.994 | 9.368 | 28.688 | 24.324 | 0.906 | 0.385 |
| **SL1344_2341** | *SL1344-2341* | 26.158 | 17.077 | 77.399 | 63.964 | 0.338 | 0.267 |
| **SL1344_2347** | *fabB* | 2095.237 | 747.058 | 1039.854 | 525.268 | 2.01 | 1.42 |
| **SL1344_2348** | *mnmC* | 24.844 | 15.215 | 31.407 | 28.613 | 0.791 | 0.532 |
| **SL1344_2349** | *yfcL* | 151.203 | 111.644 | 241.542 | 167.587 | 0.626 | 0.666 |
| **SL1344_2350** | *yfcM* | 81.406 | 40.268 | 74.948 | 63.552 | 1.09 | 0.634 |
| **SL1344_2351** | *yfcA* | 22.801 | 17.299 | 43.807 | 27.253 | 0.52 | 0.635 |
| **SL1344_2352** | *mepA* | 37.884 | 12.12 | 61.257 | 38.891 | 0.618 | 0.312 |
| **SL1344_2353** | *aroC* | 76.419 | 66.894 | 119.935 | 102.816 | 0.637 | 0.651 |
| **SL1344_2354** | *yfcB* | 274.274 | 145.52 | 182.474 | 140.279 | 1.5 | 1.04 |
| **SL1344_2355** | *yfcN* | 26.916 | 14.171 | 39.478 | 39.291 | 0.682 | 0.361 |
| **SL1344_2356** | *sixA* | 667.825 | 215.261 | 709.313 | 625.664 | 0.942 | 0.344 |
| **SL1344_2357** | *fadJ* | 52.415 | 80.878 | 81.695 | 123.911 | 0.642 | 0.653 |
| **SL1344_2358** | *fadI* | 14.087 | 49.446 | 24.277 | 53.927 | 0.58 | 0.917 |
| **SL1344_2359** | *yfcZ* | 528.388 | 962.56 | 680.868 | 575.795 | 0.776 | 1.67 |
| **SL1344_2360** | *fadL* | 396.566 | 338.916 | 207.086 | 305.382 | 1.91 | 1.11 |
| **SL1344_2361** | *vacJ* | 145.324 | 123.806 | 313.379 | 355.742 | 0.464 | 0.348 |
| **SL1344_2362** | *yfdC* | 2.576 | 18.197 | 8.371 | 15.94 | 0.308 | 1.14 |
| **SL1344_2363** | *pgtE* | 1.954 | 14.705 | 9.39 | 16.042 | 0.208 | 0.917 |

| | | | | | | | |
|---|---|---|---|---|---|---|---|
| **SL1344_2364** | *pgtA* | 10.482 | 14.607 | 29.696 | 41.305 | 0.353 | 0.354 |
| **SL1344_2365** | *pgtB* | 9.143 | 7.287 | 27.823 | 31.347 | 0.329 | 0.232 |
| **SL1344_2366** | *pgtC* | 10.758 | 13.274 | 16.15 | 22.316 | 0.666 | 0.595 |
| **SL1344_2367** | *pgtP* | 2.722 | 2.932 | 5.562 | 4.881 | 0.489 | 0.601 |
| **SL1344_2368** | *SL1344-2368* | 22.898 | 123.166 | 50.739 | 59.203 | 0.451 | 2.08 |
| **SL1344_2369A** | *ypdK* | 355.988 | 264.527 | 273.791 | 91.006 | 1.3 | 2.91 |
| **SL1344_2370** | *yfdZ* | 87.717 | 79.001 | 118.547 | 110.063 | 0.74 | 0.718 |
| **SL1344_2371** | *glk* | 320.606 | 348.204 | 346.767 | 264.966 | 0.925 | 1.31 |
| **SL1344_2372** | *SL1344-2372* | 9.53 | 14.034 | 18.965 | 16.96 | 0.503 | 0.827 |
| **SL1344_2373** | *SL1344-2373* | 4.763 | 6.872 | 16.045 | 22.628 | 0.297 | 0.304 |
| **SL1344_2374** | *SL1344-2374* | 31.345 | 39.9 | 39.107 | 45.55 | 0.802 | 0.876 |
| **SL1344_2375** | *ypeC* | 31.679 | 65.525 | 73.657 | 86.511 | 0.43 | 0.757 |
| **SL1344_2376** | *mntH* | 15.728 | 13.801 | 40.171 | 38.877 | 0.392 | 0.355 |
| **SL1344_2377** | *nupC* | 1945.246 | 148.765 | 1409.3 | 192.928 | 1.38 | 0.771 |
| **SL1344_2378** | *yfeA* | 7.676 | 9.815 | 18.101 | 19.884 | 0.424 | 0.494 |
| **SL1344_2379** | *yfeC* | 2042.45 | 1385.722 | 246.647 | 249.648 | 8.28 | 5.55 |
| **SL1344_2380** | *yfeD* | 372.94 | 274.335 | 225.448 | 268.982 | 1.65 | 1.02 |
| **SL1344_2381** | *gltX* | 245.64 | 146.563 | 382.92 | 299.264 | 0.641 | 0.49 |
| **SL1344_2387** | *yfeR* | 6.386 | 9.539 | 16.239 | 16.093 | 0.393 | 0.593 |
| **SL1344_2388** | *yfeH* | 33.893 | 14.98 | 30.783 | 19.314 | 1.1 | 0.776 |
| **SL1344_2390** | *lig* | 141.086 | 57.562 | 131.989 | 119.815 | 1.07 | 0.48 |
| **SL1344_2391** | *zipA* | 464.079 | 238.936 | 636.655 | 588.621 | 0.729 | 0.406 |
| **SL1344_2392** | *cysZ* | 86.15 | 62.13 | 51.834 | 44.391 | 1.66 | 1.4 |
| **SL1344_2393** | *cysK* | 526.172 | 5153.59 | 375.011 | 1912.764 | 1.4 | 2.69 |

| | | | | | | | |
|---|---|---|---|---|---|---|---|
| **SL1344_2394** | *ptsH* | 1487.201 | 1995.29 | 2271.925 | 1696.079 | 0.655 | 1.18 |
| **SL1344_2395** | *ptsI* | 541.245 | 288.461 | 870.906 | 846.889 | 0.621 | 0.341 |
| **SL1344_2396** | *crr* | 1954.343 | 2029.568 | 1981.981 | 1436.08 | 0.986 | 1.41 |
| **SL1344_2397** | *SL1344-2397* | 29.351 | 19.556 | 122.459 | 66.693 | 0.24 | 0.293 |
| **SL1344_2398** | *pdxK* | 35.23 | 20.869 | 43.076 | 28.697 | 0.818 | 0.727 |
| **SL1344_2399** | *ptsJ* | 7.874 | 2.525 | 15.468 | 11.089 | 0.509 | 0.228 |
| **SL1344_2400** | *yfeJ* | 32.721 | 12.093 | 22.102 | 15.078 | 1.48 | 0.802 |
| **SL1344_2403** | *cysM* | 54.132 | 52.284 | 57.85 | 102.68 | 0.936 | 0.509 |
| **SL1344_2404** | *cysA* | 10.62 | 373.869 | 25.723 | 567.145 | 0.413 | 0.659 |
| **SL1344_2405** | *cysW* | 4.257 | 433.519 | 13.829 | 289.014 | 0.308 | 1.5 |
| **SL1344_2406** | *cysU* | 5.039 | 57.092 | 26.129 | 249.152 | 0.193 | 0.229 |
| **SL1344_2407** | *cysP* | 186.197 | 593.798 | 99.596 | 313.112 | 1.87 | 1.9 |
| **SL1344_2408** | *ucpA* | 573.184 | 874.915 | 431.188 | 905.057 | 1.33 | 0.967 |
| **SL1344_2409** | *SL1344-2409* | 321.425 | 143.223 | 183.51 | 202.001 | 1.75 | 0.709 |
| **SL1344_2410** | *yfeY* | 32.166 | 42.749 | 54.136 | 76.524 | 0.594 | 0.559 |
| **SL1344_2411** | *yfeZ* | 14.47 | 10.883 | 172.996 | 152.219 | 0.0836 | 0.0715 |
| **SL1344_2412** | *SL1344-2412* | 54.053 | 59.878 | 377.937 | 297.297 | 0.143 | 0.201 |
| **SL1344_2413** | *amiA* | 22.317 | 15.09 | 56.276 | 36.685 | 0.397 | 0.411 |
| **SL1344_2414** | *hemF* | 30.452 | 9.976 | 31.62 | 22.03 | 0.963 | 0.453 |
| **SL1344_2415** | *SL1344-2415* | 13.633 | 10.925 | 38.014 | 17.878 | 0.359 | 0.611 |
| **SL1344_2416** | *SL1344-2416* | 5.79 | 5.056 | 9.816 | 11.484 | 0.59 | 0.44 |
| **SL1344_2419** | *eutL* | 2.87 | 6.184 | 11.513 | 19.123 | 0.249 | 0.323 |
| **SL1344_2420** | *eutC* | 5.279 | 3.792 | 5.994 | 5.58 | 0.881 | 0.68 |
| **SL1344_2421** | *eutB* | 15.907 | 4.595 | 7.526 | 5.45 | 2.11 | 0.843 |

| | | | | | | | |
|---|---|---|---|---|---|---|---|
| **SL1344_2426** | *eutE* | 1.855 | 2.955 | 4.901 | 5.029 | 0.378 | 0.588 |
| **SL1344_2431** | *eutQ* | 5.062 | 1.282 | 3.74 | 3.504 | 1.35 | 0.366 |
| **SL1344_2435** | *maeB* | 550.663 | 551.748 | 230.582 | 357.295 | 2.39 | 1.54 |
| **SL1344_2436** | *tal* | 68.344 | 88.192 | 26.307 | 42.535 | 2.6 | 2.07 |
| **SL1344_2437** | *tktB* | 59.342 | 48.849 | 22.318 | 33.797 | 2.66 | 1.45 |
| **SL1344_2438** | *SL1344-2438* | 280.34 | 225.854 | 278.666 | 216.772 | 1.01 | 1.04 |
| **SL1344_2440** | *yffH* | 92.697 | 57.393 | 38.829 | 34.379 | 2.39 | 1.67 |
| **SL1344_2441** | *SL1344-2441* | 7.975 | 4.294 | 9.602 | 6.831 | 0.831 | 0.629 |
| **SL1344_2442** | *aegA* | 5.129 | 10.817 | 8.951 | 13.186 | 0.573 | 0.82 |
| **SL1344_2443** | *narQ* | 1.844 | 5.159 | 12.727 | 25.33 | 0.145 | 0.204 |
| **SL1344_2444** | *acrD* | 4.068 | 1.464 | 12.431 | 14.271 | 0.327 | 0.103 |
| **SL1344_2444A** | *ypfM* | 425.213 | 2199.354 | 3371.518 | 3717.092 | 0.126 | 0.592 |
| **SL1344_2445** | *yffB* | 66.324 | 164.242 | 125.496 | 122.384 | 0.528 | 1.34 |
| **SL1344_2446** | *dapE* | 158.534 | 98.595 | 217.084 | 182.199 | 0.73 | 0.541 |
| **SL1344_2447** | *SL1344-2447* | 313.347 | 330.292 | 392.655 | 265.245 | 0.798 | 1.25 |
| **SL1344_2448** | *ypfI* | 3.636 | 4.717 | 21.8 | 25.963 | 0.167 | 0.182 |
| **SL1344_2449** | *SL1344-2449* | 42.82 | 51.488 | 57.496 | 62.518 | 0.745 | 0.824 |
| **SL1344_2450** | *purC* | 110.596 | 153 | 101.702 | 96.004 | 1.09 | 1.59 |
| **SL1344_2451** | *nlpB* | 344.471 | 254.8 | 733.389 | 686.842 | 0.47 | 0.371 |
| **SL1344_2452** | *dapA* | 1530.164 | 708.69 | 1335.484 | 873.381 | 1.15 | 0.811 |
| **SL1344_2453** | *gcvR* | 135.434 | 182.34 | 337.776 | 352.091 | 0.401 | 0.518 |
| **SL1344_2454** | *bcp* | 519.05 | 819.288 | 978.915 | 952.882 | 0.53 | 0.86 |
| **SL1344_2456** | *perM* | 49.55 | 17.069 | 101.417 | 81.954 | 0.489 | 0.208 |
| **SL1344_2457** | *SL1344-2457* | 55.798 | 48.042 | 21.006 | 40.017 | 2.66 | 1.2 |

| | | | | | | | |
|---|---|---|---|---|---|---|---|
| **SL1344_2458** | *yfgD* | 236.613 | 415.119 | 145.557 | 209.617 | 1.63 | 1.98 |
| **SL1344_2459** | *yfgE* | 71.262 | 39.726 | 194.316 | 156.596 | 0.367 | 0.254 |
| **SL1344_2460** | *uraA* | 59.791 | 71.765 | 53.957 | 48.47 | 1.11 | 1.48 |
| **SL1344_2461** | *upp* | 542.568 | 907.601 | 427.815 | 342.283 | 1.27 | 2.65 |
| **SL1344_2462** | *purM* | 47.445 | 31.007 | 81.146 | 86.497 | 0.585 | 0.358 |
| **SL1344_2463** | *purN* | 34.553 | 43.219 | 69.889 | 64.855 | 0.494 | 0.666 |
| **SL1344_2464** | *ppk* | 72.139 | 54.628 | 71.58 | 96.399 | 1.01 | 0.567 |
| **SL1344_2465** | *ppx* | 114.512 | 71.462 | 83.863 | 83.168 | 1.37 | 0.859 |
| **SL1344_2466** | *SL1344-2466* | 0.561 | 2.212 | 4.177 | 5.242 | 0.134 | 0.422 |
| **SL1344_2468** | *SL1344-2468* | 8.941 | 75.107 | 109.765 | 153.888 | 0.0815 | 0.488 |
| **SL1344_2469** | *SL1344-2469* | 9.866 | 12.986 | 9.558 | 10.624 | 1.03 | 1.22 |
| **SL1344_2471** | *guaA* | 461.326 | 190.873 | 335.886 | 185.769 | 1.37 | 1.03 |
| **SL1344_2472** | *guaB* | 138.483 | 44.003 | 73.101 | 38.6 | 1.89 | 1.14 |
| **SL1344_2473** | *xseA* | 21.091 | 6.399 | 66.48 | 45.008 | 0.317 | 0.142 |
| **SL1344_2474** | *shdA* | 0.938 | 0.6 | 6.583 | 8.569 | 0.143 | 0.07 |
| **SL1344_2477** | *ratA* | 0.497 | 0.693 | 6.121 | 10.172 | 0.0812 | 0.0681 |
| **SL1344_2478** | *sinI* | 6.104 | 2.905 | 7.616 | 5.994 | 0.801 | 0.485 |
| **SL1344_2481** | *engA* | 109.146 | 59.848 | 160.741 | 102.411 | 0.679 | 0.584 |
| **SL1344_2482** | *yfgL* | 775.803 | 283.682 | 570.489 | 389.732 | 1.36 | 0.728 |
| **SL1344_2483** | *yfgM* | 246.881 | 173.723 | 370.307 | 209.316 | 0.667 | 0.83 |
| **SL1344_2484** | *hisS* | 570.309 | 267.924 | 649.706 | 432.564 | 0.878 | 0.619 |
| **SL1344_2485** | *gcpE* | 306.446 | 229.534 | 301.532 | 272.039 | 1.02 | 0.844 |
| **SL1344_2486** | *yfgA* | 182.648 | 156.55 | 385.308 | 294.019 | 0.474 | 0.532 |
| **SL1344_2487** | *yfgB* | 244.843 | 108.123 | 173.527 | 116.522 | 1.41 | 0.928 |

| | | | | | | | |
|---|---|---|---|---|---|---|---|
| **SL1344_2488** | *ndk* | 8894.228 | 1953.092 | 1157.391 | 1048.757 | 7.68 | 1.86 |
| **SL1344_2492** | *SL1344-2492* | 0.995 | 2.259 | 5.062 | 8.537 | 0.197 | 0.265 |
| **SL1344_2493** | *pbpC* | 3.425 | 0.705 | 22.378 | 21.746 | 0.153 | 0.0324 |
| **SL1344_2494** | *SL1344-2494* | 36.692 | 30.089 | 61.124 | 78.465 | 0.6 | 0.383 |
| **SL1344_2495** | *SL1344-2495* | 213.746 | 324.695 | 237.755 | 286.044 | 0.899 | 1.14 |
| **SL1344_2496** | *SL1344-2496* | 7.963 | 4.27 | 12.537 | 9.252 | 0.635 | 0.462 |
| **SL1344_2497** | *sseBb* | 40.987 | 49.725 | 85.799 | 84.237 | 0.478 | 0.59 |
| **SL1344_2498** | *pepB* | 180.949 | 134.294 | 157.492 | 129.895 | 1.15 | 1.03 |
| **SL1344_2499** | *yfhJ* | 539.817 | 335.707 | 930.817 | 504.387 | 0.58 | 0.666 |
| **SL1344_2500** | *fdx* | 338.077 | 162.158 | 341.564 | 219.48 | 0.99 | 0.739 |
| **SL1344_2501** | *hscA* | 143.301 | 30.391 | 188.368 | 87.833 | 0.761 | 0.346 |
| **SL1344_2502** | *hscB* | 69.06 | 29.66 | 181.015 | 88.093 | 0.382 | 0.337 |
| **SL1344_2503** | *iscA* | 231.297 | 99.932 | 606.434 | 327.086 | 0.381 | 0.306 |
| **SL1344_2504** | *nifU* | 127.107 | 233.241 | 330.819 | 214.111 | 0.384 | 1.09 |
| **SL1344_2505** | *iscS* | 946.525 | 309.651 | 505.325 | 168.774 | 1.87 | 1.83 |
| **SL1344_2506** | *yfhP* | 550.807 | 75.579 | 772.59 | 281.402 | 0.713 | 0.269 |
| **SL1344_2507** | *SL1344-2507* | 444.362 | 166.708 | 542.621 | 278.816 | 0.819 | 0.598 |
| **SL1344_2508** | *suhB* | 167.644 | 64.129 | 452.659 | 512.687 | 0.37 | 0.125 |
| **SL1344_2511** | *asrB* | 2.313 | 7.641 | 4.114 | 6.82 | 0.562 | 1.12 |
| **SL1344_2513** | *SL1344-2513* | 3.119 | 5.789 | 25.71 | 25.33 | 0.121 | 0.229 |
| **SL1344_2514** | *SL1344-2514* | 8.152 | 11.284 | 34.552 | 32.503 | 0.236 | 0.347 |
| **SL1344_2515** | *csiE* | 4.91 | 19.853 | 19.632 | 44.933 | 0.25 | 0.442 |
| **SL1344_2516** | *hcaT* | 25.859 | 7.995 | 60.365 | 31.899 | 0.428 | 0.251 |
| **SL1344_2517** | *glyA* | 699.287 | 1062.249 | 680.708 | 846.857 | 1.03 | 1.25 |

| | | | | | | | |
|---|---|---|---|---|---|---|---|
| **SL1344_2518** | *hmpA* | 45.874 | 30.784 | 22.029 | 19.205 | 2.08 | 1.6 |
| **SL1344_2519** | *cadC* | 27.126 | 16.51 | 55.259 | 50.549 | 0.491 | 0.327 |
| **SL1344_2520** | *cadB* | 8.621 | 66.949 | 8.664 | 30.931 | 0.995 | 2.16 |
| **SL1344_2521** | *cadA* | 5.715 | 61.258 | 12.67 | 39.984 | 0.451 | 1.53 |
| **SL1344_2522** | *yjdL* | 1.54 | 5.028 | 7.85 | 13.736 | 0.196 | 0.366 |
| **SL1344_2523** | *glnB* | 296.146 | 530.325 | 349.748 | 311.971 | 0.847 | 1.7 |
| **SL1344_2524** | *yfhA* | 25.483 | 27.656 | 65.469 | 75.699 | 0.389 | 0.365 |
| **SL1344_2525** | *yfhG* | 15.398 | 24.808 | 56.504 | 96.146 | 0.273 | 0.258 |
| **SL1344_2526** | *yfhK* | 10.173 | 14.707 | 56.681 | 53.418 | 0.179 | 0.275 |
| **SL1344_2527** | *purG* | 31.546 | 20.609 | 18.492 | 20.466 | 1.71 | 1.01 |
| **SL1344_2529** | *yfhD* | 4.291 | 3.038 | 21.621 | 16.197 | 0.198 | 0.188 |
| **SL1344_2530** | *yfhC* | 15.978 | 12.199 | 87.137 | 95.226 | 0.183 | 0.128 |
| **SL1344_2531** | *yfhB* | 74.551 | 75.08 | 162.753 | 143.97 | 0.458 | 0.521 |
| **SL1344_2532** | *SL1344-2532* | 42.505 | 50.891 | 65.894 | 79.779 | 0.645 | 0.638 |
| **SL1344_2533** | *murQ* | 135.405 | 117.554 | 221.786 | 171.929 | 0.611 | 0.684 |
| **SL1344_2534** | *yfhH* | 7.321 | 24.997 | 28.623 | 34.374 | 0.256 | 0.727 |
| **SL1344_2535** | *SL1344-2535* | 3.823 | 2.981 | 34.478 | 25.49 | 0.111 | 0.117 |
| **SL1344_2537** | *SL1344-2537* | 2.682 | 12.107 | 4.408 | 6.442 | 0.608 | 1.88 |
| **SL1344_2539** | *acpS* | 16.935 | 13.39 | 276.01 | 266.982 | 0.0614 | 0.0502 |
| **SL1344_2540** | *pdxJ* | 140.384 | 240.49 | 163.54 | 171.696 | 0.858 | 1.4 |
| **SL1344_2541** | *recO* | 23.873 | 32.938 | 80.533 | 49.485 | 0.296 | 0.666 |
| **SL1344_2542** | *era* | 494.854 | 177.711 | 247.49 | 168.558 | 2 | 1.05 |
| **SL1344_2543** | *rnc* | 195.663 | 162.612 | 221.577 | 195.774 | 0.883 | 0.831 |
| **SL1344_2544** | *lepB* | 196.829 | 124.322 | 296.292 | 196.158 | 0.664 | 0.634 |

| | | | | | | | |
|---|---|---|---|---|---|---|---|
| **SL1344_2545** | *lepA* | 449.91 | 163.327 | 343.842 | 306.01 | 1.31 | 0.534 |
| **SL1344_2546** | *gogB* | 6.656 | 13.113 | 4.03 | 7.526 | 1.65 | 1.74 |
| **SL1344_2547** | *sarA* | 20.791 | 48.847 | 5.132 | 18.986 | 4.05 | 2.57 |
| **SL1344_2549** | *SL1344-2549* | 13.5 | 117.247 | 3.144 | 28.739 | 4.29 | 4.08 |
| **SL1344_2556** | *SL1344-2556* | 9.781 | 6.977 | 22.159 | 16.946 | 0.441 | 0.412 |
| **SL1344_2561** | *SL1344-2561* | 9.509 | 15.207 | 16.122 | 17.382 | 0.59 | 0.875 |
| **SL1344_2563** | *gipA* | 84.273 | 47.087 | 218.546 | 162.874 | 0.386 | 0.289 |
| **SL1344_2566** | *SL1344-2566* | 6.413 | 22.863 | 13.54 | 24.783 | 0.474 | 0.923 |
| **SL1344_2567** | *SL1344-2567* | 8.859 | 6.61 | 15.744 | 13.441 | 0.563 | 0.492 |
| **SL1344_2568** | *SL1344-2568* | 11.056 | 9.969 | 12.153 | 8.893 | 0.91 | 1.12 |
| **SL1344_2569** | *SL1344-2569* | 5.964 | 2.676 | 8.954 | 5.54 | 0.666 | 0.483 |
| **SL1344_2570** | *SL1344-2570* | 2.209 | 2.624 | 23.033 | 31.168 | 0.0959 | 0.0842 |
| **SL1344_2574** | *SL1344-2574* | 10.307 | 46.024 | 12.661 | 14.555 | 0.814 | 3.16 |
| **SL1344_2577** | *SL1344-2577* | 31.105 | 96.537 | 42.05 | 45.436 | 0.74 | 2.12 |
| **SL1344_2578** | *SL1344-2578* | 38.623 | 14.593 | 58.084 | 28.68 | 0.665 | 0.509 |
| **SL1344_2583** | *SL1344-2583* | 14.739 | 19.942 | 129.837 | 82.635 | 0.114 | 0.241 |
| **SL1344_2584** | *SL1344-2584* | 55.4 | 49.999 | 74.747 | 40.711 | 0.741 | 1.23 |
| **SL1344_2585** | *SL1344-2585* | 19.423 | 75.107 | 13.441 | 57.677 | 1.45 | 1.3 |
| **SL1344_2587** | *SL1344-2587* | 17.378 | 10.713 | 8.111 | 8.392 | 2.14 | 1.28 |
| **SL1344_2589** | *SL1344-2589* | 4.711 | 7.966 | 7.534 | 11.545 | 0.625 | 0.69 |
| **SL1344_2590** | *gpP* | 5.04 | 3.337 | 4.758 | 3.105 | 1.06 | 1.07 |
| **SL1344_2591** | *gpO* | 1.111 | 5.481 | 9.099 | 10.835 | 0.122 | 0.506 |
| **SL1344_2593** | *SL1344-2593* | 81.392 | 75.107 | 305.029 | 253.248 | 0.267 | 0.297 |
| **SL1344_2594** | *SL1344-2594* | 334.399 | 595.405 | 348.187 | 366.089 | 0.96 | 1.63 |

| | | | | | | | |
|---|---|---|---|---|---|---|---|
| **SL1344_2595** | *SL1344-2595* | 15.442 | 30.889 | 81.726 | 48.474 | 0.189 | 0.637 |
| **SL1344_2597** | *recEb* | 5.529 | 9.223 | 6.136 | 7.266 | 0.901 | 1.27 |
| **SL1344_2597A** | *SL1344-2597A* | 9.599 | 24.512 | 4.391 | 10.848 | 2.19 | 2.26 |
| **SL1344_2599** | *SL1344-2599* | 11.423 | 20.764 | 12.576 | 12.811 | 0.908 | 1.62 |
| **SL1344_2600** | *SL1344-2600* | 32.966 | 15.54 | 39.98 | 24.14 | 0.825 | 0.644 |
| **SL1344_2601** | *rseC* | 8.756 | 25.933 | 49.88 | 70.874 | 0.176 | 0.366 |
| **SL1344_2602** | *rseB* | 33.958 | 36.818 | 99.174 | 98.282 | 0.342 | 0.375 |
| **SL1344_2603** | *rseA* | 42.191 | 141.371 | 253.26 | 366.359 | 0.167 | 0.386 |
| **SL1344_2604** | *rpoE* | 52.103 | 94.356 | 215.3 | 540.12 | 0.242 | 0.175 |
| **SL1344_2605** | *nadB* | 7.185 | 8.424 | 8.303 | 9.028 | 0.865 | 0.933 |
| **SL1344_2606** | *yfiC* | 29.036 | 26.268 | 111.702 | 70.866 | 0.26 | 0.371 |
| **SL1344_2607** | *srmB* | 167.074 | 159.837 | 144.173 | 110.335 | 1.16 | 1.45 |
| **SL1344_2608** | *yfiE* | 19.995 | 16.325 | 25.328 | 26.838 | 0.789 | 0.608 |
| **SL1344_2610** | *yfiD* | 97.116 | 679.15 | 139.82 | 289.957 | 0.695 | 2.34 |
| **SL1344_2611** | *ung* | 170.377 | 183.855 | 126.225 | 133.404 | 1.35 | 1.38 |
| **SL1344_2612** | *yfiF* | 258.106 | 83.618 | 182.662 | 81.388 | 1.41 | 1.03 |
| **SL1344_2613** | *trxC* | 147.281 | 242.123 | 349.031 | 574.535 | 0.422 | 0.421 |
| **SL1344_2614** | *yfiP* | 12.256 | 17.18 | 24.316 | 31.741 | 0.504 | 0.541 |
| **SL1344_2615** | *yfiQ* | 42.066 | 50.409 | 78.661 | 106.12 | 0.535 | 0.475 |
| **SL1344_2616** | *pssA* | 146.72 | 143.015 | 212.513 | 241.307 | 0.69 | 0.593 |
| **SL1344_2618** | *kgtP* | 18.186 | 33.906 | 17.012 | 34.411 | 1.07 | 0.985 |
| **SL1344_2620** | *clpB* | 898.54 | 249.147 | 1951.553 | 465.475 | 0.46 | 0.535 |
| **SL1344_2621** | *yfiH* | 28.061 | 28.993 | 204.964 | 186.857 | 0.137 | 0.155 |
| **SL1344_2622** | *rluD* | 163.644 | 73.984 | 226.158 | 184.213 | 0.724 | 0.402 |

| | | | | | | | |
|---|---|---|---|---|---|---|---|
| **SL1344_2623** | *yfiO* | 275.358 | 276.417 | 379.593 | 338.209 | 0.725 | 0.817 |
| **SL1344_2624** | *raiA* | 789.782 | 12295.394 | 514.684 | 2100.634 | 1.53 | 5.85 |
| **SL1344_2625** | *pheA* | 99.167 | 92.16 | 72.239 | 58.896 | 1.37 | 1.56 |
| **SL1344_2626** | *SL1344-2626* | 6.903 | 9.429 | 18.138 | 22.077 | 0.381 | 0.427 |
| **SL1344_2627** | *tyrA* | 87.461 | 47.258 | 103.565 | 69.661 | 0.845 | 0.678 |
| **SL1344_2628** | *aroF* | 46.261 | 23.182 | 41.096 | 36.009 | 1.13 | 0.644 |
| **SL1344_2630** | *yfiN* | 2.618 | 2.451 | 10.626 | 11.208 | 0.246 | 0.219 |
| **SL1344_2631** | *int* | 29.597 | 51.178 | 151.423 | 151.267 | 0.195 | 0.338 |
| **SL1344_2632** | *SL1344-2632* | 423.964 | 446.051 | 274.681 | 180.878 | 1.54 | 2.47 |
| **SL1344_2633** | *SL1344-2633* | 157.012 | 116.022 | 255.214 | 209.693 | 0.615 | 0.553 |
| **SL1344_2644** | *SL1344-2644* | 216.531 | 690.169 | 167.158 | 284.504 | 1.3 | 2.43 |
| **SL1344_2645** | *gpB* | 76.493 | 95.665 | 129.947 | 291.36 | 0.589 | 0.328 |
| **SL1344_2646** | *rplS* | 6604.765 | 3652.82 | 6073.329 | 2175.261 | 1.09 | 1.68 |
| **SL1344_2647** | *trmD* | 47.787 | 15.854 | 4624.804 | 2020.443 | 0.0103 | 0.00785 |
| **SL1344_2648** | *rimM* | 314.949 | 158.592 | 3971.453 | 2481.391 | 0.0793 | 0.0639 |
| **SL1344_2649** | *rpsP* | 4671.6 | 5432.686 | 3320.763 | 1530.646 | 1.41 | 3.55 |
| **SL1344_2650** | *ffh* | 221.957 | 147.879 | 349.944 | 363.517 | 0.634 | 0.407 |
| **SL1344_2651** | *corE* | 63.38 | 38.219 | 242.928 | 130.877 | 0.261 | 0.292 |
| **SL1344_2652** | *corB* | 28.072 | 23.879 | 58.813 | 74.697 | 0.477 | 0.32 |
| **SL1344_2653** | *SL1344-2653* | 13.675 | 45.46 | 32.885 | 41.096 | 0.416 | 1.11 |
| **SL1344_2654** | *grpE* | 3234.657 | 2113.033 | 4282.942 | 1951.244 | 0.755 | 1.08 |
| **SL1344_2655** | *ppnK* | 46.332 | 75.605 | 59.043 | 79.221 | 0.785 | 0.954 |
| **SL1344_2656** | *recN* | 40.033 | 30.982 | 86.477 | 64.506 | 0.463 | 0.48 |
| **SL1344_2657** | *smpA* | 558.592 | 1108.807 | 631.196 | 548.196 | 0.885 | 2.02 |

| | | | | | | | |
|---|---|---|---|---|---|---|---|
| **SL1344_2658** | *rnfH* | 111.676 | 86.02 | 106.417 | 172.908 | 1.05 | 0.497 |
| **SL1344_2659** | *yfjG* | 118.129 | 175.871 | 294.28 | 297.516 | 0.401 | 0.591 |
| **SL1344_2660** | *smpB* | 542.186 | 150.83 | 361.238 | 208.048 | 1.5 | 0.725 |
| **SL1344_2661** | *bapA* | 0.263 | 0.089 | 2.186 | 2.299 | 0.12 | 0.0387 |
| **SL1344_2670** | *gpE* | 11.722 | 5.388 | 17.034 | 9.416 | 0.688 | 0.572 |
| **SL1344_2674** | *sopE* | 510.971 | 334.273 | 126.215 | 108.353 | 4.05 | 3.09 |
| **SL1344_2675** | *SL1344-2675* | 4.062 | 19.506 | 6.149 | 6.993 | 0.661 | 2.79 |
| **SL1344_2676** | *SL1344-2676* | 4.275 | 2.687 | 6.903 | 4.94 | 0.619 | 0.544 |
| **SL1344_2691** | *SL1344-2691* | 4.154 | 0.251 | 5.957 | 2.3 | 0.697 | 0.109 |
| **SL1344_2695** | *SL1344-2695* | 157.488 | 140.076 | 56.365 | 102.673 | 2.79 | 1.36 |
| **SL1344_2696** | *SL1344-2696* | 88.792 | 523.951 | 122.429 | 281.172 | 0.725 | 1.86 |
| **SL1344_2697** | *SL1344-2697* | 220.021 | 292.06 | 81.734 | 110.788 | 2.69 | 2.64 |
| **SL1344_2698** | *SL1344-2698* | 193.727 | 242.39 | 98.836 | 78.329 | 1.96 | 3.09 |
| **SL1344_2699** | *SL1344-2699* | 32.38 | 40.406 | 80.874 | 70.367 | 0.4 | 0.574 |
| **SL1344_2700** | *SL1344-2700* | 33.825 | 58.304 | 71.684 | 79.956 | 0.472 | 0.729 |
| **SL1344_2702** | *SL1344-2702* | 11.659 | 4.519 | 7.102 | 6.566 | 1.64 | 0.688 |
| **SL1344_2703** | *SL1344-2703* | 34.011 | 19.989 | 30.811 | 6.045 | 1.1 | 3.31 |
| **SL1344_2706** | *cIIc* | 13.58 | 4.001 | 10.682 | 3.224 | 1.27 | 1.24 |
| **SL1344_2707** | *apl* | 66.326 | 16.391 | 22.743 | 9.419 | 2.92 | 1.74 |
| **SL1344_2708** | *cI* | 78.001 | 107.346 | 109.019 | 142.764 | 0.715 | 0.752 |
| **SL1344_2709** | *SL1344-2709* | 26.251 | 28.176 | 36.82 | 51.091 | 0.713 | 0.551 |
| **SL1344_2710** | *SL1344-2710* | 14.994 | 27.22 | 17.211 | 19.531 | 0.871 | 1.39 |
| **SL1344_2712** | *SL1344-2712* | 12.511 | 5.967 | 15.69 | 11.776 | 0.797 | 0.507 |
| **SL1344_2713** | *SL1344-2713* | 42.496 | 206.547 | 22.333 | 58.852 | 1.9 | 3.51 |

| | | | | | | | |
|---|---|---|---|---|---|---|---|
| **SL1344_2714** | *SL1344-2714* | 43.257 | 184.297 | 109.67 | 158.092 | 0.394 | 1.17 |
| **SL1344_2715** | *SL1344-2715* | 15.311 | 84.376 | 37.735 | 83.855 | 0.406 | 1.01 |
| **SL1344_2719** | *SL1344-2719* | 2.359 | 6.777 | 3416.528 | 2499.618 | 0.000691 | 0.00271 |
| **SL1344_2722** | *SL1344-2722* | 3.276 | 4.193 | 8.626 | 7.634 | 0.38 | 0.549 |
| **SL1344_2726** | *SL1344-2726* | 73.77 | 75.142 | 43.226 | 39.792 | 1.71 | 1.89 |
| **SL1344_2727** | *SL1344-2727* | 20.732 | 48.962 | 18.267 | 35.421 | 1.13 | 1.38 |
| **SL1344_2730** | *SL1344-2730* | 83.167 | 129.579 | 43.602 | 38.453 | 1.91 | 3.37 |
| **SL1344_2731** | *SL1344-2731* | 31.585 | 63.952 | 20.218 | 21.699 | 1.56 | 2.95 |
| **SL1344_2732** | *SL1344-2732* | 162.53 | 197.285 | 350.535 | 268.875 | 0.464 | 0.734 |
| **SL1344_2738** | *SL1344-2738* | 61.113 | 67.343 | 12.394 | 15.778 | 4.93 | 4.27 |
| **SL1344_2745** | *SL1344-2745* | 2.115 | 3.12 | 2.179 | 2.179 | 0.971 | 1.43 |
| **SL1344_2746** | *SL1344-2746* | 4.841 | 7.364 | 7.258 | 5.969 | 0.667 | 1.23 |
| **SL1344_2751** | *SL1344-2751* | 98.935 | 73.557 | 52.501 | 66.444 | 1.88 | 1.11 |
| **SL1344_2752** | *SL1344-2752* | 111.195 | 149.969 | 48.62 | 77.84 | 2.29 | 1.93 |
| **SL1344_2756** | *fljB* | 1002.841 | 20.348 | 538.829 | 24.131 | 1.86 | 0.843 |
| **SL1344_2757** | *hin* | 10.434 | 16.857 | 12.385 | 17.681 | 0.842 | 0.953 |
| **SL1344_2759** | *iroC* | 1.264 | 0.391 | 5.061 | 3.282 | 0.25 | 0.119 |
| **SL1344_2760** | *iroD* | 2.805 | 3.77 | 5.182 | 2.991 | 0.541 | 1.26 |
| **SL1344_2761** | *iroE* | 5.945 | 2.253 | 4.978 | 3.41 | 1.19 | 0.661 |
| **SL1344_2763** | *pipB2* | 10.737 | 72.156 | 19.129 | 64.385 | 0.561 | 1.12 |
| **SL1344_2764** | *virK* | 50.538 | 126.315 | 20.811 | 85.276 | 2.43 | 1.48 |
| **SL1344_2765** | *mig-14* | 164.385 | 280.654 | 56.5 | 105.179 | 2.91 | 2.67 |
| **SL1344_2767** | *nixA* | 15.645 | 19.655 | 43.053 | 52.34 | 0.363 | 0.376 |
| **SL1344_2768** | *tctE* | 27.423 | 77.917 | 120.789 | 271.943 | 0.227 | 0.287 |

| | | | | | | | |
|---|---|---|---|---|---|---|---|
| **SL1344_2769** | *tctD* | 131.543 | 476.048 | 431.907 | 732.565 | 0.305 | 0.65 |
| **SL1344_2770** | *SL1344-2770* | 454.671 | 444.99 | 283.216 | 382.733 | 1.61 | 1.16 |
| **SL1344_2771** | *SL1344-2771* | 227.524 | 100.39 | 341.609 | 317.827 | 0.666 | 0.316 |
| **SL1344_2772** | *SL1344-2772* | 66.266 | 77.809 | 160.59 | 169.954 | 0.413 | 0.458 |
| **SL1344_2773** | *csiD* | 20.871 | 281.182 | 9.369 | 142.109 | 2.23 | 1.98 |
| **SL1344_2774** | *ygaF* | 5.038 | 39.88 | 11.524 | 78.043 | 0.437 | 0.511 |
| **SL1344_2775** | *gabD* | 13.522 | 34.879 | 8.806 | 37.745 | 1.54 | 0.924 |
| **SL1344_2776** | *gabT* | 12.125 | 21.72 | 9.77 | 24.97 | 1.24 | 0.87 |
| **SL1344_2777** | *gabP* | 2.197 | 7.574 | 7.47 | 15.36 | 0.294 | 0.493 |
| **SL1344_2779** | *ygaU* | 19.337 | 63.184 | 18.797 | 26.758 | 1.03 | 2.36 |
| **SL1344_2780** | *yqaE* | 23.082 | 77.433 | 29.305 | 60.067 | 0.788 | 1.29 |
| **SL1344_2782** | *ygaP* | 25.001 | 33.238 | 26.881 | 33.658 | 0.93 | 0.988 |
| **SL1344_2783** | *stpA* | 1072.861 | 1675.658 | 310.272 | 440.512 | 3.46 | 3.8 |
| **SL1344_2784** | *SL1344-2784* | 15.522 | 33.406 | 90.481 | 181.395 | 0.172 | 0.184 |
| **SL1344_2785** | *ygaC* | 10.793 | 11.821 | 34.923 | 66.612 | 0.309 | 0.177 |
| **SL1344_2786** | *ygaM* | 25.319 | 141.661 | 37.851 | 189.579 | 0.669 | 0.747 |
| **SL1344_2787** | *SL1344-2787* | 3.813 | 2.955 | 24.807 | 38.396 | 0.154 | 0.077 |
| **SL1344_2788** | *SL1344-2788* | 2.74 | 16.375 | 4.978 | 5.877 | 0.551 | 2.79 |
| **SL1344_2792** | *nrdE* | 3.146 | 1.459 | 5.848 | 8.691 | 0.538 | 0.168 |
| **SL1344_2794** | *proV* | 84.83 | 79.273 | 84.793 | 92.153 | 1 | 0.86 |
| **SL1344_2795** | *proW* | 168.357 | 97.465 | 143.367 | 88.633 | 1.17 | 1.1 |
| **SL1344_2796** | *proX* | 297.933 | 151.614 | 123.503 | 67.487 | 2.41 | 2.25 |
| **SL1344_2797** | *SL1344-2797* | 8.292 | 6.486 | 22.503 | 16.14 | 0.368 | 0.402 |
| **SL1344_2798** | *emrR* | 64.88 | 146.034 | 229.09 | 381.078 | 0.283 | 0.383 |

| | | | | | | | |
|---|---|---|---|---|---|---|---|
| **SL1344_2799** | *emrA* | 27.604 | 50.277 | 74.617 | 72.062 | 0.37 | 0.698 |
| **SL1344_2800** | *emrB* | 18.693 | 6.762 | 53.332 | 33.024 | 0.351 | 0.205 |
| **SL1344_2802** | *luxS* | 430.536 | 372.667 | 217.968 | 249.332 | 1.98 | 1.49 |
| **SL1344_2803** | *gshA* | 239.477 | 147.401 | 199.397 | 194.331 | 1.2 | 0.759 |
| **SL1344_2805** | *yqaB* | 46.144 | 78.458 | 407.092 | 284.815 | 0.113 | 0.275 |
| **SL1344_2806** | *csrA* | 628.542 | 880.619 | 570.771 | 730.435 | 1.1 | 1.21 |
| **SL1344_2807** | *alaS* | 519.205 | 386.255 | 534.889 | 391.135 | 0.971 | 0.988 |
| **SL1344_2808** | *oraA* | 11.461 | 4.888 | 41.923 | 33.348 | 0.273 | 0.147 |
| **SL1344_2809** | *recA* | 1330.257 | 647.291 | 1131.548 | 790.047 | 1.18 | 0.819 |
| **SL1344_2810** | *ygaD* | 87.365 | 152.98 | 526.255 | 357.536 | 0.166 | 0.428 |
| **SL1344_2811** | *mltB* | 45.93 | 52.402 | 67.369 | 73.141 | 0.682 | 0.716 |
| **SL1344_2812** | *srlA* | 62.868 | 289.211 | 65.202 | 180.484 | 0.964 | 1.6 |
| **SL1344_2813** | *srlE* | 52.069 | 102.592 | 70.799 | 125.496 | 0.735 | 0.817 |
| **SL1344_2814** | *slrB* | 15.818 | 70.083 | 44.432 | 75.734 | 0.356 | 0.925 |
| **SL1344_2815** | *gutD* | 49.48 | 169.007 | 34.739 | 91.26 | 1.42 | 1.85 |
| **SL1344_2816** | *gutM* | 5.426 | 32.499 | 34.249 | 49.029 | 0.158 | 0.663 |
| **SL1344_2817** | *srlR* | 145.156 | 137.009 | 186.618 | 160.661 | 0.778 | 0.853 |
| **SL1344_2818** | *gutQ* | 131.318 | 99.074 | 97.433 | 84.501 | 1.35 | 1.17 |
| **SL1344_2819** | *norR* | 11.364 | 15.966 | 31.577 | 31.142 | 0.36 | 0.513 |
| **SL1344_2822** | *hypF* | 10.486 | 10.138 | 23.479 | 23.499 | 0.447 | 0.431 |
| **SL1344_2825** | *hycI* | 6.661 | 6.21 | 18.872 | 14.22 | 0.353 | 0.437 |
| **SL1344_2827** | *hycG* | 8.71 | 3.808 | 9.707 | 7.682 | 0.897 | 0.496 |
| **SL1344_2828** | *hycF* | 8.721 | 2.004 | 15.446 | 9.529 | 0.565 | 0.21 |
| **SL1344_2829** | *hycE* | 4.327 | 0.716 | 9.181 | 6.122 | 0.471 | 0.117 |

| | | | | | | | |
|---|---|---|---|---|---|---|---|
| **SL1344_2835** | *hypB* | 63.805 | 145.237 | 63.472 | 81.51 | 1.01 | 1.78 |
| **SL1344_2836** | *hypC* | 8.89 | 48.836 | 40.962 | 76.611 | 0.217 | 0.637 |
| **SL1344_2837** | *hypD* | 20.576 | 28.009 | 19.614 | 53.077 | 1.05 | 0.528 |
| **SL1344_2838** | *hypE* | 22.893 | 42.32 | 34.459 | 77.033 | 0.664 | 0.549 |
| **SL1344_2839** | *fhlA* | 3.872 | 7.623 | 8.517 | 14.735 | 0.455 | 0.517 |
| **SL1344_2841** | *sitA* | 32.37 | 5.409 | 28.97 | 7.217 | 1.12 | 0.75 |
| **SL1344_2842** | *sitB* | 15.843 | 4.386 | 23.895 | 17.06 | 0.663 | 0.257 |
| **SL1344_2843** | *sitC* | 17.463 | 4.74 | 19.399 | 8.032 | 0.9 | 0.59 |
| **SL1344_2844** | *sitD* | 15.548 | 4.487 | 12.918 | 13.585 | 1.2 | 0.33 |
| **SL1344_2845** | *avrA* | 52.726 | 88.292 | 61.477 | 115.503 | 0.858 | 0.764 |
| **SL1344_2846** | *sprB* | 329.954 | 611.202 | 136.256 | 171.203 | 2.42 | 3.57 |
| **SL1344_2847** | *hilC* | 108.19 | 343.477 | 105.507 | 245.981 | 1.03 | 1.4 |
| **SL1344_2848** | *orgC* | 332.168 | 198.057 | 89.723 | 119.933 | 3.7 | 1.65 |
| **SL1344_2849** | *orgB* | 150.724 | 158.816 | 173.367 | 207.242 | 0.869 | 0.766 |
| **SL1344_2850** | *orgA* | 50.414 | 40.586 | 245.158 | 276.928 | 0.206 | 0.147 |
| **SL1344_2851** | *prgK* | 174.775 | 172.876 | 761.791 | 753.08 | 0.229 | 0.23 |
| **SL1344_2852** | *prgJ* | 4884.116 | 4490.959 | 1467.171 | 1128.819 | 3.33 | 3.98 |
| **SL1344_2853** | *prgI* | 687.434 | 1400.174 | 357.828 | 607.257 | 1.92 | 2.31 |
| **SL1344_2854** | *prgH* | 123.309 | 168.928 | 300.658 | 349.947 | 0.41 | 0.483 |
| **SL1344_2855** | *hilD* | 69.569 | 391.744 | 133.655 | 314.299 | 0.521 | 1.25 |
| **SL1344_2856** | *hilA* | 56.274 | 95.402 | 30.149 | 48.605 | 1.87 | 1.96 |
| **SL1344_2857** | *iagB* | 29.536 | 125.762 | 35.619 | 73.688 | 0.829 | 1.71 |
| **SL1344_2858** | *sptP* | 286.578 | 189.476 | 179.912 | 116.435 | 1.59 | 1.63 |
| **SL1344_2859** | *sicP* | 115.678 | 134.831 | 93.207 | 130.184 | 1.24 | 1.04 |

| | | | | | | | |
|---|---|---|---|---|---|---|---|
| **SL1344_2860** | *iacP* | 151.195 | 95.066 | 122.351 | 133.712 | 1.24 | 0.711 |
| **SL1344_2861** | *sipA* | 568.904 | 211.996 | 134.555 | 79.151 | 4.23 | 2.68 |
| **SL1344_2862** | *sipD* | 122.404 | 103.877 | 142.117 | 98.59 | 0.861 | 1.05 |
| **SL1344_2863** | *sipC* | 1094.566 | 1124.12 | 599.636 | 470.367 | 1.83 | 2.39 |
| **SL1344_2864** | *sipB* | 704.121 | 507.755 | 371.21 | 347.079 | 1.9 | 1.46 |
| **SL1344_2865** | *sicA* | 416.857 | 480.656 | 462.777 | 473.672 | 0.901 | 1.01 |
| **SL1344_2866** | *spaS* | 5.859 | 8.574 | 46.986 | 25.353 | 0.125 | 0.338 |
| **SL1344_2867** | *spaR* | 9.417 | 3.607 | 28.058 | 25.736 | 0.336 | 0.14 |
| **SL1344_2868** | *spaQ* | 29.484 | 26.583 | 82.67 | 62.531 | 0.357 | 0.425 |
| **SL1344_2869** | *spaP* | 48.934 | 25.697 | 50.338 | 40.37 | 0.972 | 0.637 |
| **SL1344_2870** | *spaO* | 123.192 | 118.515 | 178.344 | 89.106 | 0.691 | 1.33 |
| **SL1344_2871** | *invJ* | 115.813 | 136.581 | 117.558 | 99.968 | 0.985 | 1.37 |
| **SL1344_2872** | *invI* | 11.466 | 17.005 | 125.285 | 108.478 | 0.0915 | 0.157 |
| **SL1344_2873** | *invC* | 54.764 | 23.881 | 141.763 | 151.211 | 0.386 | 0.158 |
| **SL1344_2874** | *invB* | 283.35 | 569.511 | 373.001 | 309.584 | 0.76 | 1.84 |
| **SL1344_2875** | *invA* | 29.425 | 62.171 | 70.848 | 88.021 | 0.415 | 0.706 |
| **SL1344_2876** | *invE* | 29.782 | 38.357 | 64.829 | 80.142 | 0.459 | 0.479 |
| **SL1344_2877** | *invG* | 139.837 | 172.329 | 60.224 | 110.759 | 2.32 | 1.56 |
| **SL1344_2878** | *invF* | 67.378 | 128.624 | 108.461 | 237.443 | 0.621 | 0.542 |
| **SL1344_2879** | *invH* | 56.528 | 356.346 | 77.334 | 70.249 | 0.731 | 5.07 |
| **SL1344_2880** | *SL1344-2880* | 98.526 | 299.596 | 146.872 | 139.431 | 0.671 | 2.15 |
| **SL1344_2882** | *SL1344-2882* | 13.577 | 22.674 | 17.099 | 23.144 | 0.794 | 0.98 |
| **SL1344_2884** | *SL1344-2884* | 87.063 | 514.016 | 115.039 | 272.365 | 0.757 | 1.89 |
| **SL1344_2885** | *SL1344-2885* | 27.579 | 30.532 | 201.61 | 191.37 | 0.137 | 0.16 |

| | | | | | | | |
|---|---|---|---|---|---|---|---|
| **SL1344_2886** | *pphB* | 3.424 | 5.591 | 9.711 | 12.697 | 0.353 | 0.44 |
| **SL1344_2888** | *mutS* | 108.734 | 51.676 | 88.455 | 71.426 | 1.23 | 0.723 |
| **SL1344_2891** | *SL1344-2891* | 3.553 | 5.103 | 6.454 | 10.599 | 0.551 | 0.481 |
| **SL1344_2892** | *SL1344-2892* | 17.319 | 23.81 | 38.691 | 44.287 | 0.448 | 0.538 |
| **SL1344_2893** | *SL1344-2893* | 11.515 | 15.807 | 16.508 | 21.128 | 0.698 | 0.748 |
| **SL1344_2894** | *hyi* | 19.809 | 12.14 | 19.849 | 26.463 | 0.998 | 0.459 |
| **SL1344_2895** | *ygbL* | 13.71 | 13.306 | 12.003 | 17.633 | 1.14 | 0.755 |
| **SL1344_2896** | *ygbK* | 10.873 | 11.633 | 15.438 | 25.059 | 0.704 | 0.464 |
| **SL1344_2897** | *ygbJ* | 33.505 | 21.79 | 60.279 | 46.526 | 0.556 | 0.468 |
| **SL1344_2898** | *ygbI* | 8.202 | 22.852 | 46.571 | 57.998 | 0.176 | 0.394 |
| **SL1344_2899** | *SL1344-2899* | 18.562 | 10.245 | 54.515 | 59.462 | 0.34 | 0.172 |
| **SL1344_2901** | *SL1344-2901* | 3.233 | 0.667 | 4.769 | 9.194 | 0.678 | 0.0725 |
| **SL1344_2903** | *rpoS* | 905.918 | 2638.444 | 3621.356 | 5341.914 | 0.25 | 0.494 |
| **SL1344_2904** | *nlpD* | 95.003 | 160.996 | 518.789 | 1470.964 | 0.183 | 0.109 |
| **SL1344_2905** | *pcm* | 17.088 | 23.65 | 112.041 | 136.242 | 0.153 | 0.174 |
| **SL1344_2906** | *surE* | 35.268 | 88.999 | 116.839 | 127.272 | 0.302 | 0.699 |
| **SL1344_2907** | *truD* | 43.747 | 29.476 | 190.883 | 147.974 | 0.229 | 0.199 |
| **SL1344_2908** | *ispF* | 71.033 | 80.634 | 77.06 | 73.393 | 0.922 | 1.1 |
| **SL1344_2909** | *ispD* | 26.975 | 21.239 | 119.876 | 85.969 | 0.225 | 0.247 |
| **SL1344_2910** | *ftsB* | 17.265 | 33.575 | 323.036 | 413.287 | 0.0534 | 0.0812 |
| **SL1344_2912** | *cysC* | 1.465 | 127.287 | 16.797 | 406.212 | 0.0872 | 0.313 |
| **SL1344_2913** | *cysN* | 7.728 | 377.518 | 39.924 | 500.048 | 0.194 | 0.755 |
| **SL1344_2914** | *cysD* | 81.01 | 792.683 | 89.664 | 509.378 | 0.903 | 1.56 |
| **SL1344_2915** | *iap* | 12.89 | 5.457 | 13.967 | 22.077 | 0.923 | 0.247 |

| | | | | | | | |
|---|---|---|---|---|---|---|---|
| **SL1344_2920** | *yghJ* | 23.7 | 31.666 | 14.283 | 20.252 | 1.66 | 1.56 |
| **SL1344_2921** | *SL1344-2921* | 52.336 | 37.709 | 14.311 | 16.636 | 3.66 | 2.27 |
| **SL1344_2922** | *SL1344-2922* | 4.942 | 10.004 | 6.261 | 11.942 | 0.789 | 0.838 |
| **SL1344_2924** | *sopD* | 41.262 | 48.699 | 25.773 | 21.213 | 1.6 | 2.3 |
| **SL1344_2925** | *cysH* | 1.691 | 304.198 | 7.607 | 339.787 | 0.222 | 0.895 |
| **SL1344_2926** | *cysI* | 9.503 | 376.638 | 14.27 | 394.511 | 0.666 | 0.955 |
| **SL1344_2927** | *cysJ* | 14.667 | 225.642 | 31.381 | 216.253 | 0.467 | 1.04 |
| **SL1344_2928** | *ptpS* | 98.168 | 48.346 | 102.095 | 64.877 | 0.962 | 0.745 |
| **SL1344_2929** | *SL1344-2929* | 7.937 | 13.553 | 54.616 | 59.568 | 0.145 | 0.228 |
| **SL1344_2930** | *ygcF* | 81.657 | 36.541 | 198.517 | 188.249 | 0.411 | 0.194 |
| **SL1344_2931** | *eno* | 3196.314 | 1978.324 | 2065.469 | 1180.202 | 1.55 | 1.68 |
| **SL1344_2932** | *pyrG* | 1302.493 | 367.929 | 538.721 | 432.429 | 2.42 | 0.851 |
| **SL1344_2933** | *mazG* | 75.674 | 60.039 | 74.547 | 91.252 | 1.02 | 0.658 |
| **SL1344_2936** | *SL1344-2936* | 161.927 | 250.151 | 149.341 | 226.896 | 1.08 | 1.1 |
| **SL1344_2937** | *relA* | 138.279 | 111.817 | 226.373 | 238.738 | 0.611 | 0.468 |
| **SL1344_2938** | *rumA* | 33.662 | 48.707 | 92.37 | 123.17 | 0.364 | 0.395 |
| **SL1344_2939** | *barA* | 11.272 | 13.175 | 21.71 | 27.45 | 0.519 | 0.48 |
| **SL1344_2940** | *SL1344-2940* | 25.48 | 90.04 | 36.751 | 80.279 | 0.693 | 1.12 |
| **SL1344_2941** | *gudD* | 21.409 | 135.89 | 15.983 | 109.751 | 1.34 | 1.24 |
| **SL1344_2942** | *ygcY* | 4.679 | 114.079 | 15.769 | 183.496 | 0.297 | 0.622 |
| **SL1344_2943** | *gudT* | 1.568 | 128.284 | 7.437 | 115.574 | 0.211 | 1.11 |
| **SL1344_2944** | *SL1344-2944* | 217.704 | 249.865 | 301.549 | 231.794 | 0.722 | 1.08 |
| **SL1344_2945** | *yqcB* | 15.649 | 9.556 | 74.339 | 73 | 0.211 | 0.131 |
| **SL1344_2946** | *yqcC* | 13.274 | 8.451 | 41.924 | 48.284 | 0.317 | 0.175 |

| | | | | | | | |
|---|---|---|---|---|---|---|---|
| **SL1344_2947** | *syd* | 29.489 | 62.913 | 264.547 | 273.008 | 0.111 | 0.23 |
| **SL1344_2948** | *queF* | 128.22 | 71.707 | 175.114 | 110.813 | 0.732 | 0.647 |
| **SL1344_2949** | *ygdH* | 306.943 | 447.694 | 345.289 | 498.934 | 0.889 | 0.897 |
| **SL1344_2950** | *sdaC* | 6927.76 | 2219.075 | 2176.736 | 1298.365 | 3.18 | 1.71 |
| **SL1344_2951** | *sdaB* | 723.011 | 630.837 | 1063.411 | 759.512 | 0.68 | 0.831 |
| **SL1344_2952** | *exo* | 21.69 | 16.422 | 56.047 | 54.402 | 0.387 | 0.302 |
| **SL1344_2953** | *fucO* | 7.316 | 16.28 | 9.545 | 14.73 | 0.766 | 1.11 |
| **SL1344_2954** | *fucA* | 4.659 | 5.039 | 5.089 | 13.283 | 0.916 | 0.379 |
| **SL1344_2955** | *fucP* | 1.753 | 7.334 | 8.328 | 14.999 | 0.21 | 0.489 |
| **SL1344_2956** | *fucI* | 84.392 | 34.011 | 12.755 | 19.359 | 6.62 | 1.76 |
| **SL1344_2957** | *fucK* | 7.592 | 5.369 | 8.689 | 17.977 | 0.874 | 0.299 |
| **SL1344_2958** | *fucU* | 5.738 | 17.528 | 7.287 | 15.033 | 0.787 | 1.17 |
| **SL1344_2959** | *fucR* | 72.016 | 56.924 | 258.706 | 240.258 | 0.278 | 0.237 |
| **SL1344_2960** | *ygdE* | 99.733 | 23.6 | 97.857 | 60.085 | 1.02 | 0.393 |
| **SL1344_2961** | *ygdD* | 23.02 | 21.815 | 115.852 | 67.713 | 0.199 | 0.322 |
| **SL1344_2962** | *gcvA* | 10.64 | 8.373 | 49.195 | 64.187 | 0.216 | 0.13 |
| **SL1344_2963** | *ygdI* | 577.924 | 2611.957 | 219.138 | 524.299 | 2.64 | 4.98 |
| **SL1344_2964** | *csdA* | 37.401 | 25.55 | 95.043 | 71.975 | 0.394 | 0.355 |
| **SL1344_2965** | *ygdK* | 12.932 | 9.805 | 50.372 | 52.387 | 0.257 | 0.187 |
| **SL1344_2967** | *ygdL* | 71.224 | 45.01 | 160.422 | 129.373 | 0.444 | 0.348 |
| **SL1344_2968** | *mltA* | 135.802 | 153.822 | 125.218 | 109.972 | 1.08 | 1.4 |
| **SL1344_2969** | *amiC* | 45.647 | 58.04 | 167.605 | 145.169 | 0.272 | 0.4 |
| **SL1344_2970** | *argA* | 3.466 | 9.141 | 12.163 | 20.276 | 0.285 | 0.451 |
| **SL1344_2971** | *recD* | 21.021 | 17.154 | 47.867 | 27.524 | 0.439 | 0.623 |

| | | | | | | | |
|---|---|---|---|---|---|---|---|
| **SL1344_2972** | *recB* | 12.654 | 11.145 | 38.53 | 40.469 | 0.328 | 0.275 |
| **SL1344_2973** | *ptr* | 22.58 | 14.009 | 30.892 | 33.05 | 0.731 | 0.424 |
| **SL1344_2974** | *recC* | 13.236 | 12.083 | 39.158 | 38.341 | 0.338 | 0.315 |
| **SL1344_2979** | *thyA* | 181.753 | 75.636 | 171.139 | 80.9 | 1.06 | 0.935 |
| **SL1344_2980** | *lgt* | 67.371 | 61.887 | 134.693 | 128.181 | 0.5 | 0.483 |
| **SL1344_2981** | *ptsP* | 26.897 | 33.33 | 143.495 | 164.818 | 0.187 | 0.202 |
| **SL1344_2982** | *nudH* | 96.093 | 93.129 | 260.95 | 266.514 | 0.368 | 0.349 |
| **SL1344_2983** | *mutH* | 22.368 | 30.883 | 233.487 | 186.656 | 0.0958 | 0.165 |
| **SL1344_2984** | *ygdQ* | 61.516 | 20.768 | 126.5 | 188.994 | 0.486 | 0.11 |
| **SL1344_2985** | *ygdR* | 55.951 | 459.998 | 87.395 | 285.509 | 0.64 | 1.61 |
| **SL1344_2986** | *tas* | 63.004 | 59.2 | 82.632 | 78.761 | 0.762 | 0.752 |
| **SL1344_2987** | *ygeD* | 19.485 | 11.478 | 95.936 | 73.22 | 0.203 | 0.157 |
| **SL1344_2988** | *aas* | 59.907 | 55.803 | 65.345 | 74.081 | 0.917 | 0.753 |
| **SL1344_2989** | *galR* | 74.611 | 89.703 | 163.082 | 184.713 | 0.458 | 0.486 |
| **SL1344_2990** | *SL1344-2990* | 14.755 | 17.088 | 73.576 | 92.059 | 0.201 | 0.186 |
| **SL1344_2991** | *lysA* | 2.531 | 6.032 | 14.246 | 20.082 | 0.178 | 0.3 |
| **SL1344_2992** | *lysR* | 4.87 | 10.465 | 15.777 | 14.621 | 0.309 | 0.716 |
| **SL1344_2993** | *ygeA* | 185.609 | 158.716 | 115.018 | 83.123 | 1.61 | 1.91 |
| **SL1344_2994** | *araE* | 0.876 | 4.985 | 17.934 | 24.537 | 0.0488 | 0.203 |
| **SL1344_2995** | *kduD* | 19.343 | 22.674 | 21.166 | 20.339 | 0.914 | 1.11 |
| **SL1344_2996** | *kduI* | 15.205 | 19.911 | 30.232 | 35.474 | 0.503 | 0.561 |
| **SL1344_2997** | *yqeF* | 242.15 | 158.428 | 761.098 | 868.879 | 0.318 | 0.182 |
| **SL1344_2998** | *SL1344-2998* | 10.414 | 14.014 | 58.077 | 49.953 | 0.179 | 0.281 |
| **SL1344_2999** | *SL1344-2999* | 3.922 | 8.168 | 27.456 | 31.988 | 0.143 | 0.255 |

| | | | | | | | |
|---|---|---|---|---|---|---|---|
| **SL1344_3000** | *SL1344-3000* | 62.708 | 28.978 | 85.612 | 48.24 | 0.732 | 0.601 |
| **SL1344_3001** | *yohL* | 15.612 | 22.175 | 168.837 | 159.421 | 0.0925 | 0.139 |
| **SL1344_3002** | *yohM* | 12.871 | 2.113 | 25.779 | 16.87 | 0.499 | 0.125 |
| **SL1344_3009** | *SL1344-3009* | 12.624 | 31.808 | 33.407 | 34.568 | 0.378 | 0.92 |
| **SL1344_3011** | *SL1344-3011* | 51.331 | 102.97 | 104.381 | 234.153 | 0.492 | 0.44 |
| **SL1344_3012** | *SL1344-3012* | 51.146 | 168.263 | 90.548 | 222.806 | 0.565 | 0.755 |
| **SL1344_3013** | *SL1344-3013* | 9.096 | 12.559 | 12.347 | 10.007 | 0.737 | 1.26 |
| **SL1344_3014** | *SL1344-3014* | 23.319 | 29.664 | 144.028 | 156.955 | 0.162 | 0.189 |
| **SL1344_3015** | *idi* | 52.147 | 136.665 | 191.287 | 202.066 | 0.273 | 0.676 |
| **SL1344_3016** | *lysS* | 1591.892 | 985.905 | 1434.852 | 779.361 | 1.11 | 1.27 |
| **SL1344_3017-1** | *prfB-1* | 267.134 | 129.066 | 443.887 | 323.624 | 0.602 | 0.399 |
| **SL1344_3017-2** | *prfB-2* | 801.641 | 383.725 | 1372.709 | 745.478 | 0.584 | 0.515 |
| **SL1344_3018** | *recJ* | 52.742 | 21.458 | 123.979 | 136.456 | 0.425 | 0.157 |
| **SL1344_3019** | *dsbC* | 297.726 | 192.374 | 155.049 | 196.834 | 1.92 | 0.977 |
| **SL1344_3020** | *xerD* | 27.848 | 30.484 | 134.177 | 160.464 | 0.208 | 0.19 |
| **SL1344_3021** | *fldB* | 106.595 | 69.455 | 271.354 | 289.034 | 0.393 | 0.24 |
| **SL1344_3022** | *ygfX* | 123.679 | 52.577 | 489.665 | 465.647 | 0.253 | 0.113 |
| **SL1344_3023** | *ygfY* | 1074.145 | 966.309 | 575.348 | 567.704 | 1.87 | 1.7 |
| **SL1344_3024** | *ygfZ* | 688.248 | 573.154 | 270.805 | 305.279 | 2.54 | 1.88 |
| **SL1344_3025** | *yqfA* | 33.633 | 35.763 | 122.731 | 119.978 | 0.274 | 0.298 |
| **SL1344_3026** | *yqfB* | 75.321 | 173.759 | 111.431 | 135.541 | 0.676 | 1.28 |
| **SL1344_3027** | *bglA* | 42.146 | 38.517 | 42.84 | 33.554 | 0.984 | 1.15 |
| **SL1344_3029** | *gcvP* | 713.195 | 802.575 | 431.249 | 553.564 | 1.65 | 1.45 |
| **SL1344_3030** | *gcvH* | 1438.575 | 2744.571 | 1409.961 | 1836.041 | 1.02 | 1.49 |

| | | | | | | | |
|---|---|---|---|---|---|---|---|
| **SL1344_3031** | *gcvT* | 990.573 | 947.452 | 627.019 | 592.481 | 1.58 | 1.6 |
| **SL1344_3032** | *visC* | 68.543 | 61.293 | 87.772 | 80.958 | 0.781 | 0.757 |
| **SL1344_3033** | *ubiH* | 40.618 | 19.212 | 177.659 | 149.646 | 0.229 | 0.128 |
| **SL1344_3034** | *pepP* | 226.679 | 177.343 | 158.482 | 176.508 | 1.43 | 1 |
| **SL1344_3035** | *ygfB* | 57.763 | 91.047 | 132.596 | 207.42 | 0.436 | 0.439 |
| **SL1344_3036** | *ygfE* | 698.852 | 680.419 | 1008.132 | 1006.132 | 0.693 | 0.676 |
| **SL1344_3038** | *serA* | 137.832 | 171.791 | 83.485 | 112.56 | 1.65 | 1.53 |
| **SL1344_3039** | *rpiA* | 937.334 | 1071.232 | 869.978 | 827.868 | 1.08 | 1.29 |
| **SL1344_3040** | *iciA* | 54.162 | 33.022 | 64.467 | 51.333 | 0.84 | 0.643 |
| **SL1344_3041** | *yggE* | 93.823 | 122.11 | 135.882 | 160.254 | 0.69 | 0.762 |
| **SL1344_3042** | *yggA* | 15.45 | 14.866 | 38.885 | 26.878 | 0.397 | 0.553 |
| **SL1344_3043** | *yggB* | 118.595 | 206.118 | 111.647 | 125.837 | 1.06 | 1.64 |
| **SL1344_3044** | *fba* | 1765.582 | 2043.283 | 1327.208 | 1257.299 | 1.33 | 1.63 |
| **SL1344_3045** | *pgk* | 1814.836 | 1412.025 | 1073.714 | 936.743 | 1.69 | 1.51 |
| **SL1344_3046** | *epd* | 143.322 | 142.149 | 437.359 | 407.599 | 0.328 | 0.349 |
| **SL1344_3047** | *SL1344-3047* | 13.062 | 38.322 | 9.928 | 29.855 | 1.32 | 1.28 |
| **SL1344_3048** | *SL1344-3048* | 2.965 | 10.338 | 9.781 | 21.757 | 0.303 | 0.475 |
| **SL1344_3052** | *tktA* | 1831.965 | 1009.528 | 777.76 | 760.249 | 2.36 | 1.33 |
| **SL1344_3053** | *yggG* | 81.11 | 83.257 | 112.106 | 77.379 | 0.724 | 1.08 |
| **SL1344_3054** | *speB* | 1506.661 | 462.412 | 838.254 | 217.819 | 1.8 | 2.12 |
| **SL1344_3055** | *SL1344-3055* | 4.119 | 5.191 | 7.563 | 9.634 | 0.545 | 0.539 |
| **SL1344_3057** | *SL1344-3057* | 5.638 | 7.018 | 10.809 | 12.329 | 0.522 | 0.569 |
| **SL1344_3058** | *SL1344-3058* | 5.588 | 7.223 | 13.463 | 12.791 | 0.415 | 0.565 |
| **SL1344_3059** | *SL1344-3059* | 2.09 | 4.064 | 9.052 | 6.599 | 0.231 | 0.616 |

| | | | | | | | |
|---|---|---|---|---|---|---|---|
| **SL1344_3060** | *SL1344-3060* | 314.44 | 208.871 | 178.54 | 210.775 | 1.76 | 0.991 |
| **SL1344_3062** | *speA* | 581.707 | 133.59 | 316.172 | 173.597 | 1.84 | 0.77 |
| **SL1344_3063** | *yqgB* | 22.871 | 20.097 | 327.464 | 220.54 | 0.0698 | 0.0911 |
| **SL1344_3065** | *metK* | 294.793 | 174.853 | 296.975 | 151.96 | 0.993 | 1.15 |
| **SL1344_3066** | *galP* | 278.991 | 369.853 | 229.165 | 209.657 | 1.22 | 1.76 |
| **SL1344_3067** | *sprT* | 9.509 | 6.829 | 40.592 | 43.6 | 0.234 | 0.157 |
| **SL1344_3068** | *endA* | 175.075 | 40.544 | 61.964 | 29.131 | 2.83 | 1.39 |
| **SL1344_3069** | *yggJ* | 148.972 | 55.812 | 255.763 | 182.953 | 0.582 | 0.305 |
| **SL1344_3070** | *gshB* | 370.152 | 247.043 | 205.977 | 126.021 | 1.8 | 1.96 |
| **SL1344_3071** | *yqgE* | 34.215 | 76.102 | 77.276 | 72.236 | 0.443 | 1.05 |
| **SL1344_3072** | *yqgF* | 85.598 | 49.426 | 81.31 | 86.684 | 1.05 | 0.57 |
| **SL1344_3073** | *SL1344-3073* | 4.739 | 3.861 | 31.512 | 29.282 | 0.15 | 0.132 |
| **SL1344_3074** | *pilT* | 2.896 | 6.379 | 9.938 | 8.947 | 0.291 | 0.713 |
| **SL1344_3075** | *yggS* | 158.02 | 137.876 | 255.926 | 194.014 | 0.617 | 0.711 |
| **SL1344_3076** | *yggT* | 215.48 | 156.437 | 271.563 | 222.258 | 0.793 | 0.704 |
| **SL1344_3077** | *yggU* | 72.416 | 36.699 | 237.467 | 226.417 | 0.305 | 0.162 |
| **SL1344_3078** | *yggV* | 78.328 | 102.604 | 156.521 | 141.409 | 0.5 | 0.726 |
| **SL1344_3079** | *yggW* | 24.886 | 12.743 | 49.176 | 36.3 | 0.506 | 0.351 |
| **SL1344_3081** | *ansB* | 1040.679 | 3332.977 | 887.521 | 2481.066 | 1.17 | 1.34 |
| **SL1344_3082** | *yggN* | 76.377 | 65.093 | 200.017 | 164.785 | 0.382 | 0.395 |
| **SL1344_3083** | *yggL* | 609.503 | 463.252 | 706.533 | 422.131 | 0.863 | 1.1 |
| **SL1344_3084** | *trmB* | 102.686 | 77.941 | 255.472 | 206.225 | 0.402 | 0.378 |
| **SL1344_3085** | *mutY* | 61.724 | 14.922 | 112.257 | 85.992 | 0.55 | 0.174 |
| **SL1344_3086** | *yggX* | 478.488 | 514.349 | 394.26 | 448.184 | 1.21 | 1.15 |

| | | | | | | | |
|---|---|---|---|---|---|---|---|
| **SL1344_3087** | *mltC* | 19.186 | 26.494 | 30.891 | 37.894 | 0.621 | 0.699 |
| **SL1344_3088** | *nupG* | 831.264 | 228.199 | 780.992 | 208.55 | 1.06 | 1.09 |
| **SL1344_3089** | *speC* | 8.175 | 9.936 | 19.33 | 19.742 | 0.423 | 0.503 |
| **SL1344_3092** | *ripA* | 3.973 | 9.901 | 6.271 | 7.203 | 0.634 | 1.37 |
| **SL1344_3093** | *ripB* | 15.583 | 22.931 | 12.355 | 13.097 | 1.26 | 1.75 |
| **SL1344_3094** | *ripC* | 5.056 | 7.666 | 9.269 | 12.735 | 0.545 | 0.602 |
| **SL1344_3095** | *SL1344-3095* | 35.22 | 47.824 | 34.252 | 43.406 | 1.03 | 1.1 |
| **SL1344_3096** | *SL1344-3096* | 1.055 | 3.636 | 28.55 | 74.426 | 0.0369 | 0.0488 |
| **SL1344_3098** | *SL1344-3098* | 56.685 | 257.566 | 33.746 | 115.981 | 1.68 | 2.22 |
| **SL1344_3100** | *SL1344-3100* | 2.486 | 4.399 | 11.183 | 11.751 | 0.222 | 0.374 |
| **SL1344_3101** | *SL1344-3101* | 32.044 | 27.39 | 33.07 | 22.889 | 0.969 | 1.2 |
| **SL1344_3106** | *SL1344-3106* | 6.15 | 34.379 | 13.189 | 33.102 | 0.466 | 1.04 |
| **SL1344_3107** | *SL1344-3107* | 1.077 | 14.935 | 5.545 | 9.848 | 0.194 | 1.52 |
| **SL1344_3108** | *exuT* | 178.989 | 340.471 | 124.416 | 217.461 | 1.44 | 1.57 |
| **SL1344_3109** | *uxuA* | 966.642 | 1225.129 | 434.82 | 564.03 | 2.22 | 2.17 |
| **SL1344_3110** | *uxuB* | 174.971 | 322.974 | 98.157 | 152.492 | 1.78 | 2.12 |
| **SL1344_3111** | *uxaC* | 218.093 | 338.325 | 158.536 | 213.126 | 1.38 | 1.59 |
| **SL1344_3112** | *SL1344-3112* | 272.607 | 354.815 | 299.189 | 322.11 | 0.911 | 1.1 |
| **SL1344_3113** | *gsp* | 94.8 | 62.783 | 70.217 | 61.039 | 1.35 | 1.03 |
| **SL1344_3114** | *yghU* | 47.519 | 60.254 | 63.333 | 71.728 | 0.75 | 0.84 |
| **SL1344_3117** | *hybG* | 15.928 | 42.616 | 14.394 | 51.271 | 1.11 | 0.831 |
| **SL1344_3119** | *hybE* | 21.91 | 22.813 | 86.343 | 98.792 | 0.254 | 0.231 |
| **SL1344_3120** | *hybD* | 17.818 | 12.505 | 61.836 | 127 | 0.288 | 0.0985 |
| **SL1344_3121** | *hybC* | 116.304 | 88.699 | 63.018 | 101.809 | 1.85 | 0.871 |

| | | | | | | | |
|---|---|---|---|---|---|---|---|
| **SL1344_3122** | *hybB* | 41.321 | 45.117 | 67.792 | 89.804 | 0.61 | 0.502 |
| **SL1344_3123** | *hybA* | 67.111 | 97.449 | 65.22 | 122.854 | 1.03 | 0.793 |
| **SL1344_3124** | *hypO* | 57.872 | 101.515 | 56.95 | 109.14 | 1.02 | 0.93 |
| **SL1344_3126** | *SL1344-3126* | 166.565 | 0.041 | 87.86 | 10.06 | 1.9 | 0.00411 |
| **SL1344_3127** | *yqhA* | 160.124 | 243.777 | 167.262 | 187.618 | 0.957 | 1.3 |
| **SL1344_3128** | *SL1344-3128* | 31.547 | 89.245 | 132.948 | 230.964 | 0.237 | 0.386 |
| **SL1344_3129** | *SL1344-3129* | 6.337 | 21.184 | 36.627 | 47.769 | 0.173 | 0.443 |
| **SL1344_3130** | *SL1344-3130* | 26.601 | 38.797 | 24.603 | 43.82 | 1.08 | 0.885 |
| **SL1344_3131** | *yghA* | 61.469 | 224.047 | 46.736 | 98.71 | 1.32 | 2.27 |
| **SL1344_3132** | *exbD* | 48.356 | 28.422 | 167.766 | 114.083 | 0.288 | 0.249 |
| **SL1344_3133** | *exbB* | 71.114 | 30.91 | 185.304 | 138.165 | 0.384 | 0.224 |
| **SL1344_3135** | *metC* | 34.231 | 38.534 | 24.558 | 18.562 | 1.39 | 2.08 |
| **SL1344_3136** | *yghB* | 76.684 | 199.735 | 217.114 | 197.167 | 0.353 | 1.01 |
| **SL1344_3137** | *yqhC* | 11.607 | 18.302 | 74.26 | 98.006 | 0.156 | 0.187 |
| **SL1344_3138** | *yqhD* | 267.493 | 133.822 | 107.402 | 69.815 | 2.49 | 1.92 |
| **SL1344_3139** | *dkgA* | 58.265 | 108.933 | 89.691 | 111.608 | 0.65 | 0.976 |
| **SL1344_3141** | *SL1344-3141* | 618.296 | 838.183 | 59.894 | 84.096 | 10.3 | 9.97 |
| **SL1344_3142** | *ygiR* | 48.075 | 13.028 | 74.159 | 31.893 | 0.648 | 0.408 |
| **SL1344_3143** | *SL1344-3143* | 16.423 | 11.752 | 9.252 | 12.433 | 1.78 | 0.945 |
| **SL1344_3145** | *ygiK* | 3.213 | 11.129 | 8.878 | 6.211 | 0.362 | 1.79 |
| **SL1344_3146** | *sufI* | 55.885 | 21.567 | 130.177 | 93.772 | 0.429 | 0.23 |
| **SL1344_3147** | *plsC* | 87.187 | 66.27 | 125.009 | 80.629 | 0.697 | 0.822 |
| **SL1344_3148** | *parC* | 179.024 | 102.137 | 204.642 | 150.178 | 0.875 | 0.68 |
| **SL1344_3150** | *ygiW* | 635.473 | 1247.749 | 349.48 | 323.246 | 1.82 | 3.86 |

| | | | | | | | |
|---|---|---|---|---|---|---|---|
| **SL1344_3151** | *ygiX* | 42.154 | 72.35 | 53.98 | 44.401 | 0.781 | 1.63 |
| **SL1344_3152** | *ygiY* | 69.411 | 15.821 | 17.098 | 18.931 | 4.06 | 0.836 |
| **SL1344_3153** | *mdaB* | 52.888 | 94.318 | 98.041 | 98.169 | 0.539 | 0.961 |
| **SL1344_3154** | *ygiN* | 469.797 | 205.575 | 167.262 | 141.312 | 2.81 | 1.45 |
| **SL1344_3155** | *parE* | 264.827 | 130.509 | 249.51 | 225.81 | 1.06 | 0.578 |
| **SL1344_3156** | *yqiA* | 188.161 | 91.513 | 374.799 | 274.816 | 0.502 | 0.333 |
| **SL1344_3157** | *cpdA* | 63.841 | 59.724 | 290.803 | 394.629 | 0.22 | 0.151 |
| **SL1344_3158** | *yqiB* | 26.449 | 11.9 | 166.923 | 141.129 | 0.158 | 0.0843 |
| **SL1344_3159** | *nudF* | 30.112 | 33.957 | 96.031 | 62.529 | 0.314 | 0.543 |
| **SL1344_3160** | *tolC* | 421.7 | 388.403 | 192.176 | 205.11 | 2.19 | 1.89 |
| **SL1344_3161** | *ygiB* | 115.041 | 279.171 | 317.349 | 521.176 | 0.363 | 0.536 |
| **SL1344_3162** | *ygiC* | 253.915 | 189.221 | 286.734 | 340.269 | 0.886 | 0.556 |
| **SL1344_3163** | *ygiD* | 6.411 | 5.157 | 64.179 | 44.779 | 0.0999 | 0.115 |
| **SL1344_3164** | *ygiE* | 34.186 | 42.272 | 86.872 | 81.377 | 0.394 | 0.519 |
| **SL1344_3165** | *assT* | 17.788 | 7.987 | 5.784 | 7.239 | 3.08 | 1.1 |
| **SL1344_3166** | *SL1344-3166* | 6.166 | 8.604 | 6.934 | 7.124 | 0.889 | 1.21 |
| **SL1344_3168** | *ribB* | 346.839 | 195.119 | 674.308 | 521.804 | 0.514 | 0.374 |
| **SL1344_3169** | *yqiC* | 160.811 | 299.067 | 223.892 | 256.537 | 0.718 | 1.17 |
| **SL1344_3170** | *glgS* | 23.396 | 115.96 | 137.223 | 174.663 | 0.17 | 0.664 |
| **SL1344_3172** | *yqiK* | 12.121 | 7.774 | 10.753 | 12.823 | 1.13 | 0.606 |
| **SL1344_3173** | *rfaE* | 329.903 | 98.332 | 230.397 | 169.387 | 1.43 | 0.581 |
| **SL1344_3174** | *glnE* | 35.487 | 21.31 | 68.936 | 91.555 | 0.515 | 0.233 |
| **SL1344_3175** | *ygiF* | 82.654 | 66.14 | 104.332 | 99.482 | 0.792 | 0.665 |
| **SL1344_3176** | *ygiM* | 233.312 | 283.256 | 136.607 | 201.844 | 1.71 | 1.4 |

| | | | | | | | |
|---|---|---|---|---|---|---|---|
| **SL1344_3177** | *cca* | 55 | 26.288 | 55.581 | 50.616 | 0.99 | 0.519 |
| **SL1344_3178** | *bacA* | 60.199 | 38.752 | 139.283 | 116.293 | 0.432 | 0.333 |
| **SL1344_3179** | *folB* | 169.919 | 128.36 | 183.652 | 175.913 | 0.925 | 0.73 |
| **SL1344_3180** | *ygiH* | 92.66 | 83.693 | 192.796 | 183.435 | 0.481 | 0.456 |
| **SL1344_3182** | *gcp* | 176.24 | 74.931 | 185.147 | 110.641 | 0.952 | 0.677 |
| **SL1344_3183** | *rpsU* | 2450.263 | 1797.525 | 5909.581 | 3742.569 | 0.415 | 0.48 |
| **SL1344_3184** | *dnaG* | 22.918 | 10.48 | 422.342 | 348.045 | 0.0543 | 0.0301 |
| **SL1344_3185** | *rpoD* | 682.721 | 340.29 | 880.296 | 337.313 | 0.776 | 1.01 |
| **SL1344_3186** | *mug* | 46.702 | 41.457 | 102.506 | 120.751 | 0.456 | 0.343 |
| **SL1344_3187** | *yqjH* | 37.844 | 27.279 | 65.43 | 27.296 | 0.578 | 0.999 |
| **SL1344_3188** | *yqjI* | 71.983 | 83.872 | 105.977 | 89.029 | 0.679 | 0.942 |
| **SL1344_3189** | *SL1344-3189* | 748.85 | 4.952 | 428.866 | 13.371 | 1.75 | 0.37 |
| **SL1344_3190** | *aer* | 372.874 | 8.9 | 373.028 | 16.962 | 1 | 0.525 |
| **SL1344_3191** | *oat* | 2.34 | 6.749 | 4.834 | 12.277 | 0.484 | 0.55 |
| **SL1344_3192** | *fadH* | 30.667 | 47.807 | 40.937 | 83.997 | 0.749 | 0.569 |
| **SL1344_3193** | *ygjO* | 60.808 | 21.717 | 118.275 | 64.541 | 0.514 | 0.336 |
| **SL1344_3194** | *ygjP* | 177.748 | 30.61 | 87.613 | 74.753 | 2.03 | 0.409 |
| **SL1344_3196** | *ygjR* | 327.91 | 334.319 | 97.874 | 135.077 | 3.35 | 2.48 |
| **SL1344_3197** | *ygjT* | 17.227 | 7.511 | 65.322 | 74.208 | 0.264 | 0.101 |
| **SL1344_3198** | *ygjU* | 238.251 | 562.91 | 363.082 | 703.672 | 0.656 | 0.8 |
| **SL1344_3199** | *yqjA* | 215.171 | 232.278 | 270.516 | 180.119 | 0.795 | 1.29 |
| **SL1344_3201** | *yqjC* | 110.047 | 242.591 | 206.892 | 529.248 | 0.532 | 0.458 |
| **SL1344_3201A** | *yqjD* | 684.411 | 1124.574 | 708.168 | 903.545 | 0.966 | 1.24 |
| **SL1344_3202** | *yqjE* | 95.245 | 236.284 | 913.561 | 1543.021 | 0.104 | 0.153 |

| | | | | | | | |
|---|---|---|---|---|---|---|---|
| **SL1344_3203** | *yqjK* | 27.229 | 23.354 | 260.211 | 564.414 | 0.105 | 0.0414 |
| **SL1344_3205** | *yqjG* | 65.971 | 58.166 | 29.559 | 34.198 | 2.23 | 1.7 |
| **SL1344_3206** | *yhaH* | 56.768 | 104.262 | 121.235 | 182.002 | 0.468 | 0.573 |
| **SL1344_3207** | *yhaJ* | 150.659 | 124.668 | 328.05 | 184.832 | 0.459 | 0.674 |
| **SL1344_3209** | *yhaL* | 196.597 | 360.719 | 100.791 | 200.904 | 1.95 | 1.8 |
| **SL1344_3210** | *yhaN* | 14.404 | 3.839 | 19.302 | 19.384 | 0.746 | 0.198 |
| **SL1344_3212** | *tdcG* | 8.933 | 14.451 | 31.194 | 35.409 | 0.286 | 0.408 |
| **SL1344_3213** | *tdcE* | 19.164 | 14.493 | 23.738 | 32.165 | 0.807 | 0.451 |
| **SL1344_3214** | *tdcD* | 10.429 | 22.73 | 9.546 | 18.379 | 1.09 | 1.24 |
| **SL1344_3215** | *tdcC* | 19.909 | 65.315 | 24.11 | 32.946 | 0.826 | 1.98 |
| **SL1344_3216** | *tdcB* | 8.072 | 45.554 | 29.832 | 60.374 | 0.271 | 0.755 |
| **SL1344_3217** | *tdcA* | 6.934 | 18.762 | 24.124 | 38.006 | 0.287 | 0.494 |
| **SL1344_3220** | *garK* | 8.006 | 96.274 | 15.888 | 198.494 | 0.504 | 0.485 |
| **SL1344_3221** | *garRb* | 33.683 | 540.124 | 18.906 | 417.064 | 1.78 | 1.3 |
| **SL1344_3222** | *garL* | 21.878 | 408.128 | 11.397 | 211.135 | 1.92 | 1.93 |
| **SL1344_3223** | *garD* | 16.907 | 215.314 | 12.95 | 121.54 | 1.31 | 1.77 |
| **SL1344_3224** | *SL1344-3224* | 8.742 | 28.414 | 51.872 | 44.353 | 0.169 | 0.641 |
| **SL1344_3225** | *agaR* | 49.825 | 61.252 | 143.367 | 102.659 | 0.348 | 0.597 |
| **SL1344_3226** | *gatY* | 7.339 | 2.944 | 5.366 | 4.129 | 1.37 | 0.713 |
| **SL1344_3230** | *SL1344-3230* | 8.283 | 5.187 | 13.187 | 21.765 | 0.628 | 0.238 |
| **SL1344_3232** | *gatB* | 3.946 | 15.514 | 1.509 | 8.314 | 2.61 | 1.87 |
| **SL1344_3233** | *gatC* | 3.317 | 2.475 | 9.287 | 7.356 | 0.357 | 0.337 |
| **SL1344_3235** | *gatR* | 127.643 | 241.941 | 332.208 | 367.931 | 0.384 | 0.658 |
| **SL1344_3236** | *yraL* | 62.072 | 42.592 | 104.539 | 59.216 | 0.594 | 0.719 |

| | | | | | | | |
|---|---|---|---|---|---|---|---|
| **SL1344_3237** | *yraM* | 72.841 | 112.636 | 71.192 | 84.963 | 1.02 | 1.33 |
| **SL1344_3239** | *yraO* | 29.347 | 60.885 | 91.333 | 183.694 | 0.321 | 0.331 |
| **SL1344_3240** | *yraP* | 161.551 | 207.725 | 87.24 | 157.708 | 1.85 | 1.32 |
| **SL1344_3241** | *yraR* | 27.075 | 13.523 | 33.538 | 40.804 | 0.807 | 0.331 |
| **SL1344_3242** | *yhbO* | 17.336 | 67.366 | 39.64 | 45.982 | 0.437 | 1.47 |
| **SL1344_3245** | *yhbS* | 421.29 | 428.912 | 459.543 | 406.818 | 0.917 | 1.05 |
| **SL1344_3246** | *yhbT* | 135.189 | 243.063 | 515.849 | 606.451 | 0.262 | 0.401 |
| **SL1344_3247** | *yhbU* | 6.419 | 20.215 | 12.019 | 23.669 | 0.534 | 0.854 |
| **SL1344_3248** | *yhbV* | 10.708 | 35.055 | 30.582 | 46.625 | 0.35 | 0.752 |
| **SL1344_3249** | *yhbW* | 45.512 | 44.133 | 36.624 | 40.778 | 1.24 | 1.08 |
| **SL1344_3252** | *mtr* | 19.68 | 12.714 | 33.835 | 22.096 | 0.582 | 0.575 |
| **SL1344_3253** | *deaD* | 51.396 | 25.445 | 133.051 | 62.865 | 0.386 | 0.405 |
| **SL1344_3254** | *nlpI* | 75.314 | 39.122 | 1238.465 | 1984.276 | 0.0608 | 0.0197 |
| **SL1344_3255** | *pnp* | 1118.596 | 732.025 | 888.594 | 744.948 | 1.26 | 0.983 |
| **SL1344_3256** | *rpsO* | 2611.349 | 5733.688 | 7364.558 | 4087.834 | 0.355 | 1.4 |
| **SL1344_3257** | *truB* | 92.08 | 38.797 | 343.019 | 206.812 | 0.268 | 0.188 |
| **SL1344_3258** | *rbfA* | 159.619 | 141.119 | 194.723 | 187.416 | 0.82 | 0.753 |
| **SL1344_3259** | *infB* | 367.96 | 389.669 | 526.054 | 507.936 | 0.699 | 0.767 |
| **SL1344_3260** | *nusA* | 730.812 | 371.515 | 935.559 | 629.824 | 0.781 | 0.59 |
| **SL1344_3261** | *yhbC* | 412.961 | 286.236 | 869.354 | 868.894 | 0.475 | 0.329 |
| **SL1344_3262** | *argG* | 28.012 | 22.117 | 20.481 | 22.991 | 1.37 | 0.962 |
| **SL1344_3264** | *secG* | 645.094 | 770.499 | 4185.847 | 4864.228 | 0.154 | 0.158 |
| **SL1344_3265** | *glmM* | 1373.769 | 647.015 | 728.783 | 437.171 | 1.89 | 1.48 |
| **SL1344_3266** | *folP* | 111.138 | 76.995 | 259.294 | 184.431 | 0.429 | 0.417 |

| | | | | | | | |
|---|---|---|---|---|---|---|---|
| **SL1344_3268** | *ftsH* | 1545.264 | 883.784 | 2397.343 | 1651.658 | 0.645 | 0.535 |
| **SL1344_3269** | *ftsJ* | 206.944 | 130.835 | 1056.39 | 593.442 | 0.196 | 0.22 |
| **SL1344_3270** | *yhbY* | 194.496 | 595.996 | 621.501 | 548.218 | 0.313 | 1.09 |
| **SL1344_3271** | *greA* | 913.728 | 271.942 | 487.659 | 549.371 | 1.87 | 0.495 |
| **SL1344_3272** | *dacB* | 40 | 28.034 | 90.429 | 83.26 | 0.442 | 0.337 |
| **SL1344_3273** | *obgE* | 146.396 | 63.556 | 212.973 | 127.243 | 0.687 | 0.499 |
| **SL1344_3274** | *yhbE* | 7.963 | 1.951 | 169.034 | 120.87 | 0.0471 | 0.0161 |
| **SL1344_3275** | *rpmA* | 6908.763 | 3580.08 | 15456.98 | 6704.095 | 0.447 | 0.534 |
| **SL1344_3276** | *rplU* | 2228.706 | 1506.278 | 15065.039 | 8803.883 | 0.148 | 0.171 |
| **SL1344_3277** | *ispB* | 213.27 | 200.075 | 314.169 | 334.449 | 0.679 | 0.598 |
| **SL1344_3278** | *nlp* | 6.783 | 25.272 | 15.681 | 23.087 | 0.433 | 1.09 |
| **SL1344_3279** | *murA* | 604.816 | 171.241 | 600.208 | 270.553 | 1.01 | 0.633 |
| **SL1344_3280** | *yrbA* | 247.851 | 79.589 | 201.397 | 171.883 | 1.23 | 0.463 |
| **SL1344_3281** | *yrbB* | 101.049 | 51.76 | 441.445 | 289.495 | 0.229 | 0.179 |
| **SL1344_3282** | *yrbC* | 137.562 | 184.812 | 204.569 | 276.384 | 0.672 | 0.669 |
| **SL1344_3283** | *yrbD* | 282.675 | 234.131 | 242.452 | 153.352 | 1.17 | 1.53 |
| **SL1344_3284** | *yrbE* | 106.898 | 41.525 | 199.579 | 122.994 | 0.536 | 0.338 |
| **SL1344_3285** | *yrbF* | 110.234 | 200.801 | 271.304 | 163.482 | 0.406 | 1.23 |
| **SL1344_3286** | *yrbG* | 41.676 | 15.407 | 310.935 | 206.347 | 0.134 | 0.0747 |
| **SL1344_3287** | *yrbH* | 99.017 | 87.938 | 191.301 | 165.651 | 0.518 | 0.531 |
| **SL1344_3288** | *yrbI* | 376.255 | 92.375 | 408.103 | 303.92 | 0.922 | 0.304 |
| **SL1344_3289** | *yrbK* | 131.235 | 87.034 | 287.605 | 262.693 | 0.456 | 0.331 |
| **SL1344_3290** | *yhbN* | 97.697 | 117.413 | 130.322 | 134.182 | 0.75 | 0.875 |
| **SL1344_3291** | *yhbG* | 187.123 | 141.758 | 317.047 | 279.056 | 0.59 | 0.508 |

| | | | | | | | |
|---|---|---|---|---|---|---|---|
| **SL1344_3292** | *rpoN* | 76.243 | 43.83 | 173.46 | 195.707 | 0.44 | 0.224 |
| **SL1344_3293** | *hpf* | 329.885 | 1335.153 | 331.288 | 600.861 | 0.996 | 2.22 |
| **SL1344_3294** | *ptsN* | 100.703 | 342.318 | 166.533 | 303.759 | 0.605 | 1.13 |
| **SL1344_3295** | *yhbJ* | 284.895 | 137.475 | 241.125 | 230.279 | 1.18 | 0.597 |
| **SL1344_3296** | *ptsO* | 9.974 | 30.896 | 48.839 | 85.29 | 0.204 | 0.362 |
| **SL1344_3297** | *yrbL* | 4.115 | 6.662 | 20.61 | 25.516 | 0.2 | 0.261 |
| **SL1344_3298** | *mtgA* | 54.079 | 9.797 | 127.929 | 108.689 | 0.423 | 0.0901 |
| **SL1344_3299** | *yhbL* | 895.971 | 457.843 | 216.037 | 202.341 | 4.15 | 2.26 |
| **SL1344_3300** | *arcB* | 113.931 | 111.39 | 126.037 | 128.509 | 0.904 | 0.867 |
| **SL1344_3301** | *yhcC* | 3.946 | 2.267 | 13.103 | 11.075 | 0.301 | 0.205 |
| **SL1344_3302** | *gltB* | 25.318 | 14.836 | 34.098 | 24.618 | 0.742 | 0.603 |
| **SL1344_3303** | *gltD* | 25.238 | 17.065 | 32.331 | 29.597 | 0.781 | 0.577 |
| **SL1344_3305** | *codB* | 12.698 | 13.886 | 25.724 | 17.912 | 0.494 | 0.775 |
| **SL1344_3306** | *codA* | 37.846 | 38.073 | 58.421 | 53.435 | 0.648 | 0.713 |
| **SL1344_3307** | *yhcH* | 28.965 | 635.882 | 81.333 | 820.479 | 0.356 | 0.775 |
| **SL1344_3308** | *nanK* | 54.397 | 538.579 | 102.943 | 937.509 | 0.528 | 0.574 |
| **SL1344_3309** | *nanE* | 324.196 | 4578.718 | 188.767 | 2039.502 | 1.72 | 2.25 |
| **SL1344_3310** | *nanT* | 65.03 | 1086.835 | 160.964 | 2004.232 | 0.404 | 0.542 |
| **SL1344_3311** | *nanA* | 1096.884 | 5957.643 | 454.156 | 4138.988 | 2.42 | 1.44 |
| **SL1344_3312** | *yhcK* | 40.584 | 311.506 | 156.31 | 221.67 | 0.26 | 1.41 |
| **SL1344_3313** | *sspB* | 227.325 | 306.707 | 762.049 | 893.748 | 0.298 | 0.343 |
| **SL1344_3314** | *sspA* | 1180.182 | 1101.433 | 1496.381 | 1528.526 | 0.789 | 0.721 |
| **SL1344_3315** | *SL1344-3315* | 62.024 | 78.139 | 45.314 | 51.633 | 1.37 | 1.51 |
| **SL1344_3316** | *rpsI* | 6150.34 | 2376.936 | 16464.97 | 6700.98 | 0.374 | 0.355 |

| | | | | | | | |
|---|---|---|---|---|---|---|---|
| **SL1344_3317** | *rplM* | 6300.544 | 2294.964 | 12151.285 | 4549.565 | 0.519 | 0.504 |
| **SL1344_3318** | *yhcM* | 71.98 | 62.7 | 102.715 | 83.863 | 0.701 | 0.748 |
| **SL1344_3319** | *yhcB* | 652.6 | 823.645 | 937.55 | 1069.122 | 0.696 | 0.77 |
| **SL1344_3320** | *degQ* | 250.018 | 213.113 | 166.633 | 161.263 | 1.5 | 1.32 |
| **SL1344_3321** | *degS* | 20.174 | 22.039 | 47.321 | 43.82 | 0.426 | 0.503 |
| **SL1344_3322** | *SL1344-3322* | 2.992 | 6.319 | 6.855 | 5.153 | 0.436 | 1.23 |
| **SL1344_3324** | *oadA2* | 8.199 | 0.613 | 2.704 | 2.056 | 3.03 | 0.298 |
| **SL1344_3326** | *ttdB* | 11.494 | 10.126 | 14.151 | 6.729 | 0.812 | 1.5 |
| **SL1344_3327** | *ttdA* | 4.538 | 9.75 | 5.018 | 5.588 | 0.904 | 1.74 |
| **SL1344_3329** | *SL1344-3329* | 27.197 | 36.564 | 101.713 | 141.551 | 0.267 | 0.258 |
| **SL1344_3330** | *SL1344-3330* | 53.369 | 200.501 | 126.755 | 168.369 | 0.421 | 1.19 |
| **SL1344_3331** | *mdh* | 14321.065 | 11694.441 | 3382.564 | 4030.113 | 4.23 | 2.9 |
| **SL1344_3332** | *argR* | 597.725 | 475.282 | 511.373 | 489.788 | 1.17 | 0.97 |
| **SL1344_3333** | *yhcN* | 25.337 | 31.434 | 70.054 | 71.804 | 0.362 | 0.438 |
| **SL1344_3334** | *SL1344-3334* | 71.831 | 31.845 | 51.011 | 24.269 | 1.41 | 1.31 |
| **SL1344_3335** | *yhcO* | 11.709 | 40.364 | 33.347 | 58.277 | 0.351 | 0.693 |
| **SL1344_3339** | *yhcS* | 14.512 | 13.092 | 39.85 | 40.558 | 0.364 | 0.323 |
| **SL1344_3340** | *tldD* | 116.669 | 118.214 | 147.928 | 140.073 | 0.789 | 0.844 |
| **SL1344_3341** | *yhdP* | 31.143 | 25.732 | 50.221 | 56.964 | 0.62 | 0.452 |
| **SL1344_3342** | *cafA* | 107.114 | 69.548 | 140.734 | 124.233 | 0.761 | 0.56 |
| **SL1344_3343** | *maf-b* | 17.639 | 9.619 | 120.559 | 103.635 | 0.146 | 0.0928 |
| **SL1344_3344** | *mreD* | 17.085 | 11.613 | 218.11 | 83.79 | 0.0783 | 0.139 |
| **SL1344_3345** | *mreC* | 16.246 | 7.429 | 84.209 | 80.022 | 0.193 | 0.0928 |
| **SL1344_3346** | *mreB* | 718.837 | 299.124 | 673.029 | 460.741 | 1.07 | 0.649 |

| | | | | | | | |
|---|---|---|---|---|---|---|---|
| **SL1344_3348** | *yhdA* | 19.731 | 14.719 | 152.157 | 77.208 | 0.13 | 0.191 |
| **SL1344_3349** | *yhdH* | 207.575 | 428.778 | 71.242 | 115.984 | 2.91 | 3.7 |
| **SL1344_3350** | *SL1344-3350* | 57.251 | 87.852 | 61.199 | 60.17 | 0.935 | 1.46 |
| **SL1344_3351** | *SL1344-3351* | 62.253 | 55.551 | 51.015 | 50.453 | 1.22 | 1.1 |
| **SL1344_3352** | *accB* | 2687.625 | 1221.061 | 1932.26 | 896.721 | 1.39 | 1.36 |
| **SL1344_3353** | *accC* | 696.17 | 472.521 | 1317.597 | 625.999 | 0.528 | 0.755 |
| **SL1344_3354** | *yhdT* | 16.808 | 25.473 | 70.209 | 45.671 | 0.239 | 0.558 |
| **SL1344_3355** | *panF* | 27.559 | 8.807 | 56.922 | 28.775 | 0.484 | 0.306 |
| **SL1344_3356** | *prmA* | 204.697 | 59.075 | 153.608 | 72.454 | 1.33 | 0.815 |
| **SL1344_3357** | *yhdG* | 12.868 | 14.506 | 579.183 | 371.418 | 0.0222 | 0.0391 |
| **SL1344_3358** | *fis* | 547.099 | 651.355 | 1732.956 | 764.197 | 0.316 | 0.852 |
| **SL1344_3359** | *yhdJ* | 4.281 | 2.46 | 29.726 | 15.245 | 0.144 | 0.161 |
| **SL1344_3360** | *yhdU* | 123.77 | 186.337 | 125.989 | 220.394 | 0.982 | 0.845 |
| **SL1344_3361** | *SL1344-3361* | 3.326 | 2.559 | 14.268 | 15.382 | 0.233 | 0.166 |
| **SL1344_3363** | *acrE* | 2.658 | 3.172 | 1.548 | 2.234 | 1.72 | 1.42 |
| **SL1344_3364** | *acrF* | 3.117 | 3.801 | 6.284 | 8.013 | 0.496 | 0.474 |
| **SL1344_3365** | *yhdV* | 8.799 | 20.835 | 39.071 | 40.08 | 0.225 | 0.52 |
| **SL1344_3366** | *yrdA* | 119.775 | 195.239 | 148.534 | 134.574 | 0.806 | 1.45 |
| **SL1344_3368** | *SL1344-3368* | 50.232 | 59.218 | 110.895 | 91.313 | 0.453 | 0.649 |
| **SL1344_3369** | *yrdC* | 51.963 | 86.303 | 99.331 | 97.728 | 0.523 | 0.883 |
| **SL1344_3370** | *yrdD* | 24.201 | 42.717 | 377.428 | 381.518 | 0.0641 | 0.112 |
| **SL1344_3371** | *smg* | 164.346 | 422.765 | 695.058 | 877.477 | 0.236 | 0.482 |
| **SL1344_3372** | *dprA* | 6.682 | 20.013 | 142.22 | 180.191 | 0.047 | 0.111 |
| **SL1344_3373** | *def* | 516.384 | 450.007 | 1110.955 | 727.072 | 0.465 | 0.619 |

| | | | | | | | |
|---|---|---|---|---|---|---|---|
| **SL1344_3374** | *fmt* | 92.038 | 67.017 | 479.403 | 303.766 | 0.192 | 0.221 |
| **SL1344_3375** | *fmu* | 26.752 | 11.548 | 36.62 | 29.033 | 0.731 | 0.398 |
| **SL1344_3376** | *sapG* | 66.33 | 30.923 | 48.778 | 41.58 | 1.36 | 0.744 |
| **SL1344_3377** | *mscL* | 903.215 | 712.416 | 565.504 | 397.023 | 1.6 | 1.79 |
| **SL1344_3378** | *arfA* | 328.419 | 269.566 | 260.185 | 386.413 | 1.26 | 0.698 |
| **SL1344_3379** | *zntR* | 53.358 | 44.389 | 179.209 | 226.464 | 0.298 | 0.196 |
| **SL1344_3380** | *yhdN* | 18.448 | 22.121 | 201.841 | 132.623 | 0.0914 | 0.167 |
| **SL1344_3381** | *rplQ* | 6111.048 | 4199.073 | 18842.14 | 8972.611 | 0.324 | 0.468 |
| **SL1344_3382** | *rpoA* | 2122.387 | 1592.659 | 10854.993 | 4741.566 | 0.196 | 0.336 |
| **SL1344_3383** | *rpsD* | 4412.792 | 1983.789 | 13763.478 | 4805.435 | 0.321 | 0.413 |
| **SL1344_3384** | *rpsK* | 6624.912 | 2254.817 | 13170.111 | 3925.339 | 0.503 | 0.574 |
| **SL1344_3385** | *rpsM* | 3938.83 | 1592.688 | 13307.444 | 5700.853 | 0.296 | 0.279 |
| **SL1344_3386** | *rpmJ* | 993.656 | 1732.507 | 5171.633 | 3809.715 | 0.192 | 0.455 |
| **SL1344_3387** | *secY* | 1049.945 | 513.428 | 8095.993 | 4172.445 | 0.13 | 0.123 |
| **SL1344_3388** | *rplO* | 5219.855 | 2862.209 | 22158.794 | 6891.021 | 0.236 | 0.415 |
| **SL1344_3389** | *rpmD* | 3096.523 | 3356.849 | 8863.276 | 5166.478 | 0.349 | 0.65 |
| **SL1344_3390** | *rpsE* | 10212.32 | 1445.452 | 17835.841 | 5619.178 | 0.573 | 0.257 |
| **SL1344_3391** | *rplR* | 7778.708 | 2170.339 | 12945.365 | 4786.383 | 0.601 | 0.453 |
| **SL1344_3392** | *rplF* | 4243.039 | 1340.172 | 11148.81 | 4099.986 | 0.381 | 0.327 |
| **SL1344_3393** | *rpsH* | 3237.013 | 1077.436 | 8457.02 | 2850.887 | 0.383 | 0.378 |
| **SL1344_3394** | *rpsN* | 4832.853 | 700.441 | 8827.622 | 2734.815 | 0.547 | 0.256 |
| **SL1344_3395** | *rplE* | 14720.324 | 3350.929 | 22595.46 | 5886.962 | 0.651 | 0.569 |
| **SL1344_3396** | *rplX* | 5281.453 | 1748.038 | 13666.303 | 6402.937 | 0.386 | 0.273 |
| **SL1344_3397** | *rplN* | 5798.661 | 3002.992 | 18752.963 | 8892.868 | 0.309 | 0.338 |

| | | | | | | | |
|---|---|---|---|---|---|---|---|
| **SL1344_3398** | *rpsQ* | 8560.202 | 6960.308 | 12437.241 | 12871.29 | 0.688 | 0.541 |
| **SL1344_3399** | *rpmC* | 4376.992 | 3531.437 | 15928.315 | 10494.492 | 0.275 | 0.337 |
| **SL1344_3400** | *rplP* | 6989.832 | 4261.176 | 8470.173 | 4757.169 | 0.825 | 0.896 |
| **SL1344_3401** | *rpsC* | 15580.092 | 4527.289 | 15649.689 | 7622.117 | 0.996 | 0.594 |
| **SL1344_3402** | *rplV* | 11114.853 | 5522.591 | 17549.56 | 7887.961 | 0.633 | 0.7 |
| **SL1344_3403** | *rpsS* | 3243.385 | 2200.085 | 7209.466 | 3533.488 | 0.45 | 0.623 |
| **SL1344_3404** | *rplB* | 2506.399 | 1424.143 | 7315.3 | 3395.684 | 0.343 | 0.419 |
| **SL1344_3405** | *rplW* | 4216.085 | 1476.263 | 15449.348 | 5228.611 | 0.273 | 0.282 |
| **SL1344_3406** | *rplD* | 11061.63 | 2630.155 | 13833.511 | 4036.787 | 0.8 | 0.652 |
| **SL1344_3407** | *rplC* | 4429.524 | 1405.341 | 8247.821 | 2543.536 | 0.537 | 0.553 |
| **SL1344_3408** | *rpsJ* | 8900.215 | 2574.561 | 14400.128 | 5461.005 | 0.618 | 0.471 |
| **SL1344_3410** | *bfr* | 89.598 | 270.374 | 131.495 | 278.944 | 0.681 | 0.969 |
| **SL1344_3412** | *tufA* | 8398.956 | 4932.718 | 6708.917 | 3161.203 | 1.25 | 1.56 |
| **SL1344_3413** | *fusA* | 8418.307 | 3051.469 | 6109.983 | 3210.445 | 1.38 | 0.95 |
| **SL1344_3414** | *rpsG* | 11272.568 | 3180.824 | 13600.551 | 3041.245 | 0.829 | 1.05 |
| **SL1344_3415** | *rpsL* | 1191.464 | 793.219 | 3758.896 | 1538.934 | 0.317 | 0.515 |
| **SL1344_3416** | *yheL* | 51.59 | 28.106 | 517.715 | 81.687 | 0.0996 | 0.344 |
| **SL1344_3417** | *yheM* | 15.918 | 10.098 | 84.133 | 125.296 | 0.189 | 0.0806 |
| **SL1344_3418** | *yheN* | 46.958 | 55.19 | 99.838 | 115.708 | 0.47 | 0.477 |
| **SL1344_3419** | *yheO* | 140.375 | 211.838 | 136.54 | 129.908 | 1.03 | 1.63 |
| **SL1344_3420** | *fkpA* | 2177.832 | 1215.33 | 1006.809 | 593.045 | 2.16 | 2.05 |
| **SL1344_3421** | *slyX* | 140.012 | 204.996 | 209.486 | 194.314 | 0.668 | 1.05 |
| **SL1344_3422** | *slyD* | 4670.045 | 3629.872 | 2700.324 | 1662.53 | 1.73 | 2.18 |
| **SL1344_3423** | *yheV* | 107.198 | 82.235 | 168.688 | 175.988 | 0.635 | 0.467 |

| | | | | | | | |
|---|---|---|---|---|---|---|---|
| **SL1344_3424** | *kefB* | 34.579 | 31.939 | 157.736 | 127.387 | 0.219 | 0.251 |
| **SL1344_3425** | *yheR* | 22.627 | 26.001 | 63.762 | 99.825 | 0.355 | 0.26 |
| **SL1344_3426** | *yheS* | 22.058 | 28.2 | 78.559 | 103.558 | 0.281 | 0.272 |
| **SL1344_3428** | *SL1344-3428* | 291.798 | 379.131 | 236.648 | 227.296 | 1.23 | 1.67 |
| **SL1344_3429** | *yheT* | 27.99 | 43.246 | 90.074 | 112.269 | 0.311 | 0.385 |
| **SL1344_3430** | *yheU* | 115.145 | 73.923 | 133.22 | 120.894 | 0.864 | 0.611 |
| **SL1344_3431** | *prkB* | 54.976 | 56.215 | 45.152 | 53.499 | 1.22 | 1.05 |
| **SL1344_3432** | *yhfA* | 479.255 | 947.091 | 513.821 | 564.889 | 0.933 | 1.68 |
| **SL1344_3433** | *crp* | 1024.165 | 505.378 | 1106.962 | 929.758 | 0.925 | 0.544 |
| **SL1344_3434** | *yhfK* | 38.556 | 38.343 | 81.983 | 61.559 | 0.47 | 0.623 |
| **SL1344_3435** | *argD* | 30.375 | 49.983 | 22.894 | 25.449 | 1.33 | 1.96 |
| **SL1344_3436** | *pabA* | 28.233 | 20.865 | 57.067 | 61.262 | 0.495 | 0.341 |
| **SL1344_3437** | *fic* | 11.78 | 40.046 | 32.454 | 84.887 | 0.363 | 0.472 |
| **SL1344_3438** | *yhfG* | 28.54 | 121.467 | 31.148 | 76.998 | 0.916 | 1.58 |
| **SL1344_3439** | *ppiA* | 226.137 | 348.535 | 400.202 | 430.029 | 0.565 | 0.81 |
| **SL1344_3440** | *yhfC* | 101.312 | 1.439 | 71.684 | 4.807 | 1.41 | 0.299 |
| **SL1344_3441** | *nirB* | 4.305 | 17.032 | 10.848 | 19.379 | 0.397 | 0.879 |
| **SL1344_3442** | *nirD* | 2.353 | 13.313 | 8.33 | 14.345 | 0.283 | 0.928 |
| **SL1344_3444** | *cysG* | 71.688 | 64.952 | 117.021 | 93.565 | 0.613 | 0.694 |
| **SL1344_3447** | *yhfL* | 41.54 | 517.518 | 74.618 | 109.017 | 0.557 | 4.75 |
| **SL1344_3448** | *trpS* | 461.657 | 419.024 | 265.978 | 283.289 | 1.74 | 1.48 |
| **SL1344_3449** | *gph* | 117.453 | 106.02 | 236.017 | 187.41 | 0.498 | 0.566 |
| **SL1344_3450** | *rpe* | 584.523 | 731.881 | 181.429 | 158.66 | 3.22 | 4.61 |
| **SL1344_3451** | *dam* | 92.646 | 23.161 | 141.397 | 75.223 | 0.655 | 0.308 |

| | | | | | | | |
|---|---|---|---|---|---|---|---|
| **SL1344_3452** | *damX* | 178.139 | 191.929 | 332.841 | 252.534 | 0.535 | 0.76 |
| **SL1344_3453** | *aroB* | 205.414 | 124.05 | 682.278 | 460.441 | 0.301 | 0.269 |
| **SL1344_3454** | *aroK* | 548.852 | 368.644 | 829.717 | 915.229 | 0.661 | 0.403 |
| **SL1344_3460** | *mrcA* | 21.362 | 10.268 | 81.753 | 56.755 | 0.261 | 0.181 |
| **SL1344_3461** | *nudE* | 131.193 | 201.652 | 219.372 | 349.778 | 0.598 | 0.577 |
| **SL1344_3462** | *igaA* | 45.319 | 92.002 | 88.117 | 110.76 | 0.514 | 0.831 |
| **SL1344_3463** | *yrfG* | 46.984 | 15.353 | 176.905 | 86.198 | 0.266 | 0.178 |
| **SL1344_3464** | *hslR* | 302.451 | 50.593 | 1007.671 | 157.042 | 0.3 | 0.322 |
| **SL1344_3465** | *hslO* | 560.44 | 130.739 | 632.273 | 149.308 | 0.886 | 0.876 |
| **SL1344_3466** | *yhgE* | 12.739 | 10.382 | 46.406 | 44.964 | 0.275 | 0.231 |
| **SL1344_3467** | *pckA* | 3641.513 | 4502.755 | 1188.797 | 2019.29 | 3.06 | 2.23 |
| **SL1344_3468** | *envZ* | 15.925 | 15.233 | 120.161 | 176.775 | 0.133 | 0.0862 |
| **SL1344_3469** | *ompR* | 309.455 | 546.532 | 705.486 | 576.363 | 0.439 | 0.948 |
| **SL1344_3470** | *greB* | 68.935 | 46.065 | 42.193 | 43.665 | 1.63 | 1.05 |
| **SL1344_3471** | *yhgF* | 191.822 | 75.472 | 177.7 | 100.496 | 1.08 | 0.751 |
| **SL1344_3472** | *feoA* | 13.76 | 36.696 | 90.862 | 54.614 | 0.151 | 0.672 |
| **SL1344_3473** | *feoB* | 100.75 | 32.177 | 88.005 | 33.98 | 1.14 | 0.947 |
| **SL1344_3474** | *yhgG* | 24.227 | 8.036 | 104.954 | 67.844 | 0.231 | 0.118 |
| **SL1344_3475** | *SL1344-3475* | 6.534 | 46.611 | 48.102 | 43.469 | 0.136 | 1.07 |
| **SL1344_3476** | *bioH* | 82.918 | 46.847 | 180.278 | 105.277 | 0.46 | 0.445 |
| **SL1344_3478** | *yhgI* | 458.859 | 140.294 | 407.203 | 151.789 | 1.13 | 0.924 |
| **SL1344_3479** | *gntT* | 181.134 | 30.576 | 188.271 | 32.863 | 0.962 | 0.93 |
| **SL1344_3480** | *malQ* | 20.557 | 27.058 | 42.962 | 66.908 | 0.478 | 0.404 |
| **SL1344_3481** | *malP* | 90.003 | 85.78 | 47.34 | 69.305 | 1.9 | 1.24 |

| | | | | | | | |
|---|---|---|---|---|---|---|---|
| **SL1344_3482** | *malT* | 59.326 | 120.809 | 118.413 | 206.673 | 0.501 | 0.585 |
| **SL1344_3483** | *SL1344-3483* | 18.23 | 36.722 | 135.835 | 118.178 | 0.134 | 0.311 |
| **SL1344_3484** | *SL1344-3484* | 452.009 | 463.639 | 193.628 | 177.311 | 2.33 | 2.61 |
| **SL1344_3485** | *rtcA* | 4.991 | 5.068 | 35.631 | 25.909 | 0.14 | 0.196 |
| **SL1344_3486** | *rtcB* | 53.605 | 15.916 | 66.034 | 25.072 | 0.812 | 0.635 |
| **SL1344_3487** | *SL1344-3487* | 20.836 | 1.488 | 52.31 | 9.942 | 0.398 | 0.15 |
| **SL1344_3488** | *rtcR* | 31.316 | 9.963 | 42.404 | 20.973 | 0.739 | 0.475 |
| **SL1344_3491** | *glpG* | 160.202 | 70.477 | 199.955 | 181.85 | 0.801 | 0.388 |
| **SL1344_3492** | *glpE* | 1284.536 | 1474.21 | 600.432 | 575.19 | 2.14 | 2.56 |
| **SL1344_3493** | *glpD* | 2274.178 | 3028.636 | 3064.532 | 3564.414 | 0.742 | 0.85 |
| **SL1344_3494** | *SL1344-3494* | 4.12 | 9.593 | 12.604 | 17.049 | 0.327 | 0.563 |
| **SL1344_3496** | *SL1344-3496* | 3.786 | 7.17 | 12.012 | 9.519 | 0.315 | 0.753 |
| **SL1344_3498** | *SL1344-3498* | 10.452 | 3.647 | 4.762 | 4.96 | 2.19 | 0.735 |
| **SL1344_3499** | *SL1344-3499* | 3.593 | 4.58 | 6.488 | 9.768 | 0.554 | 0.469 |
| **SL1344_3500** | *SL1344-3500* | 114.474 | 107.79 | 69.787 | 125.372 | 1.64 | 0.86 |
| **SL1344_3501** | *glgP* | 81.441 | 99.309 | 109.634 | 139.769 | 0.743 | 0.711 |
| **SL1344_3502** | *glgA* | 66.294 | 67.832 | 97.828 | 137.317 | 0.678 | 0.494 |
| **SL1344_3503** | *glgC* | 70.065 | 138.772 | 95.633 | 156.23 | 0.733 | 0.888 |
| **SL1344_3504** | *glgX* | 19.193 | 14.76 | 49.566 | 99.712 | 0.387 | 0.148 |
| **SL1344_3505** | *glgB* | 58.572 | 86.9 | 79.78 | 109.706 | 0.734 | 0.792 |
| **SL1344_3506** | *asd* | 480.239 | 246.216 | 200.416 | 187.555 | 2.4 | 1.31 |
| **SL1344_3507** | *gntU* | 15.803 | 2.333 | 62.543 | 28.452 | 0.253 | 0.082 |
| **SL1344_3508** | *gntK* | 27.269 | 4.331 | 39.869 | 8.739 | 0.684 | 0.496 |
| **SL1344_3509** | *gntR* | 142.459 | 98.003 | 102.056 | 85.5 | 1.4 | 1.15 |

| | | | | | | | |
|---|---|---|---|---|---|---|---|
| **SL1344_3510** | *yhhW* | 26.961 | 44.077 | 27.911 | 23.692 | 0.966 | 1.86 |
| **SL1344_3511** | *yhhX* | 65.981 | 69.594 | 28.867 | 27.44 | 2.29 | 2.54 |
| **SL1344_3512** | *yhhY* | 23.605 | 95.425 | 17.591 | 35.7 | 1.34 | 2.67 |
| **SL1344_3513** | *SL1344-3513* | 5.37 | 2.368 | 3.251 | 1.811 | 1.65 | 1.31 |
| **SL1344_3514** | *SL1344-3514* | 23.616 | 9.284 | 16.275 | 10.535 | 1.45 | 0.881 |
| **SL1344_3515** | *SL1344-3515* | 9.135 | 5.598 | 11.431 | 3.06 | 0.799 | 1.83 |
| **SL1344_3516** | *SL1344-3516* | 19.045 | 4.403 | 20.362 | 11.564 | 0.935 | 0.381 |
| **SL1344_3517** | *ggt* | 4.075 | 8.312 | 12.544 | 12.79 | 0.325 | 0.65 |
| **SL1344_3518** | *yhhA* | 6.814 | 58.56 | 7.22 | 28.006 | 0.944 | 2.09 |
| **SL1344_3519** | *ugpQ* | 28.599 | 63.615 | 83.969 | 82.228 | 0.341 | 0.774 |
| **SL1344_3520** | *ugpC* | 55.27 | 94.506 | 66.731 | 85.134 | 0.828 | 1.11 |
| **SL1344_3521** | *ugpE* | 6.787 | 17.689 | 13.727 | 30.524 | 0.494 | 0.58 |
| **SL1344_3522** | *ugpA* | 6.399 | 7.966 | 25.348 | 31.939 | 0.252 | 0.249 |
| **SL1344_3523** | *ugpB* | 135.783 | 327.401 | 92.149 | 212.983 | 1.47 | 1.54 |
| **SL1344_3525** | *phd* | 34.27 | 83.833 | 45.274 | 75.188 | 0.757 | 1.11 |
| **SL1344_3526** | *livF* | 10.197 | 3.43 | 13.052 | 7.213 | 0.781 | 0.475 |
| **SL1344_3527** | *livG* | 6.089 | 1.063 | 9.427 | 8.123 | 0.646 | 0.131 |
| **SL1344_3528** | *livM* | 2.918 | 2.076 | 15.818 | 9.176 | 0.184 | 0.226 |
| **SL1344_3529** | *livH* | 4.534 | 2.642 | 12.991 | 9.755 | 0.349 | 0.271 |
| **SL1344_3530** | *livK* | 36.85 | 9.621 | 15.628 | 14.224 | 2.36 | 0.676 |
| **SL1344_3531** | *yhhK* | 53.953 | 106.283 | 134.407 | 100.997 | 0.401 | 1.05 |
| **SL1344_3532** | *livJ* | 11.796 | 16.882 | 25.128 | 31.384 | 0.469 | 0.538 |
| **SL1344_3533** | *rpoH* | 449.462 | 977.199 | 1444.738 | 1844.095 | 0.311 | 0.53 |
| **SL1344_3534** | *ftsX* | 121.864 | 107.571 | 192.853 | 187.5 | 0.632 | 0.574 |

| | | | | | | | |
|---|---|---|---|---|---|---|---|
| **SL1344_3535** | *ftsE* | 153.339 | 170.511 | 264.018 | 298.603 | 0.581 | 0.571 |
| **SL1344_3536** | *ftsY* | 113.496 | 154.384 | 284.792 | 335.833 | 0.399 | 0.46 |
| **SL1344_3537** | *rsmD* | 30.837 | 33.042 | 94.377 | 76.668 | 0.327 | 0.431 |
| **SL1344_3538** | *yhhL* | 8.023 | 27.657 | 60.918 | 47.029 | 0.132 | 0.588 |
| **SL1344_3539** | *yhhM* | 29.719 | 19.035 | 34.514 | 34.129 | 0.861 | 0.558 |
| **SL1344_3540** | *yhhN* | 208.455 | 361.695 | 286.506 | 212.056 | 0.728 | 1.71 |
| **SL1344_3541** | *zntA* | 56.206 | 57.473 | 45.931 | 64.873 | 1.22 | 0.886 |
| **SL1344_3542** | *tcp* | 201.527 | 1.986 | 97.061 | 12.251 | 2.08 | 0.162 |
| **SL1344_3543** | *yhhP* | 25.988 | 12.443 | 46.915 | 76.566 | 0.554 | 0.163 |
| **SL1344_3544** | *yhhQ* | 5.244 | 2.349 | 17.544 | 8.931 | 0.299 | 0.263 |
| **SL1344_3545** | *SL1344-3545* | 320.901 | 804.674 | 217.492 | 288.283 | 1.48 | 2.79 |
| **SL1344_3546** | *yhhS* | 20.606 | 11.002 | 84.043 | 64.834 | 0.245 | 0.17 |
| **SL1344_3547** | *yhhT* | 7.216 | 7.644 | 11.742 | 13.264 | 0.615 | 0.576 |
| **SL1344_3548** | *acpT* | 12.37 | 19.737 | 103.873 | 117.887 | 0.119 | 0.167 |
| **SL1344_3549** | *nikR* | 129.432 | 85.957 | 136.413 | 89.257 | 0.949 | 0.963 |
| **SL1344_3550** | *yhhJ* | 6.945 | 17.172 | 15.038 | 19.107 | 0.462 | 0.899 |
| **SL1344_3551** | *yhiH* | 8.614 | 16.05 | 24.574 | 46.144 | 0.351 | 0.348 |
| **SL1344_3552** | *yhiI* | 20.951 | 54.901 | 43.963 | 81.343 | 0.477 | 0.675 |
| **SL1344_3553** | *yhiN* | 37.782 | 16.196 | 80.607 | 65.224 | 0.469 | 0.248 |
| **SL1344_3554** | *pitA* | 567.732 | 125.003 | 318.611 | 170.325 | 1.78 | 0.734 |
| **SL1344_3555** | *uspB* | 15.503 | 100.615 | 90.245 | 190.12 | 0.172 | 0.529 |
| **SL1344_3556** | *uspA* | 1841.285 | 4261.102 | 1029.112 | 1967.545 | 1.79 | 2.17 |
| **SL1344_3557** | *yhiP* | 240.796 | 878.504 | 273.205 | 468.803 | 0.881 | 1.87 |
| **SL1344_3558** | *yhiQ* | 53.345 | 5.825 | 304.301 | 79.067 | 0.175 | 0.0737 |

| | | | | | | | |
|---|---|---|---|---|---|---|---|
| **SL1344_3559** | *prlC* | 571.836 | 135.277 | 365.786 | 103.39 | 1.56 | 1.31 |
| **SL1344_3560** | *SL1344-3560* | 5.631 | 17.487 | 11.722 | 22.525 | 0.48 | 0.776 |
| **SL1344_3561** | *yhiR* | 68.393 | 41.878 | 116.581 | 127.003 | 0.587 | 0.33 |
| **SL1344_3562** | *gor* | 307.259 | 246.495 | 233.7 | 205.94 | 1.31 | 1.2 |
| **SL1344_3563** | *SL1344-3563* | 15.366 | 8.796 | 29.456 | 26.286 | 0.522 | 0.335 |
| **SL1344_3564** | *SL1344-3564* | 5.919 | 2.68 | 22.265 | 35.957 | 0.266 | 0.0745 |
| **SL1344_3565** | *SL1344-3565* | 20.851 | 13.347 | 14.574 | 9.832 | 1.43 | 1.36 |
| **SL1344_3566** | *SL1344-3566* | 51.447 | 27.821 | 59.663 | 28.159 | 0.862 | 0.988 |
| **SL1344_3567** | *SL1344-3567* | 17.018 | 18.706 | 38.331 | 33.212 | 0.444 | 0.563 |
| **SL1344_3568** | *treF* | 9.686 | 15.583 | 35.32 | 37.631 | 0.274 | 0.414 |
| **SL1344_3569** | *SL1344-3569* | 100.184 | 207.657 | 115.956 | 263.412 | 0.864 | 0.788 |
| **SL1344_3571** | *yhjB* | 3.239 | 10.04 | 5.587 | 5.533 | 0.58 | 1.81 |
| **SL1344_3572** | *yhjC* | 10.655 | 9.75 | 25.647 | 25.065 | 0.415 | 0.389 |
| **SL1344_3573** | *yhjD* | 12.828 | 15.865 | 25.984 | 28.831 | 0.494 | 0.55 |
| **SL1344_3574** | *yhjE* | 2.282 | 3.033 | 13.221 | 18.532 | 0.173 | 0.164 |
| **SL1344_3575** | *yhjG* | 13.643 | 42.08 | 28.068 | 35.031 | 0.486 | 1.2 |
| **SL1344_3576** | *yhjH* | 63.202 | 1.417 | 155.128 | 5.384 | 0.407 | 0.263 |
| **SL1344_3577** | *kdgK* | 65.432 | 74.75 | 73.688 | 74.772 | 0.888 | 1 |
| **SL1344_3578** | *yhjJ* | 42.765 | 39.176 | 88.689 | 99.039 | 0.482 | 0.396 |
| **SL1344_3579** | *dctA* | 1719.718 | 1484.259 | 1376.469 | 1222.581 | 1.25 | 1.21 |
| **SL1344_3580** | *yhjK* | 15.6 | 12.789 | 48.909 | 45.197 | 0.319 | 0.283 |
| **SL1344_3581** | *bcsC* | 24.242 | 11.174 | 48.093 | 41.369 | 0.504 | 0.27 |
| **SL1344_3582** | *bcsZ* | 66.34 | 19.732 | 84.406 | 59.924 | 0.786 | 0.329 |
| **SL1344_3583** | *bcsB* | 109.23 | 40.677 | 65.266 | 52.771 | 1.67 | 0.771 |

| | | | | | | | |
|---|---|---|---|---|---|---|---|
| **SL1344_3584** | *bcsA* | 37.952 | 20.834 | 64.01 | 53.682 | 0.593 | 0.388 |
| **SL1344_3585** | *bcsQ* | 162.804 | 70.189 | 155.457 | 134.605 | 1.05 | 0.521 |
| **SL1344_3586** | *bcsR* | 459.627 | 271.085 | 483.864 | 318.592 | 0.95 | 0.851 |
| **SL1344_3587** | *bcsE* | 27.338 | 26.438 | 52.303 | 65.015 | 0.523 | 0.407 |
| **SL1344_3589** | *bcsG* | 63.528 | 26.318 | 48.386 | 46.616 | 1.31 | 0.565 |
| **SL1344_3591** | *yhjV* | 7.154 | 2.409 | 12.582 | 6.198 | 0.569 | 0.389 |
| **SL1344_3592** | *dppF* | 21.775 | 18.448 | 19.37 | 60.805 | 1.12 | 0.303 |
| **SL1344_3593** | *dppD* | 12.332 | 25.646 | 23.166 | 41.355 | 0.532 | 0.62 |
| **SL1344_3594** | *dppC* | 13.045 | 23.728 | 33.182 | 53.284 | 0.393 | 0.445 |
| **SL1344_3595** | *dppB* | 5.339 | 27.142 | 26.284 | 73.295 | 0.203 | 0.37 |
| **SL1344_3596** | *dppA* | 110.329 | 252.557 | 69.658 | 200.481 | 1.58 | 1.26 |
| **SL1344_3598** | *SL1344-3598* | 10.681 | 6.046 | 9.43 | 8.489 | 1.13 | 0.712 |
| **SL1344_3599** | *SL1344-3599* | 6.577 | 4.386 | 15.361 | 13.481 | 0.428 | 0.325 |
| **SL1344_3600** | *yhjW* | 13.784 | 14.955 | 42.493 | 38.297 | 0.324 | 0.39 |
| **SL1344_3608** | *tag* | 144.121 | 123.303 | 174.775 | 182.463 | 0.825 | 0.676 |
| **SL1344_3609** | *yiaC* | 131.275 | 116.762 | 56.404 | 76.813 | 2.33 | 1.52 |
| **SL1344_3610** | *bisC* | 27.239 | 16.379 | 55.099 | 45.105 | 0.494 | 0.363 |
| **SL1344_3611** | *yiaD* | 151.744 | 112.829 | 62.922 | 49.24 | 2.41 | 2.29 |
| **SL1344_3612** | *yiaE* | 205.814 | 149.856 | 77.786 | 95.649 | 2.65 | 1.57 |
| **SL1344_3613** | *yiaF* | 177.75 | 149.628 | 474.463 | 443.346 | 0.375 | 0.337 |
| **SL1344_3614** | *yiaG* | 301.464 | 244.736 | 30.546 | 92.229 | 9.87 | 2.65 |
| **SL1344_3615** | *cspA* | 1209.177 | 2511.038 | 3133.548 | 2750.483 | 0.386 | 0.913 |
| **SL1344_3616** | *SL1344-3616* | 22.429 | 28.546 | 19.597 | 43.139 | 1.14 | 0.662 |
| **SL1344_3617** | *SL1344-3617* | 34.591 | 45.627 | 71.389 | 93.233 | 0.485 | 0.489 |

| | | | | | | | |
|---|---|---|---|---|---|---|---|
| **SL1344_3618** | *SL1344-3618* | 135.243 | 287.437 | 57.496 | 110.736 | 2.35 | 2.6 |
| **SL1344_3619** | *yafP* | 15.684 | 11.482 | 16.696 | 18.392 | 0.939 | 0.624 |
| **SL1344_3620** | *glyS* | 615.764 | 340.862 | 372.547 | 225.178 | 1.65 | 1.51 |
| **SL1344_3621** | *glyQ* | 670.934 | 398.357 | 543.915 | 260.32 | 1.23 | 1.53 |
| **SL1344_3622** | *SL1344-3622* | 3.907 | 36.368 | 39.272 | 120.575 | 0.0995 | 0.302 |
| **SL1344_3623** | *yiaH* | 3.804 | 6.146 | 14.754 | 14.444 | 0.258 | 0.426 |
| **SL1344_3625** | *xylB* | 46.135 | 17.064 | 38.281 | 30.759 | 1.21 | 0.555 |
| **SL1344_3626** | *xylA* | 31.678 | 30.694 | 44.484 | 44.337 | 0.712 | 0.692 |
| **SL1344_3627** | *xylR* | 12.1 | 19.039 | 23.955 | 30.864 | 0.505 | 0.617 |
| **SL1344_3628** | *bax* | 178.444 | 258.151 | 368.931 | 462.27 | 0.484 | 0.558 |
| **SL1344_3629** | *malS* | 3.444 | 3.153 | 4.701 | 5.806 | 0.733 | 0.543 |
| **SL1344_3630** | *avtA* | 93.5 | 27.948 | 52.946 | 28.18 | 1.77 | 0.992 |
| **SL1344_3631** | *ysaA* | 14.112 | 13.489 | 77.884 | 76.209 | 0.181 | 0.177 |
| **SL1344_3632** | *yiaJ* | 41.68 | 63.775 | 64.435 | 79.781 | 0.647 | 0.799 |
| **SL1344_3633** | *yiaK* | 7.229 | 4.902 | 7.893 | 5.736 | 0.916 | 0.855 |
| **SL1344_3635** | *SL1344-3635* | 4.124 | 3.864 | 4.303 | 4.613 | 0.958 | 0.838 |
| **SL1344_3643** | *SL1344-3643* | 7.254 | 15.838 | 27.76 | 34.876 | 0.261 | 0.454 |
| **SL1344_3644** | *SL1344-3644* | 8.413 | 16.588 | 9.528 | 20.124 | 0.883 | 0.824 |
| **SL1344_3645** | *aldB* | 18.308 | 61.082 | 23.662 | 47.886 | 0.774 | 1.28 |
| **SL1344_3646** | *SL1344-3646* | 5.839 | 38.175 | 18.449 | 38.324 | 0.316 | 0.996 |
| **SL1344_3647** | *selB* | 48.993 | 39.725 | 48.951 | 62.583 | 1 | 0.635 |
| **SL1344_3648** | *selA* | 29.3 | 27.707 | 85.794 | 95.552 | 0.342 | 0.29 |
| **SL1344_3649** | *yibF* | 130.247 | 320.672 | 178.915 | 269.024 | 0.728 | 1.19 |
| **SL1344_3650** | *mtlA* | 124.256 | 134.907 | 126.54 | 129.004 | 0.982 | 1.05 |

| | | | | | | | |
|---|---|---|---|---|---|---|---|
| **SL1344_3651** | *mtlD* | 146.054 | 126.274 | 166.451 | 151.028 | 0.877 | 0.836 |
| **SL1344_3652** | *mtlR* | 67.708 | 96.335 | 242.099 | 147.405 | 0.28 | 0.654 |
| **SL1344_3653** | *SL1344-3653* | 3.383 | 33.363 | 29.356 | 33.39 | 0.115 | 0.999 |
| **SL1344_3654** | *yibL* | 57.401 | 67.459 | 62.007 | 76.799 | 0.926 | 0.878 |
| **SL1344_3656** | *sadA* | 1.876 | 1.365 | 4.364 | 6.643 | 0.43 | 0.205 |
| **SL1344_3657** | *lctP* | 249.074 | 2043.102 | 253.879 | 1363.3 | 0.981 | 1.5 |
| **SL1344_3658** | *lctR* | 40.072 | 402.087 | 156.468 | 1148.23 | 0.256 | 0.35 |
| **SL1344_3659** | *lctD* | 430.563 | 2761.343 | 272.53 | 1326.77 | 1.58 | 2.08 |
| **SL1344_3660** | *yibK* | 92.163 | 52.953 | 113.726 | 184.708 | 0.81 | 0.287 |
| **SL1344_3661** | *SL1344-3661* | 18.612 | 13.198 | 114.551 | 84.47 | 0.162 | 0.156 |
| **SL1344_3662** | *SL1344-3662* | 6.478 | 24.379 | 11.498 | 16.463 | 0.563 | 1.48 |
| **SL1344_3663** | *SL1344-3663* | 2.561 | 9.866 | 7.357 | 14.581 | 0.348 | 0.677 |
| **SL1344_3665** | *cysE* | 104.85 | 132.484 | 478.937 | 287.499 | 0.219 | 0.461 |
| **SL1344_3666** | *gpsA* | 63.547 | 96.297 | 432.561 | 580.007 | 0.147 | 0.166 |
| **SL1344_3667** | *secB* | 3826.764 | 3242.492 | 1289.539 | 1714.992 | 2.97 | 1.89 |
| **SL1344_3668** | *grxC* | 730.769 | 876.448 | 1857.515 | 1363.033 | 0.393 | 0.643 |
| **SL1344_3669** | *yibN* | 1274.46 | 795.786 | 943.17 | 937.265 | 1.35 | 0.849 |
| **SL1344_3670** | *pmgI* | 119.27 | 71.235 | 66.673 | 52.755 | 1.79 | 1.35 |
| **SL1344_3671** | *yibP* | 96.905 | 23.786 | 114.616 | 114.172 | 0.845 | 0.208 |
| **SL1344_3672** | *yigQ* | 6.823 | 7.275 | 32.232 | 24.656 | 0.212 | 0.295 |
| **SL1344_3673** | *yibD* | 9.437 | 3.943 | 18.354 | 14.554 | 0.514 | 0.271 |
| **SL1344_3674** | *tdh* | 2092.977 | 1231.742 | 920.428 | 887.217 | 2.27 | 1.39 |
| **SL1344_3675** | *kbl* | 775.214 | 829.552 | 752.408 | 802.628 | 1.03 | 1.03 |
| **SL1344_3676** | *rfaD* | 237.158 | 359.281 | 426.337 | 480.493 | 0.556 | 0.748 |

| | | | | | | | |
|---|---|---|---|---|---|---|---|
| **SL1344_3677** | *rfaF* | 35.449 | 33.848 | 172.123 | 128.861 | 0.206 | 0.263 |
| **SL1344_3678** | *rfaC* | 69.246 | 51.265 | 173.348 | 118.765 | 0.399 | 0.432 |
| **SL1344_3679** | *rfaL* | 81.021 | 106.763 | 90.327 | 75.92 | 0.897 | 1.41 |
| **SL1344_3680** | *rfaK* | 130.062 | 118.592 | 85.07 | 80.828 | 1.53 | 1.47 |
| **SL1344_3681** | *rfaZ* | 74.833 | 104.131 | 45.311 | 49.79 | 1.65 | 2.09 |
| **SL1344_3682** | *rfaY* | 274.747 | 82.729 | 76.979 | 50.623 | 3.57 | 1.63 |
| **SL1344_3683** | *rfaJ* | 28.748 | 22.472 | 46.796 | 41.948 | 0.614 | 0.536 |
| **SL1344_3684** | *rfaI* | 139.813 | 58.429 | 230.109 | 113.955 | 0.608 | 0.513 |
| **SL1344_3685** | *rfaB* | 39.847 | 31.995 | 109.45 | 63.357 | 0.364 | 0.505 |
| **SL1344_3686** | *yibR* | 51.858 | 17.151 | 99.254 | 86.176 | 0.522 | 0.199 |
| **SL1344_3687** | *rfaP* | 62.532 | 50.121 | 171.304 | 113.562 | 0.365 | 0.441 |
| **SL1344_3688** | *rfaG* | 83.977 | 49.036 | 188.098 | 142.601 | 0.446 | 0.344 |
| **SL1344_3689** | *rfaQ* | 62.626 | 28.326 | 179.382 | 160.118 | 0.349 | 0.177 |
| **SL1344_3690** | *kdtA* | 98.379 | 77.123 | 179.153 | 129.748 | 0.549 | 0.594 |
| **SL1344_3691** | *coaD* | 136.024 | 142.136 | 166.814 | 177.563 | 0.815 | 0.8 |
| **SL1344_3692** | *mutM* | 50.498 | 7.558 | 206.112 | 52.686 | 0.245 | 0.143 |
| **SL1344_3693** | *rpmG* | 2052.776 | 1862.487 | 3696.824 | 2984.07 | 0.555 | 0.624 |
| **SL1344_3694** | *rpmB* | 1043.02 | 3115.201 | 4987.302 | 3726.804 | 0.209 | 0.836 |
| **SL1344_3695** | *radC* | 53.062 | 34.419 | 313.858 | 243.531 | 0.169 | 0.141 |
| **SL1344_3696** | *dfp* | 44.686 | 68.966 | 225.182 | 184.522 | 0.198 | 0.374 |
| **SL1344_3697** | *dut* | 583.116 | 311.466 | 901.232 | 319.15 | 0.647 | 0.976 |
| **SL1344_3698** | *slmA* | 62.07 | 51.158 | 149.011 | 151.877 | 0.417 | 0.337 |
| **SL1344_3699** | *pyrE* | 113.963 | 154.054 | 92.675 | 126.506 | 1.23 | 1.22 |
| **SL1344_3700** | *rph* | 203.754 | 125.892 | 495.686 | 309.975 | 0.411 | 0.406 |

| | | | | | | | |
|---|---|---|---|---|---|---|---|
| **SL1344_3701** | *yicC* | 246.438 | 253.978 | 380.985 | 297.422 | 0.647 | 0.854 |
| **SL1344_3702** | *SL1344-3702* | 8.969 | 7.135 | 21.137 | 24.066 | 0.424 | 0.296 |
| **SL1344_3703** | *SL1344-3703* | 11.812 | 2.237 | 5.202 | 3.591 | 2.27 | 0.623 |
| **SL1344_3704** | *SL1344-3704* | 80.746 | 35.441 | 41.757 | 41.902 | 1.93 | 0.846 |
| **SL1344_3705** | *ligB* | 4.038 | 4.841 | 16.369 | 15.044 | 0.247 | 0.322 |
| **SL1344_3706** | *gmk* | 32.538 | 25.072 | 388.86 | 299.135 | 0.0837 | 0.0838 |
| **SL1344_3707** | *rpoZ* | 6184.739 | 2353.881 | 3369.128 | 1176.615 | 1.84 | 2 |
| **SL1344_3708** | *spoT* | 48.852 | 26.023 | 355.024 | 237.461 | 0.138 | 0.11 |
| **SL1344_3709** | *spoU* | 106.035 | 37.658 | 110.746 | 87.534 | 0.957 | 0.43 |
| **SL1344_3710** | *recG* | 10.975 | 5.423 | 50.302 | 31.321 | 0.218 | 0.173 |
| **SL1344_3711** | *SL1344-3711* | 3.543 | 7.904 | 75.806 | 170.444 | 0.0467 | 0.0464 |
| **SL1344_3712** | *gltS* | 350.356 | 711.347 | 354.673 | 469.542 | 0.988 | 1.51 |
| **SL1344_3713** | *yicE* | 4.89 | 3.274 | 10.866 | 8.841 | 0.45 | 0.37 |
| **SL1344_3714** | *hilE* | 43.859 | 75.539 | 81.577 | 109.865 | 0.538 | 0.688 |
| **SL1344_3715** | *yicI* | 11.997 | 9.21 | 9.212 | 10.27 | 1.3 | 0.897 |
| **SL1344_3720** | *rhuM* | 97.916 | 50.328 | 63.258 | 83.275 | 1.55 | 0.604 |
| **SL1344_3723** | *misL* | 3.323 | 3.107 | 9.298 | 11.432 | 0.357 | 0.272 |
| **SL1344_3725** | *marT* | 5.327 | 12.759 | 13.355 | 20.93 | 0.399 | 0.61 |
| **SL1344_3726** | *slsA* | 17.471 | 18.878 | 28.842 | 28.013 | 0.606 | 0.674 |
| **SL1344_3727** | *cigR* | 48.589 | 64.62 | 93.487 | 73.594 | 0.52 | 0.878 |
| **SL1344_3728** | *mgtB* | 2.952 | 3.018 | 14.642 | 14.027 | 0.202 | 0.215 |
| **SL1344_3729** | *mgtC* | 7.91 | 11.728 | 52.424 | 123.256 | 0.151 | 0.0951 |
| **SL1344_3729A** | *mgtM* | 239.596 | 2371.027 | 761.211 | 1053.506 | 0.315 | 2.25 |
| **SL1344_3730** | *yicL* | 20.584 | 22.146 | 19.608 | 22.974 | 1.05 | 0.964 |

| | | | | | | | |
|---|---|---|---|---|---|---|---|
| **SL1344_3731** | *SL1344-3731* | 29.157 | 109.614 | 19.937 | 26.027 | 1.46 | 4.21 |
| **SL1344_3732** | *SL1344-3732* | 1.75 | 8.32 | 5.203 | 9.36 | 0.336 | 0.889 |
| **SL1344_3733** | *SL1344-3733* | 3.84 | 9.07 | 12.851 | 22.523 | 0.299 | 0.403 |
| **SL1344_3734** | *SL1344-3734* | 5.174 | 9.434 | 9.775 | 17.726 | 0.529 | 0.532 |
| **SL1344_3735** | *SL1344-3735* | 11.207 | 12.062 | 9.175 | 15.571 | 1.22 | 0.775 |
| **SL1344_3736** | *SL1344-3736* | 16.199 | 39.469 | 9.587 | 11.791 | 1.69 | 3.35 |
| **SL1344_3738** | *SL1344-3738* | 34.091 | 24.205 | 61.516 | 66.783 | 0.554 | 0.362 |
| **SL1344_3739** | *SL1344-3739* | 9.06 | 51.826 | 7.071 | 20.149 | 1.28 | 2.57 |
| **SL1344_3740** | *SL1344-3740* | 61.035 | 84.645 | 61.369 | 82.187 | 0.995 | 1.03 |
| **SL1344_3744** | *SL1344-3744* | 7.125 | 20.574 | 15.708 | 28.88 | 0.454 | 0.712 |
| **SL1344_3745** | *SL1344-3745* | 7.016 | 13.1 | 4.248 | 9.492 | 1.65 | 1.38 |
| **SL1344_3746** | *SL1344-3746* | 3.025 | 11.218 | 3.497 | 4.493 | 0.865 | 2.5 |
| **SL1344_3747** | *SL1344-3747* | 2.033 | 3.985 | 19.357 | 24.065 | 0.105 | 0.166 |
| **SL1344_3748** | *SL1344-3748* | 21.169 | 73.177 | 26.637 | 34.107 | 0.795 | 2.15 |
| **SL1344_3749** | *SL1344-3749* | 13.154 | 129.666 | 14.685 | 45.503 | 0.896 | 2.85 |
| **SL1344_3750** | *SL1344-3750* | 25.476 | 87.825 | 29.188 | 84.677 | 0.873 | 1.04 |
| **SL1344_3751** | *SL1344-3751* | 12.253 | 64.181 | 44.417 | 108.447 | 0.276 | 0.592 |
| **SL1344_3752** | *yicN* | 42.599 | 70.123 | 103.332 | 99.651 | 0.412 | 0.704 |
| **SL1344_3753** | *uhpT* | 5.145 | 2.59 | 22.968 | 10.543 | 0.224 | 0.246 |
| **SL1344_3754** | *uhpC* | 8.418 | 1.791 | 47.681 | 56.49 | 0.177 | 0.0317 |
| **SL1344_3755** | *uhpB* | 2.914 | 1.675 | 36.486 | 29.15 | 0.0799 | 0.0574 |
| **SL1344_3756** | *uhpA* | 91.947 | 23.249 | 148.461 | 91.479 | 0.619 | 0.254 |
| **SL1344_3757** | *SL1344-3757* | 67.717 | 61.179 | 50.9 | 39.44 | 1.33 | 1.55 |
| **SL1344_3758** | *SL1344-3758* | 77.353 | 48.55 | 72.609 | 43.603 | 1.07 | 1.11 |

| | | | | | | | |
|---|---|---|---|---|---|---|---|
| **SL1344_3759** | *SL1344-3759* | 527.029 | 163.886 | 68.726 | 58.071 | 7.67 | 2.82 |
| **SL1344_3760** | *SL1344-3760* | 15.213 | 17.481 | 31.191 | 37.007 | 0.488 | 0.472 |
| **SL1344_3761** | *ilvN* | 36.005 | 25.713 | 52.469 | 47.859 | 0.686 | 0.537 |
| **SL1344_3762** | *ilvB* | 41.285 | 21.546 | 53.476 | 44.135 | 0.772 | 0.488 |
| **SL1344_3763** | *ivbL* | 1218.567 | 406.066 | 3286.583 | 1028.699 | 0.371 | 0.395 |
| **SL1344_3766** | *emrD* | 28.024 | 14.924 | 200.049 | 182.025 | 0.14 | 0.082 |
| **SL1344_3767** | *SL1344-3767* | 10.956 | 15.032 | 59.076 | 58.666 | 0.185 | 0.256 |
| **SL1344_3768** | *dsdC* | 47.215 | 27.753 | 42.824 | 32.762 | 1.1 | 0.847 |
| **SL1344_3769** | *dsdX* | 247.838 | 12.913 | 275.591 | 26.383 | 0.899 | 0.489 |
| **SL1344_3770** | *dsdA* | 982.367 | 174.089 | 546.215 | 103.733 | 1.8 | 1.68 |
| **SL1344_3771** | *yidF* | 36.453 | 69.43 | 66.959 | 111.286 | 0.544 | 0.624 |
| **SL1344_3772** | *yidG* | 12.393 | 20.613 | 123.225 | 136.347 | 0.101 | 0.151 |
| **SL1344_3773** | *yidH* | 73.483 | 77.599 | 186.007 | 173.629 | 0.395 | 0.447 |
| **SL1344_3774** | *yidE* | 15.564 | 15.307 | 49.989 | 42.102 | 0.311 | 0.364 |
| **SL1344_3775** | *hslS* | 117.708 | 17.044 | 314.089 | 97.827 | 0.375 | 0.174 |
| **SL1344_3776** | *hslT* | 4512.02 | 389.989 | 3867.5 | 448.978 | 1.17 | 0.869 |
| **SL1344_3777** | *yidQ* | 72.407 | 179.568 | 115.419 | 209.694 | 0.627 | 0.856 |
| **SL1344_3778** | *yidR* | 26.196 | 17.241 | 26.582 | 32.337 | 0.985 | 0.533 |
| **SL1344_3779** | *ccmH1* | 2.892 | 11.728 | 9.819 | 40.599 | 0.295 | 0.289 |
| **SL1344_3780** | *ccmG1* | 6.577 | 31.329 | 16.957 | 78.126 | 0.388 | 0.401 |
| **SL1344_3781** | *ccmF1* | 4.81 | 13.414 | 12.43 | 60.547 | 0.387 | 0.222 |
| **SL1344_3782** | *ccmE1* | 8.879 | 34.861 | 16.726 | 123.967 | 0.531 | 0.281 |
| **SL1344_3783** | *ccmC1* | 7.78 | 53.735 | 20.398 | 137.472 | 0.381 | 0.391 |
| **SL1344_3784** | *ccmB1* | 3.784 | 8.283 | 24.113 | 118.87 | 0.157 | 0.0697 |

| | | | | | | | |
|---|---|---|---|---|---|---|---|
| **SL1344_3785** | *ccmA1* | 1.322 | 11.608 | 33.498 | 189.729 | 0.0395 | 0.0612 |
| **SL1344_3786** | *yhjA* | 5.873 | 70.4 | 15.503 | 31.531 | 0.379 | 2.23 |
| **SL1344_3790** | *torR* | 15.375 | 44.268 | 161.986 | 208.573 | 0.0949 | 0.212 |
| **SL1344_3792** | *torS* | 1.774 | 1.964 | 9.589 | 11.091 | 0.185 | 0.177 |
| **SL1344_3793** | *dgoT* | 3.849 | 2.389 | 7.018 | 7.373 | 0.548 | 0.324 |
| **SL1344_3794** | *dgoD* | 22.359 | 19.418 | 18.716 | 20.16 | 1.19 | 0.963 |
| **SL1344_3795** | *dgoA* | 7.28 | 9.906 | 9.279 | 10.524 | 0.784 | 0.941 |
| **SL1344_3796** | *dgoK* | 6.196 | 8.048 | 29.114 | 26.654 | 0.213 | 0.302 |
| **SL1344_3797** | *dgoR* | 58.508 | 26.617 | 72.411 | 61.288 | 0.808 | 0.434 |
| **SL1344_3798** | *yidA* | 205.011 | 42.131 | 89.393 | 66.277 | 2.29 | 0.636 |
| **SL1344_3799** | *SL1344-3799* | 0.375 | 3.501 | 3.122 | 3.886 | 0.12 | 0.901 |
| **SL1344_3800** | *SL1344-3800* | 6.048 | 7.064 | 7.204 | 10.004 | 0.84 | 0.706 |
| **SL1344_3801** | *SL1344-3801* | 26.331 | 27.224 | 48.588 | 69.275 | 0.542 | 0.393 |
| **SL1344_3802** | *gyrB* | 712.931 | 254.538 | 437.225 | 268.57 | 1.63 | 0.948 |
| **SL1344_3803** | *recF* | 13.614 | 4.813 | 153.846 | 79.133 | 0.0885 | 0.0608 |
| **SL1344_3804** | *dnaN* | 584.202 | 273.321 | 287.971 | 162.005 | 2.03 | 1.69 |
| **SL1344_3805** | *dnaA* | 103.812 | 57.534 | 152.475 | 136.944 | 0.681 | 0.42 |
| **SL1344_3806** | *rpmH* | 35667.755 | 16175.069 | 23993.154 | 10413.358 | 1.49 | 1.55 |
| **SL1344_3807** | *rnpA* | 42.587 | 32.499 | 518.909 | 358.653 | 0.0821 | 0.0906 |
| **SL1344_3809** | *yidC* | 624.218 | 287.242 | 525.505 | 333.351 | 1.19 | 0.862 |
| **SL1344_3810** | *thdF* | 56.4 | 23.02 | 57.107 | 29.271 | 0.988 | 0.786 |
| **SL1344_3812** | *SL1344-3812* | 11.064 | 10.878 | 24.267 | 22.672 | 0.456 | 0.48 |
| **SL1344_3813** | *SL1344-3813* | 17.708 | 85.899 | 18.993 | 20.846 | 0.932 | 4.12 |
| **SL1344_3815** | *yidZ* | 5.981 | 7.44 | 24.791 | 22.97 | 0.241 | 0.324 |

| | | | | | | | |
|---|---|---|---|---|---|---|---|
| **SL1344_3816** | *yieE* | 22.159 | 19.075 | 48.358 | 40.618 | 0.458 | 0.47 |
| **SL1344_3817** | *yieF* | 222.814 | 323.363 | 135.77 | 154.29 | 1.64 | 2.1 |
| **SL1344_3818** | *yieG* | 21.147 | 12.303 | 65.255 | 41.581 | 0.324 | 0.296 |
| **SL1344_3819** | *yieH* | 26.842 | 17.873 | 70.823 | 67.381 | 0.379 | 0.265 |
| **SL1344_3820** | *phoU* | 212.235 | 84.511 | 146.527 | 130.719 | 1.45 | 0.647 |
| **SL1344_3821** | *pstB* | 45.734 | 30.759 | 76.87 | 69.35 | 0.595 | 0.444 |
| **SL1344_3822** | *pstA* | 8.172 | 2.901 | 40.951 | 30.095 | 0.2 | 0.0964 |
| **SL1344_3823** | *pstC* | 18.868 | 12.045 | 70.414 | 43.95 | 0.268 | 0.274 |
| **SL1344_3824** | *pstS* | 84.385 | 82.201 | 291.486 | 314.766 | 0.289 | 0.261 |
| **SL1344_3825** | *SL1344-3825* | 2.068 | 4.307 | 17.53 | 19.762 | 0.118 | 0.218 |
| **SL1344_3827** | *SL1344-3827* | 26.062 | 12.42 | 24.343 | 22.405 | 1.07 | 0.554 |
| **SL1344_3828** | *glmS* | 295.681 | 250.489 | 374.752 | 357.054 | 0.789 | 0.702 |
| **SL1344_3829** | *glmU* | 347.351 | 328.844 | 518.098 | 392.575 | 0.67 | 0.838 |
| **SL1344_3831** | *atpC* | 1112.149 | 1395.246 | 3237.025 | 3983.347 | 0.344 | 0.35 |
| **SL1344_3832** | *atpD* | 5894.269 | 2785.724 | 3784.146 | 2764.323 | 1.56 | 1.01 |
| **SL1344_3833** | *atpG* | 1555.907 | 1788.235 | 3452.759 | 3319.663 | 0.451 | 0.539 |
| **SL1344_3834** | *atpA* | 2351.844 | 1857.924 | 3083.138 | 2653.759 | 0.763 | 0.7 |
| **SL1344_3835** | *atpH* | 2967.256 | 793.327 | 7427.976 | 5177.303 | 0.399 | 0.153 |
| **SL1344_3836** | *atpF* | 1242.83 | 2119.057 | 4562.028 | 5238.746 | 0.272 | 0.404 |
| **SL1344_3837** | *atpE* | 14059.155 | 18838.204 | 18781.993 | 9026.56 | 0.749 | 2.09 |
| **SL1344_3838** | *atpB* | 1005.796 | 1341.336 | 3090.212 | 2074.371 | 0.325 | 0.647 |
| **SL1344_3839** | *atpI* | 45.988 | 99.622 | 3544.668 | 2551.85 | 0.013 | 0.039 |
| **SL1344_3840** | *gidB* | 113.127 | 37.25 | 96.147 | 38.417 | 1.18 | 0.97 |
| **SL1344_3841** | *gidA* | 250.777 | 69.857 | 283.359 | 118.515 | 0.885 | 0.589 |

| | | | | | | | |
|---|---|---|---|---|---|---|---|
| **SL1344_3842** | *mioC* | 242.244 | 247.573 | 784.806 | 661.215 | 0.309 | 0.374 |
| **SL1344_3843** | *asnCb* | 18.313 | 45.051 | 59.814 | 45.46 | 0.306 | 0.991 |
| **SL1344_3844** | *asnA* | 16.036 | 11.234 | 44.468 | 33.821 | 0.361 | 0.332 |
| **SL1344_3845** | *yieM* | 8.957 | 6.02 | 37.658 | 27.389 | 0.238 | 0.22 |
| **SL1344_3846** | *yieN* | 19.771 | 25.536 | 38.452 | 60.875 | 0.514 | 0.419 |
| **SL1344_3847** | *kup* | 72.085 | 41.417 | 88.252 | 45.318 | 0.817 | 0.914 |
| **SL1344_3848** | *rbsD* | 16.208 | 3.725 | 74.756 | 24.985 | 0.217 | 0.149 |
| **SL1344_3849** | *rbsA* | 15.84 | 1.716 | 28.274 | 13.261 | 0.56 | 0.129 |
| **SL1344_3850** | *rbsC* | 48.042 | 7.957 | 162.661 | 110.432 | 0.295 | 0.0721 |
| **SL1344_3851** | *rbsB* | 323.277 | 363.62 | 332.995 | 409.222 | 0.971 | 0.889 |
| **SL1344_3852** | *rbsK* | 110.178 | 194.117 | 134.426 | 204.948 | 0.82 | 0.947 |
| **SL1344_3853** | *rbsR* | 26.368 | 31.525 | 67.378 | 119.394 | 0.391 | 0.264 |
| **SL1344_3857** | *yieP* | 45.256 | 52.68 | 91.511 | 99.528 | 0.495 | 0.529 |
| **SL1344_3858** | *yifA* | 57.8 | 89.734 | 57.077 | 103.75 | 1.01 | 0.865 |
| **SL1344_3859** | *yifE* | 614.469 | 1159.772 | 783.656 | 1012.515 | 0.784 | 1.15 |
| **SL1344_3860A** | *ilvX* | 64.998 | 85.36 | 711.214 | 588.351 | 0.0914 | 0.145 |
| **SL1344_3860B** | *ilvL* | 457.411 | 384.767 | 739.282 | 1178.168 | 0.619 | 0.327 |
| **SL1344_3861** | *ilvG* | 46.148 | 15.322 | 152.115 | 191.507 | 0.303 | 0.08 |
| **SL1344_3862** | *ilvM* | 14.969 | 16.68 | 45.317 | 101.903 | 0.33 | 0.164 |
| **SL1344_3863** | *ilvE* | 300.682 | 145.331 | 80.394 | 62.631 | 3.74 | 2.32 |
| **SL1344_3864** | *ilvD* | 95.044 | 37.777 | 87.213 | 54.85 | 1.09 | 0.689 |
| **SL1344_3865** | *ilvA* | 65.056 | 35.177 | 123.648 | 68.765 | 0.526 | 0.512 |
| **SL1344_3866** | *SL1344-3866* | 2.234 | 26.738 | 47.564 | 79.227 | 0.047 | 0.337 |
| **SL1344_3867** | *SL1344-3867* | 35.655 | 255.179 | 60.156 | 78.263 | 0.593 | 3.26 |

| | | | | | | | |
|---|---|---|---|---|---|---|---|
| **SL1344_3868** | *ilvY* | 22.598 | 42.896 | 39.474 | 49.283 | 0.572 | 0.87 |
| **SL1344_3869** | *ilvC* | 70.624 | 56.27 | 70.129 | 86.117 | 1.01 | 0.653 |
| **SL1344_3870** | *ppiC* | 47.649 | 113.369 | 290.801 | 234.243 | 0.164 | 0.484 |
| **SL1344_3872** | *rep* | 25.402 | 11.656 | 51.364 | 39.916 | 0.495 | 0.292 |
| **SL1344_3873** | *gppA* | 65.026 | 26.208 | 193.333 | 129.884 | 0.336 | 0.202 |
| **SL1344_3874** | *rhlB* | 264.925 | 147.272 | 372.743 | 269.328 | 0.711 | 0.547 |
| **SL1344_3875** | *trxA* | 1930.992 | 6167.261 | 2636.218 | 3019.935 | 0.732 | 2.04 |
| **SL1344_3876** | *rho* | 679.044 | 466.324 | 489.041 | 436.274 | 1.39 | 1.07 |
| **SL1344_3877** | *rfe* | 34.047 | 36.845 | 145.77 | 149.628 | 0.234 | 0.246 |
| **SL1344_3878** | *wzzE* | 119.293 | 142.214 | 275.369 | 227.447 | 0.433 | 0.625 |
| **SL1344_3879** | *rffE* | 51.501 | 57.436 | 123.783 | 129.125 | 0.416 | 0.445 |
| **SL1344_3880** | *rffD* | 316.406 | 125.487 | 201.543 | 149.017 | 1.57 | 0.842 |
| **SL1344_3881** | *rffG* | 98.489 | 65.114 | 207.086 | 128.38 | 0.476 | 0.507 |
| **SL1344_3882** | *rffH* | 32.097 | 22.674 | 52.249 | 63.94 | 0.614 | 0.355 |
| **SL1344_3883** | *rffC* | 22.427 | 12.142 | 106.154 | 70.511 | 0.211 | 0.172 |
| **SL1344_3884** | *rffA* | 99.024 | 78.666 | 107.874 | 93.423 | 0.918 | 0.842 |
| **SL1344_3885** | *wzxE* | 22.949 | 24.522 | 75.981 | 55.721 | 0.302 | 0.44 |
| **SL1344_3886** | *SL1344-3886* | 26.089 | 49.63 | 34.713 | 31.634 | 0.752 | 1.57 |
| **SL1344_3887** | *rffT* | 49.742 | 12.964 | 83.077 | 49.479 | 0.599 | 0.262 |
| **SL1344_3888** | *rffM* | 56.798 | 40.758 | 61.623 | 47.607 | 0.922 | 0.856 |
| **SL1344_3889** | *yifK* | 100.921 | 96.633 | 220.481 | 136.664 | 0.458 | 0.707 |
| **SL1344_3890** | *hemY* | 132.668 | 142.129 | 152.925 | 184.395 | 0.868 | 0.771 |
| **SL1344_3891** | *hemX* | 84.643 | 115.404 | 125.416 | 136.869 | 0.675 | 0.843 |
| **SL1344_3892** | *hemD* | 73.334 | 26.805 | 136.402 | 146.933 | 0.538 | 0.182 |

| | | | | | | | |
|---|---|---|---|---|---|---|---|
| **SL1344_3893** | *hemC* | 234.798 | 157.792 | 423.809 | 391.03 | 0.554 | 0.404 |
| **SL1344_3894** | *cyaA* | 29.609 | 27.187 | 81.253 | 90.649 | 0.364 | 0.3 |
| **SL1344_3898** | *cyaY* | 307.405 | 316.161 | 269.093 | 325.925 | 1.14 | 0.97 |
| **SL1344_3900** | *yifL* | 510.987 | 372.116 | 550.98 | 336.844 | 0.927 | 1.1 |
| **SL1344_3901** | *dapF* | 177.44 | 168.657 | 342.834 | 313.186 | 0.518 | 0.539 |
| **SL1344_3902** | *yigA* | 133.522 | 82.24 | 285.924 | 238.749 | 0.467 | 0.344 |
| **SL1344_3903** | *xerC* | 19.535 | 14.689 | 91.212 | 87.049 | 0.214 | 0.169 |
| **SL1344_3904** | *yigB* | 26.584 | 38.232 | 69.384 | 81.439 | 0.383 | 0.469 |
| **SL1344_3905** | *uvrD* | 52.9 | 34.781 | 95.545 | 97.241 | 0.554 | 0.358 |
| **SL1344_3906** | *corA* | 173.599 | 130.177 | 977.339 | 1064.987 | 0.178 | 0.122 |
| **SL1344_3909** | *rarD* | 24.012 | 27.9 | 56.132 | 80.323 | 0.428 | 0.347 |
| **SL1344_3910** | *yigI* | 138.12 | 322.051 | 146.76 | 247.957 | 0.941 | 1.3 |
| **SL1344_3911** | *pldA* | 219.978 | 202.607 | 200.369 | 175.175 | 1.1 | 1.16 |
| **SL1344_3912** | *recQ* | 18.578 | 18.735 | 153.026 | 145.505 | 0.121 | 0.129 |
| **SL1344_3913** | *rhtC* | 41.941 | 47.319 | 85.766 | 86.203 | 0.489 | 0.549 |
| **SL1344_3914** | *rhtB* | 10.676 | 13.801 | 22.509 | 13.511 | 0.474 | 1.02 |
| **SL1344_3915** | *pldB* | 9.255 | 12.106 | 85.287 | 102.754 | 0.109 | 0.118 |
| **SL1344_3916** | *yigL* | 86.242 | 58.68 | 104.796 | 97.832 | 0.823 | 0.6 |
| **SL1344_3917** | *yigM* | 27.164 | 3.477 | 55.595 | 14.722 | 0.489 | 0.236 |
| **SL1344_3918** | *metR* | 8.439 | 6.631 | 20.288 | 10.29 | 0.416 | 0.644 |
| **SL1344_3919** | *metE* | 29.166 | 7.478 | 26.648 | 29.239 | 1.09 | 0.256 |
| **SL1344_3920** | *SL1344-3920* | 12.773 | 2.935 | 25.228 | 13.475 | 0.506 | 0.218 |
| **SL1344_3921** | *dlhH* | 37.715 | 45.431 | 27.421 | 56.804 | 1.38 | 0.8 |
| **SL1344_3922** | *udp* | 6771.851 | 411.252 | 4861.407 | 327.493 | 1.39 | 1.26 |

| | | | | | | | |
|---|---|---|---|---|---|---|---|
| **SL1344_3923** | *yigN* | 43.144 | 54.522 | 99.586 | 103.474 | 0.433 | 0.527 |
| **SL1344_3924** | *ubiE* | 466.508 | 348.834 | 423.559 | 419.388 | 1.1 | 0.832 |
| **SL1344_3925** | *yigP* | 130.306 | 125.491 | 462.276 | 330.564 | 0.282 | 0.38 |
| **SL1344_3926** | *aarF* | 46.822 | 75.4 | 165.602 | 175.558 | 0.283 | 0.429 |
| **SL1344_3927** | *tatA* | 907.647 | 876.541 | 1773.817 | 1174.617 | 0.512 | 0.746 |
| **SL1344_3928** | *tatB* | 59.41 | 102.094 | 630.528 | 867.663 | 0.0942 | 0.118 |
| **SL1344_3929** | *tatC* | 657.437 | 899.538 | 239.864 | 217.299 | 2.74 | 4.14 |
| **SL1344_3930** | *tatD* | 18.391 | 25.497 | 45.535 | 39.386 | 0.404 | 0.647 |
| **SL1344_3931** | *rfaH* | 25.784 | 31.855 | 74.029 | 107 | 0.348 | 0.298 |
| **SL1344_3932** | *yigC* | 194.833 | 134.893 | 377.126 | 391.619 | 0.517 | 0.344 |
| **SL1344_3933** | *fre* | 397.412 | 352.025 | 471.662 | 384.209 | 0.843 | 0.916 |
| **SL1344_3934** | *SL1344-3934* | 9.996 | 8.652 | 18.353 | 18.598 | 0.545 | 0.465 |
| **SL1344_3935** | *fadA* | 158.208 | 511.913 | 119.657 | 474.187 | 1.32 | 1.08 |
| **SL1344_3936** | *fadB* | 272.267 | 982.239 | 320.187 | 741.661 | 0.85 | 1.32 |
| **SL1344_3937** | *pepQ* | 423.337 | 310.845 | 188.465 | 245.056 | 2.25 | 1.27 |
| **SL1344_3938** | *yigZ* | 10.203 | 18.471 | 110.847 | 162.214 | 0.092 | 0.114 |
| **SL1344_3939** | *trkH* | 23.319 | 18.27 | 89.654 | 88.722 | 0.26 | 0.206 |
| **SL1344_3940** | *hemG* | 431.598 | 320.672 | 197.589 | 179.393 | 2.18 | 1.79 |
| **SL1344_3941** | *mobB* | 5.277 | 21.751 | 27.923 | 36.268 | 0.189 | 0.6 |
| **SL1344_3942** | *mobA* | 57.778 | 78.137 | 81.732 | 78.736 | 0.707 | 0.992 |
| **SL1344_3943** | *yihD* | 240.505 | 389.989 | 204.962 | 265.878 | 1.17 | 1.47 |
| **SL1344_3944** | *rodA* | 88.162 | 79.599 | 301.405 | 290.979 | 0.293 | 0.274 |
| **SL1344_3945** | *dsbA* | 1331.494 | 1762.339 | 590.585 | 630.664 | 2.25 | 2.79 |
| **SL1344_3946** | *yihG* | 44.542 | 11.374 | 46.764 | 40.883 | 0.952 | 0.278 |

| | | | | | | | |
|---|---|---|---|---|---|---|---|
| **SL1344_3947** | *polA* | 310.436 | 181.073 | 156.305 | 131.131 | 1.99 | 1.38 |
| **SL1344_3948** | *engB* | 548.041 | 534.307 | 321.979 | 259.64 | 1.7 | 2.06 |
| **SL1344_3950** | *yihI* | 159.917 | 151.202 | 529.43 | 271.684 | 0.302 | 0.557 |
| **SL1344_3951** | *hemN* | 125.799 | 116.636 | 150.254 | 153.801 | 0.837 | 0.758 |
| **SL1344_3951A** | *yshB* | 110.923 | 362.78 | 1943.206 | 1277.775 | 0.0571 | 0.284 |
| **SL1344_3952** | *glnG* | 18.178 | 5.162 | 19.878 | 15.965 | 0.914 | 0.323 |
| **SL1344_3953** | *glnL* | 3.27 | 1.037 | 30.244 | 24.755 | 0.108 | 0.0419 |
| **SL1344_3954** | *glnA* | 236.274 | 171.259 | 84.648 | 74.611 | 2.79 | 2.3 |
| **SL1344_3955** | *typA* | 795.262 | 161.028 | 618.86 | 292.665 | 1.29 | 0.55 |
| **SL1344_3961** | *SL1344-3961* | 14.223 | 21.632 | 12.327 | 13.186 | 1.15 | 1.64 |
| **SL1344_3964** | *yihP* | 2.311 | 1.918 | 12.336 | 9.283 | 0.187 | 0.207 |
| **SL1344_3965** | *yihQ* | 6.48 | 5.844 | 9.361 | 7.917 | 0.692 | 0.738 |
| **SL1344_3967** | *yihS* | 5.862 | 5.258 | 8.253 | 6.677 | 0.71 | 0.787 |
| **SL1344_3968** | *yihT* | 6.532 | 20.507 | 6.932 | 6.904 | 0.942 | 2.97 |
| **SL1344_3969** | *yihU* | 4.421 | 3.337 | 3.836 | 4.798 | 1.15 | 0.695 |
| **SL1344_3970** | *yihV* | 6.49 | 17.175 | 12.066 | 16.655 | 0.538 | 1.03 |
| **SL1344_3971** | *yihW* | 55.808 | 49.662 | 67.136 | 69.68 | 0.831 | 0.713 |
| **SL1344_3972** | *yihX* | 463.196 | 504.378 | 208.48 | 216.441 | 2.22 | 2.33 |
| **SL1344_3973** | *rbn* | 16.138 | 11.999 | 70.041 | 82.452 | 0.23 | 0.146 |
| **SL1344_3974** | *yihZ* | 123.795 | 142.41 | 143.203 | 160.033 | 0.864 | 0.89 |
| **SL1344_3975** | *yiiD* | 389.128 | 49.333 | 62.126 | 65.234 | 6.26 | 0.756 |
| **SL1344_3976** | *SL1344-3976* | 11.591 | 47.409 | 44.113 | 58.672 | 0.263 | 0.808 |
| **SL1344_3978** | *SL1344-3978* | 19.859 | 16.237 | 22.353 | 17.341 | 0.888 | 0.936 |
| **SL1344_3979** | *SL1344-3979* | 5.123 | 52.542 | 50.087 | 72.846 | 0.102 | 0.721 |

| | | | | | | | |
|---|---|---|---|---|---|---|---|
| **SL1344_3980** | *SL1344-3980* | 22.376 | 93.033 | 50.499 | 70.127 | 0.443 | 1.33 |
| **SL1344_3981** | *fdhE* | 201.579 | 130.557 | 448.215 | 367.206 | 0.45 | 0.356 |
| **SL1344_3982** | *fdoI* | 893.035 | 158.075 | 1065.561 | 333.751 | 0.838 | 0.474 |
| **SL1344_3983** | *fdoH* | 1217.778 | 200.071 | 1548.064 | 733.96 | 0.787 | 0.273 |
| **SL1344_3984** | *fdoG* | 762.988 | 285.061 | 1270.568 | 721.41 | 0.601 | 0.395 |
| **SL1344_3986** | *fdhD* | 63.508 | 51.361 | 91.382 | 86.865 | 0.695 | 0.591 |
| **SL1344_3987** | *SL1344-3987* | 0.803 | 3.757 | 15.42 | 17.642 | 0.0521 | 0.213 |
| **SL1344_3990** | *SL1344-3990* | 45.92 | 10.925 | 47.789 | 42.313 | 0.961 | 0.258 |
| **SL1344_3991** | *SL1344-3991* | 22.341 | 24.353 | 97.854 | 81.364 | 0.228 | 0.299 |
| **SL1344_3993** | *SL1344-3993* | 25.708 | 42.979 | 23.208 | 35.416 | 1.11 | 1.21 |
| **SL1344_3994** | *rhaD* | 4.575 | 1.068 | 17.661 | 32.589 | 0.259 | 0.0328 |
| **SL1344_3995** | *rhaA* | 3.054 | 2.807 | 3.584 | 4.605 | 0.852 | 0.61 |
| **SL1344_3997** | *rhaS* | 13.72 | 16.091 | 39.653 | 32.932 | 0.346 | 0.489 |
| **SL1344_3998** | *rhaR* | 10.877 | 34.371 | 42.723 | 69.404 | 0.255 | 0.495 |
| **SL1344_4004** | *sodA* | 7990.858 | 2302.536 | 1812.752 | 866.155 | 4.41 | 2.66 |
| **SL1344_4005** | *yiiM* | 89.478 | 70.683 | 107.421 | 110.487 | 0.833 | 0.64 |
| **SL1344_4006** | *SL1344-4006* | 81.168 | 99.799 | 140.109 | 155.209 | 0.579 | 0.643 |
| **SL1344_4007** | *cpxA* | 47.993 | 56.882 | 169.088 | 167.914 | 0.284 | 0.339 |
| **SL1344_4008** | *cpxR* | 145.234 | 228.197 | 163.365 | 210.657 | 0.889 | 1.08 |
| **SL1344_4009** | *cpxP* | 4.372 | 71.551 | 816.992 | 2045.645 | 0.00535 | 0.035 |
| **SL1344_4010** | *fieF* | 21.239 | 29.001 | 138.842 | 184.139 | 0.153 | 0.157 |
| **SL1344_4011** | *pfkA* | 637.922 | 515.846 | 428.241 | 305.311 | 1.49 | 1.69 |
| **SL1344_4012** | *sbp* | 27.265 | 568.287 | 33.525 | 344.805 | 0.813 | 1.65 |
| **SL1344_4013** | *cdh-a* | 259.954 | 179.501 | 115.49 | 94.828 | 2.25 | 1.89 |

| | | | | | | | |
|---|---|---|---|---|---|---|---|
| **SL1344_4014** | *SL1344-4014* | 12.061 | 13.248 | 15.464 | 14.924 | 0.78 | 0.888 |
| **SL1344_4015** | *SL1344-4015* | 13.504 | 9.991 | 22.028 | 14.81 | 0.613 | 0.675 |
| **SL1344_4016** | *SL1344-4016* | 65.79 | 45.413 | 56.81 | 67.636 | 1.16 | 0.671 |
| **SL1344_4017** | *SL1344-4017* | 23.17 | 37.821 | 74.063 | 92.803 | 0.313 | 0.408 |
| **SL1344_4020** | *SL1344-4020* | 20.175 | 61.397 | 12.082 | 20.466 | 1.67 | 3 |
| **SL1344_4021** | *ydeV* | 5.091 | 14.475 | 29.789 | 31.085 | 0.171 | 0.466 |
| **SL1344_4022** | *ydeW* | 54.816 | 34.932 | 60.408 | 51.733 | 0.907 | 0.675 |
| **SL1344_4023** | *ego* | 42.045 | 31.796 | 39.809 | 48.31 | 1.06 | 0.658 |
| **SL1344_4024** | *ydeY* | 23.871 | 15.246 | 34.057 | 29.691 | 0.701 | 0.513 |
| **SL1344_4025** | *ydeZ* | 19.082 | 14.935 | 19.316 | 17.664 | 0.988 | 0.846 |
| **SL1344_4026** | *yneA* | 20.773 | 25.466 | 17.448 | 19.948 | 1.19 | 1.28 |
| **SL1344_4027** | *yneB* | 135.147 | 141.633 | 35.351 | 35.771 | 3.82 | 3.96 |
| **SL1344_4028** | *yneC* | 26.577 | 51.826 | 29.259 | 30.018 | 0.908 | 1.73 |
| **SL1344_4029** | *SL1344-4029* | 20.737 | 10.225 | 41.698 | 39.35 | 0.497 | 0.26 |
| **SL1344_4030** | *tpiA* | 1993.802 | 1797.603 | 1515.436 | 1320.391 | 1.32 | 1.36 |
| **SL1344_4031** | *yiiQ* | 19.434 | 37.258 | 44.547 | 62.816 | 0.436 | 0.593 |
| **SL1344_4032** | *yiiR* | 26.355 | 8.879 | 64.164 | 45.765 | 0.411 | 0.194 |
| **SL1344_4033** | *fpr* | 42.316 | 48.808 | 136.17 | 135.751 | 0.311 | 0.36 |
| **SL1344_4034** | *glpX* | 166.459 | 125.681 | 304.035 | 374.616 | 0.547 | 0.335 |
| **SL1344_4035** | *glpK* | 8999.387 | 11067.09 | 5523.721 | 5631.696 | 1.63 | 1.97 |
| **SL1344_4036** | *glpF* | 5126.12 | 5412.595 | 4582.75 | 4275.965 | 1.12 | 1.27 |
| **SL1344_4037** | *yiiU* | 4579.424 | 3399.362 | 2522.963 | 2320.361 | 1.82 | 1.47 |
| **SL1344_4038** | *menG* | 279.042 | 388.673 | 1134.547 | 1156.554 | 0.246 | 0.336 |
| **SL1344_4039** | *menA* | 73.261 | 53.32 | 177.59 | 79.868 | 0.413 | 0.668 |

| | | | | | | | |
|---|---|---|---|---|---|---|---|
| **SL1344_4040** | *hslU* | 1591.404 | 267.285 | 956.534 | 209.114 | 1.66 | 1.28 |
| **SL1344_4041** | *hslV* | 1367.602 | 161.919 | 1806.399 | 310.363 | 0.757 | 0.522 |
| **SL1344_4042** | *ftsN* | 31.327 | 44.092 | 121.017 | 111.223 | 0.259 | 0.396 |
| **SL1344_4043** | *cytR* | 356.32 | 202.473 | 376.304 | 222.935 | 0.947 | 0.908 |
| **SL1344_4044** | *priA* | 14.105 | 9.62 | 28.686 | 22.771 | 0.492 | 0.422 |
| **SL1344_4045** | *rpmE* | 4664.961 | 1016.806 | 3284.992 | 2066.325 | 1.42 | 0.492 |
| **SL1344_4047** | *SL1344-4047* | 1.487 | 1.937 | 7.485 | 5.481 | 0.199 | 0.353 |
| **SL1344_4048** | *metJ* | 294.11 | 295.614 | 297.554 | 286.648 | 0.988 | 1.03 |
| **SL1344_4049** | *metB* | 19.069 | 12.186 | 121.016 | 82.158 | 0.158 | 0.148 |
| **SL1344_4050** | *metL* | 46.056 | 25.833 | 84.117 | 74.951 | 0.548 | 0.345 |
| **SL1344_4053** | *SL1344-4053* | 4.367 | 8.586 | 79.903 | 59.039 | 0.0547 | 0.145 |
| **SL1344_4054** | *SL1344-4054* | 6.995 | 3.451 | 10.911 | 10.87 | 0.641 | 0.317 |
| **SL1344_4055** | *metF* | 32.554 | 11.375 | 24.94 | 11.804 | 1.31 | 0.964 |
| **SL1344_4056** | *katG* | 82.97 | 164.424 | 71.191 | 91.416 | 1.17 | 1.8 |
| **SL1344_4058** | *gldA* | 42.466 | 68.144 | 52.854 | 84.911 | 0.803 | 0.803 |
| **SL1344_4059** | *talC* | 82.408 | 122.397 | 106.931 | 115.46 | 0.771 | 1.06 |
| **SL1344_4060** | *ptsA* | 14.29 | 19.547 | 29.797 | 31.678 | 0.48 | 0.617 |
| **SL1344_4061** | *frwC* | 6.084 | 8.125 | 19.447 | 20.933 | 0.313 | 0.388 |
| **SL1344_4062** | *frwB* | 94.6 | 186.476 | 53.819 | 74.495 | 1.76 | 2.5 |
| **SL1344_4063** | *pflD* | 9.84 | 8.939 | 30.882 | 38.625 | 0.319 | 0.231 |
| **SL1344_4064** | *pflC* | 5.589 | 9.828 | 57.657 | 89.811 | 0.0969 | 0.109 |
| **SL1344_4065** | *frwD* | 19.731 | 13.451 | 26.729 | 44.806 | 0.738 | 0.3 |
| **SL1344_4066** | *yijO* | 12.645 | 6.626 | 33.318 | 36.268 | 0.38 | 0.183 |
| **SL1344_4068** | *yijP* | 366.705 | 106.622 | 159.283 | 99.574 | 2.3 | 1.07 |

| | | | | | | | |
|---|---|---|---|---|---|---|---|
| **SL1344_4069** | *ppc* | 183.074 | 121.756 | 119.121 | 104.848 | 1.54 | 1.16 |
| **SL1344_4070** | *argE* | 114.946 | 139.054 | 103.356 | 99.927 | 1.11 | 1.39 |
| **SL1344_4071** | *argC* | 14.313 | 14.619 | 37.732 | 32.816 | 0.379 | 0.446 |
| **SL1344_4072** | *argB* | 32.987 | 31.516 | 90.227 | 76.457 | 0.366 | 0.412 |
| **SL1344_4073** | *argH* | 55.454 | 30.824 | 90.476 | 53.046 | 0.613 | 0.581 |
| **SL1344_4074** | *oxyR* | 108.007 | 178.278 | 190.766 | 305.106 | 0.566 | 0.584 |
| **SL1344_4075** | *udhA* | 275.86 | 566.455 | 383.745 | 482.368 | 0.719 | 1.17 |
| **SL1344_4076** | *yijC* | 355.818 | 288.876 | 399.782 | 377.471 | 0.89 | 0.765 |
| **SL1344_4077** | *yijD* | 273.116 | 343.885 | 169.054 | 232.721 | 1.62 | 1.48 |
| **SL1344_4078** | *trmA* | 35.484 | 18.905 | 70.056 | 37.663 | 0.507 | 0.502 |
| **SL1344_4079** | *btuB* | 448.459 | 166.142 | 190.586 | 163.345 | 2.35 | 1.02 |
| **SL1344_4080** | *murI* | 275.268 | 317.832 | 267.131 | 205.548 | 1.03 | 1.55 |
| **SL1344_4081** | *murB* | 109.185 | 44.158 | 256.152 | 118.45 | 0.426 | 0.373 |
| **SL1344_4082** | *birA* | 51.326 | 37.154 | 62.304 | 60.811 | 0.824 | 0.611 |
| **SL1344_4083** | *coaA* | 83.034 | 123.382 | 169.975 | 134.064 | 0.489 | 0.92 |
| **SL1344_4085** | *tufB* | 31448.244 | 25388.461 | 22788.905 | 15887.346 | 1.38 | 1.6 |
| **SL1344_4086** | *secE* | 950.193 | 752.131 | 3279.337 | 1663.868 | 0.29 | 0.452 |
| **SL1344_4087** | *nusG* | 1966.532 | 885.896 | 1984.033 | 1327.223 | 0.991 | 0.667 |
| **SL1344_4088** | *rplK* | 4433.092 | 1428.288 | 5257.965 | 2313.164 | 0.843 | 0.617 |
| **SL1344_4089** | *rplA* | 5993.073 | 1398.633 | 10721.016 | 3119.755 | 0.559 | 0.448 |
| **SL1344_4090** | *rplJ* | 8424.143 | 6683.02 | 26471.392 | 11488.001 | 0.318 | 0.582 |
| **SL1344_4091** | *rplL* | 47918.002 | 15393.433 | 30437.904 | 16239.451 | 1.57 | 0.948 |
| **SL1344_4092** | *rpoB* | 1306.287 | 555.262 | 998.784 | 616.569 | 1.31 | 0.901 |
| **SL1344_4093** | *rpoC* | 1614.196 | 923.633 | 1382.845 | 1048.018 | 1.17 | 0.881 |

| | | | | | | | |
|---|---|---|---|---|---|---|---|
| **SL1344_4096** | *sseK1* | 9.192 | 37.005 | 7.728 | 12.412 | 1.19 | 2.98 |
| **SL1344_4098** | *thiH* | 46.927 | 15.836 | 20.734 | 17.569 | 2.26 | 0.901 |
| **SL1344_4099** | *thiG* | 6.296 | 9.705 | 12.273 | 16.245 | 0.513 | 0.597 |
| **SL1344_4101** | *thiF* | 7.409 | 9.948 | 27.672 | 22.777 | 0.268 | 0.437 |
| **SL1344_4103** | *thiC* | 5.682 | 6.458 | 12.477 | 22.266 | 0.455 | 0.29 |
| **SL1344_4104** | *rsd* | 196.83 | 1108.093 | 158.466 | 462.018 | 1.24 | 2.4 |
| **SL1344_4105** | *nudC* | 50.017 | 50.181 | 71.128 | 83.939 | 0.703 | 0.598 |
| **SL1344_4106** | *hemE* | 141.566 | 118.478 | 211.685 | 168.389 | 0.669 | 0.704 |
| **SL1344_4107** | *nfi* | 60.163 | 51.724 | 137.18 | 118.987 | 0.439 | 0.435 |
| **SL1344_4108** | *yjaG* | 57.492 | 75.963 | 269.996 | 245.716 | 0.213 | 0.309 |
| **SL1344_4109** | *hupA* | 8817.142 | 22915.44 | 8559.492 | 8470.649 | 1.03 | 2.71 |
| **SL1344_4110** | *yjaH* | 44.332 | 55.752 | 1041.739 | 1150.675 | 0.0426 | 0.0485 |
| **SL1344_4112** | *hydH* | 16.852 | 24.571 | 86.297 | 86.148 | 0.195 | 0.285 |
| **SL1344_4113** | *hydG* | 52.141 | 54.376 | 81.739 | 113.382 | 0.638 | 0.48 |
| **SL1344_4114** | *purD* | 32.901 | 8.595 | 15.448 | 16.475 | 2.13 | 0.522 |
| **SL1344_4115** | *purH* | 25.651 | 17.283 | 36.383 | 22.977 | 0.705 | 0.752 |
| **SL1344_4116** | *yjaB* | 77.034 | 136.508 | 118.982 | 154.181 | 0.647 | 0.885 |
| **SL1344_4117** | *metA* | 15.912 | 6.217 | 12.564 | 11.621 | 1.27 | 0.535 |
| **SL1344_4118** | *aceB* | 22.983 | 27.387 | 15.258 | 21.296 | 1.51 | 1.29 |
| **SL1344_4119** | *aceA* | 33.294 | 36.695 | 16.534 | 25.68 | 2.01 | 1.43 |
| **SL1344_4120** | *aceK* | 0.439 | 2.174 | 7.283 | 11.054 | 0.0603 | 0.197 |
| **SL1344_4121** | *SL1344-4121* | 40.96 | 33.62 | 58.006 | 42.855 | 0.706 | 0.785 |
| **SL1344_4122** | *iclR* | 79.715 | 75.607 | 141.716 | 139.329 | 0.562 | 0.543 |
| **SL1344_4123** | *metH* | 70.667 | 41.415 | 56.214 | 54.028 | 1.26 | 0.767 |

| | | | | | | | |
|---|---|---|---|---|---|---|---|
| **SL1344_4124** | *yjbB* | 30.794 | 33.135 | 70.937 | 73.84 | 0.434 | 0.449 |
| **SL1344_4125** | *pepE* | 141.723 | 173.701 | 167.469 | 174.158 | 0.846 | 0.997 |
| **SL1344_4127** | *SL1344-4127* | 92.519 | 172.824 | 78.719 | 134.954 | 1.18 | 1.28 |
| **SL1344_4128** | *yjbC* | 119.75 | 99.53 | 177.973 | 131.344 | 0.673 | 0.758 |
| **SL1344_4129** | *yjbD* | 12.142 | 66.277 | 103.455 | 96.45 | 0.117 | 0.687 |
| **SL1344_4130** | *SL1344-4130* | 8.986 | 6.499 | 25.493 | 17.48 | 0.352 | 0.372 |
| **SL1344_4134** | *SL1344-4134* | 5.959 | 5.612 | 5.678 | 6.344 | 1.05 | 0.885 |
| **SL1344_4135** | *SL1344-4135* | 3.153 | 2.338 | 7.39 | 9.832 | 0.427 | 0.238 |
| **SL1344_4139** | *SL1344-4139* | 45.505 | 13.314 | 29.868 | 13.355 | 1.52 | 0.997 |
| **SL1344_4140** | *gtrBb* | 30.616 | 7.022 | 49.87 | 25.661 | 0.614 | 0.274 |
| **SL1344_4142** | *SL1344-4142* | 11.38 | 12.325 | 10.113 | 15.852 | 1.13 | 0.778 |
| **SL1344_4146** | *SL1344-4146* | 2.589 | 1.242 | 4.627 | 3.532 | 0.56 | 0.352 |
| **SL1344_4147** | *SL1344-4147* | 10.238 | 36.577 | 11.947 | 10.949 | 0.857 | 3.34 |
| **SL1344_4148** | *SL1344-4148* | 3.834 | 6.608 | 5.052 | 4.744 | 0.759 | 1.39 |
| **SL1344_4155** | *SL1344-4155* | 97.126 | 32.642 | 15.861 | 16.328 | 6.12 | 2 |
| **SL1344_4156** | *lysC* | 72.831 | 40.259 | 53.152 | 55.79 | 1.37 | 0.722 |
| **SL1344_4157** | *pgi* | 390.216 | 293.646 | 221.437 | 142.171 | 1.76 | 2.07 |
| **SL1344_4158** | *yjbE* | 15.59 | 8.678 | 29.204 | 40.995 | 0.534 | 0.212 |
| **SL1344_4161** | *yjbH* | 2.032 | 1.265 | 11.896 | 14.517 | 0.171 | 0.0871 |
| **SL1344_4162** | *yjbA* | 5.329 | 10.261 | 22.325 | 36.533 | 0.239 | 0.281 |
| **SL1344_4163** | *malG* | 47.701 | 70.159 | 27.837 | 66.105 | 1.71 | 1.06 |
| **SL1344_4164** | *malF* | 15.594 | 32.448 | 20.74 | 41.487 | 0.752 | 0.782 |
| **SL1344_4166** | *malE* | 161.157 | 485.899 | 103.045 | 311.665 | 1.56 | 1.56 |
| **SL1344_4167** | *malK* | 67.14 | 99.029 | 100.486 | 145.073 | 0.668 | 0.683 |

| | | | | | | | |
|---|---|---|---|---|---|---|---|
| **SL1344_4168** | *lamB* | 236.865 | 1171.727 | 147.007 | 599.88 | 1.61 | 1.95 |
| **SL1344_4169** | *malM* | 162.301 | 412.87 | 86.989 | 196.986 | 1.87 | 2.1 |
| **SL1344_4170** | *ubiC* | 60.383 | 111.593 | 331.932 | 405.505 | 0.182 | 0.275 |
| **SL1344_4171** | *ubiA* | 397.002 | 353.508 | 263.086 | 270.703 | 1.51 | 1.31 |
| **SL1344_4172** | *plsB* | 239.443 | 118.567 | 202.615 | 164.927 | 1.18 | 0.719 |
| **SL1344_4173** | *dgkA* | 100.582 | 100.096 | 103.737 | 108.248 | 0.97 | 0.925 |
| **SL1344_4174** | *lexA* | 716.748 | 568.743 | 1352.218 | 1106.985 | 0.53 | 0.514 |
| **SL1344_4175** | *dinF* | 9.687 | 7.182 | 88.983 | 58.423 | 0.109 | 0.123 |
| **SL1344_4176** | *yjbJ* | 452.434 | 1729.592 | 254.763 | 660.661 | 1.78 | 2.62 |
| **SL1344_4177** | *zur* | 50.935 | 57.739 | 188.795 | 161.239 | 0.27 | 0.358 |
| **SL1344_4178** | *SL1344-4178* | 4.786 | 13.594 | 23.949 | 31.74 | 0.2 | 0.428 |
| **SL1344_4179** | *yjbN* | 29.627 | 12.665 | 67.307 | 98.918 | 0.44 | 0.128 |
| **SL1344_4180** | *pspG* | 16.144 | 19.582 | 51.771 | 54.463 | 0.312 | 0.36 |
| **SL1344_4181** | *qor* | 46.742 | 52.398 | 43.491 | 44.647 | 1.07 | 1.17 |
| **SL1344_4182** | *dnaB* | 34.279 | 27.381 | 142.861 | 124.736 | 0.24 | 0.22 |
| **SL1344_4183** | *alr-b* | 43.135 | 29.791 | 64.582 | 58.275 | 0.668 | 0.511 |
| **SL1344_4184** | *tyrB* | 66.433 | 57.596 | 129.619 | 86.347 | 0.513 | 0.667 |
| **SL1344_4185** | *aphA* | 265.877 | 524.45 | 123.99 | 233.525 | 2.14 | 2.25 |
| **SL1344_4186** | *yjbQ* | 14.479 | 18.922 | 82.685 | 149.422 | 0.175 | 0.127 |
| **SL1344_4187** | *yjbR* | 47.919 | 149.38 | 50.199 | 82.967 | 0.955 | 1.8 |
| **SL1344_4190** | *uvrA* | 132.967 | 56.299 | 109.453 | 70.509 | 1.21 | 0.798 |
| **SL1344_4192** | *ssb* | 1202.058 | 541.224 | 1345.383 | 822.444 | 0.893 | 0.658 |
| **SL1344_4193** | *siiA* | 24.127 | 23.104 | 64.549 | 21.505 | 0.374 | 1.07 |
| **SL1344_4194** | *siiB* | 72.405 | 34.133 | 79.373 | 31.211 | 0.912 | 1.09 |

| | | | | | | | |
|---|---|---|---|---|---|---|---|
| **SL1344_4195** | *siiC* | 38.701 | 60.961 | 58.27 | 28.996 | 0.664 | 2.1 |
| **SL1344_4196** | *siiD* | 6.114 | 6.972 | 51.828 | 41.774 | 0.118 | 0.167 |
| **SL1344_4197** | *siiE* | 9.578 | 61.81 | 30.246 | 46.698 | 0.317 | 1.32 |
| **SL1344_4198** | *siiF* | 11.026 | 21.917 | 10.994 | 14.353 | 1 | 1.53 |
| **SL1344_4200** | *yjcC* | 6.503 | 5.01 | 7.786 | 8.09 | 0.835 | 0.619 |
| **SL1344_4201** | *soxS* | 424.957 | 752.853 | 2831.944 | 2389.152 | 0.15 | 0.315 |
| **SL1344_4203** | *SL1344-4203* | 0.973 | 12.506 | 15.644 | 38.455 | 0.0622 | 0.325 |
| **SL1344_4204** | *yjcD* | 30.43 | 7.659 | 55.86 | 24.967 | 0.545 | 0.307 |
| **SL1344_4205** | *yjcE* | 65.628 | 40.061 | 93.445 | 76.221 | 0.702 | 0.526 |
| **SL1344_4206** | *SL1344-4206* | 29.597 | 31.33 | 80.402 | 75.232 | 0.368 | 0.416 |
| **SL1344_4209** | *actP* | 31.606 | 92.55 | 34.321 | 107.75 | 0.921 | 0.859 |
| **SL1344_4210** | *yjcH* | 54.684 | 186.789 | 35.728 | 141.235 | 1.53 | 1.32 |
| **SL1344_4211** | *acs* | 378.856 | 1143.306 | 179.337 | 878.818 | 2.11 | 1.3 |
| **SL1344_4213** | *nrfA* | 4.243 | 44.921 | 8.281 | 36.68 | 0.512 | 1.22 |
| **SL1344_4214** | *nrfB* | 2.506 | 20.154 | 23.262 | 31.727 | 0.108 | 0.635 |
| **SL1344_4216** | *nrfD* | 1.484 | 4.194 | 7.191 | 12.658 | 0.206 | 0.331 |
| **SL1344_4219** | *gltP* | 67.006 | 70.927 | 143.422 | 141.477 | 0.467 | 0.501 |
| **SL1344_4220** | *yjcO* | 87.728 | 115.369 | 110.688 | 119.63 | 0.793 | 0.964 |
| **SL1344_4221** | *fdhF* | 3.031 | 5.035 | 20.29 | 31.394 | 0.149 | 0.16 |
| **SL1344_4223** | *lpxO* | 48.905 | 97.205 | 31.938 | 53.757 | 1.53 | 1.81 |
| **SL1344_4225** | *phnB* | 5.866 | 12.256 | 28.092 | 41.659 | 0.209 | 0.294 |
| **SL1344_4226** | *phnA* | 142.877 | 167.219 | 404.073 | 401.253 | 0.354 | 0.417 |
| **SL1344_4227** | *proP* | 34.501 | 75.081 | 90.332 | 152.28 | 0.382 | 0.493 |
| **SL1344_4228** | *basS* | 173.438 | 137.757 | 78.778 | 78.022 | 2.2 | 1.77 |

| | | | | | | | |
|---|---|---|---|---|---|---|---|
| **SL1344_4229** | *basR* | 266.242 | 335.938 | 188.477 | 220.356 | 1.41 | 1.52 |
| **SL1344_4230** | *yjdB* | 101.646 | 73.4 | 74.387 | 86.95 | 1.37 | 0.844 |
| **SL1344_4231** | *adiC* | 31.853 | 28.113 | 25.448 | 17.402 | 1.25 | 1.62 |
| **SL1344_4232** | *adiY* | 12.895 | 27.405 | 11.947 | 22.592 | 1.08 | 1.21 |
| **SL1344_4233** | *adiA* | 6.125 | 10.693 | 11.458 | 12.925 | 0.535 | 0.827 |
| **SL1344_4234** | *melR* | 13.577 | 25.007 | 66.92 | 71.552 | 0.203 | 0.349 |
| **SL1344_4235** | *melA* | 28.724 | 13.193 | 15.859 | 10.306 | 1.81 | 1.28 |
| **SL1344_4237** | *fumB* | 24.332 | 23.995 | 26.593 | 27.541 | 0.915 | 0.871 |
| **SL1344_4238** | *dcuB* | 43.745 | 49.253 | 50.248 | 59.969 | 0.871 | 0.821 |
| **SL1344_4240** | *dcuR* | 35.845 | 95.608 | 67.104 | 94.566 | 0.534 | 1.01 |
| **SL1344_4241** | *dcuS* | 5.404 | 5.919 | 46.383 | 67.499 | 0.117 | 0.0877 |
| **SL1344_4242** | *SL1344-4242* | 4.433 | 16.32 | 3.304 | 13.393 | 1.34 | 1.22 |
| **SL1344_4243** | *SL1344-4243* | 3.493 | 14.646 | 32.24 | 55.376 | 0.108 | 0.264 |
| **SL1344_4244** | *SL1344-4244* | 4.359 | 6.855 | 14.911 | 21.336 | 0.292 | 0.321 |
| **SL1344_4245** | *SL1344-4245* | 2.625 | 9.673 | 14.907 | 28.652 | 0.176 | 0.338 |
| **SL1344_4247** | *SL1344-4247* | 15.746 | 21.998 | 20.73 | 20.763 | 0.76 | 1.06 |
| **SL1344_4250** | *rtsB* | 3.289 | 38.734 | 11.947 | 43.236 | 0.275 | 0.896 |
| **SL1344_4251** | *rtsA* | 94.738 | 236.91 | 25.122 | 80.21 | 3.77 | 2.95 |
| **SL1344_4253** | *SL1344-4253* | 9.463 | 37.712 | 55.104 | 84.05 | 0.172 | 0.449 |
| **SL1344_4254** | *SL1344-4254* | 33.928 | 24.333 | 59.299 | 74.797 | 0.572 | 0.325 |
| **SL1344_4255** | *phoN* | 35.375 | 95.663 | 28.083 | 60.527 | 1.26 | 1.58 |
| **SL1344_4258** | *SL1344-4258* | 10.31 | 32.07 | 37.456 | 43.565 | 0.275 | 0.736 |
| **SL1344_4259** | *yjdC* | 84.886 | 167.337 | 60.732 | 121.818 | 1.4 | 1.37 |
| **SL1344_4260** | *dipZ* | 13.687 | 12.974 | 27.344 | 29.571 | 0.501 | 0.439 |

| | | | | | | | |
|---|---|---|---|---|---|---|---|
| **SL1344_4261** | *cutA* | 9.526 | 45.348 | 36.048 | 48.983 | 0.264 | 0.926 |
| **SL1344_4262** | *dcuA* | 488.058 | 394.074 | 142.651 | 187.469 | 3.42 | 2.1 |
| **SL1344_4263** | *aspA* | 3912.142 | 9231.483 | 2428.362 | 5075.82 | 1.61 | 1.82 |
| **SL1344_4264** | *fxsA* | 219.685 | 116.005 | 523.596 | 131.344 | 0.42 | 0.883 |
| **SL1344_4265** | *yjeH* | 3.432 | 3.888 | 36.419 | 37.573 | 0.0942 | 0.103 |
| **SL1344_4266** | *groES* | 15947.848 | 4996.555 | 17066.055 | 6404.099 | 0.934 | 0.78 |
| **SL1344_4267** | *groEL* | 10938.594 | 5208.19 | 5779.82 | 2924.596 | 1.89 | 1.78 |
| **SL1344_4268** | *yjeI* | 1122.306 | 967.657 | 1231.724 | 1485.893 | 0.911 | 0.651 |
| **SL1344_4269** | *yjeJ* | 13.453 | 11.892 | 12.95 | 12.766 | 1.04 | 0.932 |
| **SL1344_4270** | *yjeK* | 38.312 | 23.996 | 59.91 | 64.171 | 0.639 | 0.374 |
| **SL1344_4271** | *efp* | 2530.324 | 3706.139 | 2772.65 | 1830.81 | 0.913 | 2.02 |
| **SL1344_4273** | *ecnB* | 493.689 | 766.743 | 407.67 | 825.37 | 1.21 | 0.929 |
| **SL1344_4274** | *rflM* | 14.396 | 32.969 | 24.016 | 40.01 | 0.599 | 0.824 |
| **SL1344_4275** | *sugE* | 30.714 | 28.449 | 199.722 | 109.413 | 0.154 | 0.26 |
| **SL1344_4276** | *blc* | 8.868 | 27.769 | 18.793 | 39.891 | 0.472 | 0.696 |
| **SL1344_4277** | *frdD* | 359.77 | 798.683 | 742.522 | 1088.249 | 0.485 | 0.734 |
| **SL1344_4278** | *frdC* | 386.557 | 521.153 | 562.064 | 867.69 | 0.688 | 0.601 |
| **SL1344_4279** | *frdB* | 789.741 | 1385.875 | 553.378 | 963.732 | 1.43 | 1.44 |
| **SL1344_4280** | *frdA* | 682.371 | 753.627 | 507.148 | 868.116 | 1.35 | 0.868 |
| **SL1344_4281** | *epmA* | 181.033 | 112.186 | 236.087 | 97.16 | 0.767 | 1.15 |
| **SL1344_4282** | *yjeM* | 8.31 | 4.707 | 10.159 | 6.226 | 0.818 | 0.756 |
| **SL1344_4284** | *yjeP* | 13.638 | 11.368 | 36.656 | 39.734 | 0.372 | 0.286 |
| **SL1344_4285** | *psd* | 178.805 | 117.861 | 141 | 93.621 | 1.27 | 1.26 |
| **SL1344_4286** | *yjeQ* | 155.763 | 119.748 | 182.969 | 183.979 | 0.851 | 0.651 |

| | | | | | | | |
|---|---|---|---|---|---|---|---|
| **SL1344_4287** | *orn* | 357.118 | 254.021 | 384.544 | 256.048 | 0.929 | 0.992 |
| **SL1344_4288** | *SL1344-4288* | 84.118 | 76.375 | 42.372 | 34.751 | 1.99 | 2.2 |
| **SL1344_4289** | *yjeS* | 10.437 | 6.862 | 61.183 | 44.031 | 0.171 | 0.156 |
| **SL1344_4290** | *yjeF* | 37.054 | 36.647 | 80.482 | 89.218 | 0.46 | 0.411 |
| **SL1344_4291** | *yjeE* | 51.635 | 45.642 | 67.184 | 112.625 | 0.769 | 0.405 |
| **SL1344_4292** | *amiB* | 46.145 | 55.087 | 119.69 | 113.457 | 0.386 | 0.486 |
| **SL1344_4293** | *mutL* | 18.68 | 23.516 | 104.071 | 115.842 | 0.179 | 0.203 |
| **SL1344_4294** | *miaA* | 23.902 | 36.335 | 992.716 | 718.526 | 0.0241 | 0.0506 |
| **SL1344_4295** | *hfq* | 1765.869 | 2823.212 | 1254.579 | 1198.143 | 1.41 | 2.36 |
| **SL1344_4296** | *hflX* | 186.132 | 84.376 | 572.909 | 449.281 | 0.325 | 0.188 |
| **SL1344_4297** | *hflK* | 411.213 | 872.562 | 316.83 | 366.962 | 1.3 | 2.38 |
| **SL1344_4298** | *hflC* | 422.901 | 411.512 | 634.738 | 554.116 | 0.666 | 0.743 |
| **SL1344_4299** | *purA* | 1570.861 | 568.415 | 2696.053 | 868.037 | 0.583 | 0.655 |
| **SL1344_4300** | *nsrR* | 59.889 | 59.079 | 352.192 | 428.862 | 0.17 | 0.138 |
| **SL1344_4301** | *rnr* | 168.288 | 126.979 | 195.506 | 166.932 | 0.861 | 0.761 |
| **SL1344_4302** | *yjfH* | 326.944 | 147.658 | 141.213 | 93.577 | 2.32 | 1.58 |
| **SL1344_4310** | *aidB* | 5.143 | 21.081 | 21.156 | 48.566 | 0.243 | 0.434 |
| **SL1344_4311** | *yjfN* | 384.118 | 1055.707 | 2769.085 | 2204.491 | 0.139 | 0.479 |
| **SL1344_4312** | *yjfO* | 39.072 | 31.653 | 1540.369 | 1238.307 | 0.0254 | 0.0256 |
| **SL1344_4313** | *yjfP* | 33.701 | 68.747 | 35.555 | 64.928 | 0.948 | 1.06 |
| **SL1344_4314** | *yjfQ* | 45.57 | 95.464 | 96.621 | 94.924 | 0.472 | 1.01 |
| **SL1344_4315** | *yjfR* | 4.891 | 8.686 | 6.865 | 9.081 | 0.712 | 0.957 |
| **SL1344_4316** | *sgaT* | 5.166 | 4.476 | 5.076 | 6.538 | 1.02 | 0.685 |
| **SL1344_4317** | *sgaB* | 30.178 | 26.008 | 13.353 | 17.542 | 2.26 | 1.48 |

| | | | | | | | |
|---|---|---|---|---|---|---|---|
| **SL1344_4318** | *ptxA* | 11.839 | 20.187 | 9.095 | 14.091 | 1.3 | 1.43 |
| **SL1344_4319** | *sgaH* | 13.185 | 17.345 | 49.441 | 24.847 | 0.267 | 0.698 |
| **SL1344_4320** | *sgaU* | 3.669 | 4.376 | 14.337 | 22.482 | 0.256 | 0.195 |
| **SL1344_4324** | *rpsF* | 4875.314 | 1504.713 | 3642.829 | 1776.106 | 1.34 | 0.847 |
| **SL1344_4325** | *priB* | 36.644 | 51.394 | 4046.367 | 1830.299 | 0.00906 | 0.0281 |
| **SL1344_4326** | *rpsR* | 3431.975 | 714.522 | 3454.96 | 1000.558 | 0.993 | 0.714 |
| **SL1344_4327** | *rplI* | 1976.697 | 1077.457 | 4498.224 | 1422.413 | 0.439 | 0.757 |
| **SL1344_4329** | *ytfB* | 26.153 | 55.08 | 132.772 | 160.204 | 0.197 | 0.344 |
| **SL1344_4330** | *fklB* | 1413.969 | 761.264 | 883.124 | 426.393 | 1.6 | 1.79 |
| **SL1344_4331** | *cycA* | 229.011 | 477.741 | 297.309 | 618.566 | 0.77 | 0.772 |
| **SL1344_4333** | *ytfF* | 11.827 | 19.294 | 47.047 | 89.807 | 0.251 | 0.215 |
| **SL1344_4334** | *ytfG* | 9.273 | 7.371 | 7.177 | 8.544 | 1.29 | 0.863 |
| **SL1344_4335** | *ytfH* | 7.495 | 12.831 | 59.644 | 32.801 | 0.126 | 0.391 |
| **SL1344_4336** | *cpdB* | 262.599 | 175.232 | 141.228 | 98.481 | 1.86 | 1.78 |
| **SL1344_4337** | *cysQ* | 106.805 | 96.386 | 128.856 | 166.837 | 0.829 | 0.578 |
| **SL1344_4338** | *ytfJ* | 20.156 | 32.792 | 34.943 | 52.041 | 0.577 | 0.63 |
| **SL1344_4339** | *ytfK* | 1058.635 | 3743.141 | 986.601 | 2588.77 | 1.07 | 1.45 |
| **SL1344_4340** | *ytfL* | 283.111 | 104.259 | 121.393 | 84.698 | 2.33 | 1.23 |
| **SL1344_4341** | *msrA* | 700.513 | 222.054 | 517.715 | 131.035 | 1.35 | 1.69 |
| **SL1344_4342** | *ytfM* | 41.579 | 36.992 | 111.783 | 123.446 | 0.372 | 0.3 |
| **SL1344_4343** | *ytfN* | 47.074 | 50.458 | 111.754 | 149.205 | 0.421 | 0.338 |
| **SL1344_4344** | *ytfP* | 482.82 | 621.655 | 379.611 | 418.189 | 1.27 | 1.49 |
| **SL1344_4347** | *ppa* | 1106.522 | 1289.329 | 1651.962 | 1338.214 | 0.67 | 0.963 |
| **SL1344_4348** | *fbp* | 1274.368 | 732.505 | 1015.625 | 815.86 | 1.25 | 0.898 |

| | | | | | | | |
|---|---|---|---|---|---|---|---|
| **SL1344_4349** | *mpl* | 192.425 | 96.265 | 180.092 | 157.197 | 1.07 | 0.612 |
| **SL1344_4350** | *iolR* | 106.607 | 111.819 | 44.179 | 44.183 | 2.41 | 2.53 |
| **SL1344_4351** | *iolT1* | 7.513 | 8.881 | 18.143 | 24.92 | 0.414 | 0.356 |
| **SL1344_4352** | *iolT2* | 5.067 | 11.503 | 14.516 | 16.574 | 0.349 | 0.694 |
| **SL1344_4353** | *iolB* | 17.247 | 41.569 | 44.603 | 50.059 | 0.387 | 0.83 |
| **SL1344_4354** | *iolA* | 14.386 | 10.117 | 18.087 | 22.333 | 0.795 | 0.453 |
| **SL1344_4357** | *iolG1* | 2.928 | 5.853 | 6.594 | 7.797 | 0.444 | 0.751 |
| **SL1344_4361** | *iolC* | 21.9 | 22.428 | 50.119 | 18.515 | 0.437 | 1.21 |
| **SL1344_4362** | *iolD* | 2.914 | 6.837 | 7.573 | 7.316 | 0.385 | 0.935 |
| **SL1344_4364** | *SL1344-4364* | 3.57 | 0.388 | 7.244 | 6.838 | 0.493 | 0.0568 |
| **SL1344_4365** | *iolI2* | 2.684 | 6.419 | 3.426 | 5.334 | 0.783 | 1.2 |
| **SL1344_4366** | *iolH* | 5.886 | 2.075 | 7.776 | 8.579 | 0.757 | 0.242 |
| **SL1344_4367** | *yjgA* | 555.591 | 337.888 | 619.31 | 400.659 | 0.897 | 0.843 |
| **SL1344_4368** | *pmbA* | 262.065 | 84.411 | 169.328 | 131.545 | 1.55 | 0.642 |
| **SL1344_4369** | *cybC* | 1277.65 | 1524.942 | 539.757 | 412.477 | 2.37 | 3.7 |
| **SL1344_4371** | *SL1344-4371* | 44.518 | 58.652 | 68.129 | 67.075 | 0.653 | 0.874 |
| **SL1344_4373** | *SL1344-4373* | 50.586 | 25.563 | 46.036 | 28.909 | 1.1 | 0.884 |
| **SL1344_4374** | *SL1344-4374* | 12.573 | 8.964 | 20.783 | 17.469 | 0.605 | 0.513 |
| **SL1344_4375** | *SL1344-4375* | 15.347 | 17.095 | 17.257 | 17.356 | 0.889 | 0.985 |
| **SL1344_4376** | *selA2* | 48.191 | 17.446 | 30.941 | 28.176 | 1.56 | 0.619 |
| **SL1344_4377** | *SL1344-4377* | 51.046 | 38.463 | 40.727 | 33.38 | 1.25 | 1.15 |
| **SL1344_4378** | *SL1344-4378* | 5.969 | 3.803 | 26.254 | 24.229 | 0.227 | 0.157 |
| **SL1344_4379** | *relB* | 249.445 | 125.126 | 217.706 | 188.058 | 1.15 | 0.665 |
| **SL1344_4381** | *nrdG* | 2.546 | 18.87 | 5.55 | 22.411 | 0.459 | 0.842 |

| | | | | | | | |
|---|---|---|---|---|---|---|---|
| **SL1344_4382** | *nrdD* | 28.034 | 57.018 | 16.086 | 50.223 | 1.74 | 1.14 |
| **SL1344_4384** | *treC* | 10.922 | 14.938 | 12.663 | 14.305 | 0.863 | 1.04 |
| **SL1344_4386** | *treR* | 7.618 | 9.758 | 57.165 | 114.543 | 0.133 | 0.0852 |
| **SL1344_4387** | *mgtA* | 31.815 | 8.437 | 79.04 | 57.533 | 0.403 | 0.147 |
| **SL1344_4387A** | *mgtL* | 3512.199 | 3590.011 | 5292.637 | 1398.342 | 0.664 | 2.57 |
| **SL1344_4389** | *yjgF* | 3133.63 | 2105.847 | 1040.986 | 1584.931 | 3.01 | 1.33 |
| **SL1344_4390** | *pyrI* | 9.738 | 33.422 | 13.809 | 27.267 | 0.705 | 1.23 |
| **SL1344_4391** | *pyrB* | 14.229 | 50.653 | 12.254 | 30.663 | 1.16 | 1.65 |
| **SL1344_4393** | *SL1344-4393* | 30.868 | 5.147 | 9.675 | 6.428 | 3.19 | 0.801 |
| **SL1344_4394** | *SL1344-4394* | 7.378 | 0.436 | 16.696 | 5.253 | 0.442 | 0.083 |
| **SL1344_4395** | *SL1344-4395* | 64.554 | 0.812 | 31.384 | 1.901 | 2.06 | 0.427 |
| **SL1344_4396** | *SL1344-4396* | 74.548 | 0 | 33.422 | 2.566 | 2.23 | 0 |
| **SL1344_4397** | *SL1344-4397* | 94.1 | 0.39 | 28.122 | 1.109 | 3.35 | 0.352 |
| **SL1344_4398** | *yjgK* | 27.441 | 52.405 | 27.218 | 38.323 | 1.01 | 1.37 |
| **SL1344_4399** | *argI* | 8.069 | 7.919 | 7.204 | 8.036 | 1.12 | 0.985 |
| **SL1344_4400** | *yjgD* | 398.604 | 337.823 | 571.406 | 565.1 | 0.698 | 0.598 |
| **SL1344_4401** | *miaE* | 15.581 | 19.494 | 42.94 | 43.986 | 0.363 | 0.443 |
| **SL1344_4402** | *ytgA* | 11.621 | 29.212 | 14.219 | 9.147 | 0.817 | 3.19 |
| **SL1344_4403** | *yjgM* | 58.137 | 49.801 | 45.087 | 46.822 | 1.29 | 1.06 |
| **SL1344_4405** | *valS* | 488.126 | 161.789 | 349.106 | 259.008 | 1.4 | 0.625 |
| **SL1344_4406** | *holC* | 24.131 | 19.303 | 363.1 | 208.461 | 0.0665 | 0.0926 |
| **SL1344_4407** | *pepA* | 338.919 | 341.861 | 535.019 | 395.097 | 0.633 | 0.865 |
| **SL1344_4409** | *yjgP* | 80.515 | 111.605 | 237.352 | 299.347 | 0.339 | 0.373 |
| **SL1344_4410** | *yjgQ* | 105.544 | 27.133 | 200.092 | 145.162 | 0.527 | 0.187 |

| | | | | | | | |
|---|---|---|---|---|---|---|---|
| **SL1344_4411** | *idnR* | 2.015 | 11.03 | 6.889 | 11.714 | 0.292 | 0.942 |
| **SL1344_4414** | *idnD* | 4.703 | 9.162 | 4.098 | 5.553 | 1.15 | 1.65 |
| **SL1344_4415** | *idnK* | 4.125 | 16.141 | 11.07 | 15.391 | 0.373 | 1.05 |
| **SL1344_4416** | *yjgB* | 13.986 | 31.21 | 34.015 | 54.379 | 0.411 | 0.574 |
| **SL1344_4418** | *SL1344-4418* | 37.729 | 32.521 | 36.209 | 41.969 | 1.04 | 0.775 |
| **SL1344_4419** | *SL1344-4419* | 47.654 | 126.149 | 97.678 | 155.477 | 0.488 | 0.811 |
| **SL1344_4420** | *SL1344-4420* | 39.917 | 51.187 | 58.791 | 57.577 | 0.679 | 0.889 |
| **SL1344_4421** | *SL1344-4421* | 67.726 | 43.552 | 43.191 | 52.824 | 1.57 | 0.824 |
| **SL1344_4422** | *SL1344-4422* | 26.645 | 35.5 | 83.718 | 101.48 | 0.318 | 0.35 |
| **SL1344_4423** | *SL1344-4423* | 57.568 | 65.28 | 126.562 | 188.05 | 0.455 | 0.347 |
| **SL1344_4424** | *SL1344-4424* | 77.767 | 71.702 | 41.026 | 43.342 | 1.9 | 1.65 |
| **SL1344_4425** | *SL1344-4425* | 81.965 | 71.215 | 64.893 | 64.417 | 1.26 | 1.11 |
| **SL1344_4426** | *SL1344-4426* | 153.532 | 86.972 | 104.137 | 87.653 | 1.47 | 0.992 |
| **SL1344_4427** | *SL1344-4427* | 81.38 | 155.219 | 253.448 | 204.658 | 0.321 | 0.758 |
| **SL1344_4429** | *yeeN* | 377.181 | 514.75 | 133 | 104.142 | 2.84 | 4.94 |
| **SL1344_4430** | *yjhP* | 25.208 | 35.761 | 32.987 | 22.905 | 0.764 | 1.56 |
| **SL1344_4431** | *SL1344-4431* | 9.177 | 61.166 | 145.312 | 231.479 | 0.0632 | 0.264 |
| **SL1344_4433** | *SL1344-4433* | 317.308 | 583.576 | 212.1 | 213.652 | 1.5 | 2.73 |
| **SL1344_4434** | *SL1344-4434* | 2.012 | 4.177 | 4.402 | 4.931 | 0.457 | 0.847 |
| **SL1344_4435** | *SL1344-4435* | 3.459 | 16.637 | 10.551 | 18.736 | 0.328 | 0.888 |
| **SL1344_4436** | *SL1344-4436* | 12.716 | 10.48 | 15.505 | 21.277 | 0.82 | 0.493 |
| **SL1344_4437** | *uxuR* | 63.171 | 91.574 | 264.136 | 231.573 | 0.239 | 0.395 |
| **SL1344_4438** | *trpS2* | 3.678 | 2.08 | 12.584 | 21.627 | 0.292 | 0.0962 |
| **SL1344_4442** | *yjiE* | 22.011 | 12.347 | 31.307 | 24.897 | 0.703 | 0.496 |

| | | | | | | | |
|---|---|---|---|---|---|---|---|
| **SL1344_4443** | *iadA* | 118.692 | 96.552 | 52.25 | 33.434 | 2.27 | 2.89 |
| **SL1344_4446** | *yjiJ* | 9.141 | 73.073 | 14.579 | 41.992 | 0.627 | 1.74 |
| **SL1344_4447** | *yjiN* | 4.095 | 11.872 | 15.272 | 27.144 | 0.268 | 0.437 |
| **SL1344_4448** | *yjiO* | 6.958 | 17.197 | 25.568 | 27.508 | 0.272 | 0.625 |
| **SL1344_4449** | *SL1344-4449* | 46.117 | 69.339 | 54.318 | 80.502 | 0.849 | 0.861 |
| **SL1344_4450** | *SL1344-4450* | 23.445 | 78.49 | 20.81 | 60.421 | 1.13 | 1.3 |
| **SL1344_4454** | *symE* | 21.509 | 3.268 | 12.701 | 13.433 | 1.69 | 0.243 |
| **SL1344_4455** | *hsdS* | 23.3 | 17.078 | 26.894 | 17.817 | 0.866 | 0.959 |
| **SL1344_4456** | *hsdM* | 149.214 | 56.941 | 126.911 | 70.651 | 1.18 | 0.806 |
| **SL1344_4457** | *hsdR* | 13.66 | 26.065 | 50.342 | 48.337 | 0.271 | 0.539 |
| **SL1344_4458** | *mrr* | 68.51 | 142.659 | 58.209 | 88.444 | 1.18 | 1.61 |
| **SL1344_4459** | *SL1344-4459* | 17.676 | 71.564 | 185.68 | 187.806 | 0.0952 | 0.381 |
| **SL1344_4460** | *SL1344-4460* | 29.695 | 90.695 | 107.407 | 210.507 | 0.276 | 0.431 |
| **SL1344_4461** | *yjiA* | 512.585 | 152.106 | 350.778 | 72.562 | 1.46 | 2.1 |
| **SL1344_4462** | *yjiX* | 269.857 | 9.003 | 514.435 | 105.842 | 0.525 | 0.0851 |
| **SL1344_4463** | *cstAb* | 1072.764 | 43.83 | 1422.941 | 61.588 | 0.754 | 0.712 |
| **SL1344_4464** | *tsr* | 521.566 | 1.883 | 413.927 | 15.916 | 1.26 | 0.118 |
| **SL1344_4465** | *SL1344-4465* | 11.578 | 26.756 | 25.19 | 39.293 | 0.46 | 0.681 |
| **SL1344_4466** | *SL1344-4466* | 0.56 | 10.935 | 66.939 | 22.635 | 0.00836 | 0.483 |
| **SL1344_4467** | *SL1344-4467* | 4.484 | 74.794 | 4.5 | 13.921 | 0.997 | 5.37 |
| **SL1344_4468** | *SL1344-4468* | 0.379 | 9.767 | 3.217 | 11.5 | 0.118 | 0.849 |
| **SL1344_4469** | *SL1344-4469* | 1.414 | 16.335 | 3.083 | 8.088 | 0.459 | 2.02 |
| **SL1344_4470** | *SL1344-4470* | 1.059 | 4.035 | 3.037 | 6.011 | 0.349 | 0.671 |
| **SL1344_4471** | *SL1344-4471* | 7.319 | 13.389 | 11.415 | 12.795 | 0.641 | 1.05 |

| | | | | | | | |
|---|---|---|---|---|---|---|---|
| **SL1344_4472** | *mdoB* | 76.416 | 88.369 | 105.352 | 102.404 | 0.725 | 0.863 |
| **SL1344_4473** | *yjjA* | 18.45 | 18.257 | 61.133 | 56.941 | 0.302 | 0.321 |
| **SL1344_4474** | *dnaC* | 34.089 | 39.08 | 106.166 | 80.793 | 0.321 | 0.484 |
| **SL1344_4475** | *dnaT* | 39.901 | 15.368 | 86.817 | 79.163 | 0.46 | 0.194 |
| **SL1344_4476** | *SL1344-4476* | 3.746 | 12.054 | 24.651 | 18.007 | 0.152 | 0.669 |
| **SL1344_4477** | *yjjP* | 4.089 | 7.384 | 18.628 | 20.6 | 0.22 | 0.358 |
| **SL1344_4480** | *SL1344-4480* | 115.036 | 292.536 | 224.733 | 279.821 | 0.512 | 1.05 |
| **SL1344_4481** | *fhuF* | 7.127 | 1.983 | 12.356 | 4.873 | 0.577 | 0.407 |
| **SL1344_4482** | *SL1344-4482* | 6.725 | 4.152 | 14.067 | 16.369 | 0.478 | 0.254 |
| **SL1344_4484** | *rsmC* | 81.054 | 41.381 | 103.798 | 58.837 | 0.781 | 0.703 |
| **SL1344_4485** | *holD* | 27.975 | 11.647 | 199.012 | 221.639 | 0.141 | 0.0526 |
| **SL1344_4486** | *rimI* | 16.288 | 37.891 | 217.135 | 169.144 | 0.075 | 0.224 |
| **SL1344_4487** | *yjjG* | 50.376 | 54.277 | 221.5 | 116.401 | 0.227 | 0.466 |
| **SL1344_4488** | *prfC* | 318.682 | 108.15 | 208.649 | 142.153 | 1.53 | 0.761 |
| **SL1344_4489** | *osmY* | 680.927 | 1157.242 | 169.234 | 480.603 | 4.02 | 2.41 |
| **SL1344_4490** | *yjjU* | 11.629 | 10.64 | 42.783 | 43.63 | 0.272 | 0.244 |
| **SL1344_4490A** | *ytjA* | 382.206 | 625.208 | 413.287 | 343.429 | 0.925 | 1.82 |
| **SL1344_4491** | *yjjV* | 3.9 | 7.734 | 37.972 | 32.301 | 0.103 | 0.239 |
| **SL1344_4493** | *yjjI* | 12.39 | 21.707 | 20.005 | 22.82 | 0.619 | 0.951 |
| **SL1344_4494** | *deoC* | 557.451 | 69.897 | 1055.042 | 78.475 | 0.528 | 0.891 |
| **SL1344_4495** | *deoA* | 425.099 | 21.388 | 344.114 | 38.69 | 1.24 | 0.553 |
| **SL1344_4496** | *deoB* | 1767.029 | 189.059 | 1649.425 | 264.545 | 1.07 | 0.715 |
| **SL1344_4497** | *deoD* | 1476.817 | 308.458 | 1359.698 | 280.172 | 1.09 | 1.1 |
| **SL1344_4501** | *SL1344-4501* | 8.407 | 0.296 | 4.052 | 6.588 | 2.08 | 0.0449 |

| | | | | | | | |
|---|---|---|---|---|---|---|---|
| **SL1344_4502** | *SL1344-4502* | 5.662 | 8.872 | 12.986 | 7.604 | 0.436 | 1.17 |
| **SL1344_4503** | *lplA* | 79.333 | 35.716 | 52.018 | 54.539 | 1.53 | 0.655 |
| **SL1344_4504** | *smp* | 4.497 | 14.448 | 33.341 | 37.299 | 0.135 | 0.387 |
| **SL1344_4505** | *serB* | 52.719 | 36.432 | 146.4 | 119.447 | 0.36 | 0.305 |
| **SL1344_4506** | *radA* | 52.047 | 27.543 | 121.339 | 80.491 | 0.429 | 0.342 |
| **SL1344_4507** | *nadR* | 106.315 | 104.731 | 118.107 | 107.154 | 0.9 | 0.977 |
| **SL1344_4508** | *yjjK* | 514.402 | 389.083 | 556.407 | 425.333 | 0.925 | 0.915 |
| **SL1344_4509** | *slt* | 74.128 | 63.163 | 57.957 | 75.231 | 1.28 | 0.84 |
| **SL1344_4510** | *trpR* | 47.609 | 104.84 | 141.175 | 174.206 | 0.337 | 0.602 |
| **SL1344_4511** | *yjjX* | 31.662 | 27.551 | 22.783 | 30.88 | 1.39 | 0.892 |
| **SL1344_4512** | *gpmB* | 146.89 | 105.916 | 420.477 | 332.757 | 0.349 | 0.318 |
| **SL1344_4513** | *rob* | 113.422 | 140.265 | 233.178 | 266.745 | 0.486 | 0.526 |
| **SL1344_4514** | *creA* | 234.155 | 136.186 | 296.262 | 179.913 | 0.79 | 0.757 |
| **SL1344_4515** | *creB* | 37.919 | 15.674 | 200.506 | 154.709 | 0.189 | 0.101 |
| **SL1344_4516** | *creC* | 7.092 | 5.267 | 24.128 | 23.154 | 0.294 | 0.227 |
| **SL1344_4517** | *creD* | 2.499 | 2.066 | 11.363 | 15.797 | 0.22 | 0.131 |
| **SL1344_4522** | *SL1344-4522* | 8.998 | 3.987 | 5.645 | 7.705 | 1.59 | 0.517 |
| **SL1344_4523** | *SL1344-4523* | 23.695 | 85.836 | 31.362 | 40.478 | 0.756 | 2.12 |
| **SL1344_4525** | *arcA* | 416.755 | 884.561 | 791.919 | 1373.708 | 0.526 | 0.644 |
| **SL1344_4527** | *lasT* | 33.173 | 52.377 | 24.625 | 42.621 | 1.35 | 1.23 |
| **SL1344_h2859.5** | *STM2880* | 75.524 | 331.767 | 143.093 | 280.975 | 0.528 | 1.18 |
| **SL1344_P1-0001** | *finO* | 31.906 | 42.212 | 75.497 | 113.478 | 0.423 | 0.372 |
| **SL1344_P1-0002** | *traX-P1* | 8.845 | 9.187 | 33.267 | 34.879 | 0.266 | 0.263 |
| **SL1344_P1-0003** | *traI-P1* | 1.79 | 1.268 | 21.727 | 21.223 | 0.0824 | 0.0597 |

| | | | | | | | |
|---|---|---|---|---|---|---|---|
| **SL1344_P1-0007** | *traD* | 1.998 | 1.258 | 8.153 | 7.948 | 0.245 | 0.158 |
| **SL1344_P1-0008** | *traT-P1* | 409.428 | 1011.212 | 373.988 | 588.616 | 1.09 | 1.72 |
| **SL1344_P1-0009** | *traS-P1* | 112.051 | 272.635 | 291.079 | 315.448 | 0.385 | 0.864 |
| **SL1344_P1-0010** | *traG-P1* | 7.297 | 9.253 | 36.286 | 41.908 | 0.201 | 0.221 |
| **SL1344_P1-0011** | *traH-P1* | 3.767 | 3.443 | 10.522 | 7.028 | 0.358 | 0.49 |
| **SL1344_P1-0016** | *traN-P1* | 5.767 | 5.25 | 15.694 | 16.781 | 0.367 | 0.313 |
| **SL1344_P1-0036** | *SL1344-P1-0036* | 53.287 | 83.33 | 37.744 | 39.733 | 1.41 | 2.1 |
| **SL1344_P1-0039** | *SL1344-P1-0039* | 2.222 | 1.123 | 6.853 | 5.639 | 0.324 | 0.199 |
| **SL1344_P1-0042** | *SL1344-P1-0042* | 10.962 | 27.083 | 38.054 | 39.104 | 0.288 | 0.693 |
| **SL1344_P1-0043** | *SL1344-P1-0043* | 9.526 | 15.793 | 5.438 | 4.725 | 1.75 | 3.34 |
| **SL1344_P1-0045** | *SL1344-P1-0045* | 6.337 | 2.765 | 6.411 | 8.223 | 0.988 | 0.336 |
| **SL1344_P1-0052** | *samA* | 40.163 | 17.689 | 125.912 | 53.045 | 0.319 | 0.333 |
| **SL1344_P1-0053** | *samB* | 4.736 | 6.189 | 36.882 | 25.677 | 0.128 | 0.241 |
| **SL1344_P1-0054** | *parB* | 1000.433 | 471.072 | 431.787 | 252.889 | 2.32 | 1.86 |
| **SL1344_P1-0055** | *parA-P1* | 554.444 | 399.216 | 247.386 | 132.362 | 2.24 | 3.02 |
| **SL1344_P1-0056** | *samB2* | 295.029 | 563.089 | 495.165 | 381.24 | 0.596 | 1.48 |
| **SL1344_P1-0058** | *SL1344-P1-0058* | 22.003 | 6.688 | 82.342 | 70.333 | 0.267 | 0.0951 |
| **SL1344_P1-0059** | *tlpA* | 85.556 | 61.256 | 238.11 | 146.46 | 0.359 | 0.418 |
| **SL1344_P1-0060** | *SL1344-P1-0060* | 43.579 | 25.595 | 35.329 | 67.889 | 1.23 | 0.377 |
| **SL1344_P1-0061** | *SL1344-P1-0061* | 169.754 | 624.859 | 230.722 | 358.636 | 0.736 | 1.74 |
| **SL1344_P1-0062** | *rlgA* | 44.98 | 67.591 | 191.949 | 310.811 | 0.234 | 0.217 |
| **SL1344_P1-0063** | *rlgA2* | 22.113 | 32.274 | 98.506 | 142.793 | 0.224 | 0.226 |
| **SL1344_P1-0064** | *SL1344-P1-0064* | 27.289 | 49.259 | 67.631 | 101.641 | 0.404 | 0.485 |
| **SL1344_P1-0068** | *spvB* | 1.5 | 2.221 | 3.148 | 3.186 | 0.476 | 0.697 |

| | | | | | | | |
|---|---|---|---|---|---|---|---|
| **SL1344_P1-0071** | *SL1344-P1-0071* | 23.319 | 26.539 | 79.558 | 86.183 | 0.293 | 0.308 |
| **SL1344_P1-0072** | *SL1344-P1-0072* | 105.909 | 246.892 | 493.208 | 654.619 | 0.215 | 0.377 |
| **SL1344_P1-0075** | *SL1344-P1-0075* | 11.474 | 6.901 | 32.206 | 34.648 | 0.356 | 0.199 |
| **SL1344_P1-0076** | *rsdB* | 77.111 | 31.795 | 49.437 | 57.869 | 1.56 | 0.549 |
| **SL1344_P1-0077** | *SL1344-P1-0077* | 7.144 | 15.898 | 103.268 | 97.04 | 0.0692 | 0.164 |
| **SL1344_P1-0078** | *SL1344-P1-0078* | 82.994 | 129.498 | 269.491 | 305.102 | 0.308 | 0.424 |
| **SL1344_P1-0079** | *ccdB* | 71.962 | 87.361 | 278.535 | 268.575 | 0.258 | 0.325 |
| **SL1344_P1-0080** | *ccdA* | 1520.133 | 1088.03 | 407.844 | 393.375 | 3.73 | 2.77 |
| **SL1344_P1-0081** | *SL1344-P1-0081* | 32.205 | 128.745 | 33.507 | 65.357 | 0.961 | 1.97 |
| **SL1344_P1-0084** | *repA2* | 74.262 | 71.319 | 434.518 | 324.363 | 0.171 | 0.22 |
| **SL1344_P1-0087** | *pefA* | 55.202 | 75.885 | 15.055 | 17.61 | 3.67 | 4.31 |
| **SL1344_P1-0089** | *pefD* | 5.125 | 2.945 | 7.965 | 8.443 | 0.643 | 0.349 |
| **SL1344_P1-0090** | *SL1344-P1-0090* | 22.583 | 19.004 | 24.447 | 18.588 | 0.924 | 1.02 |
| **SL1344_P1-0091** | *SL1344-P1-0091* | 77.034 | 86.813 | 137.885 | 204.968 | 0.559 | 0.424 |
| **SL1344_P1-0092** | *pefI* | 16.119 | 15.329 | 35 | 29.855 | 0.461 | 0.513 |
| **SL1344_P1-0094** | *srgA* | 5.974 | 5.928 | 4.494 | 4.806 | 1.33 | 1.23 |
| **SL1344_P1-0096** | *rcK* | 141.091 | 89.842 | 43.036 | 38.088 | 3.28 | 2.36 |
| **SL1344_P1-0099** | *repA-P1* | 18.523 | 8.946 | 142.148 | 167.333 | 0.13 | 0.0535 |
| **SL1344_P1-0102** | *repC-P1* | 70.504 | 288.123 | 802.215 | 975.114 | 0.0879 | 0.295 |
| **SL1344_P1-0104** | *SL1344-P1-0104* | 5.608 | 14.082 | 54.077 | 58.369 | 0.104 | 0.241 |
| **SL1344_P2-0002** | *repZ* | 12.504 | 7.184 | 175.597 | 155.874 | 0.0712 | 0.0461 |
| **SL1344_P2-0003** | *yacA* | 246.424 | 181.39 | 193.015 | 204.893 | 1.28 | 0.885 |
| **SL1344_P2-0004** | *yacB* | 77.876 | 106.856 | 229.286 | 204.834 | 0.34 | 0.522 |
| **SL1344_P2-0005** | *yacCb* | 29.632 | 50.075 | 111.451 | 142.681 | 0.266 | 0.351 |

| | | | | | | | |
|---|---|---|---|---|---|---|---|
| **SL1344_P2-0006** | *yadA* | 26.177 | 67.114 | 78.693 | 96.662 | 0.333 | 0.694 |
| **SL1344_P2-0007** | *yaeA* | 233.489 | 142.438 | 207.699 | 140.32 | 1.12 | 1.02 |
| **SL1344_P2-0008** | *yaeBb* | 12.935 | 33.507 | 13.009 | 24.985 | 0.994 | 1.34 |
| **SL1344_P2-0009** | *yafA* | 9.088 | 29.682 | 18.392 | 34.241 | 0.494 | 0.867 |
| **SL1344_P2-0010** | *yafB* | 42.212 | 96.11 | 39.586 | 53.57 | 1.07 | 1.79 |
| **SL1344_P2-0011** | *yagA* | 18.369 | 74.283 | 25.704 | 45.288 | 0.715 | 1.64 |
| **SL1344_P2-0012** | *cib* | 149.827 | 99.808 | 50.228 | 51.893 | 2.98 | 1.92 |
| **SL1344_P2-0013** | *imm* | 23.133 | 26.778 | 15.037 | 17.509 | 1.54 | 1.53 |
| **SL1344_P2-0016** | *ybfAb* | 68.211 | 142.349 | 46.84 | 66.505 | 1.46 | 2.14 |
| **SL1344_P2-0017** | *resA* | 9.79 | 20.856 | 23.256 | 33.254 | 0.421 | 0.627 |
| **SL1344_P2-0019** | *parA-P2* | 437.178 | 295.511 | 287.074 | 212.873 | 1.52 | 1.39 |
| **SL1344_P2-0020** | *SL1344-P2-0020* | 676.71 | 556.505 | 333.376 | 197.041 | 2.03 | 2.82 |
| **SL1344_P2-0021** | *stbAb* | 312.85 | 203.924 | 187.333 | 163.498 | 1.67 | 1.25 |
| **SL1344_P2-0022** | *stbBb* | 86.156 | 81.002 | 119.29 | 180.449 | 0.722 | 0.449 |
| **SL1344_P2-0032** | *kicA* | 7.365 | 10.059 | 27.596 | 36.098 | 0.267 | 0.279 |
| **SL1344_P2-0033** | *ycgC* | 6.577 | 20.744 | 8.812 | 34.639 | 0.746 | 0.599 |
| **SL1344_P2-0036** | *ycjA* | 1.687 | 2.839 | 7.989 | 6.866 | 0.211 | 0.413 |
| **SL1344_P2-0037** | *psiB-P2* | 12.111 | 18.139 | 11.865 | 9.783 | 1.02 | 1.85 |
| **SL1344_P2-0038** | *psiA-P2* | 7.644 | 9.601 | 10.225 | 14.086 | 0.748 | 0.682 |
| **SL1344_P2-0039** | *SL1344-P2-0039* | 12.457 | 33.095 | 21.964 | 25.115 | 0.567 | 1.32 |
| **SL1344_P2-0041** | *ard* | 8.948 | 5.141 | 5.974 | 5.998 | 1.5 | 0.857 |
| **SL1344_P2-0044** | *ydgAb* | 229.9 | 788.334 | 432.696 | 413.136 | 0.531 | 1.91 |
| **SL1344_P2-0047A** | *SL1344-P2-0047A* | 83.33 | 212.566 | 55.256 | 439.537 | 1.51 | 0.484 |
| **SL1344_P2-0048** | *nikA* | 63.994 | 99.887 | 182.761 | 356.365 | 0.35 | 0.28 |

| | | | | | | | |
|---|---|---|---|---|---|---|---|
| **SL1344_P2-0049** | *nikB* | 7.081 | 10.808 | 63.533 | 94.136 | 0.111 | 0.115 |
| **SL1344_P2-0050** | *trbC-P2* | 3.693 | 3.532 | 11.447 | 12.258 | 0.323 | 0.288 |
| **SL1344_P2-0053** | *pndC* | 1.454 | 23.867 | 8541.161 | 7345.782 | 0.00017 | 0.00325 |
| **SL1344_P2-0054** | *pndA* | 6.278 | 37.618 | 12136.247 | 8281.981 | 0.000517 | 0.00454 |
| **SL1344_P2-0054A** | *SL1344-P2-0054A* | 9.567 | 71.686 | 22343.795 | 24850.216 | 0.000428 | 0.00288 |
| **SL1344_P2-0055** | *exc* | 27.945 | 176.568 | 75.792 | 191.826 | 0.369 | 0.92 |
| **SL1344_P2-0057** | *traX-P2* | 5.363 | 3.605 | 13.601 | 7.894 | 0.394 | 0.457 |
| **SL1344_P2-0058** | *traW-P2* | 4.035 | 2.94 | 7.15 | 9.507 | 0.564 | 0.309 |
| **SL1344_P2-0065** | *traP-P2* | 2.519 | 5.017 | 12.1 | 14.473 | 0.208 | 0.347 |
| **SL1344_P2-0071** | *sogL* | 1.1 | 0.74 | 4.794 | 3.555 | 0.229 | 0.208 |
| **SL1344_P2-0072** | *nuc* | 4.504 | 9.119 | 14.674 | 8.366 | 0.307 | 1.09 |
| **SL1344_P2-0078** | *traF-P2* | 177.829 | 246.81 | 48.564 | 54.005 | 3.66 | 4.57 |
| **SL1344_P2-0079** | *traE-P2* | 8.18 | 22.179 | 28.326 | 20.31 | 0.289 | 1.09 |
| **SL1344_P2-0080** | *rci* | 3.793 | 1.708 | 11.854 | 8.457 | 0.32 | 0.202 |
| **SL1344_P2-0088** | *pilS* | 7.893 | 3.982 | 9.791 | 17.181 | 0.806 | 0.232 |
| **SL1344_P2-0090** | *pilQ* | 2.59 | 2.67 | 12.132 | 6.239 | 0.214 | 0.428 |
| **SL1344_P2-0093** | *pilN* | 0.668 | 2.829 | 3.493 | 3.606 | 0.191 | 0.785 |
| **SL1344_P2-0100** | *trcD* | 37.793 | 84.577 | 25.538 | 34.669 | 1.48 | 2.44 |
| **SL1344_P3-0001** | *sulII* | 1238.004 | 993.727 | 1700.552 | 1587.061 | 0.728 | 0.626 |
| **SL1344_P3-0003** | *repC-P3* | 89.663 | 51.917 | 146.576 | 159.768 | 0.612 | 0.325 |
| **SL1344_P3-0004** | *repA-P3* | 110.989 | 99.522 | 117.083 | 188.451 | 0.948 | 0.528 |
| **SL1344_P3-0005** | *SL1344-P3-0005* | 577.36 | 560.6 | 909.377 | 1131.499 | 0.635 | 0.495 |
| **SL1344_P3-0006** | *SL1344-P3-0006* | 182.308 | 311.046 | 449.284 | 597.319 | 0.406 | 0.521 |
| **SL1344_P3-0008** | *mobAb* | 14.173 | 41.515 | 82.924 | 179.332 | 0.171 | 0.231 |

| | | | | | | | |
|---|---|---|---|---|---|---|---|
| **SL1344_P3-0009** | *mobBb* | 39.749 | 480.092 | 100.426 | 150.621 | 0.396 | 3.19 |
| **SL1344_P3-0011** | *mobC* | 386.113 | 333.423 | 286.483 | 429.959 | 1.35 | 0.775 |
| **SL1344_P3-0013** | *strB* | 94.739 | 133.47 | 514.325 | 584.293 | 0.184 | 0.228 |
| **SL1344_P3-0014** | *aph3''-Ib* | 574.421 | 638.165 | 1182.066 | 1277.147 | 0.486 | 0.5 |

#### Table S2: *fliF* ribosome-binding site strengths

| *fliF -luxA* sequence variant | ORF | RBS score |
|---|---|---|
| WT | tesORF | 895.811 |
| WT | *fliF* | 535.285 |
| tesORF-mut (UUG→UAAG) | *fliF* | 535.285 |
| SD swap | tesORF | 2310.505 |
| SD swap | *fliF* | 449.521 |
| 1×scr | tesORF | 653.732 |
| 1×scr | *fliF* | 1258.764 |
| 4×scr | tesORF | 597.461 |
| 4×scr | *fliF* | 3096.338 |
| sub. weak 2° str. | tesORF | 895.811 |
| sub. weak 2° str. | *fliF* | 3543.921 |
| sub. weak 2° str. tesORF-mut (UUG→UUAG) | *fliF* | 3543.921 |
| sub. weak 2° str. alt. | tesORF | 3158.511 |
| sub. weak 2° str. alt. | *fliF* | 2705.284 |
| sub. weak 2° str. tesORF-mut (UUG→UAAG) alt. | *fliF* | 2705.284 |
| sup. A | *fliF* | 467.681 |
| sup. B | *fliF* | 1258.764 |
| sup. C | *fliF* | 1576.414 |
| sup. D | *fliF* | 3877.702 |
| 3-codon-stop | tesORF | 364.177 |
| 3-codon-stop | *fliF* | 535.285 |
| 6-codon-stop | tesORF | 1404.975 |
| 6-codon-stop | *fliF* | 535.285 |
| 9-codon-stop | tesORF | 895.811 |
| 9-codon-stop | *fliF* | 535.285 |
| 12-codon-stop | tesORF | 895.811 |
| 12-codon-stop | *fliF* | 1576.414 |
| WT short | tesORF | 244.789 |
| WT short | *fliF* | 535.285 |
| tesORF-mut short | *fliF* | 535.285 |
| post-SD ins. short | tesORF | 244.789 |
| post-SD ins. short | *fliF* | 291.296 |
| post-SD del. short | tesORF | 244.789 |
| post-SD del. short | *fliF* | 218.79 |
| UGA-to-UUG short | tesORF | 244.789 |
| UGA-to-UUG short | *fliF* | 373.442 |
| AUG-to-GUG short | tesORF | 244.789 |
| AUG-to-GUG short | *fliF* | 323.471 |
| 3-codon-stop short | tesORF | 1080.896 |
| 3-codon-stop short | *fliF* | 535.285 |
| 6-codon-stop short | tesORF | 244.79 |
| 6-codon-stop short | *fliF* | 535.285 |
| 9-codon-stop short | tesORF | 256.057 |
| 9-codon-stop short | *fliF* | 535.285 |
| 12-codon-stop short | tesORF | 244.79 |
| 12-codon-stop short | *fliF* | 1576.414 |
| alt. str. A | tesORF | 571.169 |
| alt. str. A | *fliF* | 88.466 |
| alt. str. B | tesORF | 254.069 |
| alt. str. B | *fliF* | 489.21 |
| tesORF UUG→AUG | tesORF | 3698.183 |
| tesORF UUG→AUG | *fliF* | 535.285 |

#### Table S3: Distances between the *fliF* and *fliE* CDSs in motile Gammaproteobacteria

| species | order | *fliE* –*fliF* distance | accession number | *fliE* coordinates | *fliF* coordinates |
|---|---|---|---|---|---|
| *Vibrio parahaemolyticus* | Vibrionales | 15 | NC_004603.1 | complement(2363687..2363998) | complement(2361930..2363672) |
| *Photobacterium swingsii* | Vibrionales | 16 | NZ_AP024852.1 | 908400..908711 | 908727..910478 |
| *Aeromonas hydrophila* | Aeromonadales | 58 | NC_008570.1 | 1498693..1499010 | 1499068..1500780 |
| *Anaerobiospirillum succiniciproducens* | Aeromonadales | 23 | NZ_AXWV01000062.1 | 8931..9437 | 9460..11409 |
| *Psychromonas ingrahamii* | Alteromonadales | 13 | NC_008709.1 | complement(4384989..4385312) | complement(4383228..4384976) |
| *Shewanella oneidensis* | Alteromonadales | 46 | NC_004347.2 | complement(3368310..3368648) | complement(3366576..3368264) |
| *Pseudoalteromonas lipolytica* | Alteromonadales | 21 | NZ_CP032090.1 | 1050841..1051173 | 1051194..1052909 |
| *Pseudomonas aeruginosa* | Pseudomonadales | 23 | NC_002516.2 | 1192053..1192382 | 1192405..1194201 |
| *Azomonas agilis* | Pseudomonadales | 14 | NZ_VLKG01000006.1 | complement(104718..105056) | complement(102926..104704) |
| *Marinobacter nauticus* | Pseudomonadales | 16 | NC_017067.1 | 1357256..1357624 | 1357640..1359346 |
| *Marinomonas mediterranea* | Oceanospirillales | 32 | NC_015276.1 | complement(3557570..3557935) | complement(3555862..3557538) |
| *Saccharophagus degradans* | Cellvibrionales | 113 | NC_007912.1 | complement(2785603..2785968) | complement(2783820..2785490) |
| *Legionella pneumophila* | Legionellales | 172 | NZ_CP013742.1 | complement(1976994..1977308) | complement(1975188..1976822) |
| *Allochromatium vinosum* | Chromatiales | 141 | NC_013851.1 | complement(3321317..3321652) | complement(3319557..3321176) |
| *Marichromatium purpuratum* | Chromatiales | 38 | NZ_CP007031.1 | 1511629..1511964 | 1512002..1513675 |
| *Xanthomonas campestris* | Lysobacterales | 14 | NZ_CP058243.1 | 2645350..2645721 | 2645735..2647462 |
| *Dyella jiangningensis* | Lysobacterales | 31 | NZ_CP007444.1 | complement(3263249..3263572) | complement(3261509..3263218) |

#### Table S4: Strains and plasmids

| **strain** | **description** | **reference** |
|---|---|---|
| *Salmonella enterica* subsp. *enterica* serovar Typhimurium SL1344 | hisG46, derived from *Salmonella* strain 4/74 | Hoiseth and Stocker, 1981 |
| Salmonella SL1344 fliF tesORF− | SL1344, *fliF* -tesORF start codon mutant, UUG to UAAG | this study |
| *Salmonella* SL1344 Δ*flhDC* | SL1344, *flhDC* deletion, kanamycin-resistant | this study |
| *Escherichia coli* K-12 MG1655 | F− lambda− *ilvG* − *rfb* -50 *rph* −1 | Blattner *et al.* , 1997 |
| Escherichia coli MG1655 fliF-tesORF− | MG1655, *fliF* -tesORF start codon mutant, UUG to UAAG | this study |
| *Yersinia pseudotuberculosis* YPIII | serotype III strain | Gemski *et al.* , 1980 |
| Yersinia pseudotuberculosis YPIII fliF-tesORF− | YPIII, *fliF* -tesORF start codon mutant, UUG to UAA | this study |
| *Yersinia pseudotuberculosis* YPIII Δfis | YPIII, *fis* deletion mutant | Javadi et al., 2026 |
| *Salmonella* SL1344 suppressor mutant A | SL1344, *fliF* -tesORF start codon mutant, UUG to UAAG, *fliF* −21C>G | this study |
| *Salmonella* SL1344 suppressor mutant B | SL1344, *fliF* -tesORF start codon mutant, UUG to UAAG, *fliF* −29G>T | this study |
| *Salmonella* SL1344 suppressor mutant C | SL1344, *fliF* -tesORF start codon mutant, UUG to UAAG, *fliF* 11C>G | this study |
| *Salmonella* SL1344 suppressor mutant D | SL1344, *fliF* -tesORF start codon mutant, UUG to UAAG, *fliF* 13G>A | this study |
| *Salmonella* SL1344 *fliF* 13G>A | SL1344, 13G>A substitution in *fliF* CDS, A5T substitution in FliF polypeptide (suppressor mutation D) | this study |
| *Salmonella* SL1344 Δ*fliA* | SL1344, *fliA* (σ28) deletion | this study |
| *E. coli* S17-1 λ-*pir* | RP4-2 Tc::Mu-Km::Tn7 (λ*pir*) | Simon *et al.* , 1983 |

| **plasmid** | **description** | **reference** |
|---|---|---|
| *fliF* -tesORF wild-type reporter | a pWKS30-based vector carrying *fliF* tesORF promoter up to and including first ten codons of *fliF* fused to the luxAB operon | this study |
| *fliF* -tesORF start codon mutant reporter | *fliF* -tesORF reporter construct with mutant tesORF start codon, UUG to UAAG | this study |
| Promoter-less reporter construct | the pWKS30-based *luxAB* reporter plasmid without a promoter | Bryant *et al.* , 2023 |
| *fliF* -tesORF 1×scr reporter | *fliF* -tesORF reporter construct containing scrambled tesORF coding region | this study |
| *fliF* -tesORF 4×scr reporter | *fliF* -tesORF reporter construct containing the scrambled tesORF coding region extended to four times of its original length | this study |
| *fliF* -tesORF SD-swap reporter | *fliF* -tesORF reporter construct with the *fliF* -tesORF and *fliF* gene Shine–Dalgarno sequences swapped | this study |
| *fliF* -tesORF sub. weak 2° str. reporter | *fliF* -tesORF reporter construct with eight point mutations that disrupt the experimentally determined stem–loop structre | this study |
| *fliF* -tesORF start codon mutant sub. weak 2° str. reporter | *fliF* -tesORF sub. weak 2° str. reporter construct with tesORF start codon mutation, UUG to UUAG | this study |
| *fliF* -tesORF sub. weak 2° str. alt. reporter | alternative *fliF* -tesORF reporter construct with four point mutations that increase (weaken) the predicted secondary-structure minimal free energy | this study |
| *fliF* -tesORF start codon mutant sub. weak 2° str. alt. reporter | the alternative *fliF* -tesORF sub. weak 2° str.reporter construct with tesORF start codon mutation, UUG to UAAG | this study |
| *fliF* -tesORF 3 codon stop reporter | *fliF* -tesORF reporter construct with a stop codon introduced after the third codon of *fliF* -tesORF | this study |

| | | |
|---|---|---|
| *fliF* -tesORF 6 codon stop reporter | *fliF* -tesORF reporter construct with a stop codon introduced after the sixth codon of *fliF* -tesORF | this study |
| *fliF* -tesORF 9 codon stop reporter | *fliF* -tesORF reporter construct with a stop codon introduced after the ninth codon of *fliF* -tesORF | this study |
| *fliF* -tesORF 12 codon stop reporter | *fliF* -tesORF reporter construct with a stop codon introduced after the twelfth codon of *fliF* -tesORF | this study |
| *fliF* -tesORF wild-type short reporter | *fliF* -tesORF reporter construct with a *tetA* 5′ UTR in place of *fliF* 5′ UTR | this study |
| *fliF* -tesORF 3 codon stop short reporter | *fliF* -tesORF reporter construct with a *tetA* 5′ UTR and a stop codon introduced after the third codon of *fliF* -tesORF | this study |
| *fliF* -tesORF 6 codon stop short reporter | *fliF* -tesORF reporter construct with a *tetA* 5′ UTR and a stop codon introduced after the sixth codon of *fliF* -tesORF | this study |
| *fliF* -tesORF 9 codon stop short reporter | *fliF* -tesORF reporter construct with a *tetA* 5′ UTR and a stop codon introduced after the ninth codon of *fliF* -tesORF | this study |
| *fliF* -tesORF 12 codon stop short reporter | *fliF* -tesORF reporter construct with a *tetA* 5′ UTR and a stop codon introduced after the twelfth codon of *fliF* -tesORF | this study |
| *fliF-* tesORF alt. str. A reporter | *fliF* -tesORF reporter construct tesORF mutations that create an alternative predicted secondary structure | this study |
| *fliF-* tesORF alt. str. B reporter | *fliF* -tesORF reporter construct tesORF mutations that create an alternative predicted secondary structure | this study |
| *fliF* -tesORF sup. A reporter | *fliF* -tesORF start codon mutant reporter construct containing suppressor mutation A, *fliF* −21C>G | this study |
| *fliF* -tesORF sup. B reporter | *fliF* -tesORF start codon mutant reporter construct containing suppressor mutation B, *fliF* −29G>T | this study |
| *fliF* -tesORF sup. C reporter | *fliF* -tesORF start codon mutant reporter construct containing suppressor mutation C, *fliF* 11C>G | this study |
| *fliF* -tesORF sup. D reporter | *fliF* -tesORF start codon mutant reporter construct containing suppressor mutation D, *fliF* 13G>A | this study |
| *fliF* -tesORF stop to UUG | *fliF* -tesORF reporter construct with a *tetA* 5′ UTR and a tesORF stop codon mutated to UUG | this study |
| *fliF* -tesORF AUG to GUG | *fliF* -tesORF reporter construct with a *tetA* 5′ UTR and a *fliF* start codon mutated to GUG | this study |
| *fliF* -tesORF post-SD ins. | *fliF* -tesORF reporter construct with a *tetA* 5′ UTR and 3 nt inserted between the *fliF* Shine–Dalgarno sequence and the start codon | this study |
| *fliF* -tesORF post-SD del. | *fliF* -tesORF reporter construct with a *tetA* 5′ UTR and a 3-nt deletion between the *fliF* Shine–Dalgarno sequence and the start codon | this study |
| *fliF* -tesORF-Myc luxA_FLAG | a pWKS30-based vector carrying *fliF* tesORF promoter up to and including first ten codons of *fliF* fused to the *luxAB* operon. tesORF was modified to encode an N-terminal Myc tag and *luxA* modified to encode a C-terminal FLAG tag. | this study |
| pTrc99a vector | pTrc99a vector used as empty vector control (VC) | Amann *et al.* , 1988 |
| pTrc99a *fliF* -tesORF (untagged) | pTrc99a vector encoding the *fliF* -tesORF peptide | this study |
| pTrc99a *fliF* -tesORF-Myc | pTrc99a vector encoding the *fliF* -tesORF peptide with a C-terminal Myc tag | this study |
| pUC18 | cloning vector, ampicillin resistance | Norrander *et al* ., 1983 |
| pDM4 | *sacBR* , *oriT* , *oriR6K* , CHL resistance | Milton *et al.* , 1996 |

| | | |
|---|---|---|
| pBO8349 | pUC18, 86 bp region upstream of *fliF* + 30 bp of *fliF* CDS, T7 promoter sequence, *Eco* RV restriction site, runoff transcription plasmid for structure probing | this study |
| pBO8751 | pDM4, 2040 bp of the genomic region around the *fliF* -tesORF. Within the 5′-UTR, point mutations TG64,65AA were introduced. Used for *Yersinia* mutant strain generation via conjugation. | this study |

**Table S5: Inferred ribosome P-site distance from RPF read 5′ end**

***Salmonella***

| | | | | | | | | | | | | |
|---|---|---|---|---|---|---|---|---|---|---|---|---|
| mapped Ribo-Seq read length | | 24 | 25 | | 27 | 28 | 29 | 30 | 31 | 32 | 33 | 34 |
| P site distance from the 5′ end of the read | | 8 | 9 | | 11 | 12 | 13 | 14 | 15 | 16 | 17 | 18 |

***E. coli***

| | | | | | | | | | | | | |
|---|---|---|---|---|---|---|---|---|---|---|---|---|
| mapped Ribo-Seq read length | 23 | 24 | 25 | 26 | 27 | 28 | 29 | 30 | 31 | 32 | 33 | 34 |
| P site distance from the 5′ end of the read | 7 | 8 | 9 | 10 | 11 | 12 | 13 | 14 | 15 | 16 | 17 | 18 |

**Table S6: GenBank accession numbers**

| strain | accesion number |
|---|---|
| *Salmonella enterica* subsp. *enterica* serovar Typhimurium SL1344 | FQ312003.1 |
| *Salmonella enterica* subsp. *enterica* serovar Typhimurium LT2 | AE006468.2 |
| *Salmonella enterica* subsp. *enterica* serovar Typhi CT18 | AL513382.1 |
| *Salmonella bongori* N268-08 | CP006608.1 |
| *Citrobacter freundii* FDAARGOS_549 | CP033744.1 |
| *Escherichia fergusonii* RHB19-C05 | CP057657.1 |
| *Escherichia coli* strain K-12 substr. MG1655 | U00096.3 |
| *Atlantibacter hermannii* ATCC 33651 | CP042941.1 |
| *Buttiauxella agrestis* DSM9389 | AP023184.1 |
| *Pluralibacter gergoviae* FDAARGOS_186 | CP020388.1 |
| *Scandinavium goeteborgense* CCUG 66741 | CP054058.1 |
| *Kluyvera ascorbata* TP1631 | AP022665.1 |
| *Cedecea lapagei* NCTC11466 | LR134201.1 |
| *Phytobacter ursingii* CAV1151 | CP011602.1 |
| *Leclercia adecarboxylata* USDA-ARS-USMARC-60222 | CP013990.1 |
| *Cronobacter sakazakii* CS-931 | CP027107.2 |
| *Lelliottia amnigena* FDAARGOS_1445 | CP079896.1 |
| *Yokenella regensburgei* W13 | CP050811.1 |
| *Enterobacter cloacae* GGT036 | CP009756.1 |
| *Kosakonia cowanii* FBS 223 | CP035129.1 |
| *Serratia marcescens* KS10 | CP027798.1 |
| *Yersinia enterocolitica* 8081 | AM286415.1 |
| *Yersinia pseudotuberculosis* YPIII | NZ_CP009792.1 |
| *Proteus mirabilis* HI4320 | AM942759.1 |
